# Evolution of thyroglobulin: an integrated view of its origin and complexity from a structural perspective

**DOI:** 10.1101/2025.06.18.660348

**Authors:** Mauricio Gomes Pio, Wanderson Marques da Silva, Carina M. Rivolta, Héctor M. Targovnik

## Abstract

This study presents a comprehensive bioinformatics analysis of the origin and structural complexity of thyroglobulin (TG). We examine the structural and evolutionary conservation of TG in *Petromyzon marinus* (sea lamprey) by reconstructing its complete TG sequence. Based on genomic data, we assembled a 2,831-amino-acid sequence (TGPM), identifying key TG domains and generating a homology-based PDB model. Additionally, we detected a second TG transcript in sea lamprey, designated TGPM^1746^

Comparative analysis across 38 representative vertebrate species—including mammals, birds, reptiles, amphibians, ray-finned fishes, and jawless vertebrates—reveals that all TG domains are conserved throughout vertebrate evolution. Despite substantial divergence in overall amino acid sequences, tyrosine residues and cysteines—both essential for TG function—remain highly conserved. TG emerges as a structurally complex, multidomain protein featuring a conserved disordered segment at its C-terminus. This region includes the terminal portion of the ChEL domain and the hormonogenic site responsible for triiodothyronine (T_3_) synthesis, likely contributing to the conformational flexibility required for hormone production.

We further propose an evolutionary model in which a nidogen-like precursor—defined by the presence of TG type 1 modules—may have acted as the ancestral source of this essential repetitive motif within TG structure. Through genetic rearrangements and duplication events, a proto-TG likely arose, potentially shaped by environmental pressures such as ionizing radiation. Successive duplications expanded the TG architecture, culminating in the formation of 11 TG type 1 modules. The final evolutionary stage involved the integration of TG type 3 and TG type 2 modules, followed by the fusion of the ChEL domain, which enhanced TG secretion and thyroid hormone biosynthesis. Our findings demonstrate that the TG complexation process is fully established in lampreys and has remained remarkably conserved across vertebrate evolution.

## Introduction

During the evolution of chordates—including the subphyla Cephalochordata, Urochordata (or Tunicata), and Vertebrata (or Craniata)—the formation of thyroid hormones (TH) preceded the morphological differentiation of thyroid cells and their organization into follicles [Suzuki & Kondo, 1971; Suzuki et al., 1975]. The thyroid gland functions as a key endocrine organ responsible for synthesizing and releasing TH, including triiodothyronine (T_3_) and thyroxine (T_4_) [Takagi et al., 2022]. It is characterized by spheroidal follicles formed by a single layer of epithelial cells surrounding a central lumen filled with protein-rich colloid, primarily composed of thyroglobulin (TG). [Eales, 1997]. The thyroid follicle represents a functional unit and an evolutionary innovation among vertebrates. TH biosynthesis occurs at the cell–colloid interface of thyroid follicles and depends on iodide, thyroid peroxidase (TPO), a supply of hydrogen peroxide (DUOX system), an iodine acceptor protein (TG), proteolytic enzymes, and iodide recycling via iodotyrosine deiodinase (IYD) [Targovnik et al., 2020]. Iodine binds to tyrosine residues within TG, forming iodotyrosine—versatile, highly reactive, and mobile intermediates that subsequently undergo coupling reactions to generate TH [Mourouzis et al., 2020]. This process is stimulated by thyroid-stimulating hormone (TSH) binding to its G-protein-coupled receptor (TSHR) [Vassart & Dumont, 1992]. TH played a pivotal role in the evolution of both invertebrate and vertebrate animals by modulating gene expression across diverse tissues and orchestrating complex biological processes. These include the metamorphosis from larva to juveniles in Senegalese sole, amphibians, urochordatas amphioxus or lamprey, in epimorphic regeneration processes after amputation (e.g. fins in fish), smoltification of salmon, cell differentiation, energy metabolism, seasonal physiological changes, and the development of the central nervous system [Belkadi et al., 2012; Mourouzis et al., 2020; Mantzouratou et al., 2022; Takagi et al., 2022; Zwahlen et al., 2024]. Iodine, an essential component in TH signaling likely served as a potent antioxidant for early unicellular organisms 3.5 billion years ago [Mourouzis et al., 2020; Mantzouratou et al., 2022].

The emergence of the follicular thyroid cell and an endocrine system capable of concentrating iodide from aquatic environments enabled efficient TH synthesis—an essential adaptation in freshwater habitats where iodine is scarce [Takagi et al., 2022]. This advancement supported major physiological innovations such as brain development, skeletal formation, and thermoregulation [Mullur et al., 2014], ultimately driving the evolutionary diversification and increasing complexity of vertebrates through the acquisition of unique traits.

Cephalochordates, urochordates, and vertebrates share a common ancestor that likely lived around 550 million years ago [Putnam et al., 2008]. Vertebrates are classified into gnathostomes and cyclostomes (also known as agnathans or jawless fish), which diverged early in vertebrate evolution before the emergence of hinged jaws, a defining trait of gnathostomes [Schimeld et al., 2012]. The only surviving lineages of cyclostomes today fall into two categories: lampreys (also known as Hyperoartia) and hagfishes (also known as Myxini) [Xu et al., 2016]. Cyclostomes are believed to have first appeared during the Ordovician, reached their peak in the Silurian and Devonian, and then gradually declined toward extinction in the late Devonian [Xu et al., 2016]. Recently, using fossil calibrations, Marlétaz et al. [2024] estimated that the divergence between lampreys and hagfishes occurred in the late Ordovician, approximately 449 million years ago, while the split between cyclostomes and gnathostomes took place near the Cambrian-Ordovician boundary, around 493 million years ago.

The oldest known lamprey fossils date back to the late Devonian period, approximately 360 million years ago, and their morphology has remained virtually unchanged over hundreds of millions of years of evolution [Xu et al., 2016]. The most extensively studied modern lamprey species include *Petromyzon marinus* (sea lamprey), *Lethenteron japonicum*, *Lampetra fluviatilis*, and *Lampetra planeri*. Lampreys produce TG before larval metamorphosis, meaning before the thyroid follicles observed in adults are fully organized [Monaco et al., 1978]. In contrast, hagfishes contain iodinated glycoproteins of a non-TG nature, distinguished by differences in molecular size, solubility, amino acid composition, and carbohydrate and iodoamino acid content and structure [Ohmiya et al., 1989]. However, the possibility that the amino acid sequence of the hagfish iodoprotein may correspond to or be related to a TG domain cannot be ruled out.

The storage of thyroid hormones bound to TG within the follicular lumen is a characteristic feature of vertebrates. The monomeric human TG preprotein consists of a polypeptide comprising 2,768 amino acids (NCBI: NP_003226.4; Uniprot: P01266) [Malthiéry and Lissitzky, 1987; van de Graaf et al., 2001; Holzer et al., 2016]. The classical model of TG’s primary structure is organized into four distinct regions (I, II, III, and IV) [Mercken et al., 1985; Malthiéry & Lissitzky, 1987; Parma et al., 1987; Molina et al., 1996; van de Graaf et al., 2001; Holzer et al., 2016; Citterio et al., 2021]. Region I contains 10 of the 11 TG type 1 repeats, along with two modules known as the linker and hinge, plus an amino-terminal T_4_-forming site. Region II includes three TG type 2 repeats and the 11th TG type 1 repeat, while Region III encompasses all five TG type 3 repeats. The fourth region consists of a domain homologous to esterases, known as the cholinesterase-like (ChEL) domain [Swillens et al., 1986; Park & Arvan, 2004], along with a carboxi-terminal T_3_-forming site. Homology between esterases and TG was first identified by comparing the primary structures of *Torpedo californica* acetylcholinesterase and bovine TG [Schumacher et al., 1986], suggesting that both evolved from a common ancestral gene.

Tandem duplication events in the amino-terminal and amino-central regions, followed by fusion with ChEL domain, enabled TG to acquire new folding, secretion, and TH biosynthesis capabilities [Holzer et al., 2016]. Each TG monomer contains 67 tyrosine and 122 cysteine residues [Mercken et al., 1985; Malthiéry & Lissitzky, 1987; van de Graaf et al., 2001; Holzer et al., 2016]. Four hormonogenic acceptor tyrosines have been mapped at positions 24, 2573, 2587 and 2766 in human TG [Malthiéry & Lissitzky, 1987; Lamas et al., 1989; Dunn et al., 1998]. The principal T_4_-forming site involves the coupling of donor DIT^149^ to acceptor DIT^24^, whereas the predominant T_3_-forming site links MIT^2766^ from one TG monomer to DIT^2766^ in the opposing unit of the dimer [Palumbo et al., 1990; Citterio et al., 2017; 2018].

The TG type 1 domain first emerged at the root of metazoan evolution [Novinec et al., 2006; Yurchenco & Wadsworth, 2004; Takagi et al., 2022]. Each repeats comprising approximately 60 amino acids, can act as protease binders and reversible inhibitors, and have been identified in structurally diverse vertebrate and invertebrate protein families unrelated to TG, including testicans, SPARC-related modular calcium-binding proteins (SMOCs), trophoblast cell-surface antigens (TROPs), splice variants of major histocompatibility complex class II (MHC class II)–associated invariant chains, insulin-like growth factor–binding proteins (IGFBPs), equistatin, saxiphilin, and nidogen [Novinec et al., 2006]. BLAST analysis revealed 992 proteins displaying sequence similarity to the ChEL domain [Belkadi et al., 2012]. In addition to TGs were identified members of carboxylesterases, cholinesterases and neuroligins families.

This study presents a comprehensive bioinformatic investigation into the origin and structural complexity of TG. We analyzed the organization and three-dimensional architecture of TG and its alternative transcripts in the sea lamprey, a modern representative of primitive jawless vertebrates. To assess evolutionary conservation, we examined TG domain architecture and the distribution of cysteine and tyrosine residues across 38 species spanning six distinct taxonomic clades. Phylogenetic analyses were performed on both full-length TG sequences and individual domains to elucidate evolutionary relationships. Based on these findings, we propose that TG originated from a nidogen-like ancestral precursor containing a TG type-1 domain at the base of vertebrate evolution. We further present an evolutionary model detailing the domain diversification events that shaped TG structure, integrating our results with existing molecular and phylogenetic data.

## Results

### Assembly of full-length thyroglobulin from sea lamprey and comparative domain analysis

To date, the only characterized TG sequence from *Petromyzon marinus* was reported by Holzer et al. (2017); however, the ChEL domain remains incomplete. Given the pivotal role of this species in vertebrate evolution, obtaining the full-length TG sequence and examining its structural organization and phylogenetic placement is essential. Accordingly, we retrieved two peptide sequences from the NCBI, UniProt, and Ensembl databases, each exhibiting hallmark features of TG architecture. A 2475 amino-acids sequence (Supplementary Figure S1a), referred to here as TGPM^2475^, is indexed in the Uniprot database (S4R814_PETMA) and extracted from the *Petromyzon marinus* genome assembly available on ENSEMBL (ENSPMAG 00000001187, ENSPMAT00000001350.1), located on scaffold GL476337: 707,134-808,438 forward strand [Smith et al., 2013]. Alignment with human TG (NP_003226.4) using EMBOSS Needle reveals that TGPM^2475^ contains eleven TG type 1 modules, three TG type 2 modules and five TG type 3 modules, linker and hinge domains, and spacers 1, 2 and 3, along with only the first 256 amino acids of the ChEL domain. However, TGPM^2475^ lacks the signal peptide, the N-terminal hormonogenic acceptor tyrosine required for T_4_ formation, the remaining amino acids of the ChEL domain, and the C-terminal hormonogenic site for T_3_ formation. Our upstream analysis to the first amino acid in TGPM^2475^, corresponding to the first cysteine of the TG type 1-1 module in scaffold GL476337, allowed us to identify the first 11 amino acids of exon 2 (YSDSKTSLASA). Among these, three amino acids correspond to the first three amino acids of the TG type 1-1 module (ASA) and the N-terminal hormonogenic acceptor site (Y) (Supplementary Figure S2a,b). The glutamic acid residue (gag codon) overlaps the splice site between exons 1 and 2. In upstream exon 1, following the 687-nt intron, the first 25 amino-terminal amino acids of sea lamprey TG were identified (Supplementary Figure S2a,b). The SignalP 6.0 program predicts that this sequence contains a signal peptide with a cleavage site between positions 24 and 25 (MRTSPLLPATTTLYLVLWIGTISA) with a probability of 0.98 (Supplementary Figure S2c).

A second sequence of 1746 amino acids, corresponding to *Petromyzon marinus* TG was identified in the NCBI database and is referred to here as TGPM^1746^ (*Petromyzon marinus* isolate kPetMar1 chromosome 27, kPetMar1.pri, whole genome shotgun sequence, NCBI reference sequence: NC_046095, XP_032817730.1, Supplementary Figure S3a). Pairwise sequence alignment of TGPM^2475^ and TGPM^1746^ using the EMBOSS Needle program revealed that TGPM^1746^ begins at TG type 1-10 and, notably, contains the complete ChEL domain and the C-terminal hormonogenic site required for T_3_ formation—which are partially and entirely absent in TGPM^2475^, respectively.

The pairwise sequence alignment of TGPM^1746^ and the scaffold GL476337 using the Blat Search Genome, with assembly WUGSC 7.0/petMar2, revealed alignment with the initial 258 amino acids of the ChEL domain (positions 1172-1429 of TGPM^1746^) and the last 115 carboxy-terminal amino acids (positions 1632-1746 of TGPM^1746^), including the final 68 amino acids (positions 1632-1699 TGPM^1746^) of the ChEL domain and the C-terminal hormonogenic site required for T_3_ formation (Y^1745^). However, no alignment was observed between residues 1430-1631 of TGPM^1746^ (202 amino acids) with scaffold GL476337, as this region is absent from the genome data (positions 811252-833817). This absence creates a gap in the ChEL domain of the *Petromyzon marinus* TG sequence previously observed by Holzer et al. [2017]. A 26-amino-acids was identified immediately preceding the TG type 1-10 of TGPM^1746^. This sequence begins with methionine and contains 3 alanines at positions 13, 14 and 15, as well as two cysteines at positions 8 and 19. (MPSRVRACVRAPAAATFLCSALPSPG). The SignalP 6.0 program predicted that TGPM^1746^ contains a signal peptide, with a probability of 0.4996 (Supplementary Figure S3b).

To construct the complete sequence of the *Petromyzon marinus* TG protein sequence, we integrated: 36 upstream amino acids of the TGPM^2475^ identified in the scaffold GL476337, 204 downstream amino acids of the TGPM^2475^ identified in NC_046095 (TGPM^1746^), 202 of which correspond to the missing region in scaffold GL476337 and the 115 carboxy-terminal amino acids identified in the scaffold GL476337. This assembly results in the full-length sea lamprey TG (TGPM), comprising 2831 amino acids. Figure 1 shows the pairwise alignment between TG from *Petromyzon marinus* (TGPM) and *Homo sapiens* (TGHS), generated using EMBOSS Needle. The alignment highlights the positions of conserved domains, cysteine residues, and hormonogenic tyrosines located at the N- and C-terminal regions. Although the primary structures of sea lamprey TG and human TG diverge (identity: 35.1%), the hormonogenic tyrosines and over 90% of the cysteine residues are conserved at identical positions. (Figure 1).

**Figure 1.**
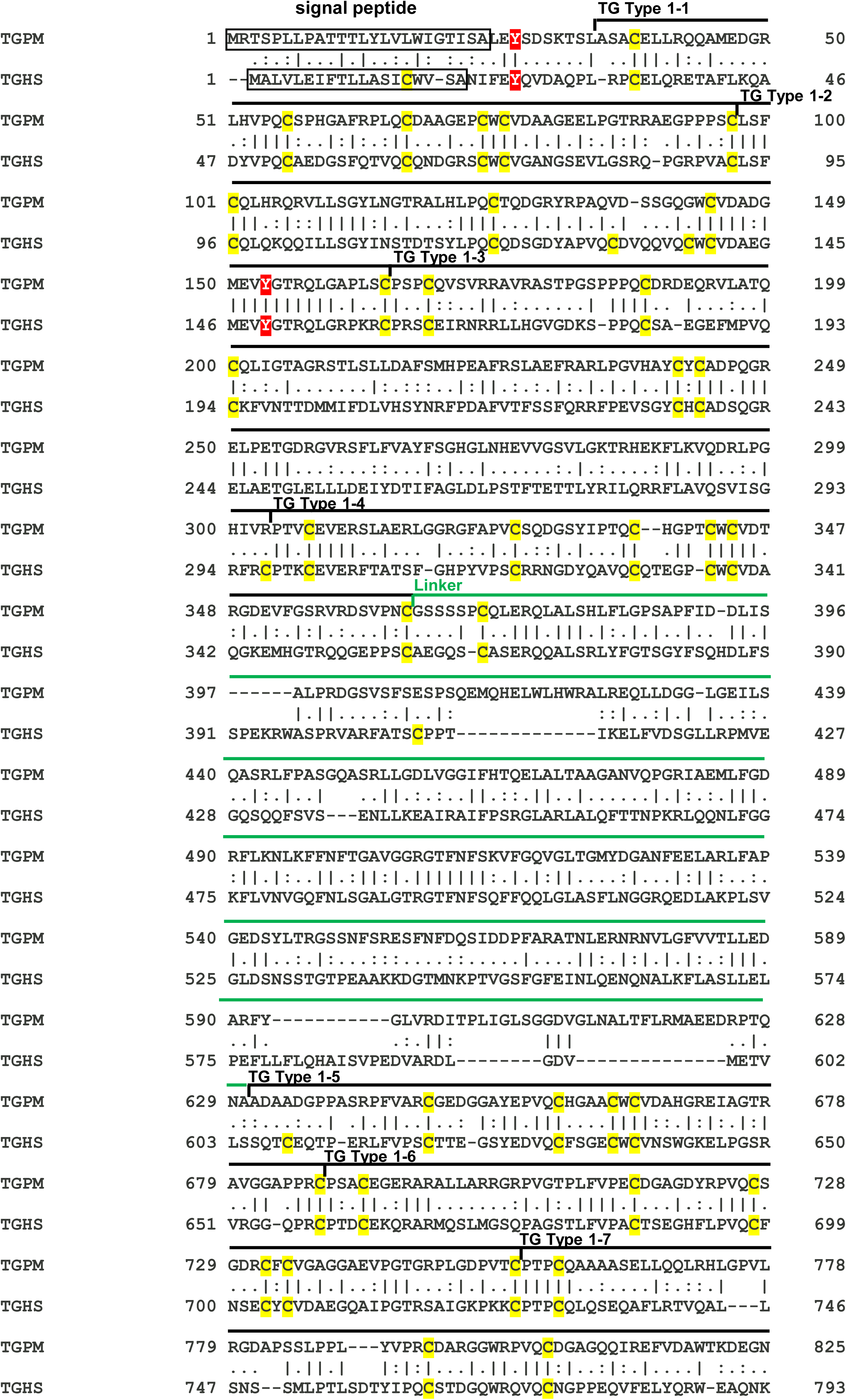
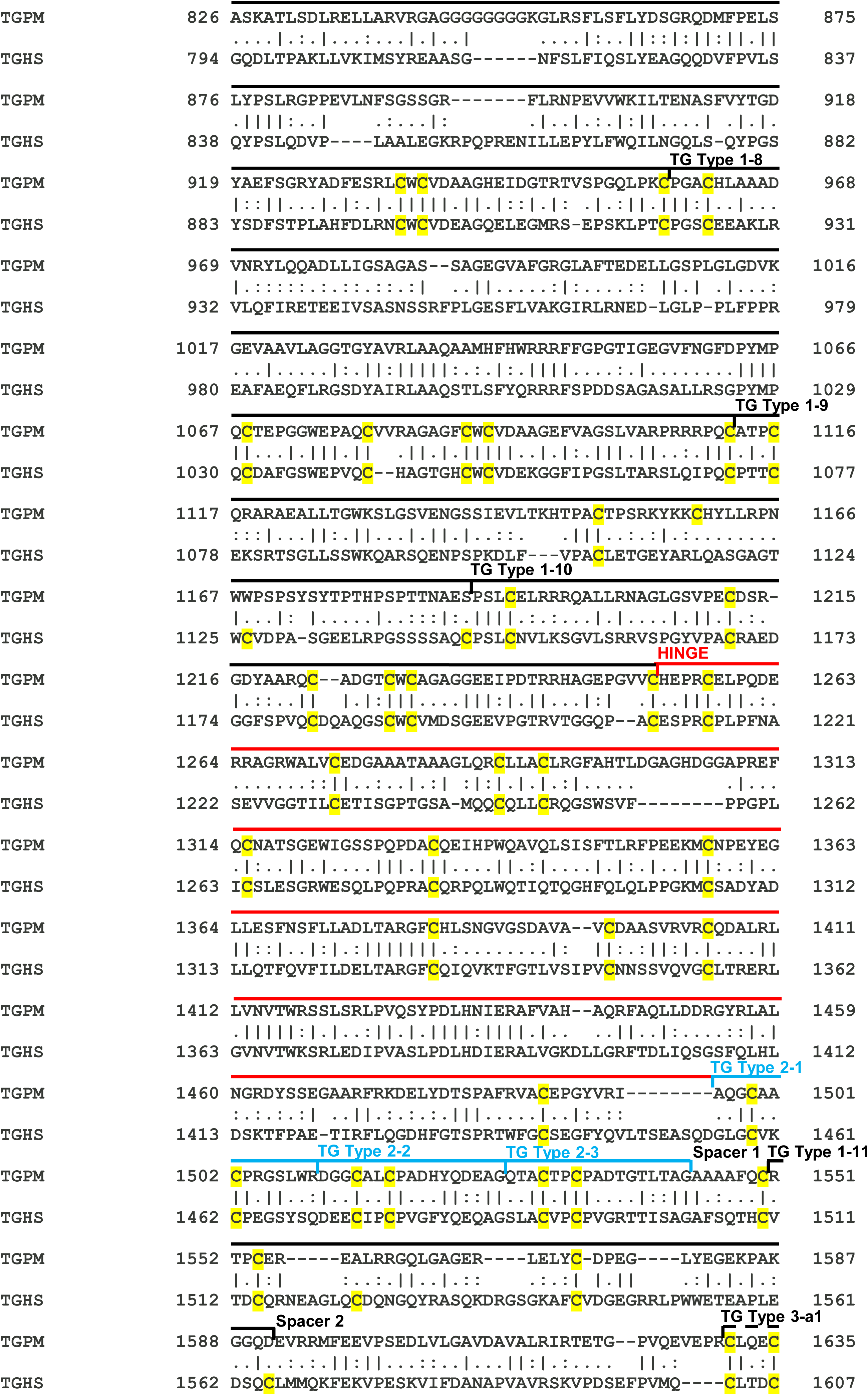
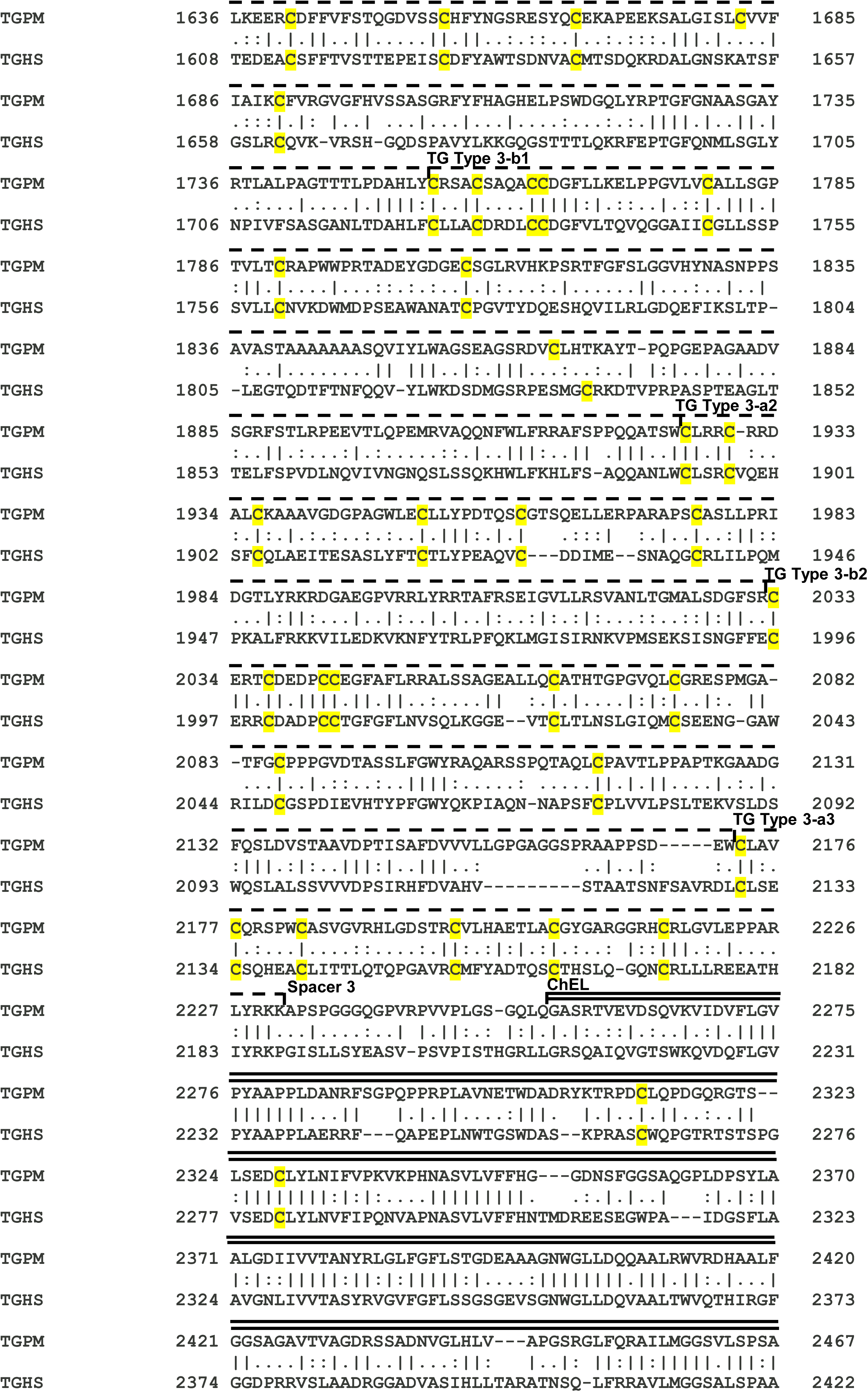
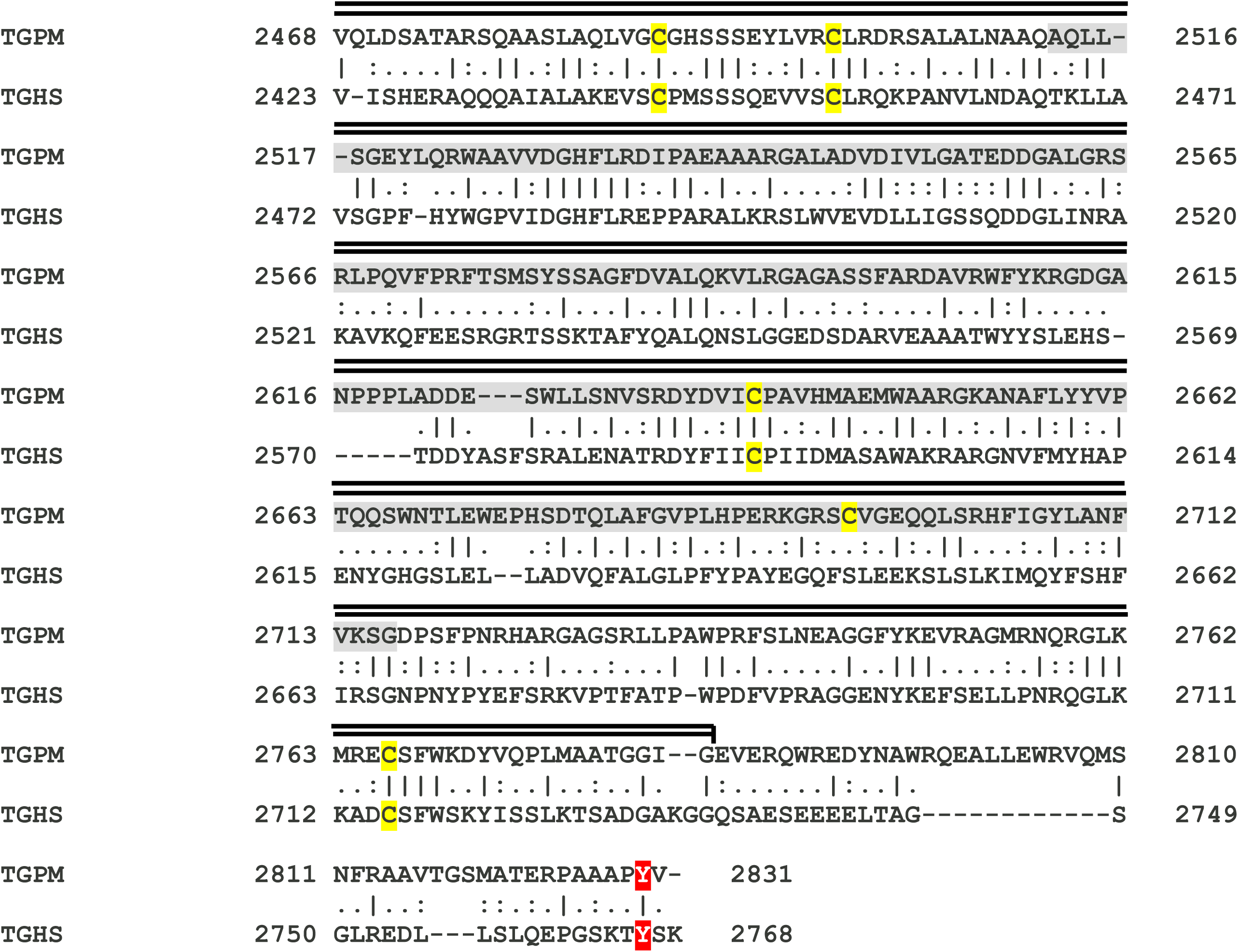
Comparative alignment of full-length thyroglobulin sequences from *Petromyzon marinus* (TGPM) and *Homo sapiens* (TGHS). The alignment was performed using the EMBOSS Needle program. Amino acids are represented using single-letter codes; signal peptides are enclosed in boxes. The figure displays the linker, hinge, and ChEL domains, along with eleven TG type 1 modules, three TG type 2 modules, five TG type 3 modules, and spacers 1, 2, and 3. Cysteine residues are highlighted in yellow. N-terminal hormonogenic acceptor and donor sites (TGPM: Y^27^ and Y^153^; TGHS: Y^24^ and Y^149^), and C-terminal hormonogenic sites (TGPM: Y^2830^; TGHS: Y^2766^) are marked in red. The 202 amino acids absent from the GL476337 scaffold are shaded in gray.

In summary, according to our results the sea lamprey TG gene is a single-copy gene, approximately 121 kb long, divided into 53 exons. Its mRNA encodes 8493 nt, corresponding to coding sequences. The sea lamprey monomeric TG preprotein consists of 24-amino-acid leader peptide followed by a 2807-amino-acid polypeptide (Figure 1).

Remarkably, the comparison of TG sequences between *Petromyzon marinus* (TGPM) and two other lamprey species, *Lampetra fluviatilis* (TGLF) and *Lampetra planeri* (TGLP), using Clustal Omega with its visualization tool Mview, reveals a high degree of sequence conservation among the three species. The sequence identity is 86.9% between TGPM and TGLF and 86.7% between TGPM and TGPLP (Supplementary Figure S4). This finding confirms that our strategy for obtaining the full-length TG of *Petromyzon marinus* was accurate. Additionally, a comparative analysis of TG organization in TGPM, TGLF and TGLP demonstrates that TG in each species contains all regions found in mammalian TG (Supplementary Figure S4). The four TG regions, along with their domains and modules, are similar in size to human TG, with strict conservation of cysteine residues. Furthermore, the amino-terminal and C-terminal hormonogenic sites are highly conserved across all three species (Figure 1, Supplementary Figure S4).

To assess the homology profile between the 11 TG type 1 modules of TGPM and TGHS, we performed a multiple alignment using the T-Coffee program. The analysis yielded a high total consistency score of 75, indicating strong sequence conservation (Supplementary Figure S4). Among the 11 TG type 1 modules, all six cysteine residues are highly conserved across both species. Figure 2a displays the canonical alignment of the 11 TG type 1 modules, highlighting the conserved cysteines. Most repeats exhibit comparable lengths, except for TG Type 1-3, TG Type 1-7, and TG Type 1-8, which contain extended sequence insertions. Remarkably, these insertions are also highly conserved in both species. In TGPM, TG Type 1-3 includes a 40-amino-acid insertion in the middle of the module and an additional 46-amino-acid insertion at its C-terminal end (40 and 45 amino acids, respectively, in human TG). Similarly, TG Type 1-7 harbors a 128-amino-acid insertion centrally (123 in human TG), while TG Type 1-8 features a 100-amino-acid insertion at the N-terminal region, identical in TGHS (Figure 2a).

**Figure 2.**
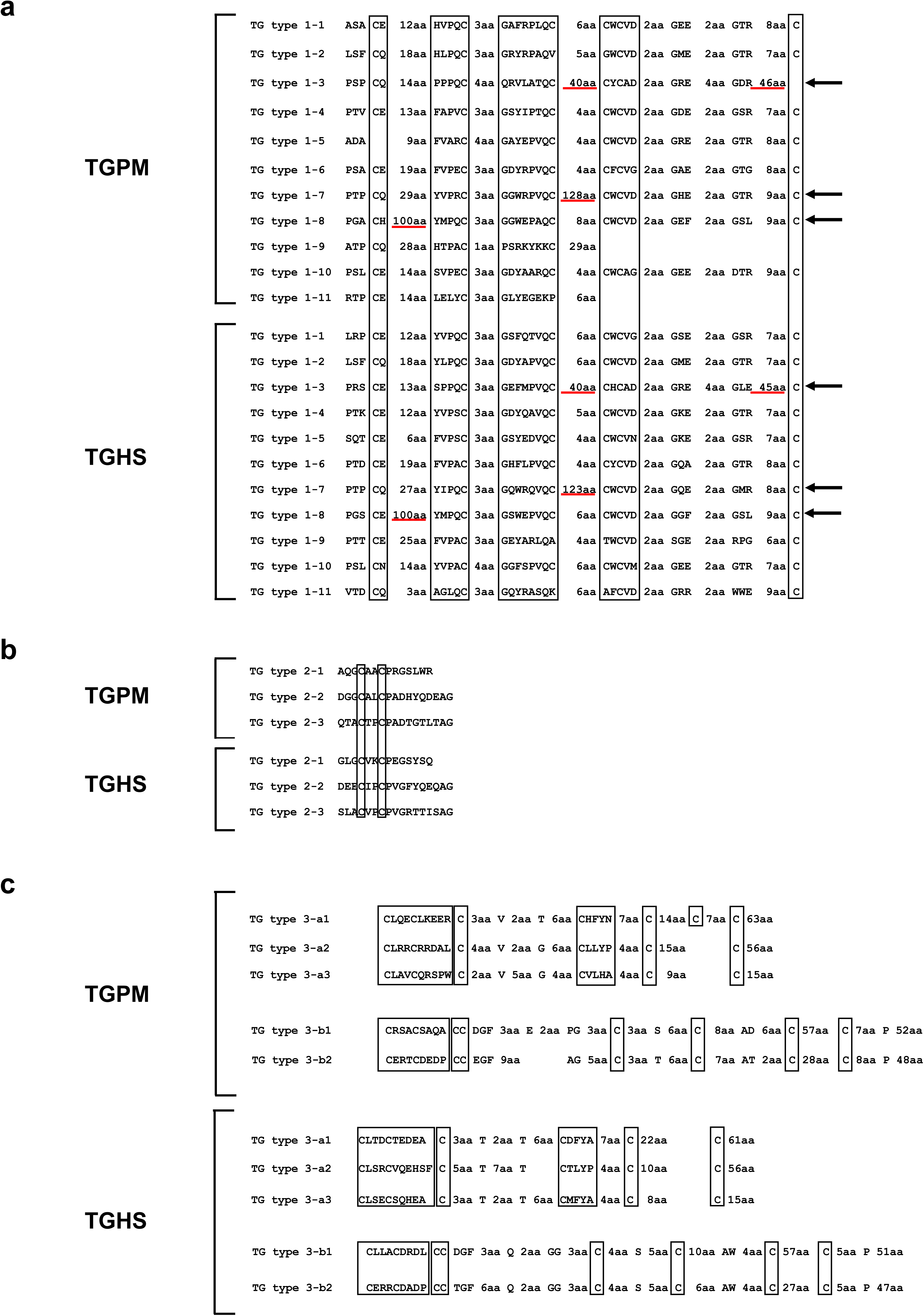
Comparative alignment of TG type 1, TG type 2, and TG type 3 modules of full-length thyroglobulins from *Petromyzon marinus* (TGPM) and *Homo sapiens* (TGHS). Repetitive domain alignments were performed following the criteria established by Parma et al. [1987], Malthiéry et al. [1987], Molina et al. [1996], and van de Graaf et al. [2001]. **a)** Alignment of eleven TG type 1 modules. Six cysteine residues are generally conserved across modules, with exceptions in TG type 1-2, TG type 1-3, TG type 1-5, TG type 1-9, and TG type 1-11 of TGPM, and in TG type 1-9 and TG type 1-11 of TGHS. TG type 1-3, TG type 1-7, and TG type 1-8 exhibit long insertions, highlighted by underlining. **b)** Alignment of three TG type 2 modules. Both cysteine residues are conserved in all modules. **c)** Alignment of five TG type 3 modules. TG type 3-a modules consistently retain six cysteines, while TG type 3-b modules preserve eight cysteines in both TGPM and TGHS. Notably, TG type 3-a1 in TGPM includes an additional cysteine residue. Cysteine positions are boxed.

A phylogenetic analysis was conducted focusing specifically on TG type 1 domains derived from TGs of *Petromyzon marinus* and *Homo sapiens*, with *Mus musculus* included as a reference taxon (Figure 3). Most nodes received strong statistical support (bootstrap values ≥70; Figure 3), and the resulting tree resolved two primary clades. The first clade encompassed TG type 1 domains 1–9 and 1–11 from *Homo sapiens* and *Mus musculus*, as well as TG type 1–11 from *Petromyzon marinus*. The second clade comprised TG type 1 domains 1–1 through 1–8 and 1–10 from all three taxa, along with TG type 1–9 from *Petromyzon marinus*. These findings are consistent with those reported by Belkadi et al. [2012], who investigated TG type 1 domain phylogeny using sequences from sea urchin, amphioxus, zebra finch, mouse, and human. Their study similarly supports a bifurcation of TG type 1 domains into two major branches. As expected, the eleven TG type 1 modules from *Homo sapiens* and *Mus musculus* cluster together in the phylogenetic tree. However, within the *Petromyzon marinus* dataset, only TG type 1–2, 1–8, and 1–11 group with their homologous counterparts from *Homo sapiens* and *Mus musculus*.

**Figure 3.**
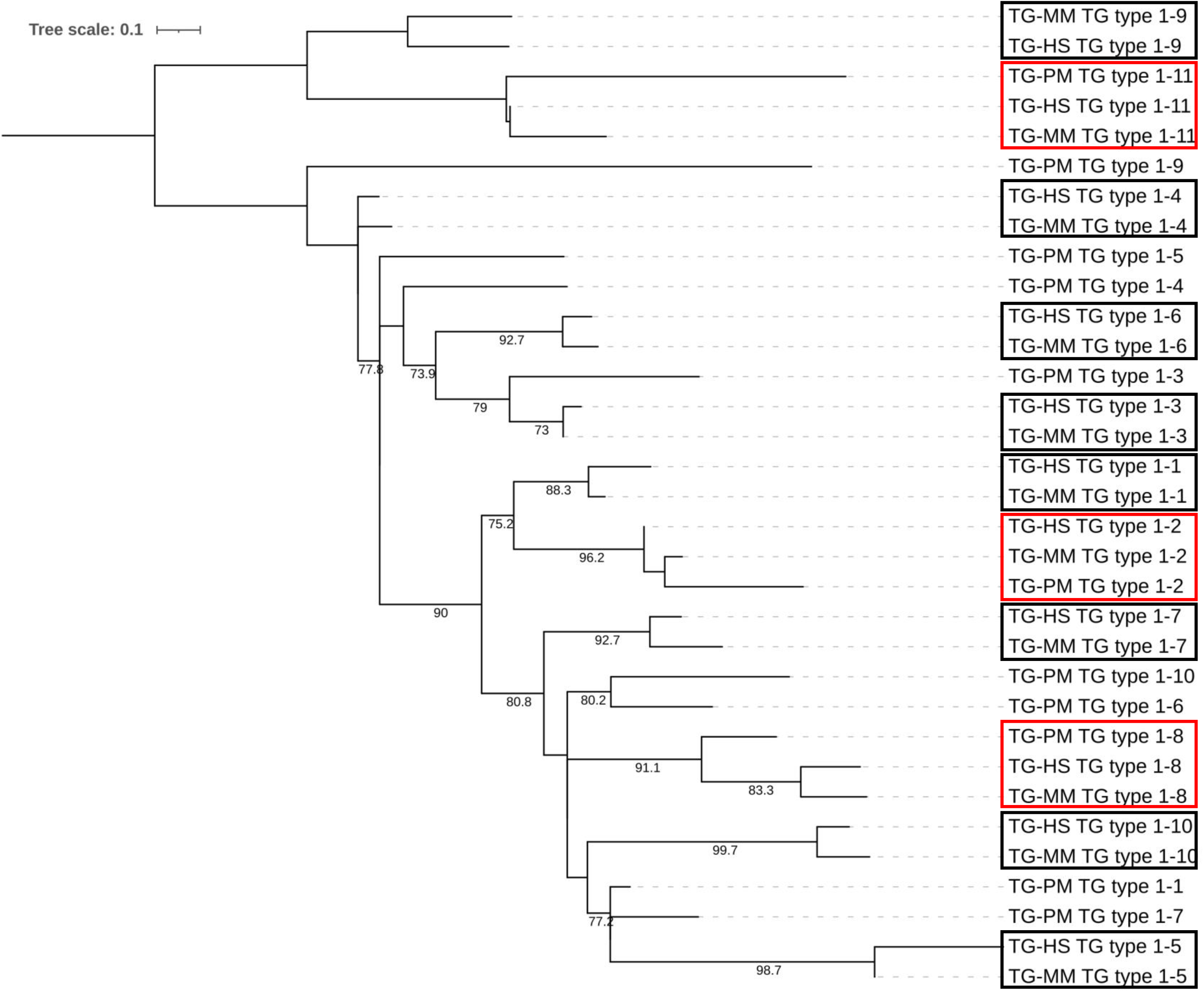
**Phylogenetic analysis of TG type 1 domains from *Homo sapiens*, *Mus musculus* and *Petromyzon marinus.*** The Maximum likelihood phylogeny, constructed from the alignment of TG type 1 amino acid sequences, depicts the evolutionary relationships among the eleven repetitive modules across the three taxa. Ultrafast bootstrap values ≥70, based on 2,000 replicates, are shown above the nodes. *Homo sapiens,* HS; *Mus musculus*, MM; *Petromyzon marinus,* PM.

The alignment of the three TG type 2 modules from TGPM and TGHS, also performed using T-Coffee, yielded a total consistency score of 94, reflecting even stronger sequence conservation than that observed in TG type 1 modules (Supplementary Figure S5). Both cysteine residues are fully conserved between the TG type 2 modules of TGPM and TGHS (Figure 2b).

The alignment of the five TG type 3 modules from TGPM and TGHS produced a total consistency score of 69, indicating moderate sequence conservation (Supplementary Figure S6). This score is lower than those obtained for TG type 1 and type 2 modules, suggesting reduced homology. All six cysteine residues are fully conserved in the TG type 3-a modules (Figure 2c), as are all eight cysteines in the TG type 3-b modules of TGPM and TGHS—except for the eighth cysteine in TG type 3-b1 and TG type 3-b2, which could not be aligned by T-Coffee (Supplementary Figure S6). Additionally, an extra cysteine residue was identified at position 1682 in the TG type 3-a1 module of TGPM (Supplementary Figure S6).

A comparative sequence analysis between positions 1189 and 2831 of TGPM and the full length TGPM^1746^ isoform, conducted using the EMBOSS Needle program, revealed five major differences at the following positions: 1388 (hinge domain), 1673 to 1710 (TG type 3a1), 1831 to 1850 (TG type 3b1), 1881 (TG type 3b1) and 2233 to 2235 (Spacer 3) of the TGPM (Figure 4). Interestingly, in the latter region, TGPM^1746^ exhibits a 39 amino acid sequence containing 8 cysteines, resembling the TG type 3b modules (SLGTSCAARCGLYSPGQPCQCNRECQRFGDCCPDAAVCL).

**Figure 4.**
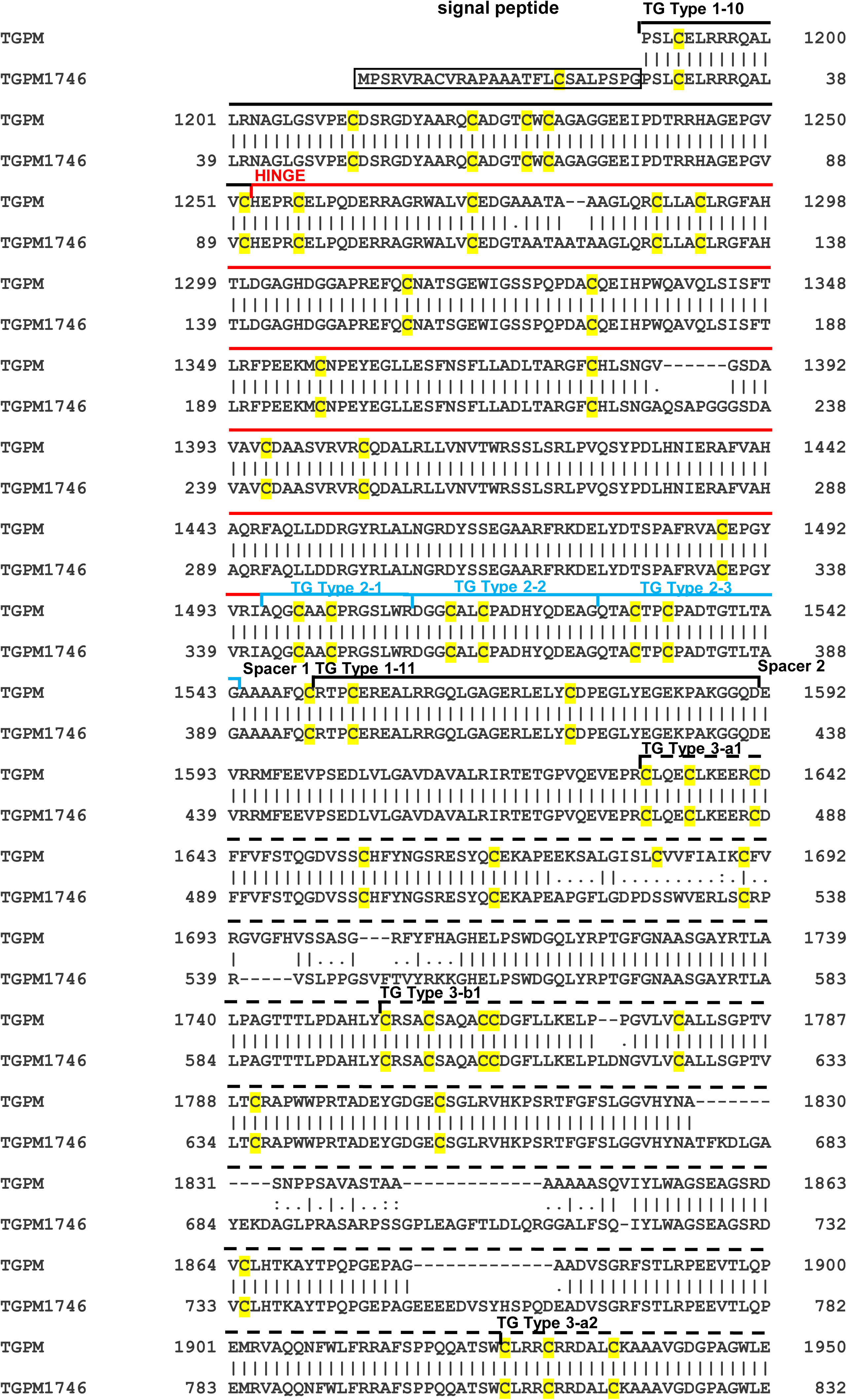
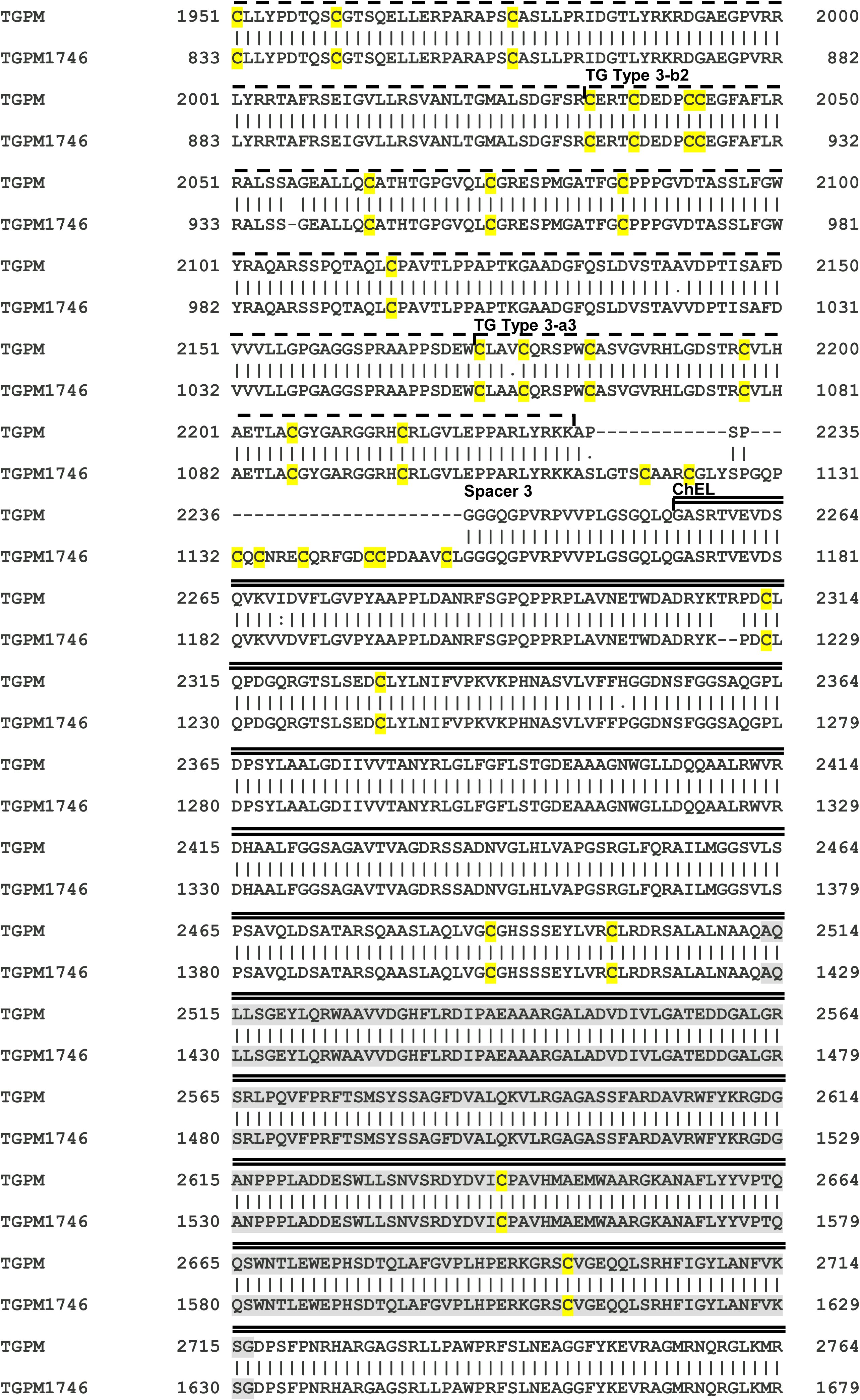
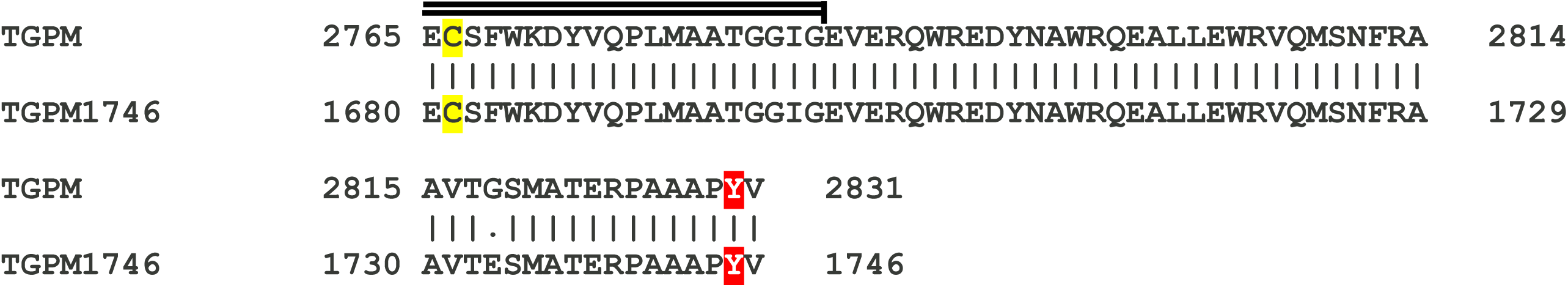
Comparative alignment of positions 1189–2831 from the full-length thyroglobulin of *Petromyzon marinus* (TGPM) and its 1746-amino acid isoform (TGPM^1746^). The alignment was performed using the EMBOSS Needle program. Amino acids are represented using single-letter codes. TGPM^1746^ begins at the TG type 1–10 module and includes the hinge domain, TG type 1–11 module, three TG type 2 modules, and five TG type 3 modules, spacers 2 and 3, the complete ChEL domain, and the C-terminal hormonogenic site. The signal peptide of TGPM^1746^ is boxed. Cysteines residues are highlighted in yellow, and the C-terminal hormonogenic sites (TGPM: Y^2830^, TGPM^1746^: Y^1745^) are marked in red. Five major differences are observed in the hinge domain, TG type 3a1, TG type 3b1, and spacer 3.

### Three-dimensional atomic structure alignment of thyroglobulins from *Petromyzon marinus* and *Homo sapiens*

To compare the three-dimensional atomic structure of the two sea lamprey TG variants with human TG, as well as with each other, we generated structural models for TGPM^2475^ and TGPM^1746^, as no PDB entries were available. Homology modeling was performed using the Swiss Model program based on the sequences of TGPM^2475^ (UniProt: S4R814_PETMA; Supplementary Figure S1b) and TGPM^1746^ (NCBI: XP_032817730.1; Supplementary Figure S3c), using the 3D structure of *Bos taurus* TG (PDB: 7N4Y; resolution: 2.61 Å) as a template (https://www.rcsb.org/structure/7N4Y) [Kim et al., 2021]. The human TG 3D structure (PDB: 6SCJ; resolution: 3.60 Å) was not selected as a template due to its lower resolution relative to PDB 7N4Y.

The structural comparison between TGPM^2475^ and TGPM^1746^ monomers (chain B) and the human TG monomer (chain A) were performed using the Matchmaker command in UCSF ChimeraX. The superposition of TGPM^2475^ and TGHS yielded an RMSD of 1.219 Å across 1,271 overlapping atom pairs, with a sequence alignment score of 4,688.8 (Supplementary Figure S8). Similarly, the alignment of TGPM^1746^ and TGHS resulted in an RMSD of 1.076 Å based on 806 overlapping atom pairs, with a sequence alignment score of 2,879.4 (Supplementary Figure S8). A consistent structural overlap was observed across the four conserved regions (I-IV) of TGPM^2475^ or TGPM^1746^, corresponding to the homologous regions in TGHS. These results supports the structural homology between the proteins, extending from the TG type 1-1 module to the first 256 amino acids of the ChEL domain in TGPM^2475^, and from the TG type 1-10 module to the carboxy-terminal end in TGPM^1746^.

We next, compared the three-dimensional atomic structures of TGPM^2475^ and TGPM^1746^ (both chain A). Structural superposition using the Matchmaker tool revealed a high degree of overlap between the complexes, with an RMSD of 0.295 Å across 1,246 aligned atom pairs and a sequence alignment score of 6,249 (Supplementary Figure S9).

Finally, a structural 3D model of TGPM we generated using the Swiss Model (Figure 5, Supplementary Video 1), based on PDB 7N4Y of *Bos taurus* TG. The full-length sea lamprey TG (TGPM, chain B) was subsequently aligned with human TG (TGHS, chain A), yielding an RMSD of 1.191 Å across 1,449 overlapping atom pairs and a sequence alignment score of 5,212.9 (Figure 5, Supplementary Videos 2 and 3). As previously observed for TGPM^2475^ and TGPM^1746^, the alignment between TGPM and TGHS revealed a consistent structural overlap across the four conserved regions (I–IV) of sea lamprey TG, corresponding to the four analogous regions in human TG.

**Figure 5.**
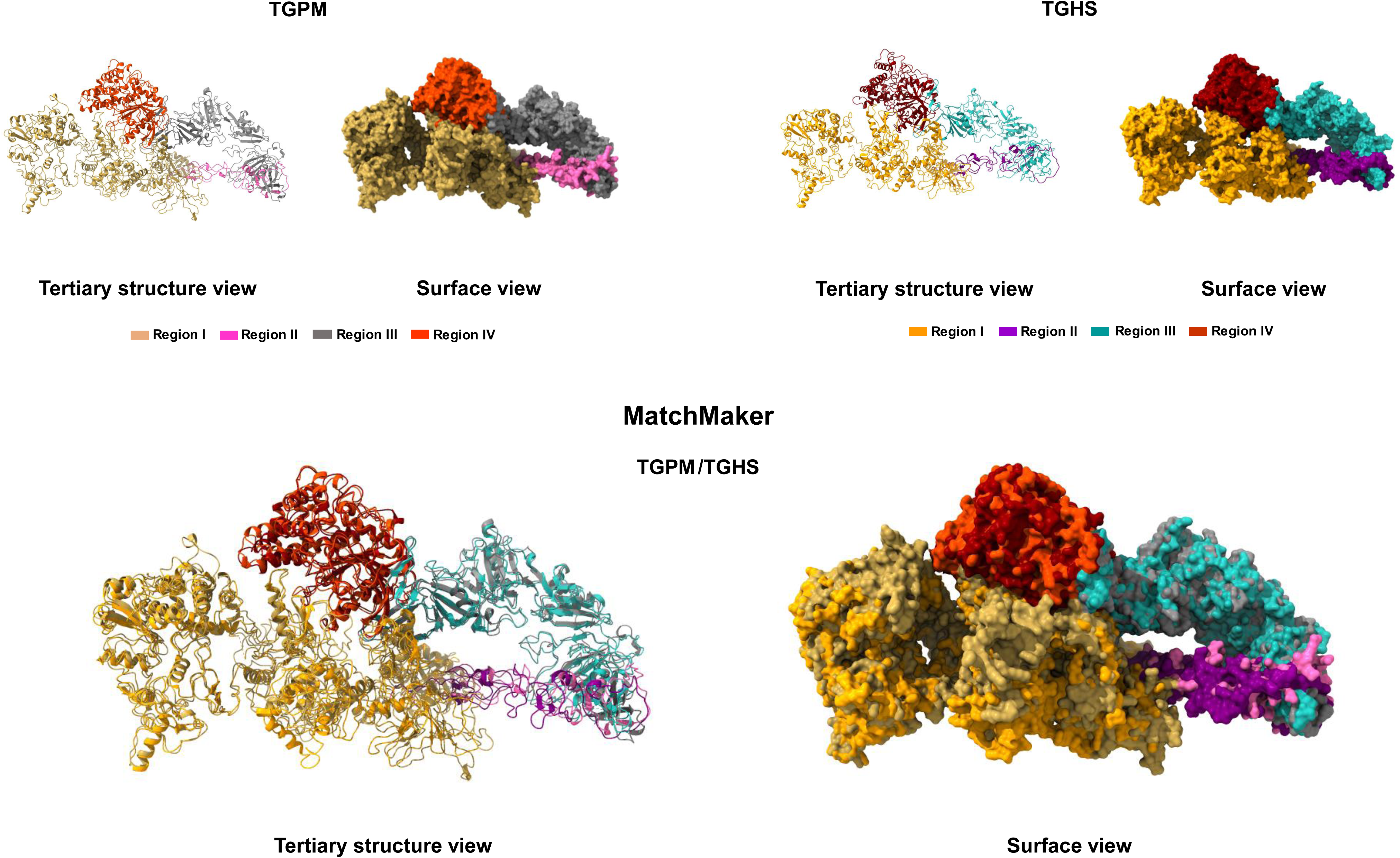
Three-dimensional atomic structure alignment of full-length thyroglobulins from *Petromyzon marinus* (TGPM) and *Homo sapiens* (TGHS). The top panel shows the homology models of TGPM and TGHS. The bottom panel illustrates the structural superposition of thyroglobulin monomers (TGPM with TGHS) across the four canonical regions (I-IV) of the classical model, performed using the Matchmaker command in UCSF ChimeraX. The left panel displays the superposition of monomers without surface rendering, while the right panel includes surface visualization.

### Comparative analysis of full-length thyroglobulin sequences from 38 vertebrate species

Comparative analysis of TG structural organization across 38 vertebrate species—including mammals (*Homo sapiens, Mus musculus*, *Bos taurus*, *Canis lupus familiaris*, *Cavia porcellus*, *Macaca fascicularis*, *Macaca mulatta*, *Pan troglodytes*, *Pongo pygmaeus*, *Gorilla gorilla*, *Panthera leo*, *Rattus norvegicus*, *Trichechus manatus latirostris*); birds (*Gallus gallus*, *Columba livia*, *Taeniopygia guttata*, *Struthio camelus*, *Larus michahellis*); reptiles (*Python bivittatus*, *Chelonia mydas*, *Crotalus tigris*, *Eublepharis macularius*); amphibians (*Xenopus tropicalis*, *Aquarana catesbeiana*); ray-finned fishes (*Danio rerio*, *Oryzias latipes*, *Stegastes partitus*, *Xiphophorus maculatus*, *Cynoglossus semilaevis*, *Cyprinus carpio*, *Carassius auratus*, *Astyanax mexicanus*, *Clupea harengus*, *Lepisosteus oculatus*); and jawless vertebrates (*Petromyzon marinus*, *Lampetra fluviatilis*, *Lampetra planeri*)—reveals that TG in each species retains the full complement of domains found in vertebrate TG (Supplementary Figure S10). As previously described, the canonical TG architecture spans from the N- to the C-terminus and includes: a signal peptide; four TG type 1 modules; a linker domain; six additional TG type 1 modules; a hinge domain; three TG type 2 modules; spacer 1; one TG type 1–11 module; spacer 2; five TG type 3 modules; spacer 3; and the ChEL domain (Supplementary Figure S10). Across the species analyzed, both the overall protein length and the dimensions of individual domains are broadly conserved, with four notable exceptions (Table 1, Supplementary Figure S10). Specifically, *Oryzias latipes*, *Stegastes partitus*, *Xiphophorus maculatus*, and *Cynoglossus semilaevis*—all members of the Actinopterygii clade—exhibit markedly reduced TG protein sizes, primarily due to a ∼95% sequence loss in the TG type 1–7 module.

**Table 1.**
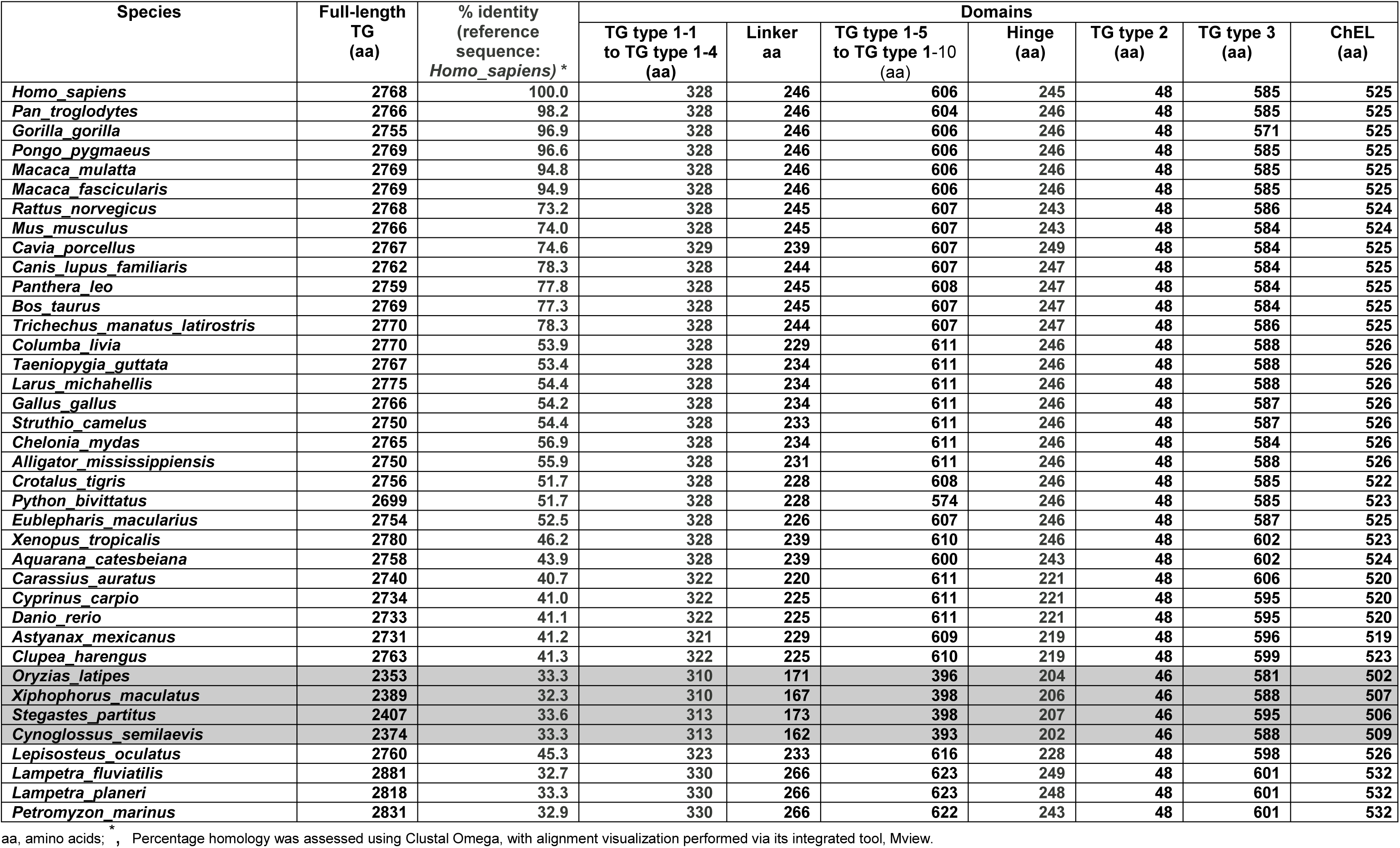
Size of full-length thyroglobulin and its constituent domains across 38 vertebrate species.

In 37 of the 38 species analyzed, the TG gene is retained as a single-copy locus, as evidenced by the ENSEMBL phylogenetic tree (https://www.ensembl.org/Homo_sapiens/Gene/Compara_Tree?collapse=none;db=core;g=ENSG00000042832;r=8:132866958-133134903). This pattern may reflect either the elimination of redundant copies via negative selection, or the functional constraints of TG, whose specialized role may preclude genetic redundancy. The sole exception is *Carassius auratus* (goldfish), which harbors two major TG gene copies. At the protein level, the second copy exhibits 93% sequence identity with the first but lacks the C-terminal region of the ChEL domain and the hormonogenic site T_3_. Additionally, a third, incomplete copy encodes only 285 amino acids corresponding to the C-terminal portion of the ChEL domain.

Cysteine residues are remarkably conserved across all eleven TG type 1, three TG type 2, and five TG type 3 modules, underscoring the structural integrity of these domains (Supplementary Figure S10). However, a few exceptions were identified beyond those previously noted in jawless vertebrates. In *Gorilla gorilla*, the third, fourth, and fifth cysteines of the TG type 1 module are absent. In *Xenopus tropicalis* and *Aquarana catesbeiana*, the third and fourth cysteines are missing from TG types 1–8. In *Python bivittatus*, the sixth cysteine is absent from TG types 1–8, and the first and second cysteines are missing from TG types 1–9. Finally, as previously noted, Oryzias latipes, Stegastes partitus, Xiphophorus maculatus, and Cynoglossus semilaevis—all Actinopterygii species—lack all six conserved cysteines from the TG type 1–7 module, owing to its complete absence in their protein architecture.

Across the alignment, 66 human TG tyrosine positions were analyzed using a strict denominator that included all 38 vertebrate taxa, treating gaps as valid but non-conserved (Figure 6a, Supplementary Table S1). Conservation values ranged from 16% to 100%, with a median of 76% and a mean of 74%, indicating that most tyrosine residues are moderately to highly conserved across vertebrates. Notably, 20 positions (30% of all sites) exhibited high conservation (≥90%), including four universally conserved tyrosine residues (positions 24, 149, 234, and 1965), which likely correspond to structurally or functionally essential motifs. Also within this group, tyrosine^2766^ can be considered, as it is present in all species except Oryzias latipes, likely due to missing C-terminal sequences (Supplementary Figure S10). In contrast, only three sites (4.5%) were highly variable (<30%), underscoring the rarity of extreme divergence. The top 10 tyrosine positions all reached conservation levels ≥95%, reinforcing the notion of strong evolutionary constraint at specific loci. Species-level conservation mirrored clade-level phylogenetic trends. The top 10 species were exclusively mammals, with several primates and rodents preserving 95–100% of human TG tyrosine positions (Figure 6b, Table 2). Archosauria (birds, Chelonia, and Alligator) present intermediate levels, while Lepidosauria exhibit slightly lower preservation rates (Figure 6b). Amphibians fall in the middle range, highlighting moderate divergence from the mammalian pattern (Figure 6b). Conversely, the bottom 10 species comprised lampreys (Hyperoartia) and ray-finned fishes (Actinopterygii), with preservation values ranging from 33% to 53% (Figure 6b, Table 3). These findings integrate both positional and taxonomic perspectives, revealing that while a subset of tyrosine residues remains nearly invariant throughout vertebrate evolution, lineage-specific divergence is concentrated in lampreys and many actinopterygians—consistent with their basal phylogenetic placement and distinct TG domain adaptations.

**Figure 6.**
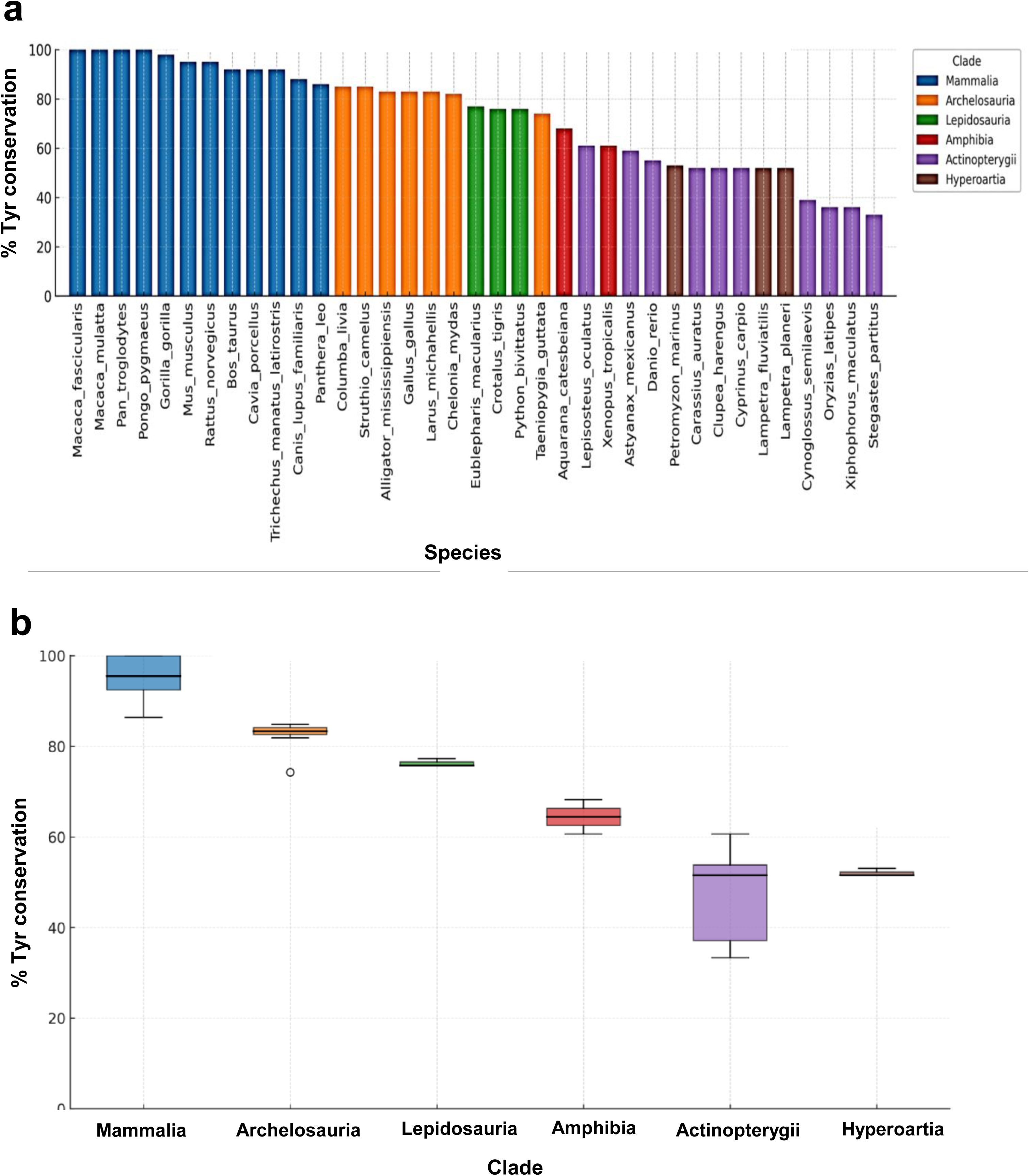
Barplot of tyrosine conservation in *Homo sapiens* thyroglobulin. A total of 66 tyrosine positions from the *Homo sapiens* thyroglobulin sequence were analyzed across all 38 species included in the multiple sequence alignment performed Clustal Omega (see Supplementary Figure S10). Alignment gaps were treated as valid but non-conserved. **a)** Percentage of tyrosine conservation per species. **b)** Percentage of tyrosine conservation per taxonomic clade.

**Table 2.**
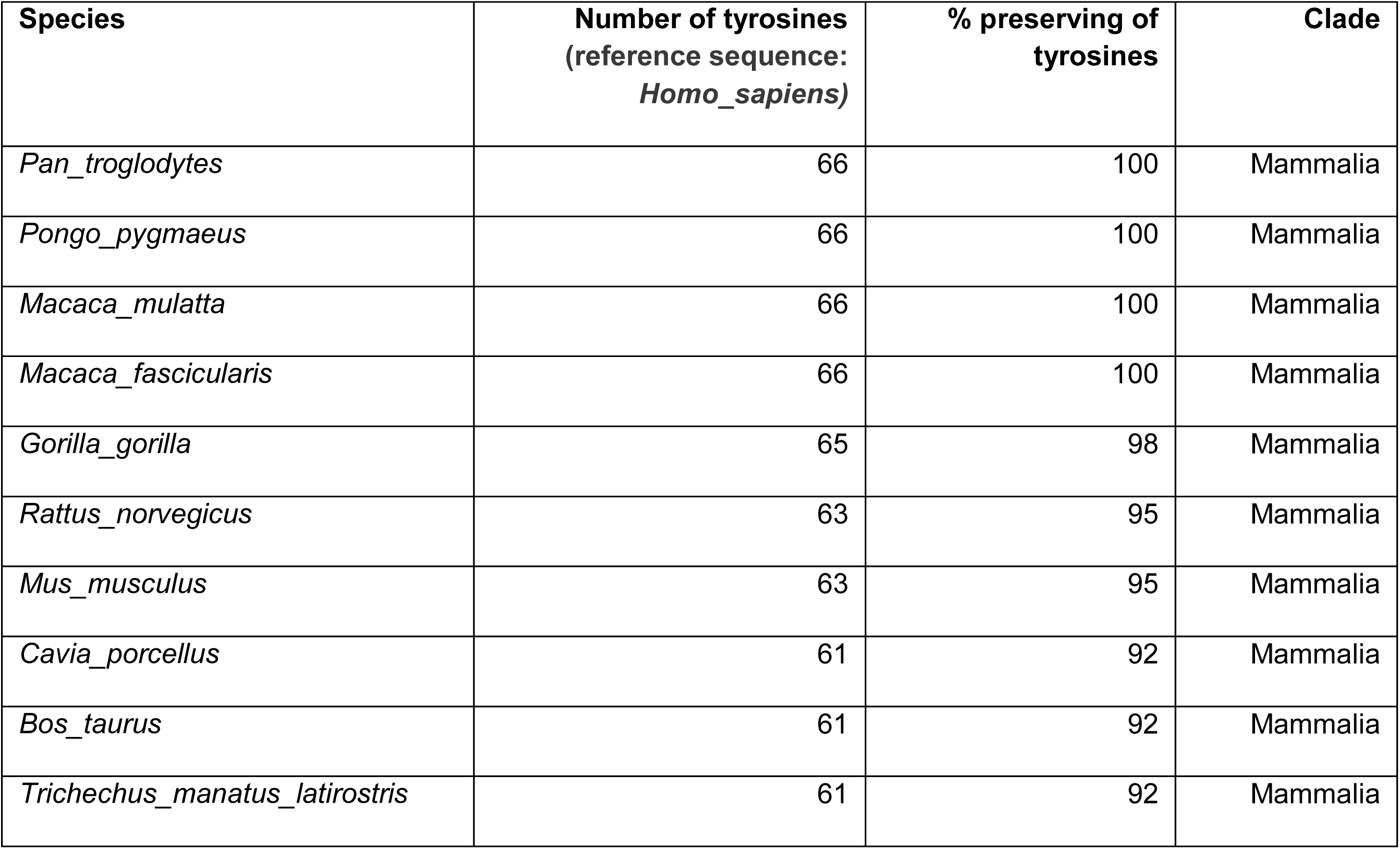
Vertebrate species showing the highest conservation of tyrosine residue sites found in human thyroglobulin.

**Table 3.** Vertebrate species showing the lowest conservation of tyrosine residue sites found in human thyroglobulin.

| **Species** | **Number of tyrosines**<br>**(reference sequence:**<br>***Homo_sapiens*)** | **% preserving of**<br>**tyrosines** | **Clade** |
| --- | --- | --- | --- |
| *Stegastes_partitus* | 22 | 33 | Actinopterygii |
| *Xiphophorus_maculatus* | 24 | 36 | Actinopterygii |
| *Oryzias_latipes* | 24 | 36 | Actinopterygii |
| *Cynoglossus_semilaevis* | 26 | 39 | Actinopterygii |
| *Lampetra_fluviatilis* | 34 | 52 | Hyperoartia |
| *Carassius_auratus* | 34 | 52 | Actinopterygii |
| *Clupea_harengus* | 34 | 52 | Actinopterygii |
| *Cyprinus_carpio* | 34 | 52 | Actinopterygii |
| *Lampetra_planeri* | 34 | 52 | Hyperoartia |
| *Petromyzon_marinus* | 35 | 53 | Hyperoartia |

A protein disorder prediction for full-length TG sequences was also performed using the AIUPred web server. Figure 7 illustrates the disorder profiles of selected representative taxa, while Supplementary Figures S18–S24 display results for all 38 species analyzed. Intrinsically disordered regions are polypeptide segments that operate through highly flexible conformational states rather than adopting a single, well-defined structure. According to our analysis, TG is a predominantly a structured protein, yet it harbors a conserved disordered region at its C-terminus, encompassing the terminal segment of the ChEL domain and the hormonogenic site responsible for T_3_ formation. This disordered region was consistently detected across the dataset, with the exception of *Python bivittatus* and *Crotalus tigris* (Supplementary Figures S21), *Cynoglossus semilaevis* (Supplementary Figures S23), and three jawless vertebrates—*Petromyzon marinus*, *Lampetra fluviatilis*, and *Lampetra planeri* (Figure 7, Supplementary Figures S24) — likely reflecting divergent C-terminal sequences. Among the otherwise highly structured TG proteins, the least structured profiles were observed in Bos taurus, Canis lupus familiaris, Panthera leo (Supplementary Figures S19), and the three jawless vertebrates (Figure 7, Supplementary Figures S24).

**Figure 7.**
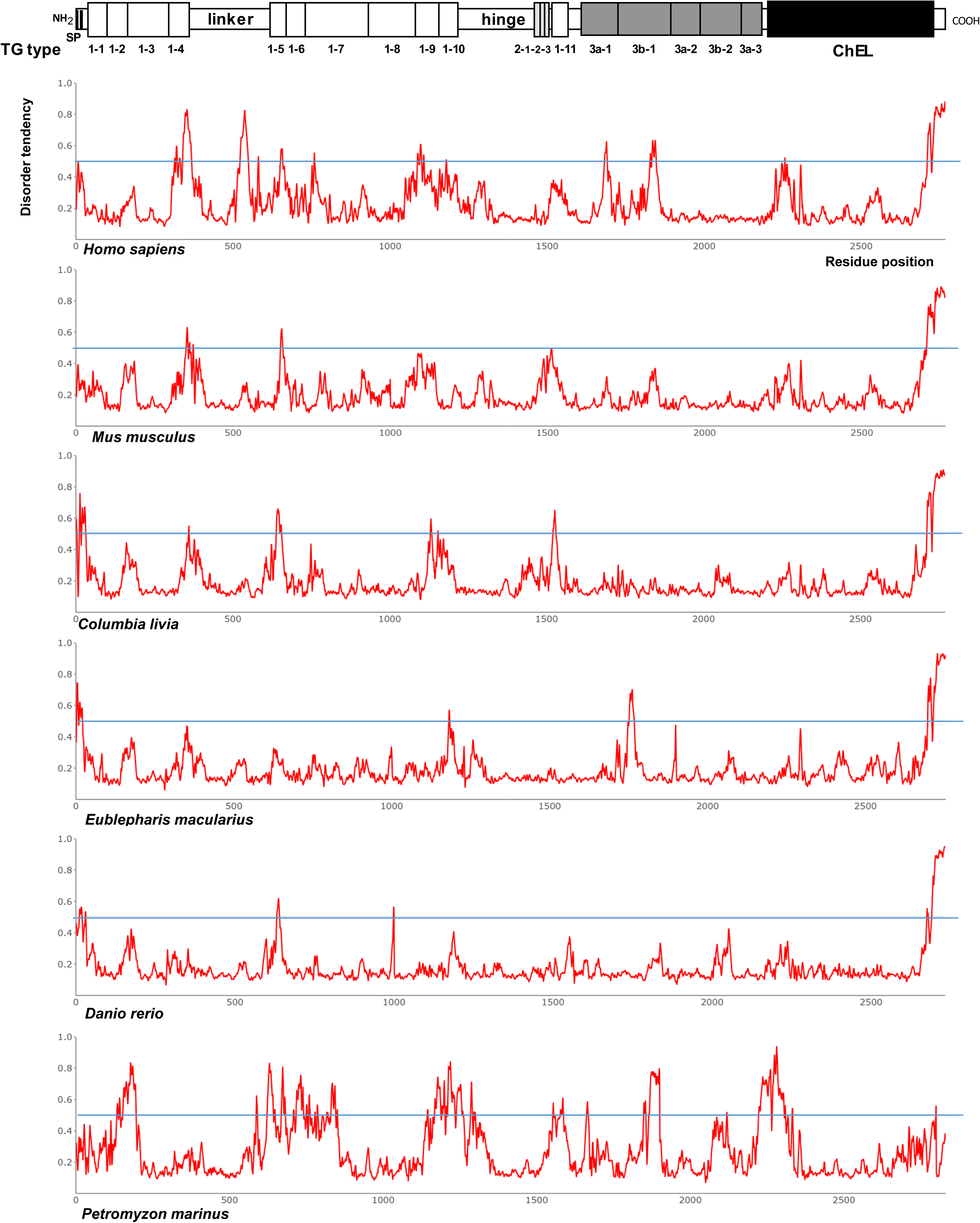
Protein disorder prediction for full-length thyroglobulin sequences from representative taxa. Disorder analysis was performed using the AIUPred web server with the “AIUPred-only disorder” setting. Residues with a predicted score ≥0.5 were classified as disordered, while those below this threshold were considered ordered. At the top, the classical model of thyroglobulin’s primary structure is shown to scale, with the signal peptide (SP), TG type-1, TG type-2, and TG type-3 modules, linker and hinge domain, spacers 1, 2, and 3, and the cholinesterase-like (ChEL) homology domain represented as boxed elements.

The full-length TG amino acid sequences were used to construct a maximum likelihood tree encompassing the 38 vertebrate species (Figure 8). The resulting tree revealed the expected overall topology, preserving established phylogenetic relationships among taxa. Most nodes received strong statistical support (bootstrap scores ≥70; Figure 8). To further explore TG evolution, we extended the analysis to include major TG domains individually (Supplementary Figure S11-S17). Phylogenetic trees derived from TG types 1-1 to 1-4 (Supplementary Figure S11), TG types 1-5 to 1-10 (Supplementary Figure S13), and TG type 3 (Supplementary Figure S15) domains recapitulated the relationships observed in the full-length TG tree, revealing consistent topologies across datasets. All four trees identified *Hyperoartia* as the earliest-branching clade. Topology tests revealed specific clustering patterns for the remaining domains analyzed. In the linker domain, the jawless clade grouped with *Oryzias latipes*, *Stegastes partitus*, *Xiphophorus maculatus*, and *Cynoglossus semilaevis* (Supplementary Figure S12). In contrast, the hinge domain placed *Hyperoartia* within the *Mammalia* clade (Supplementary Figure S14), while TG type 2 domains clustered it with *Actinopterygii* (Supplementary Figure S16). Finally, in the ChEL domain, the jawless clade grouped with *Amphibia*, *Lepidosauria*, *Archelosauria*, and *Mammalia* clades (Supplementary Figure S17). The congruent topologies of TG type 1 and TG type 3 domains support the hypothesis that TG type 3 repeats originated from TG type 1 repetitive elements.

**Figure 8.**
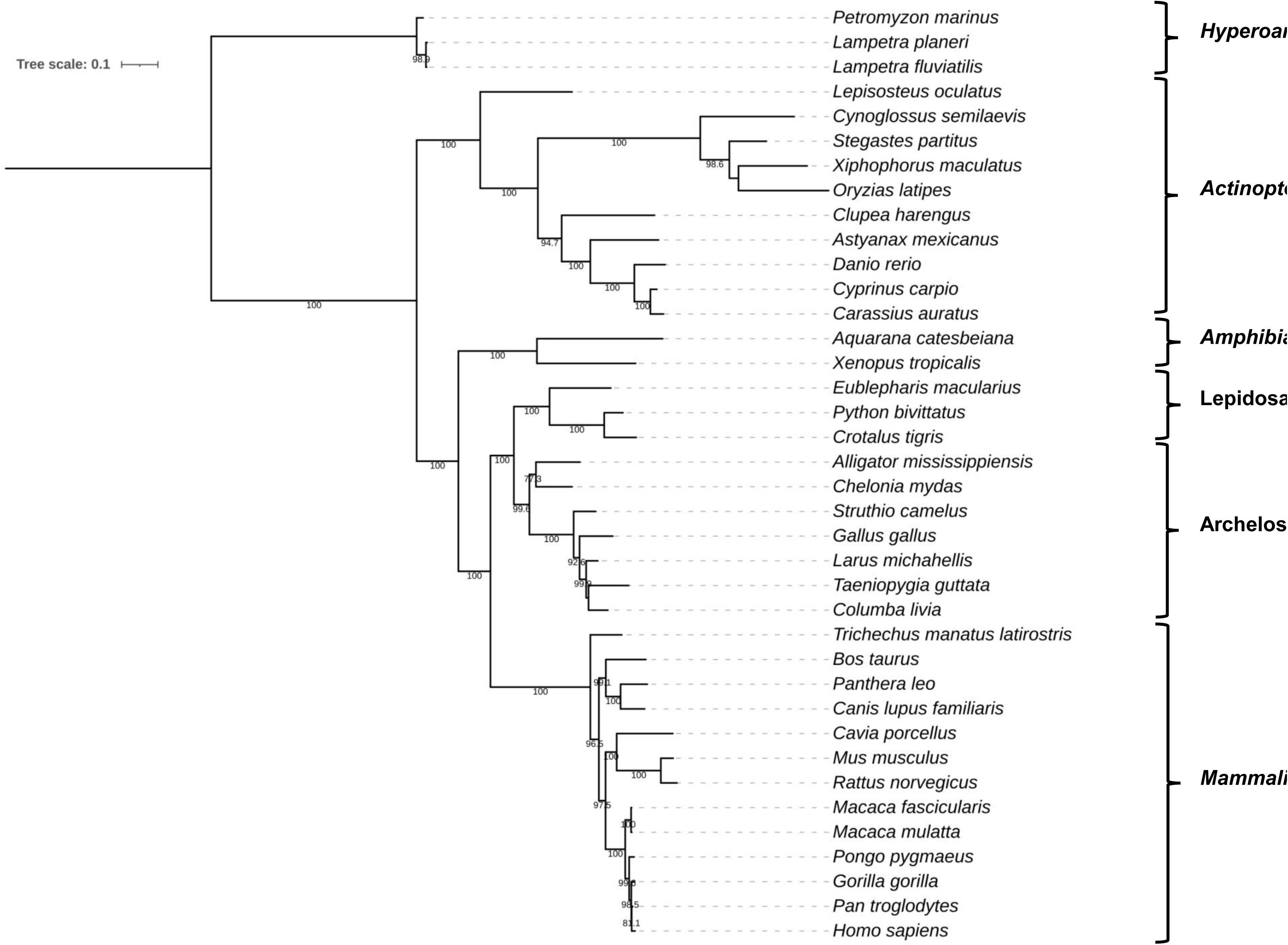
Phylogenetic analysis of full-length thyroglobulins from 38 species. The maximum likelihood phylogeny, constructed from the alignment of thyroglobulin amino acid sequences, depicts the evolutionary relationships among the sampled taxa. Ultrafast bootstrap values ≥70, based on 2,000 replicates, are shown above the nodes. Major taxonomic clades represented include *Hyperoartia, Actinopterygii, Amphibia, Lepidosauria, Archelosauria,* and *Mammalia*.

### Comparative analysis of TG type 1 and TG type 2 modules between nidogen2 and thyroglobulin

As previously noted, numerous proteins incorporate the TG type 1 module within their structure. Notably, SMOC and nidogen emerge as key representatives, given their presence at the root of vertebrate evolution. Our comprehensive analysis reveals that cyclostomes—such as the sea lamprey—exhibit a modular architecture consistent with that observed in mammalian TG proteins. This observation supports the hypothesis that an ancestral protein may have functioned as a TG type 1 precursor, facilitating its complexation during early evolutionary stages.

A targeted phylogenetic analysis was performed on TG type 1-1 and 1-2 domains derived from SMOCs, nidogens, and TGs of *Petromyzon marinus*, *Homo sapiens*, and *Mus musculus* (Figure 9). Most nodes received robust statistical support (bootstrap values ≥70), and the resulting tree resolved two principal clades. The first clade encompassed TG type 1-1 and 1-2 modules from SMOC1 and SMOC2 across all three species (Figure 9). The second clade included TG type 1-1 and 1-2 domains from nidogen-1, nidogen-2, and TG, likewise represented in *Petromyzon marinus*, *Homo sapiens*, and *Mus musculus* (Figure 9). Notably, the repetitive TG type 1 domains of TG did not cluster with those of SMOC. This phylogenetic pattern suggests that TG type 1 repeats found in nidogen and TG may have originated from a shared ancestral precursor, as indicated by the consistent co-clustering of homologous domains from both protein families within a unified evolutionary lineage.

**Figure 9.**
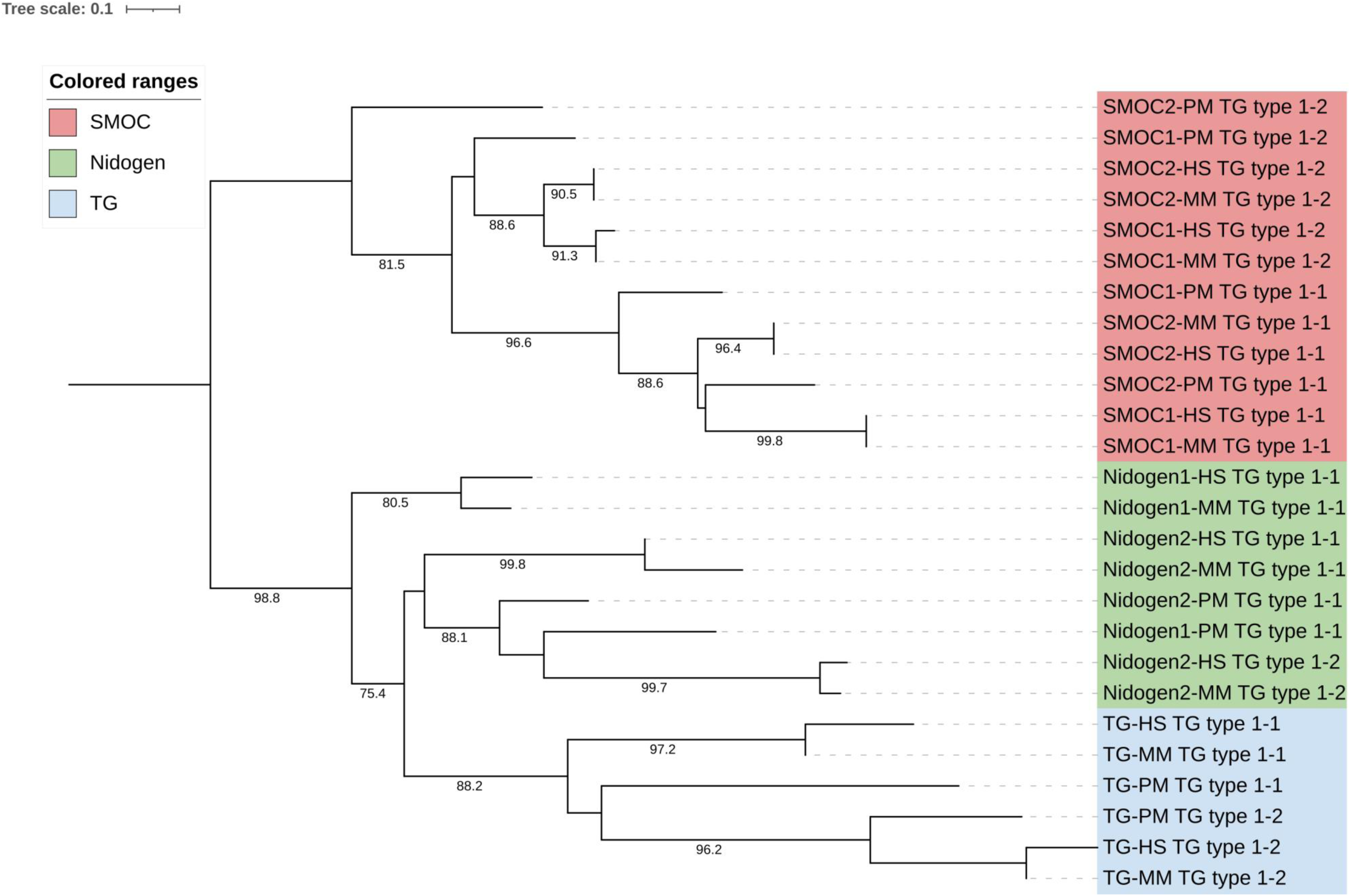
Phylogenetic analysis of TG type 1-1 and TG type 1-2 modules of SMOC1, SMOC2, nidogen1, nidogen2, and full-length thyroglobulin from *Petromyzon marinus*, *Mus musculus*, and *Homo sapiens*. The Maximum likelihood phylogeny, constructed from the alignment of TG type 1 amino acid sequences, depicts the evolutionary relationships between the both repetitive modules across the three taxa. Ultrafast bootstrap values ≥70, based on 2,000 replicates, are shown above the nodes. *Homo sapiens,* HS; *Mus musculus*, MM; *Petromyzon marinus,* PM.

Nidogen1 and nidogen2 are homologous sulfated monomeric glycoproteins found in the basal lamina, already observed in vertebrate and urochordate [Yurchenco & Wadsworth, 2004; Novinec et al., 2006; Holland and Short, 2010; Takagi et al., 2022]. In mammals, both proteins exhibit a modular structure consisting of three globular regions (G1, G2 and G3), separated by a a short flexible linker region between G1 and G2 and a rod-like domain between G2 and G3 [Bechtel et al 2012, Töpfer & Holz, 2024] (Figure 10a). The G1 globular region contains the NIDO domain (nidogen-like domain), while the G2 globular region consists of the nidogen G2 beta-barrel domain and one EGF-like domain (epidermal growth factor-like domain). The rod-like domain is composed of four EGF-like domains and one TG type 1 domain in nidogen1 (Uniprot: P14543, human) or two TG type 1 domain in nidogen2 (Uniprot: Q14112, human) of mammals (Figure 10a). However, sea lamprey nidogen2, which is related to a primitive nidogen, contains only a single TG type 1 module (Uniprot: A0AAJ7X4A4). Notably, nidogen of invertebrates such as *Drosophila melanogaster* and *Caenorhabditis elegans* do not possess TG type 1 domains. In nidogen1, the G3 globular region consists of four LDL-receptor class B repeats and an additional EGF-like domain, whereas in nidogen2, it contains five LY modules [Bechtel et al 2012, Töpfer & Holz, 2024]. Figure 10a presents the three-dimensional atomic structure of sea lamprey nidogen2 (which contains only one TG type 1 module) and human nidogen2 (which possesses two TG type 1 modules). These structures were generated using with AlphaFold server, as their crystallography remains unresolved. The SignalP 6.0 program predicts that nidogen2 sequence in both sea lamprey and human contains a signal peptide, with a cleavage site between positions 36 and 37 (MGMSRAVARARSPAGYAALPAALLCLAAALLGCARG) and a probability of 0.95 in sea lamprey. In human nidogen2, the cleavage site is predicted between positions 30 and 31 (MEGDRVAGRPVLSSLPVLLLLPLLMLRAAA) with a probability of 0.94 (data not shown).

**Figure 10.**
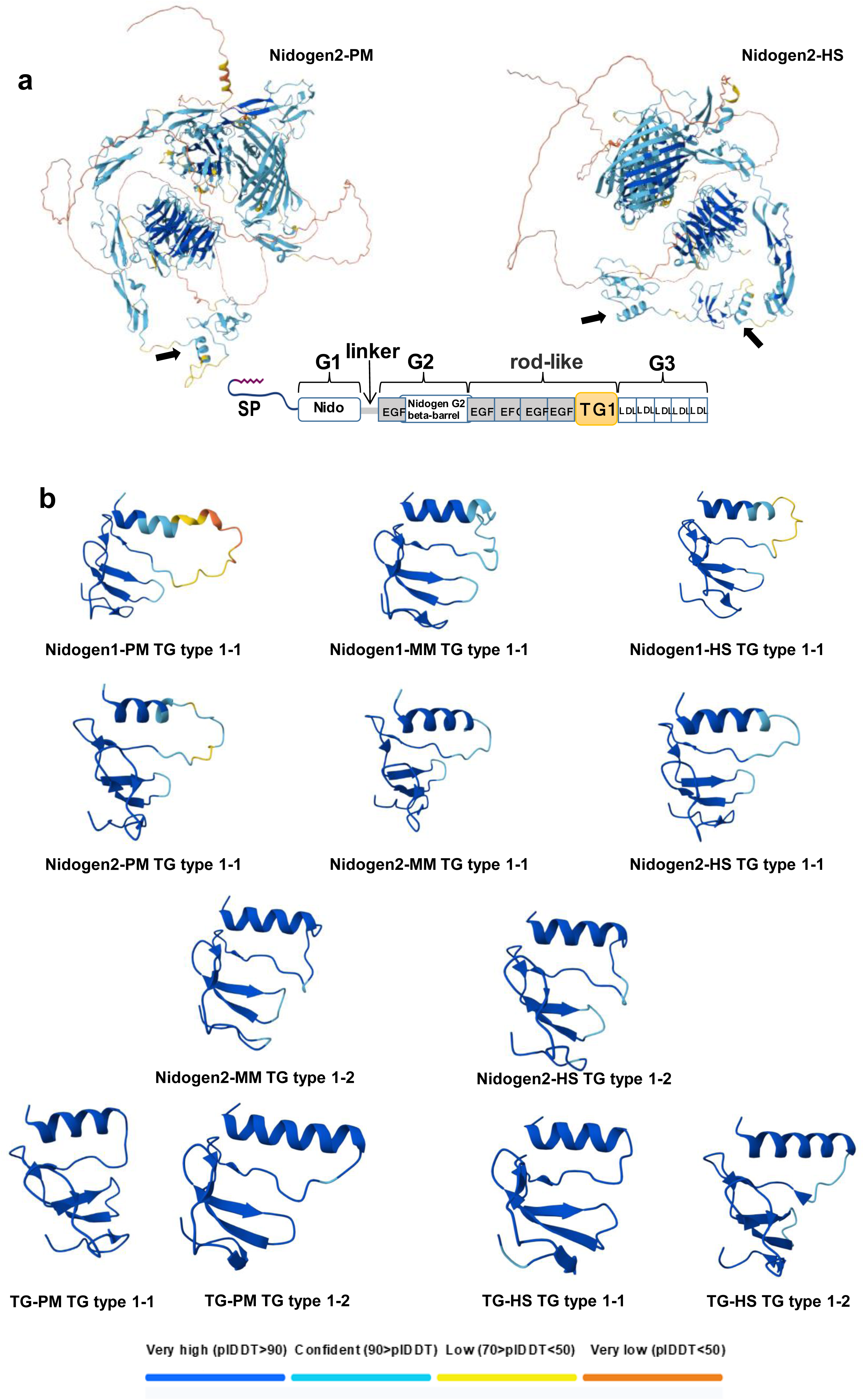
Three-dimensional atomic structure of nidogen2, and TG type 1 modules using AlphaFold server. **(a)** Predicted **3D structure** of nidogen2 from *Petromyzon marinus* and *Homo sapiens*. Arrows indicate the **TG type 1 modules**. The classical model of nidogen’s primary structure is shown. **(b)** M**odeled structure** of **TG type 1-1 and TG type 1-2 modules** of n**idogen1, nidogen2, and thyroglobulin (TG)** from *Petromyzon marinus* (**PM**), *Mus musculus* (**MM**), and *Homo sapiens* (**HS**). The pLDDT (predicted Local Distance Difference Test) score offers a per-atom confidence estimate ranging from 0 to 100, with higher values reflecting greater structural reliability. Low pLDDT scores often indicate intrinsically disordered or highly flexible regions.

To analyze the homology profile among the of TG type 1-1 and 1-2 modules of *Petromyzon marinus*, *Mus musculus* and *Homo sapiens*, including nidogen1, nidogen2, TGPM and TGHS, we performed the 3D structure modeling using AlphaFold server and an alignment using the T-Coffee program. AlphaFold modeling of TG type 1-1 and TG type 1-2 modules from nidogens and TGs in sea lamprey, mouse, and human revealed similar secondary structures, characterized by one α-helix and two β-sheets, with a very high plDDT score (Figure 10b). Additionally, the T-Coffee analysis yielded a high total consistency value of 90 points, indicating strong sequence conservation (Figure 11a). Remarkably, the six cysteine residues were well conserved across all TG type 1-1 and 1-2 repeats analyzed. Figure 11b illustrates the alignment of all TG type 1 modules around the six conserved cysteines. Notably, the structural conservation of these modules is high in both nidogens and TGs.

**Figure 11.**
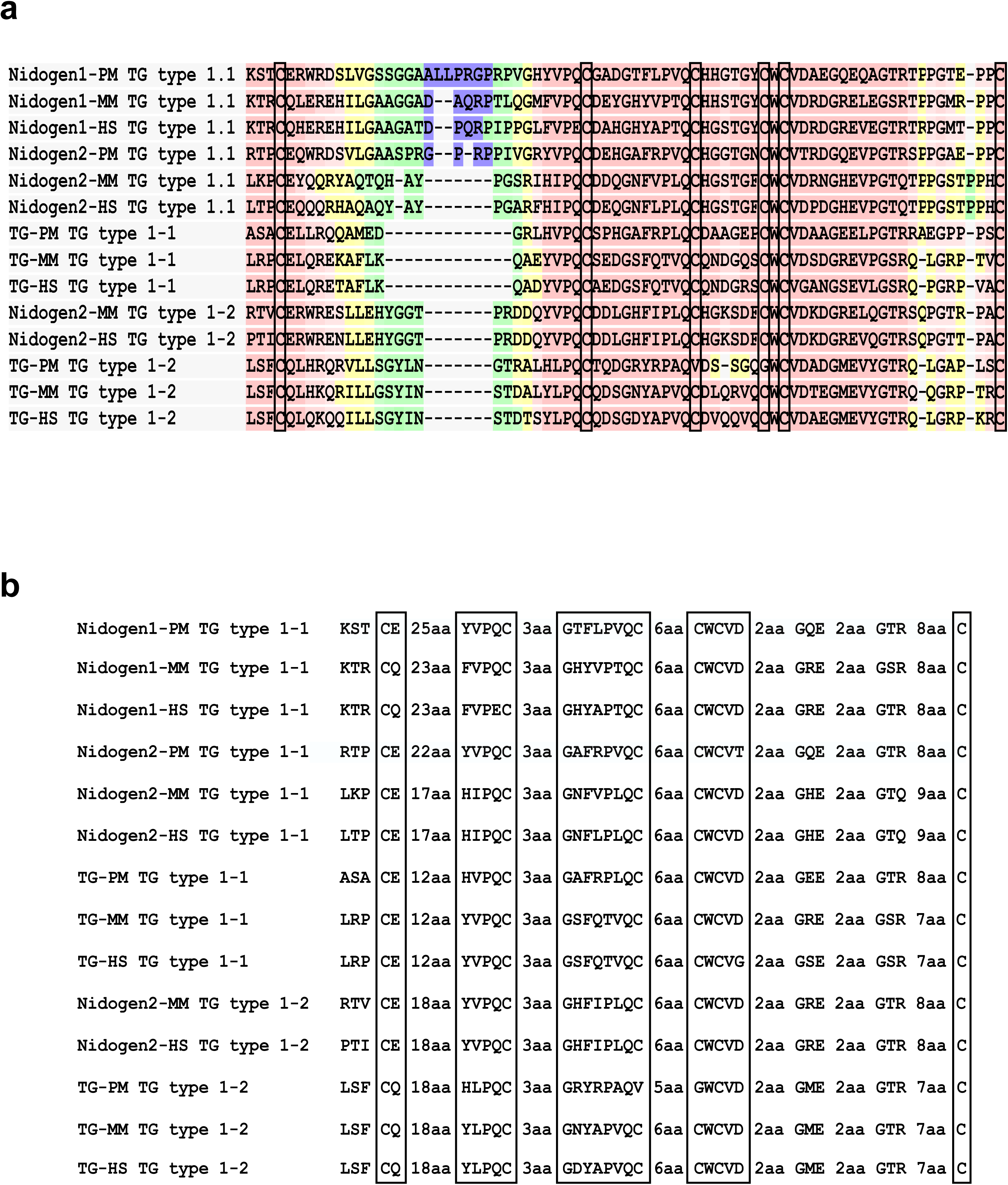
Comparative alignment of TG type 1-1 and TG type 1-2 modules of nidogen1, nidogen2, and full-length thyroglobulin from *Petromyzon marinus*, *Mus musculus*, and *Homo sapiens*. a) The TG type 1-1 and 1-2 modules were aligned using the T-Coffee program. Cysteine positions are boxed. BAD AVG GOOD b) The alignment of TG type 1-1 and TG type 1-2 modules was performed following the criteria established by Parma et al. (1987), Malthiéry et al. (1987), Molina et al. (1996), and van Graaf et al. (2001). The six conserved cysteine residues are present across all modules. Cysteine positions are boxed. Amino acids are represented using single-letter codes. *Homo sapiens,* HS; *Mus musculus*, MM; *Petromyzon marinus,* PM.

### Evolutionary model of the thyroglobulin complexation process

In our evolutionary model, we propose that a protein containing ancestral TG type 1 modules may have given rise to TG. As previously demonstrated, the TG type 1 domains of nidogen and TG cluster phylogenetically and are likely derived from a common precursor (Figure 9), here referred to as the nidogen-like ancestral precursor (Figure 12(a)-a). On one hand, this precursor gave rise to nidogen2 and, through the deletion of a TG type 1 module, to nidogen1 (Figure 12(a)-a). In addition to vertebrate clades, nidogen orthologs have also been identified in the urochordate sea squirt (*Ciona intestinalis*) [Novinec et al., 2006]. On the other hand, a later evolutionary period, the same nidogen-like ancestral precursor may have generated the initial TG type 1 repeats, leading to the emergence of a proto-TG with a limited number of such modules—subsequent to the origin of nidogens—as inferred from the observation that TG, unlike nidogen, is found exclusively in vertebrates [Novinec et al., 2006]. In contrast to nidogen1 and nidogen2, TG lacks the three globular domains (G1, G2, and G3) and the associated modules (NIDO, nidogen G2 beta-barrel, EGF-like, class B LDL receptor) that characterize nidogens. We propose two plausible mechanisms for the emergence of the first TG type 1 module: (i) a simple duplication of an existing TG type 1 module in the precursor, or (ii) a complete duplication of its precursor followed by the deletion of flanking upstream and downstream domains surrounding the TG type 1 module (Figure 12(a)-a). This process may have been facilitated by a genomic environment conducive to rearrangement. Among the most significant catalysts of such events is ionizing radiation, particularly ultraviolet (UV) radiation, given that atmospheric oxygen levels during the Early Cambrian are estimated to have been only ∼10% of present-day levels [Sperling et al., 2015], and the ozone layer was not yet fully formed. Consequently, the first secretory proto-TG likely emerged, containing three or four TG type 1 modules (Figure 12(a)-b). In addition to these motifs, the protein appears to have undergone further evolutionary modifications, including the acquisition of a signal peptide and specific tyrosine residues. These residues enabled iodine organization, allowing T_4_ synthesis via interaction between an acceptor and a donor tyrosine (Figure 12(a)-c). It is plausible that a short N-terminal segment of TG, harboring a single hormonogenic site and exhibiting a secretory profile, was sufficient to sustain primitive vertebrates on land by exerting the physiological effects of TH. This hypothesis is supported by extensive evidence of a mild hypothyroid phenotype in patients homozygous for the p.Arg296Ter variant, which retains only three TG type-1 motifs [van de Graaf et al., 1999; Rivolta et al., 2005; Caputo et al., 2007; Pardo et al., 2009; Citterio et al., 2013; Siffo et al., 2018; Zou et al., 2018]. Additional support comes from the p.A340Ter mutant, which retains four TG type 1 motifs and displays an active secretory profile, as reported by Lee et al. [2011].

**Figure 12.**
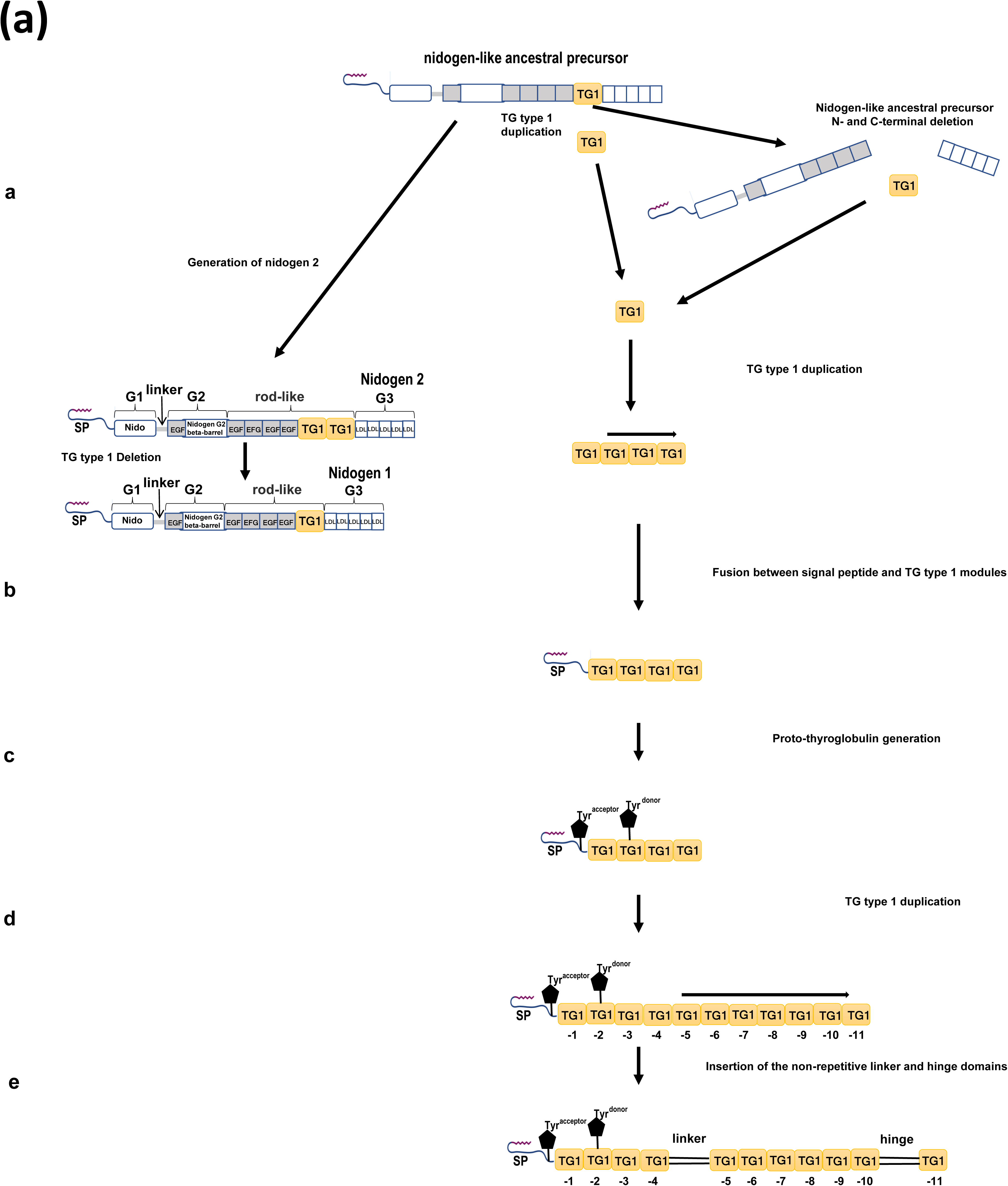
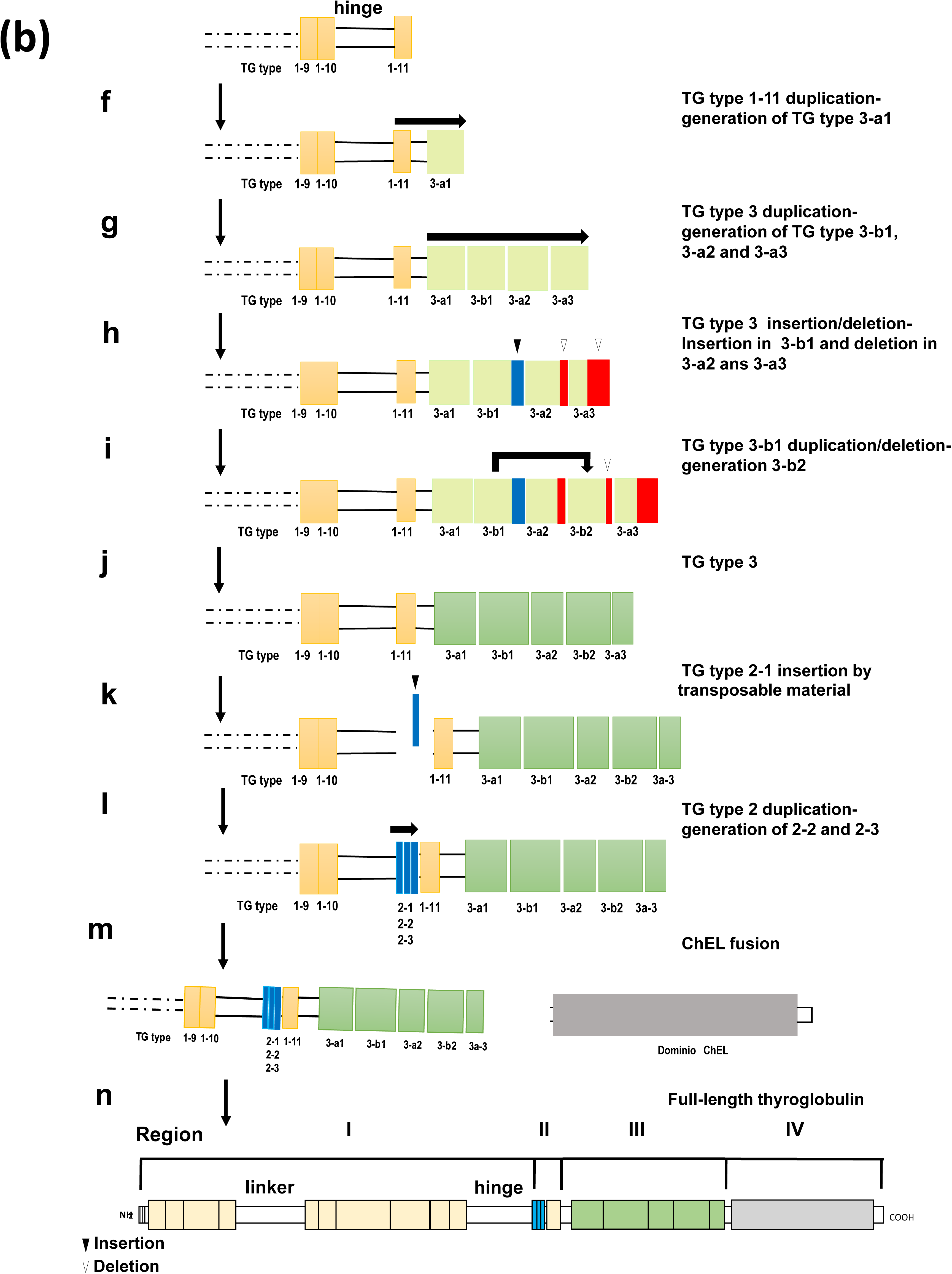
Flow chart of thyroglobulin complexation. **(a)** Schematic representation of thyroglobulin complexation, tracing its progression from the nidogen-like ancestral precursor to the acquisition of eleven TG type 1 modules and the insertion of non-repetitive linker and hinge domains. **(b)** Progression of thyroglobulin complexation from the acquisition of five TG type 3 modules and three TG type 2 modules to the fusion of the cholinesterase-like (ChEL) domain. The four canonical regions (I-IV) of the classical model of full-length thyroglobulin are shown at the bottom.

At a certain point, the proto-TG protein likely attained a stable structural and functional configuration. However, under evolutionary pressure, it underwent internal gene duplication events within the TG locus, resulting in an intragenic expansion of TG type 1 domains—ultimately producing a protein with 11 repeats— during the evolutionary remodeling that led to TG specialization in vertebrates (Figure 12(a)-d).

Insertions of genetic material following TG type 1-4 and 1-10 modules are hypothesized to have contributed to the formation of the non-repetitive linker and hinge domains, respectively (Figure 12(a)-e). These modifications represent an additional evolutionary milestone in the development of TG, progressively refining its architecture toward the definitive form observed in *Petromyzon marinus*, a modern representative of a lineage positioned at the base of vertebrate evolution.

Our structural analysis suggests a relationship between the TG type 1 and TG type 3 modules, as both contain of six cysteines and exhibit similar philogenetic clustering patterns (Supplementary Figure S11, S13, S16). BLAST analysis (NCBI) revealed no detectable homology with any genomic sequences apart from the TG Type 3 domain itself. These findings support the hypothesis that TG type 3-a1 module may have originated from the duplication of the TG type 1-11 repeat (Figure 12(b)-f). Through tandem duplications, the TG type 3-a1 module gave rise to the TG type 3-b1, TG type 3-a2 and TG type 3-a3 repeats (Figure 12(b)-g). Subsequent gene modulations involving partial deletions led to the differentiation of TG type 3-a2 and TG type 3-a3 from TG type 3-a1, while TG type 3-b1 underwent an insertion event and acquired two additional cysteines (Figure 12(b)-h). The structural similarity between the TG type 3-b1 and TG type 3-b2 modules is attributable to the conservation of eight cysteines in both. This suggests that the TG type 3-b2 module emerged via duplication of TG type 3-b1, followed by its insertion between TG type 3-a2 and TG type 3-a3 (Figure 12(b)-i). TG type 3-b2 subsequently underwent a deletion event that reduced its length— for instance, to 140 amino acids in sea lamprey TG (Figure 2c), compared to 172 amino acids in TG type 3-b1 and 134 out of 169 amino acids in human TG (Figure 2c). Thus, TG incorporates five modules that constitute its region III (Figure 12(b)-j). These evolutionary processes significantly reduced sequence conservation within region III, likely due to moderate evolutionary pressure—unlike the TG type 1 repetitive modules, whose antiproteolytic activity is essential for protein stability and preservation.

The TG Type 2 modules likely originated from the insertion of a transposable DNA fragment between the hinge and TG Type 1-11 domains (Figure 12(b)-k), followed by two duplication events (Figure 12b-l). To investigate the origins of these modules, the TG type 2-1, TG type 2-2 and TG type 2-3 modules were individually compared against the human genome and other organisms using the BLAST tool (NCBI). The analysis revealed no detectable homology with any genomic sequences other than the TG Type 2 domain itself. However, Lee et al. [2011] and Belkadi et al. [2012] classified this domain as a distant member of the GCC2/GCC3 domain superfamily. The higher conservation of TG Type 2 modules relative to TG type 3 modules**—**evidenced by the 38-species alignment (Table 1)**—**supports the hypothesis that the TG Type 2 region emerged after the formation of TG type 3 modules.

We hypothesize that the final—and perhaps most consequential —structural modulation event in TG was the incorporation of the ChEL domain and its fusion with regions I, II, and III (Figure 12(b)-m and n). The ChEL domain acts as an intramolecular chaperone, enhancing TG’s secretory efficiency [Lee et al., 2008]. Its integration not only facilitates TG secretion but also improves the protein’s capacity to elevate circulating T4 levels. Moreover, the ChEL domain is essential for TG dimerization [Lee et al., 2009]. In addition, TG has acquired the C-terminal hormonogenic site required for T_3_ synthesis. The secretion of TH in its active form, T_3_, together with elevated T_4_ levels, likely amplified the physiological functions mediated by TH. As previously noted, the TG of the *Petromyzon marinus* already exhibits a fully established complexation process.

## Discussion

In this study, we present the first complete sequence of the sea lamprey TG protein, a comprehensive analysis of TG domain conservation across major vertebrate clades, and a structural model of their complexation. A PDB model of *Petromyzon marinus* TG was generated and aligned with human TG, revealing consistent three-dimensional structural overlap across the four canonical regions (I–IV), thereby highlighting their evolutionary conservation. Additionally, a second lamprey TG transcript, TGPM^1746^ (Figure 2), was identified and characterized. This transcript initiates with the TG type 1-10 module and extends to the C-terminal end, exhibiting high homology with TGPM except in region III, where the TG type 3 repeats display notable variability. In spacer 3, the repeats resemble TG type 3b modules. This transcript was likely generated via alternative splicing from a novel promoter. Although the primary structure of TG diverges among the 38 species compared, all domains—including the repetitive (TG type 1, TG type 2, and TG type 3) and non-repetitive (linker, hinge, and ChEL) regions, as well as the N- and C-terminal hormone-related sites—remain strictly conserved across all vertebrate clades. (Figure 1, Supplementary Figure S10).

The structural organization of TG exemplifies genetic evolution through intragenic duplication events involving TG type 1, TG type 2, and TG type 3 modules, as well as gene fusion, as observed with the ChEL domain. Previous studies have established a relationship between the three families of cysteine-rich repetitive units and the intron–exon junction organization of the human TG gene [Mendive et al., 2001]. Detailed analysis of the repeat architecture revealed the following distribution: (i) TG type 1 repeats -2, -4, -7, -10, and -11 are each encoded by a single exon (exons 4, 8, 10, 16, and 22, respectively); repeats -1 and -9 span two exons (exons 2–3 and 14–15, respectively); repeats -3 and -8 span three exons (exons 5–7 and 11–13, respectively); and repeats -5 and -6 correspond to segments of exon 9. (ii) The three TG type 2 repeats are located between exons 20 and 21. (iii) The TG type 3 domain comprises two subtypes, 3a and 3b, distributed across exons 23 to 37 as follows: 3a-1 (exons 23–26), 3b-1 (exons 26–30), 3a-2 (exons 30–33), 3b-2 (exons 33–36), and 3a-3 (exons 36–37).

Protein diversity arises through a series of molecular processes—including mutation, fixation, and selection—which may be neutral, deleterious, or advantageous. Domain duplication, gene fusion, and recombination are fundamental evolutionary mechanisms that shape proteins and contribute to proteome complexity [Vogel et al., 2005]. Protein evolution is influenced not only by selective pressures on structure and function, but also by the genomic context of coding sequences, their expression profiles, their integration within biological networks, and their resistance to translation errors [Pál et al., 2006]. Many proteins are composed of domain combinations derived from diverse evolutionary origins, enabling both the modification of ancestral proteins and the emergence of novel architectures. Over time, these mechanisms have driven the development of new biological functions, thereby increasing the complexity of multicellular organisms [Patthy, 2003; Vogel et al., 2004]. Interestingly, domain combinations within proteins can be conceptualized as networks, where nodes represent functional domains from distinct superfamilies. Although the number of potential domain combinations is virtually limitless, only a restricted subset persists, shaped by strong evolutionary constraints. Notably, the reduction in protein diversity appears to correlate more closely with the rate of protein synthesis than with overall protein abundance [Pál et al., 2006]. In addition, radiation—particularly UV radiation—may act as a non-biological catalyst, inducing cycles of DNA damage and repair via homologous recombination. This process likely accelerated the rapid emergence and increasing complexity of genomes, facilitating the formation of intricate proteins such as TG within a relatively short evolutionary timeframe. All modulatory events proposed in our model occurred stochastically, within a brief evolutionary window, governed by the fundamental forces of natural selection and genetic drift. Radiation, acting as a persistent physical stressor on DNA, may have served as a catalyst, further accelerating these evolutionary dynamics.

We propose that the TG type 1 modules and the ChEL domain emerged at distinct stages of TG evolution, shaped by differing environmental pressures. TG type 1 modules likely originated earlier, during more primitive and environmentally stressful periods, which promoted a higher frequency of modulatory events and sequence variability. In contrast, the ChEL domain and its fusion with proto-TG appear to have arisen during a phase of reduced genomic rearrangement and lower environmental stress. The increasing structural complexity of genetic traits enabled vertebrates to colonize terrestrial environments and ultimately become the dominant lineage on Earth. The fusion of TG with the ChEL domain may represent a pivotal evolutionary milestone, contributing to the functional sophistication required for vertebrate diversification. The phylogenetic clustering identified in our analysis provides insight into the evolutionary origins of TG type 1 domains. The consistent co-clustering of nidogen-derived TG type 1 repeats with TG type 1 domains from TG within a single clade suggests a shared ancestral origin (Figure 9, Figure 12a). The presence of homologous TG type 1 repeats in both TG and nidogen—extracellular proteins that are structurally and functionally distinct—further supports the hypothesis that TG type 1 domains may have evolved from an ancestral nidogen-like protein. Following this divergence, the TG gene likely underwent intragenic duplication events, resulting in an expanded repertoire of TG type 1 repeats and the emergence of the modular architecture that characterizes modern TG proteins (Figure 12(a)). This evolutionary trajectory is consistent with the broader model of modular protein evolution—particularly among extracellular matrix components—where domain shuffling, duplication, and specialization have driven functional diversification across protein families. The retention of conserved cysteine-rich motifs within TG type 1 domains suggests strong selective pressure to preserve structural elements critical for protein stability and molecular interactions. Collectively, these findings support a model in which nidogen-like proteins served as evolutionary scaffolds for the origin of TG type 1 domains, followed by lineage-specific expansions that account for the diversity observed among vertebrate TGs.

The TG type 1 module predates the emergence of TG in early vertebrates and is found in several extracellular proteins, suggesting its likely involvement in the evolution of multicellular organisms by facilitating cell–cell and cell–environment interactions [Novinec et al., 2006]. A total of 170 protein sequences have been identified containing 333 TG type 1 modules [Novinec et al., 2006]. Several proteins harboring TG type 1 domains—such as nidogen, which we propose as a precursor to TG type 1 modules—play essential physiological roles [Fowlkes et al., 1997], operating across diverse tissue types. SMOC proteins are primarily localized at the basement membrane, where they contribute to calcium-binding regulation [Vannahme et al., 2002a; 2002b]. Nidogen plays a pivotal role in maintaining the three-dimensional architecture of the basal lamina and is involved in cell adhesion and nervous system development [Bechtel et al., 2012; Töpfer & Holz, 2024]. TROP proteins contribute to cell adhesion, migration, and placental development, and function as calcium signal transducers. Their overexpression in tumors has been associated with enhanced proliferative and invasive potential, positioning them as relevant targets in cancer biology [Li & Chen, 2025]. Testican proteins regulate cell adhesion and modulate the activity of cysteine proteases and metalloproteases [Bocock et al., 2003; Marr et al., 2003]. IGFBPs belong to a family of seven proteins characterized by high affinity for IGFs [Rajaram et al., 1997]. The invariant chain is essential for the proper assembly of MHC class II molecules [Holst et al., 2008]. Remarkably, in all these proteins, TG type 1 modules represent just one among several functional domains that compose their structure. In contrast, in TG, the repetitive modules constitute the majority of its architecture, which is complemented by the ChEL functional domain and the linker and hinge segments—whose functions remain unknown.

The evolutionary success of the TG type 1 module is likely due to its capacity for reversible protease binding, which provides protective advantages to proteins containing structurally resilient TG type 1 domains [Fineschi et al., 1995]. This property may have enabled such proteins to persist and function effectively in proteolytic environments during the emergence and establishment of multicellular life [Molina, 1996; Novinec et al., 2006]. In multicellular organisms, close cellular interactions generate microenvironments that are more exposed to proteolytic enzymes released by neighboring cells as part of normal metabolic activity. Notably, the release of TH involves the trafficking of TG to lysosomes [Ericson, 1981; Rousset and Mornex, 1991]. TG degradation is a slow process, requiring several hours [Brix & Herzog, 1994], and is mediated by proteolytic cleavage via thyroid-specific proteases [Dunn, 1991]. The eleven TG type 1 repeats may regulate this process in thyrocytes by selectively and reversibly inhibiting endosomal and/or lysosomal proteases, thereby playing a critical role in the physiology of TH release [Molina, 1996]. Interestingly, insertions within the TG type 1-3, TG type 1-7, and TG type 1-8 modules are highly conserved across all vertebrate TG sequences analyzed (Figures 1 and 2; Supplementary Figure S10), suggesting a shared evolutionary origin. However, in four Actinopterygii species—*Oryzias latipes*, *Stegastes partitus*, *Xiphophorus maculatus*, and *Cynoglossus semilaevis*—the TG type 1-7 insertion is largely absent from this module, indicating lineage-specific loss or divergence. These insertions likely originated from intron exonization events—as proposed by Parma et al. [1987] for the central insertion in TG type 1-7—or alternatively through random DNA insertion mechanisms. While both scenarios remain plausible, it is also likely that gene rearrangements and recombination–repair processes contributed to the emergence of this insertion pattern, acting in concert with tandem duplication and gene modulation events that shaped the structural evolution of these modules over time. These findings support the hypothesis that TG type 1 repeats underwent extensive intragenic rearrangements, retaining only those structural elements essential for their physiological function—most notably, the six conserved cysteines required for disulfide bond formation. This strongly suggests that the observed insertions originated during, or even prior to, the emergence of lampreys and have remained conserved across vertebrate lineages.

Proper protein function depends on the maintenance of a stable three-dimensional structure, which ensures the availability of sufficient active molecules within the cell [Pál et al., 2006]. Each TG monomer contains 122 cysteine residues, representing approximately 4.44% of its total amino acid composition [Mercken et al., 1985; Malthiery and Lissitzky, 1987; van de Graaf et al., 2001; Holzer et al., 2016; Citterio et al., 2021]. During biosynthesis, TG undergoes folding through the formation of up to 60 intrachain disulfide bonds, which contribute to its structural integrity and functional stability. As previously noted, TG type 1, type 2, and type 3 domains are cysteine-rich repeat modules covalently linked by intrachain disulfide bonds, which play a critical role in shaping the tertiary structure of TG. In both human and bovine TG, type 1 domains are further classified into two subgroups: type 1A and type 1B. Type 1A repeats (TG type 1–1 to TG type 1–8 and TG type 1–10) contain six cysteine residues, whereas type 1B repeats (TG type 1–9 and TG type 1–11) possess only four [Molina et al., 1996]. Conserved intradomain disulfide bonding patterns—Cys1–Cys2, Cys3–Cys4, and Cys5–Cys6—are consistently observed across all type 1A repeats [Molina et al., 1996]. In addition, TG type 1B, TG type 2, and TG type 3 repeats, as well as the six cysteine residues of the ChEL domain, also form intrachain disulfide bonds, contributing to the structural integrity of the TG molecule. Remarkably, over 95% of the cysteine residues in TG are conserved, underscoring their essential role in maintaining the protein’s structural and functional integrity (Supplementary Figure S10). As a result, TG is predominantly a well-structured protein, with the exception of a conserved disordered region at its C-terminus (Figure 7; Supplementary Figure S18-S23). This region encompasses the terminal segment of the ChEL domain and the hormonogenic site responsible for T_3_ formation. When isolated, this segment adopts a highly flexible conformational ensemble rather than a single defined structure.

Our analysis highlights both the overall conservation of TG tyrosine residues across vertebrates and a phylogenetic gradient of preservation, with mammals exhibiting the strongest selective constraints and more basal lineages showing progressively greater divergence (Figure 6). Notably, the Actinopterygii clade displayed a broad interquartile range, reflecting substantial variability in tyrosine conservation among ray-finned fishes. While some species exhibit relatively high levels of tyrosine conservation, others display markedly reduced preservation, highlighting divergent evolutionary trajectories within this group. This pattern contrasts with Mammalia, where a narrow distribution reflects consistently strong selective constraints. Hyperoartia, as a basal vertebrate lineage, shows pronounced evolutionary divergence from tetrapods and even from ray-finned fishes (Actinopterygii), consistent with its early branching position in vertebrate phylogeny. The consistently low conservation of tyrosine residues in TG suggests that, in lampreys, both the structure and function of TG have already undergone substantial modifications—potentially linked to the early evolution of the thyroid axis in primitive vertebrates. In contrast, Actinopterygii exhibit marked heterogeneity, with certain species approaching the conservation levels observed in tetrapods. Hyperoartia, however, is distinguished by its uniformly pronounced divergence, reflecting its basal position in vertebrate phylogeny.

Our study yields five key conclusions that collectively illuminate the evolutionary and structural dynamics of TG across vertebrates. First, we successfully identified and characterized the complete sequence of the sea lamprey TG protein, offering valuable insights into its structural conservation and evolutionary trajectory. In addition, we detected a second TG transcript in this species. Second, comparative analysis across 38 representative species—including mammals, birds, reptiles, amphibians, ray-finned fishes, and jawless vertebrates—demonstrates that all TG domains are conserved throughout vertebrate evolution, underscoring their functional indispensability. Third, despite substantial divergence in overall amino acid sequences, tyrosine residues and the cysteines responsible for disulfide bond formation—both critical for TG function—remain highly conserved, highlighting strong selective pressures on these molecular features. Fourth, TG is revealed to be a highly structured multidomain protein, characterized by extensive intrachain disulfide bonding and a conserved disordered segment at its C-terminus. Finally, based on our findings, we propose a stepwise model for TG complexation. This model begins with the emergence of TG type 1 modules from a nidogen-like precursor, followed by intragenic duplication events and the sequential incorporation of TG type 3 and TG type 2 modules. The process culminates in fusion with the ChEL domain, completing the architectural assembly of TG. These results provide compelling evidence that the TG complexation process is fully established in lamprey—consistent with its basal phylogenetic position—and has remained largely conserved throughout vertebrate evolution.

## Methods

### Amino acid sequence alignments using Clustal Omega, T-Coffee, EMBOSS Needle and Blat Search Genome

The TG amino acid sequence homology between multiple sequence were performed using the multiple sequence alignments Clustal Omega with its visualization tool MView (https://www.ebi.ac.uk/Tools/msa/clustalo/) and T-Coffee, (https://tcoffee.crg.eu/). T-Coffee is a versatile multiple sequence alignment tool suitable for aligning various types of biological sequences, including nucleic acids and proteins. More than just an aligner, T-Coffee acts as a framework that integrates different alignment methods with additional structural, evolutionary, or experimental data to enhance accuracy and meaningful alignments. It employs a library generated from a combination of pairwise alignments to construct multiple alignments. The coloured graphical output visually represents the level of consistency between the final alignment and the library used by T-Coffee, where the main score—the overall consistency value—ranges from 0 to 100. A score of 100 indicates complete agreement between the alignment and its reference library.

The alignment file uses a color scheme to denote consistency, from blue (low-consistency regions), which are seldom biologically meaningful, to red (high-consistency regions), which reflect strong agreement.

The amino acid sequence homology between sequence pairs was assessed using EMBOSS Needle (https://www.ebi.ac.uk/Tools/psa/emboss_needle/) and Blat Search Genome (https://genome-euro.ucsc.edu/cgi-bin/hgBlat).

### SignalP 6.0: signal peptide prediction and cleavage site identification

The SignalP 6.0 server predicts the presence of signal peptides and determines their cleavage sites in proteins (https://services.healthtech.dtu.dk/services/SignalP-6.0/). It can distinguish between five types of signal peptides: (1) Sec/SPI, standard secretory signal peptides transported by the Sec translocon and cleaved by Signal Peptidase I; (2) Sec/SPII, lipoprotein signal peptides transported by the Sec translocon and cleaved by Signal Peptidase II; (3) Tat/SPI, Tat signal peptides transported by the Tat translocon and cleaved by Signal Peptidase I; (4) Tat/SPII, Tat lipoprotein signal peptides transported by the Tat translocon and cleaved by Signal Peptidase II and (5) Sec/SPIII, Pilin and pilin-like signal peptides transported by the Sec translocon and cleaved by Signal Peptidase III.

### Modeling the three-dimensional atomic structure of proteins

The three-dimensional atomic structure of *Homo sapiens* TG has been characterized using a composite cryo-electron microscopy (cryo-EM) density map (PDB: 6SCJ, resolution: 3.60 Å; https://www.rcsb.org/structure/6SCJ) [Coscia et al., 2020]. Coscia et al., [2020] proposes a new model of the primary structure of the human TG organized in five regions: N-Terminal Domain (NTD), Core, Flap, Arm and C-Terminal Domain (CTD). However, the TG models presented by Adaixo et al. [2022] and Kim et al. [2021] adhere to the general framework of the classical model for the TG monomer. Therefore, we adapted the Coscias et al., [2020] model align with the classical model. Spacers 1, 2, and 3 are located in regions II, III, and IV, respectively. The UCSF Chimera program was used to visualize the 3D model of the TG (UCSF Resource for Biocomputing, Visualization, and Informatics at the University of California, San Francisco, https://www.cgl.ucsf.edu/chimera/) [Pettersen et al., 2021; 2004].

The PDB for TGPM^2475^ (Uniprot: S4R814_PETMA), TGPM^1746^ (NCBI: XP_032817730.1) and full-lenght assembled TG (TGPM) from *Petromyzon marinus,* were generated using homology modeling with Swiss Model (https://swissmodel.expasy.org/interactive) due to the unresolved crystallographic data.

The three-dimensional structures of nidogen2 from *Petromyzon marinus* and *Homo sapiens*, as well as TG type 1-1 and TG type 1-2 modules from nidogen1, nidogen2, and TG from *Petromyzon marinus*, *Mus musculus*, and *Homo sapiens*, were generated using the AlphaFold server (https://alphafoldserver.com/). pLDDT (predicted Local Distance Difference Test) provides a per-atom confidence estimate on a 0-100 scale, where a higher value indicate greater reliability. Regions with low pLDDT scores may correspond to unstructured or highly flexible areas.

### Structural matching using the Matchmaker command in UCSF ChimeraX

Structural matching (structural superposition) of different PDBs was performed using the Matchmaker command in UCSF ChimeraX. This command first generates a pairwise sequence alignment and then refines the alignment by optimizing residue pairs at the atomic level. The RMSD (Root-Mean-Square Deviation) value in Matchmaker quantifies the average distance between atoms—typically backbone atoms—of superimposed molecules, measured in Ångströms (Å). In globular protein conformation studies, three-dimensional structural similarity is commonly assessed by calculating the RMSD of Cα atomic coordinates after optimal rigid-body superimposition. A cutoff value of 2 Å is used, where lower RMSD values indicate higher-quality superimposition, with values approaching 0 Å representing near-perfect alignment.

### Phylogeny

Phylogenetic reconstruction of the whole TG and and of each of the component domains were performed using amino acid sequences retrieved from the NCBI database (https://www.ncbi.nlm.nih.gov) for thirty-six species: *Alligator mississippiensis* (XP_059579449.1), *Aquarana catesbeiana* (XP_073488945.1), *Astyanax mexicanus* (XP_022529282.2), *Bos Taurus* (NP_776308.1), *Canis lupus familiaris* (NP_001041569.1), *Carassius auratus* (XP_026120244.1), *Cavia porcellus* (XP_003467392.1), *Chelonia mydas* (XP_043394997.1), *Clupea harengus* (XP_031429786.1), *Columba livia* (XP_021154537.2), *Crotalus tigris* (XP_039206379.1), *Cynoglossus semilaevis* (XP_008321228.1), *Cyprinus carpio* (XP_042628676.1), *Danio rerio* (NP_001316794.1), *Eublepharis macularius (*XP_054841882.1), *Gallus gallus* (NP_001376406.2), *Homo sapiens* (NP_003226.4), *Lampetra fluviatilis* (CAL5909345.1), *Lampetra planeri* (CAL5920418.1), *Larus michahellis* (XP_074433977.1), *Lepisosteus oculatus* (XP_015212882.2), *Macaca fascicularis* (XP_045254657.2), *Macaca mulatta* (XP_028708128.1), *Mus musculus* (NP_033401.2), *Oryzias latipes* (XP_011484169.1), *Pan troglodytes* (XP_016815373.4), *Panthera leo* (XP_042780307.1), *Pongo pygmaeus* (XP_054355108.2), *Python bivittatus* (XP_025021113.1), *Rattus norvegicus* (NP_112250.2), *Stegastes partitus* (XP_008304814.1), *Struthio camelus* (XP_068789869.1), *Taeniopygia guttata* (XP_072781126.1), *Trichechus manatus latirostris* (XP_004373071.1), *Xenopus tropicalis* (NP_001316486.1) and *Xiphophorus maculatus* (XP_023185937.1). The TG sequence for *Gorilla gorilla* was obtained from UniProt (https://www.uniprot.org; accession G3QS68), while the sequence for *Petromyzon marinus* was generated in the present study.

Additionally, two phylogenetic trees were constructed: one based on the eleven TG type-1 repeats of TG from three vertebrate species (*Homo sapiens*, *Petromyzon marinus*, and *Mus musculus*), and another based on TG type 1-1 and type 1-2 repeats derived from SMOC, nidogen and TG proteins. In addition to TG sequences, the latter analysis also incorporated sequences from *Homo sapiens* SMOC1 (NCBI: NP_001030024.1), SMOC2 (NCBI: NP_071421.1), nidogen1 (UniProt: P14543) and nidogen2 (UniProt: Q14112); *Petromyzon marinus* SMOC1 (NCBI: XP_032818876.1), SMOC2 (NCBI: XP_075927210.1), nidogen1 (UniProt: A0AAJ7TJ46) and nidogen2 (UniProt: A0AAJ7X4A4); and *Mus musculus* SMOC1 (NCBI: NP_001139689.1), and SMOC2 (NCBI: NP_071710.2), nidogen1 (UniProt: P10493) and nidogen2 (UniProt: O88322).

The sequences were aligned using PRANK tool [Löytynoja, 2014]. Maximum Likelihood (ML) phylogeny was reconstructed using IQ-TREE v1.6.12 [Nguyen et al., 2015], utilizing the best substitution model determined by ModelFinder [Kalyaanamoorthy et al., 2017], and with 2000 ultrafast bootstraps [Hoang et al., 2018]. The phylogenetic tree was visualized with iTOL [Letunic and Bork, 2016].

### Tyrosine conservation in human thyroglobulin

The TG sequence was obtained from NCBI (accession NP_003226.4). Orthologous sequences from 37 additional species were aligned to the human TG sequence using Clustal Omega. Alignment positions were tracked using both the alignment column number and the ungapped human TG residue position.

All alignment positions where the human sequence contains tyrosine were considered. For each such position, conservation was calculated as the number of species (including those with gaps) that contained tyrosine divided by the total number of species (38). Gaps were counted as valid species but non-conserved. Percentages were rounded to integers for presentation. Species were assigned to clades according to updated taxonomy: Mammalia, Archosauria (birds, Chelonia, and Alligator), Lepidosauria, Amphibia, Actinopterygii, and Hyperoartia. Analyses were performed using Python 3.11 with the following libraries: pandas (2.0), matplotlib (3.7), python-docx (0.8). Data were processed in tabular format and exported to csv and docx. Figures were generated using matplotlib.

### Protein disorder prediction

A protein disorder prediction for 38 full-length TG sequences was performed using the AIUPred web server [Erdős et al., 2025] with the “AIUPred-only disorder” setting (https://aiupred.elte.hu/). Residues with a predicted score ≥0.5 were classified as disordered, while those below this threshold were considered ordered.

## Supporting information

Supplementary Material-Evolution of thyroglobulin

Supplementary Video S1-Evolution of thyroglobulin

Supplementary Video S2-Evolution of thyroglobulin

Supplementary Video S3-Evolution of thyroglobulin

## Acknowledgements

M.G.P. is a research fellow of the Consejo Nacional de Investigaciones Científicas y Técnicas (CONICET). W.M.S., C.M.R. and H.M.T. are established investigators of the CONICET.

## Author contributions

**M.G.P.** contributed to the study design and bioinformatic analysis, and conducted the structural modeling. **W.M.S.** performed the phylogenetic analysis. **C.M.R.** participated in the bioinformatic analysis, contributed to the study design, and was involved in funding acquisition. **H.M.T.** carried out the bioinformatic alignment analysis, contributed to funding acquisition, conceived and designed the study, and wrote the manuscript. All authors critically reviewed the manuscript, contributed to its revision, and approved the final version.

## Funding

This study was funded by grants from the Fondo para la Investigación Científica y Tecnológica (FONCyT-ANPCyT-MINCyT, PICT-2018-02146 to H.M.T.), CONICET (PIP 2021-

11220200102976CO to C.M.R.) and Universidad de Buenos Aires (UBACyT 2020-20020190100050BA to C.M.R.).

## Data availability

Data and material are available from the authors upon request.

## Declarations

**Conflict of interest** The authors declare that they have no conflict of interest.

**Ethical approval** Not applicable. No procedures requiring ethics approval were performed in this study.

**Consent to participate** Not applicable.

**Consent to publish** Not applicable.

