## Supplementary Material-Evolution of thyroglobulin for "Evolution of thyroglobulin: an integrated view of its origin and complexity from a structural perspective"

[1] Universidad de Buenos Aires. Facultad de Farmacia y Bioquímica. Departamento de Microbiología, Inmunología, Biotecnología y Genética/ Cátedra de Genética. Buenos Aires, Argentina.

[2] CONICET-Universidad de Buenos Aires. Instituto de Inmunología, Genética y Metabolismo (INIGEM). Buenos Aires, Argentina.

[3] CONICET-Universidad de Buenos Aires. Instituto de Fisiología y Biofísica Bernardo Houssay (IFIBIO HOUSSAY). Buenos Aires, Argentina.

**Supplementary Table S1. Comparative analysis of tyrosine residue conservation across 38 vertebrate species, relative to human thyroglobulin.**

| Position of tyrosine in human thyroglobulin | Number of species with tyrosine residues at position | Conservation_ % |
|---|---|---|
| 24 | 38 | 100 |
| 48 | 24 | 63 |
| 108 | 28 | 74 |
| 116 | 24 | 63 |
| 126 | 21 | 55 |
| 149 | 38 | 100 |
| 211 | 20 | 53 |
| 234 | 38 | 100 |
| 258 | 33 | 87 |
| 277 | 34 | 89 |
| 315 | 26 | 68 |
| 325 | 31 | 82 |
| 376 | 10 | 26 |
| 382 | 18 | 47 |
| 626 | 33 | 87 |
| 704 | 13 | 34 |
| 759 | 33 | 87 |
| 785 | 29 | 76 |
| 809 | 29 | 76 |
| 825 | 28 | 74 |
| 839 | 28 | 74 |
| 866 | 21 | 55 |
| 879 | 34 | 89 |
| 883 | 29 | 76 |
| 992 | 27 | 71 |
| 1005 | 30 | 79 |
| 1027 | 35 | 92 |
| 1114 | 22 | 58 |
| 1165 | 26 | 68 |
| 1310 | 28 | 74 |
| 1445 | 10 | 26 |
| 1467 | 15 | 39 |
| 1481 | 35 | 92 |
| 1529 | 28 | 74 |
| 1630 | 36 | 95 |
| 1677 | 27 | 71 |
| 1705 | 33 | 87 |
| 1782 | 32 | 84 |
| 1819 | 33 | 87 |
| 1916 | 26 | 68 |
| 1922 | 36 | 95 |
| 1965 | 38 | 100 |
| 2058 | 20 | 53 |
| 2063 | 36 | 95 |
| 2157 | 35 | 92 |
| 2184 | 35 | 92 |
| 2194 | 6 | 16 |
| 2233 | 37 | 97 |
| 2283 | 36 | 95 |
| 2335 | 30 | 79 |
| 2478 | 23 | 61 |
| 2540 | 35 | 92 |
| 2563 | 20 | 53 |
| 2564 | 37 | 97 |
| 2573 | 35 | 92 |
| 2587 | 17 | 45 |
| 2611 | 37 | 97 |
| 2617 | 13 | 34 |
| 2637 | 21 | 55 |
| 2640 | 26 | 68 |
| 2658 | 35 | 92 |
| 2670 | 20 | 53 |
| 2672 | 12 | 32 |
| 2697 | 37 | 97 |
| 2721 | 34 | 89 |
| 2766 | 37 | 97 |

**a**

```
CELLRQQAMEDGRLHVPQCSPHGAFRPLQCDAAGEPCWCVDAAGEELPGTRRAEGPPPSC
LSFCQLHRQRVLLSGYLNGTRALHLPQCTQDGRYRPAQVDSSGQGWCVDADGMEVYGTRQ
LGAPLSCPSPCQVSVRRAVRASTPGSPPPQCDRDEQRVLATQCQLIGTAGRSTLSLLDAF
SMHPEAFRSLAEFRARLPGVHAYCYCADPQGRELPETGDRGVRSFLFVAYFSGHGLNHEV
VGSVLGKTRHEKFLKVQDRLPGHIVRPTVCEVERSLAERLGGRGFAPVCSQDGSYIPTQC
HGPTCWCVDTRGDEVFGSRVRDSVPNCGSSSSPCQLERQLALSHLFLGPSAPFIDDLISA
LPRDGSVSFSESPSQEMQHELWLHWRALREQLLDGGLGEILSQASRLFPASGQASRLLGD
LVGGIFHTQELALTAAGANVQPGRIAEMLFGDRFLKNLKFFNFTGAVGGRGTFNFSKVFG
QVGLTGMYDGANFEELARLFAPGEDSYLTRGSSNFSRESFNFDQSIDDPFARATNLERNR
NVLGFVVTLLEDARFYGLVRDITPLIGLSGGDVGLNALTFLRMAEEDRPTQNAADAADGP
PASRPFVARCGEDGGAYEPVQCHGAACWCVDAHGREIAGTRAVGGAPPRCPSACEGERAR
ALLARRGRPVGTPLFVPECDGAGDYRPVQCSGDRCFCVGAGGAEVPGTGRPLGDPVTCPT
PCQAAAASELLQQLRHLGPVLRGDAPSSLPPLYVPRCDARGGWRPVQCDGAGQQIREFVD
AWTKDEGNASKATLSDLRELLARVRGAGGGGGGGKGLRSFLSFLYDSGRQDMFPELSLY
PSLRGPPEVLNFSGSSGRFLRNPEVVWKILTENASFVYTGDYAEFSGRYADFESRLCWCV
DAAGHEIDGTRTVSPGQLPKCPGACHLAAADVNRYLQQADLLIGSAGASSAGEGVAFGRG
LAFTEDELLGSPLGLGDVKGEVAAVLAGGTGYAVRLAAQAAMHFHWRRRFFGPGTIGEGV
FNGFDPYMPQCTEPGGWEPAQCVVRAGAGFCWCVDAAGEFVAGSLVARPRRRPQCATPCQ
RARAEALLTGWKSLGSVENGSSIEVLTKHTPACTPSRKYKKCHYLLRPNWWPSPSYSYTP
THPSPTTNAESPSLCELRRRQALLRNAGLGSVPECDSRGDYAARQCADGTCWCAGAGGEE
IPDTRRHAGEPGVVCHEPRCELPQDERRAGRWALVCEDGAAATAAAGLQRCLLACLRGFA
HTLDGAGHDGGAPREFQCNATSGEWIGSSPQPDACQEIHPWQAVQLSISFTLRFPEEKMC
NPEYEGLLESFNSFLLADLTARGFCHLSNGVGSDAVAVCDAASVRVRCQDALRLLVNVTW
RSSLSRLPVQSYPDLHNIERAFVAHAQRFAQLLDDRGYRLALNGRDYSSEGAARFRKDEL
YDTSPAFRVACEPGYVRIAQGCAACPRGSLWRDGGCALCPADHYQDEAGQTACTPCPADT
GTLTAGAAAAFQCRTPCEREALRRGQLGAGERLELYCDPEGLYEGEKPAKGGQDEVRRMF
EEVPSEDLVLGAVDAVALRIRTETGPVQEVEPRCLQECLKEERCDFFVFSTQGDVSSCHF
YNGSRESYQCEKAPEEKSALGISLCVVFIAIKCFVRGVGFHVSSASGRFYFHAGHELPSW
DGQLYRPTGFGNAASGAYRTLALPAGTTTLPDAHLYCRSACSAQACCDGFLLKELPPGVL
VCALLSGPTVLTCRAPWWPRTADEYGDGECSGLRVHKPSRTFGFSLGGVHYNASNPPSAV
ASTAAAAAAASQVIYLWAGSEAGSRDVCLHTKAYTPQPGEPAGAADVSGRFSTLRPEEVT
LQPEMRVAQQNFWLFRRAFSPPQQATSWCLRRCRRDALCKAAAVGDGPAGWLECLLYPDT
QSCGTSQELLERPARAPSCASLLPRIDGTLYRKRDGAEGPVRRLYRRTAFRSEIGVLLRS
VANLTGMALSDGFSRCERTCDEDPCCEGFAFLRRALSSAGEALLQCATHTGPGVQLCGRE
SPMGATFGCPPPGVDTASSLFGWYRAQARSSPQTAQLCPAVTLPPAPTKGAADGFQSLDV
STAAVDPTISAFDVVVLLGPGAGGSPRAAPPSDEWCLAVCQRSPWCASVGVRHLGDSTRC
VLHAETLACGYGARGGRHCRLGVLEPPARLYRKKAPSPGGGQGPVRPVVPLGSGQLQGAS
RTVEVDSQVKVIDVFLGVPYAAPPLDANRFSGPQPPRPLAVNETWDADRYKTRPDCLQPD
GQRGTSLSEDCLYLNIFVPKVKPHNASVLVFFHGGDNSFGGSAQGPLDPSYLAALGDIIV
VTANYRLGLFGFLSTGDEAAAGNWGLLDQQAALRWVRDHAALFGGSAGAVTVAGDRSSAD
NVGLHLVAPGSRGLFQRAILMGGSVLSPSAVQLDSATARSQAASLAQLVGCGHSSSEYLV
RCLRDRSALALNAAQ
```

**b**

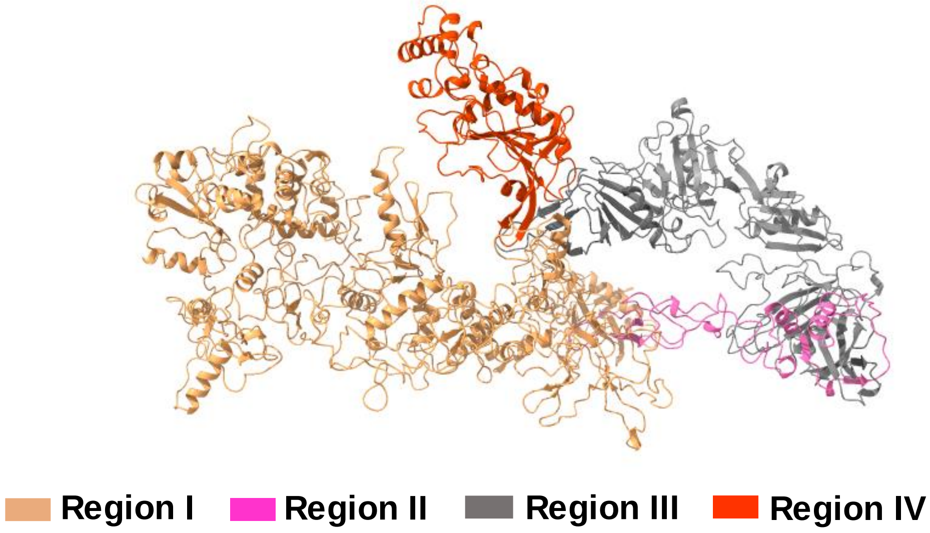


**Supplementary Figure S1. Sequence and three-dimensional atomic structure of the 2475-amino acids thyroglobulin from *Petromyson marinus* (TGPM$^{2475}$), indexed in UniProt (S4R814_PETMA). (a)** The TGPM$^{2475}$ sequence comprises eleven TG type 1 modules, three TG type 2 modules, and five TG type 3 modules, along with linker and hinge domains, spacers 1, 2 and 3, and the first 256 amino acids of the ChEL domain. Amino acids are represented using single-letter codes. **(b).** A homology-based 3D model of TGPM$^{2475}$ was generated using the Swiss-Model platform, with *Bos taurus* thyroglobulin (PDB: 7N4Y) as the structural template. The resulting PDB file was visualized using UCSF ChimeraX.

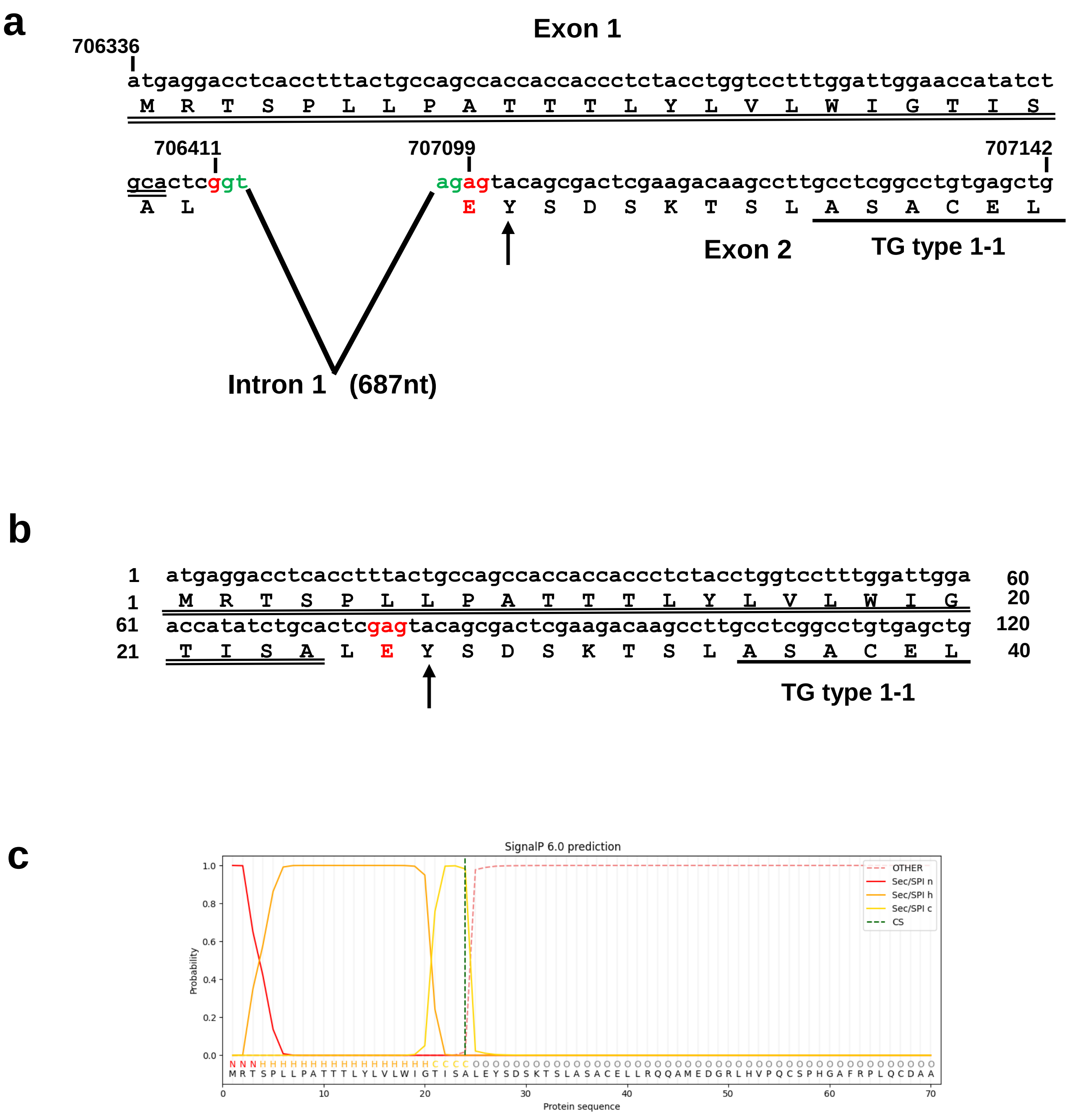


**Supplementary Figure S2. Identification of the amino-terminal end of full-length thyroglobulin from *Petromyson marinus* (TGPM). (a)** Genomic organization and deduced amino acid sequence. The genomic nucleotide and coding sequences spanning the exon 1/intron 1/exon 2 boundaries were obtained from scaffold GL476337 (ENSPMAG 00000001187, ENSPMAT00000001350.1), covering positions 706336 to 707142. **(b)** cDNA nucleotide and coding sequences were analyzed between positions 1 to 40. The nucleotide sequence is shown in the upper line, while the amino acid translation (represented using single-letter codes) appears below the second nucleotide of each codon. The canonical gt-ag splice consensus is highlighted in green, and the g\ag codon (glutamic acid residue) overlapping the exon junction is marked in red. The initial amino acid residues of the TG type 1-1 module are underlined. The amino-terminal hormonogenic acceptor site ($Y^{27}$) is indicated by an arrow, and the predicted signal peptide is double underlined. **(c)** Signal Peptide Prediction. The SignalP 6.0 program predicts that the exon 1 sequence of TGPM contains a signal peptide with a cleavage site (cs) between positions 24 and 25, with a probability of 0.98. The program identifies the following signal peptide regions: region n (amino-terminal), region h (central hydrophobic), and region c (C-terminal).

a

```
MPSRVRACVRAPAAATFLCSALPSPGPSLCELRRRQALLRNAGLGSVPECDSRGDYAARQ
CADGTCWCAGAGGEEIPDTRRHAGEPGVVCHEPRCELPQDERRAGRWALVCEDGTAATAA
TAAGLQRCLLACLRGFAHTLDGAGHDGGAPREFQCNATSGEWIGSSPQPDACQEIHPWQA
VQLSISFTLRFPEEKMCNPEYEGLLESFNSFLLADLTARGFCHLSNGAQSAPGGGSDAVA
VCDAASVRVRCQDALRLLVNVTWRSSLSRLPVQSYPDLHNIERAFVAHAQRFAQLLDDRG
YRLALNGRDYSSEGAARFRKDELYDTSPAFRVACEPGYVRIAQGCAACPRGSLWRDGGCA
LCPADHYQDEAGQTACTPCPADTGTLTAGAAAAFQCRTPCEREALRRGQLGAGERLELYC
DPEGLYEGEKPAKGGQDEVRRMFEEVPSEDLVLGAVDAVALRIRTETGPVQEVEPRCLQE
CLKEERCDFFVFSTQGDVSSCHFYNGSRESYQCEKAPEAPGFLGDPDSSWVERLSCRPRV
SLPPGSVFTVYRKKGHELPSWDGQLYRPTGFGNAASGAYRTLALPAGTTTLPDAHLYCRS
ACSAQACCDGFLLKELPLDNGVLVCALLSGPTVLTCRAPWWPRTADEYGDGECSGLRVHK
PSRTFGFSLGGVHYNATFKDLGAYEKDAGLPRASARPSSGPLEAGFTLDLQRGGALFSQI
YLWAGSEAGSRDVCLHTKAYTPQPGEPAGEEEEDVSYHSPQDEADVSGRFSTLRPEEVTL
QPEMRVAQQNFWLFRRAFSPPQQATSWCLRRCRRDALCKAAAVGDGPAGWLECLLYPDTQ
SCGTSQELLERPARAPSCASLLPRIDGTLYRKRDGAEGPVRRLYRRTAFRSEIGVLLRSV
ANLTGMALSDGFSRCERTCDEDPCCEGFAFLRRALSSGEALLQCATHTGPGVQLCGRESP
MGATFGCPPPGVDTASSLFGWYRAQARSSPQTAQLCPAVTLPPAPTKGAADGFQSLDVST
AVVDPTISAFDVVVLLGPGAGGSPRAAPPSDEWCLAACQRSPWCASVGVRHLGDSTRCVL
HAETLACGYGARGGRHCRLGVLEPPARLYRKKASLGTSCAARCGLYSPGQPCQCNRECQR
FGDCCPDAAVCLGGGQGPVRPVVPLGSGQLQGASRTVEVDSQVKVVDVFLGVPYAAPPLD
ANRFSGPQPPRPLAVNETWDADRYKPDCLQPDGQRGTSLSEDCLYLNIFVPKVKPHNASV
LVFFPGGDNSFGGSAQGPLDPSYLAALGDIIVVTANYRLGLFGFLSTGDEAAAGNWGLLD
QQAALRWVRDHAALFGGSAGAVTVAGDRSSADNVGLHLVAPGSRGLFQRAILMGGSVLSP
SAVQLDSATARSQAASLAQLVGCGHSSSEYLVRCLRDRSALALNAAQAQLLSGEYLQRWA
AVVDGHFLRDIPAEAAARGALADVDIVLGATEDDGALGRSRLPQVFPRFTSMSYSSAGFD
VALQKVLRGAGASSFARDAVRWFYKRGDGANPPPLADDESWLLSNVSRDYDVICPAVHMA
EMWAARGKANAFLYYVPTQQSWNTLEWEPHSDTQLAFGVPLHPERKGRSCVGEQQLSRHF
IGYLANFVKSGDPSFPNRHARGAGSRLLPAWPRFSLNEAGGFYKEVRAGMRNQRGLKMRE
CSFWKDYVQPLMAATGGIGEVERQWREDYNAWRQEALLEWRVQMSNFRAAVTESMATERP
AAAPYV
```

b

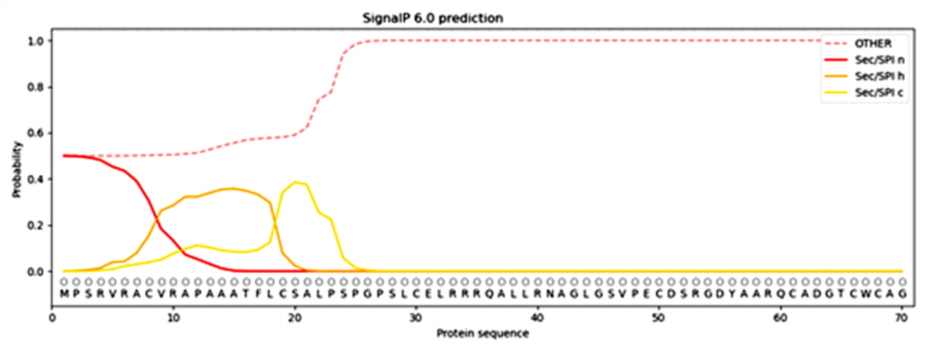


c

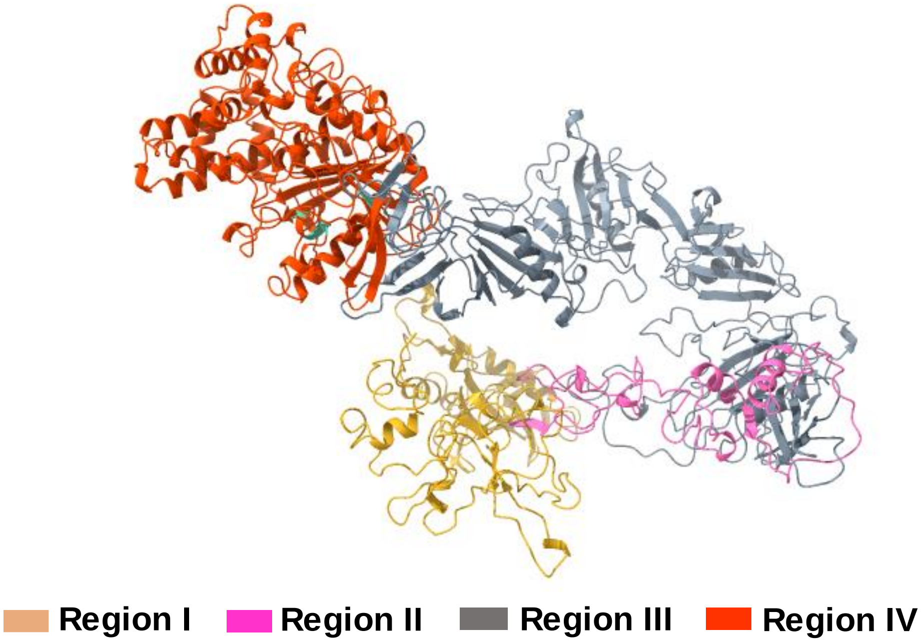

**Supplementary Figure S3. Sequence and three-dimensional atomic structure of the 1746-amino thyroglobulin isoform from *Petromyson marinus* (TGPM$^{1746}$), indexed in NCBI (XP_032817730.1).** **a)** The TGPM$^{1746}$ sequence comprises TG type 1-10 and TG type 1-11 modules, three TG type 2 modules, and five TG type 3 modules, along with linker and hinge domains, spacers 1, 2 and 3, ChEL domain. and the carboxy-terminal hormonogenic site for the formation of $T_3$. Sequence variations relative to TGPM$^{2475}$ were identified in the hinge domain and in the TG type 3-a1, TG type 3-b1, and TG type 3-a3 modules (see Figure 4). Amino acids are represented using single-letter codes. **b)** The SignalP 6.0 program predicts that the amino-terminal region is compatible with a signal peptide with a probability of 0.4996. The program identifies the following signal peptide regions: region n (amino-terminal), region h (central hydrophobic), and region c (C-terminal). **c)** A homology-based 3D model of TGPM$^{1746}$ was generated using the Swiss-Model platform, with *Bos taurus* thyroglobulin (PDB: 7N4Y) as the structural template. The resulting PDB file was visualized using UCSF ChimeraX.

```
TGPM_1-1    :  81
TGHS_1-1    :  81
TGPM_1-2    :  78
TGHS_1-2    :  78
TGPM_1-3    :  70
TGHS_1-3    :  76
TGPM_1-4    :  79
TGHS_1-4    :  81
TGPM_1-5    :  70
TGHS_1-5    :  84
TGPM_1-6    :  78
TGHS_1-6    :  76
TGPM_1-7    :  73
TGHS_1-7    :  76
TGPM_1-8    :  71
TGHS_1-8    :  73
TGPM_1-9    :  72
TGHS_1-9    :  74
TGPM_1-10   :  80
TGHS_1-10   :  76
TGPM_1-11   :  66
TGHS_1-11   :  80
cons        :  75

TGPM_1-1    ASACELLRQQAM-E-------------------------------------------------------
TGHS_1-1    LRPCELQRETAF---------------------------------------------------------
TGPM_1-2    LSFCQLHRQRVL-L--------SGY--------------------------------------------
TGHS_1-2    LSFCQLKQQIL-L--------SGY---------------------------------------------
TGPM_1-3    PSPCQVSVRRAV-R--------AS--T------------------------------------------
TGHS_1-3    PRSCEIRNRRLL----------HG---------------------------------------------
TGPM_1-4    PTVCEVERSLAE-R-------------------------------------------------------
TGHS_1-4    PTKCEVERFTAT-S-------------------------------------------------------
TGPM_1-5    ADAADGP--------------------------------------------------------------
TGHS_1-5    SQTCEQTPER-----------------------------------------------------------
TGPM_1-6    PSACEGERARAL-L--------AR--R------------------------------------------
TGHS_1-6    PTDCEKQRARMQSL--------MGS--------------------------------------------
TGPM_1-7    PTPCQAAAASEL-L--------QQ--LRHLGPV------------------------------------
TGHS_1-7    PTPCQLQSEQAF-L--------RT--VQALLS-------------------------------------
TGPM_1-8    PGACHLAAADVN-RYLQQADLLIG--SAGASSA--------GEGVAFGRGLAFTEDELLGSPLGLGDVK
TGHS_1-8    PGSCEEAKLRVL-Q--------FIRETEEIVSASNSSRFPLGESFLV--AKGIRLRNEDLGLPPLFPPR
TGPM_1-9    ATPCQRARAEAL-L--------TG--WKSLGSV------------------------------------
TGHS_1-9    PTTCEKSRTSGL-L--------SS--WKQAR--------------------------------------
TGPM_1-10   PSLCELRRRQAL-L--------RN---------------------------------------------
TGHS_1-10   PSLCNVLKSGVL-S--------RR---------------------------------------------
TGPM_1-11   RTPCEREALRRG-Q--------LG---------------------------------------------
TGHS_1-11   VTDCQRNEA------------------------------------------------------------

cons           ..

TGPM_1-1    --------------------------------------------DGRLHVPQCS-PH-G-AFRPLQCDA
TGHS_1-1    -------------------------------------------LKQADYVPQCA-ED-G-SFQTVQCQN
TGPM_1-2    ----------------------------------------L-NGTRALHLPQCT-QD-G-RYRPAQVDS
TGHS_1-2    ----------------------------------------I-NSTDTSYLPQCQ-DS-G-DYAPVQCDV
TGPM_1-3    ---------------------------------------------PGSPPPQCD-RD-EQRVLATQCQL
TGHS_1-3    --------------------------------------------VGDKSPPQCS-AE-G-EFMPVQCKF
TGPM_1-4    -------------------------------------------LGGRGFAPVCS-QD-G-SYIPTQCHG
TGHS_1-4    --------------------------------------------FGHPYVPSCR-RN-G-DYQAVQCQT
TGPM_1-5    -------------------------------------------PASRPFVARCG-EDGG-AYEPVQCHG
TGHS_1-5    -----------------------------------------------LFVPSCT-TE-G-SYEDVQCFS
TGPM_1-6    ---------------------------------------GRP-VGTPLFVPECD-GA-G-DYRPVQCSG
TGHS_1-6    ----------------------------------------Q-PAGSTLFVPACT-SE-G-HFLPVQCFN
TGPM_1-7    -----------------------------------L-RGDAPSSLPPLYVPRCD-AR-G-GWRPVQCDG
TGHS_1-7    ------------------------------------NSSML-PTLSDTYIPQCS-TD-G-QWRQVQCNG
TGPM_1-8    GEVAAVLAGGTGYAVRLAAQAAMHFHWRRRFFGPGTIGEGV-FNGFDPYMPQCT-EP-G-GWEPAQCVV
TGHS_1-8    EAFAEQFLRGSDYAIRLAAQSTLSFYQRRRFSPDDSAGASA-LLRSGPYMPQCD-AF-G-SWEPVQCHA
TGPM_1-9    -----------------------------------ENG-SS-IEVLTKHTPACT-PS-R-KYKKCHYL-
TGHS_1-9    -------------------------------------SQEN-PSPKDLFVPACL-ET-G-EYARLQAS-
TGPM_1-10   --------------------------------------------AGLGSVPECD-SR-G-DYAARQCA-
TGHS_1-10   --------------------------------------------VSPGYVPACRAED-G-GFSPVQCDQ
TGPM_1-11   --------------------------------------------AGERLELYCD-PE-G-LYEGEKPAK
TGHS_1-11   -------------------------------------------------GLQCD-QN-G-QYRASQKDR

cons                                                                *            :
```

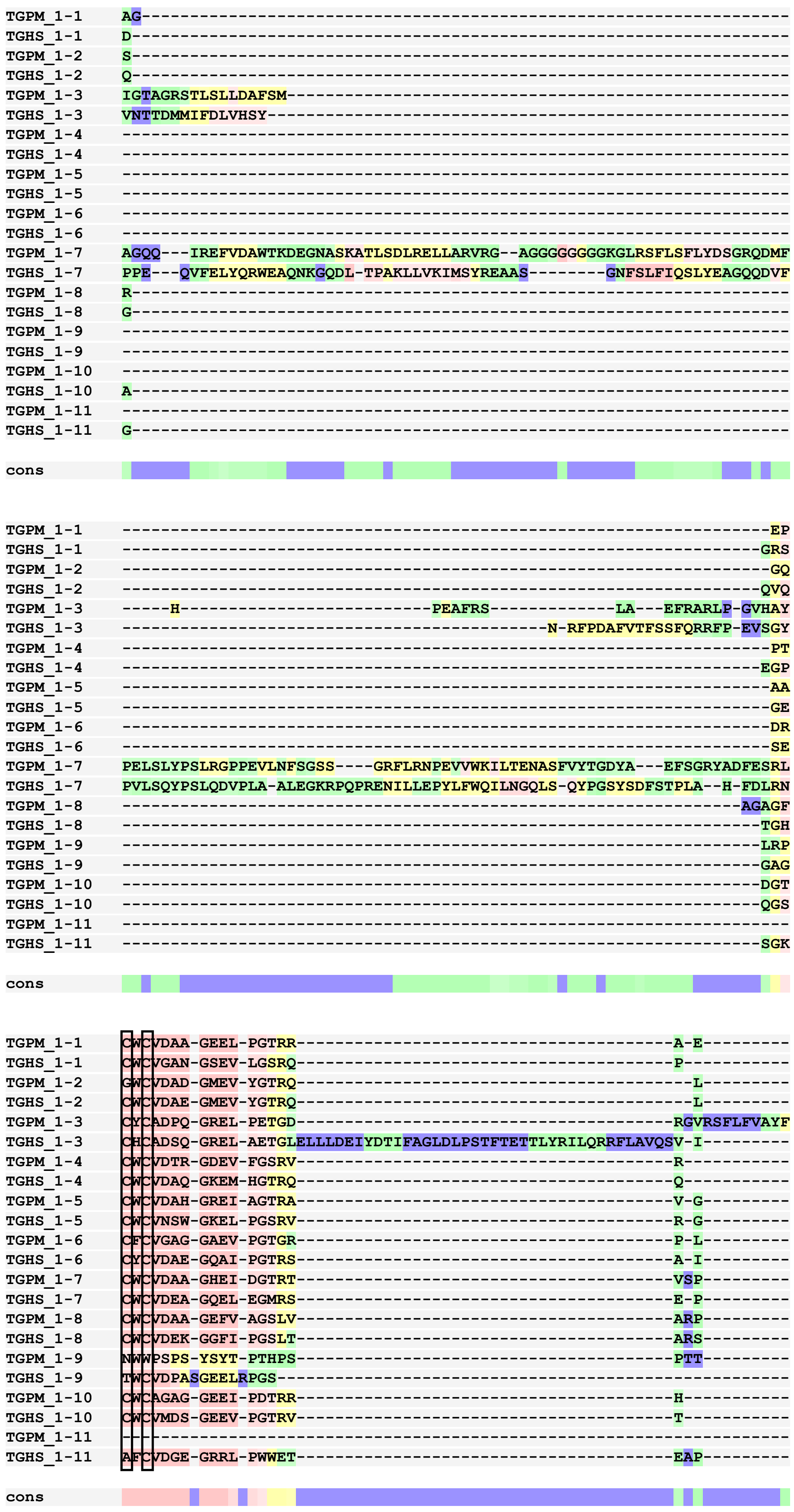
TGPM_1-1
TGHS_1-1
TGPM_1-2
TGHS_1-2
TGPM_1-3
TGHS_1-3
TGPM_1-4
TGHS_1-4
TGPM_1-5
TGHS_1-5
TGPM_1-6
TGHS_1-6
TGPM_1-7
TGHS_1-7
TGPM_1-8
TGHS_1-8
TGPM_1-9
TGHS_1-9
TGPM_1-10
TGHS_1-10
TGPM_1-11
TGHS_1-11
cons

```
TGPM_1-1    --------------GP----------------P--PSC
TGHS_1-1    --------------GR----------------P--VAC
TGPM_1-2    --------------GA----------------P--LSC
TGHS_1-2    --------------GR----------------P--KRC
TGPM_1-3    SGHGLNHEVVGSVLGKTRHEKFLKVQDRLPGHI--V-R
TGHS_1-3    --------------SG----------------R--FRC
TGPM_1-4    --------------DS----------------V--PNC
TGHS_1-4    --------------GE----------------P--PSC
TGPM_1-5    --------------GA----------------P--PRC
TGHS_1-5    ---------------G----------------Q--PRC
TGPM_1-6    --------------GD----------------P--VTC
TGHS_1-6    --------------GK----------------P--KKC
TGPM_1-7    --------------GQ----------------L--PKC
TGHS_1-7    --------------SK----------------L--PTC
TGPM_1-8    --------------RR----------------R--PQC
TGHS_1-8    --------------LQ----------------I--PQC
TGPM_1-9    --------------NA----------------E----S
TGHS_1-9    --------------SS----------------S--AQC
TGPM_1-10   --------------AG----------------EPGVVC
TGHS_1-10   --------------GG----------------Q--PAC
TGPM_1-11   --------------GG----------------Q----D
TGHS_1-11   --------------LE----------------D--SQC

cons
```

**Supplementary Figure 4. Alignment of the eleven TG type 1 modules of full-length thyroglobulin from *Petromyzon marinus* (TGPM) and *Homo sapiens* (TGHS).** TG type 1 modules were aligned using the T-Coffee program. Amino acids are represented using single-letter codes. Cysteine positions are boxed. BAD AVG GOOD

TGPM_1-1: TG type 1-1 TGPM, TGPM_1-2: TG type 1-2 TGPM, TGPM_1-3: TG type 1-3 TGPM, TGPM_1-4: TG type 1-4 TGPM, TGPM_1-5: TG type 1-5 TGPM, TGPM_1-6: TG type 1-6 TGPM, TGPM_1-7: TG type 1-7, TGPM, TGPM_1-8: TG type 1-8 TGPM, TGPM_1-9: TG type 1-9 TGPM, TGPM_1-10: TG type 1-10 TGPM, TGPM_1-11: TG type 1-11 TGPM.
TGHS_1-1: TG type 1-1 TGHS, TGHS_1-2: TG type 1-2 TGHS, TGHS_1-3: TG type 1-3 TGHS, TGHS_1-4: TG type 1-4 TGHS, TGHS_1-5: TG type 1-5 TGHS, TGHS_1-6: TG type 1-6 TGHS, TGHS_1-7: TG type 1-7 TGHS, TGHS_1-8: TG type 1-8 TGHS, TGHS_1-9: TG type 1-9 TGHS, TGHS_1-10: TG type 1-10 TGHS, TGHS_1-11: TG type 1-11 TGHS.

```
TGPM_2-1    :  97
TGHS_2-1    :  97
TGPM_2-2    :  97
TGHS_2-2    :  98
TGPM_2-3    :  97
TGHS_2-3    :  97
cons        :  94

TGPM_2-1    AQGCAACPRGSLWR---
TGHS_2-1    GLGCVKCPEGSYSQ---
TGPM_2-2    DGGCALCPADHYQDEAG
TGHS_2-2    DEECIPCPVGFYQEQAG
TGPM_2-3    QTACTPCPADTGTLTAG
TGHS_2-3    SLACVPCPVGRTTISAG

cons           *  ** .
```

**Supplementary Figure S5. Alignment of the three TG type 2 modules of full-length thyroglobulin from *Petromyzon marinus* (TGPM) and *Homo sapiens* (TGHS).** The TG type 2 modules were aligned using the T-Coffee program. Amino acids are represented using single-letter codes. Cysteine positions are boxed. BAD AVG GOOD

TGPM_2-1: TG type 2-1 TGPM, TGPM_2-2: TG type 2-2 TGPM, TGPM_2-3: TG type 2-3 TGPM.

TGHS_2-1: TG type 2-1 TGHS, TGHS_2-2: TG type 2-2 TGHS, TGHS_2-3: TG type 2-3 TGHS.

```
TGPM_3-a1   :  77
TGHS_3-a1   :  75
TGPM_3-b1   :  77
TGHS_3-b1   :  76
TGPM_3-a2   :  77
TGHS_3-a2   :  68
TGPM_3-b2   :  72
TGHS_3-b2   :  73
TGPM_3-a3   :  79
TGHS_3-a3   :  80
cons        :  69

TGPM_3-a1   CLQECLKE-ERCDFFVF---ST------QGDVSSCHFYNGSRESYQCEKAPEEK--SALGISLC-----
TGHS_3-a1   CLTDCTED-EACSFFTV---ST------TEPEISCDFYAWTSDNVACMTSDQKR--DALGNSKA-----
TGPM_3-b1   CRSACSAQ-ACCDGFLL---KE-----LPPGVLVCALLSGP-TVLTCRAPWWPRTADEYGDGECSGLRV
TGHS_3-b1   CLLACDRD-LCCDGFVL---TQ-----VQGGAIICGLLSSP-SVLLCNVKDWMDPSEAWANATCPGVTY
TGPM_3-a2   CLRRCRRD-ALCKAAAV---GD-----GPAGWLECLLYPD---TQSCGTSQELLERPARA-PSCA----
TGHS_3-a2   CLSRCVQEHSFCQLAEI---TE-----SASLYFTCTLYPE---AQVCDDIME-----SNA-QGCR----
TGPM_3-b2   CERTCDED-PCCEGFAF---LRRALSSAGEALLQCATHTGP-GVQLCGRESPMG-----ATFGCPPPGV
TGHS_3-b2   CERRCDAD-PCCTGFGFLNVSQ-----LKGGEVTCLTLNSL-GIQMCSEENG----GAWRILDCGSPDI
TGPM_3-a3   CLAVCQRS-PWCASVGV---RH------LGDSTRCVLHAE---TLACGYGAR------GG-RHCR----
TGHS_3-a3   CLSECSQH-EACLITTL---QT------QPGAVRCMFYAD---TQSCTHSLQ-------G-QNCR----

cons        *   *      *    .                 *           *                 .

TGPM_3-a1   ----VVFIA---IKCFVRGVG--------FHVSSASGRFYFHAGHELPSWDG------Q-LY----RPT
TGHS_3-a1   ----TSFGS---LRCQVKV-R--------SH-GQDSPAVYLKKGQGSTTTLQ------K-RF----EPT
TGPM_3-b1   HKPSRTFGFSLGGVHYNASNPPSAVASTAAAAAAASQVIYLWAGSEAGSRDV-CLHT-KAYTPQPGEPA
TGHS_3-b1   DQESHQVILRLGDQEFIKSLTPLE--GT-QDTFTNFQQVYLWKDSDMGSRPE-SMGCRKDTVPRPASPT
TGPM_3-a2   ----------------SLL-------------PRIDGTLYRKR-DGAEGPVR-R------LY----RRT
TGHS_3-a2   ----------------LIL-------------PQMPKALFRKK-VILEDKVK-N------FY----TRL
TGPM_3-b2   DTA---------------------------------SSLFGWYRAQARSSPQTAQLCPA-VTLPPAPTK
TGHS_3-b2   EVHTYPFGW---YQKPIAQ-N--------NA-PSFCP---------------------L-VV----LPS
TGPM_3-a3   ----------------LGV-------------LEPPARLYRKK--------------------------
TGHS_3-a3   ----------------LLL-------------REEATHIYRKP--------------------------

cons

TGPM_3-a1   GFGNAASGAYRTLALPAGTTTLPDA-----------HL-----------------Y
TGHS_3-a1   GFQNMLSGLYNPIVFSASGANLTDA-----------HL-----------------F
TGPM_3-b1   GAA-DVSGRFSTLRPE-EVTLQPEMRVAQQNFWLFRRAFSPP---QQATS-----W
TGHS_3-b1   EAG-LTTELFSPVDLN-QVIVNGNQSLSSQKHWLFKHLFSA----QQANL-----W
TGPM_3-a2   AFRSEIGVLLRSV-ANLTGMALSDG-------------FSR---------------
TGHS_3-a2   PFQKLMGISIRNK-VPMSEKSISNG-------------FFE---------------
TGPM_3-b2   ----GAADGFQSLDVS-TAAVDPTISAFDVVVLLGPGAGGSPRAAPPSDE-----W
TGHS_3-b2   LTEKVSLDSWQSLALS-SVVVDPSIRHFDVA-----HVSTA----ATSNFSAVRDL
TGPM_3-a3   --------------------------------------------------------
TGHS_3-a3   --------------------------------------------------------

cons
```

**Supplementary Figure 13. Alignment of the five TG type 3 modules of full-length thyroglobulin from *Petromyzon marinus* (TGPM) and *Homo sapiens* (TGHS).** The TG type 3 modules were aligned using the T-Coffee program. Amino acids are represented using single-letter codes. Cysteine positions are boxed. BAD AVG GOOD

TGPM_3-a1: TG type 3-a1 TGPM, TGPM_3-b1: TG type 3-b1 TGPM, TGPM_3-a2: TG type 3-a2 TGPM, TGPM_3-b2: TG type 3-b2 TGPM, TGPM_3-a3: TG type 3-a3 TGPM.
TGHS_3-a1: TG type 3-a1 TGHS, TGHS_3-b1: TG type 3-b1 TGHS, TGHS_3-a2: TG type 3-a2 TGHS, TGHS_3-b2: TG type 3-b2 TGHS, TGHS_3-a3: TG type 3-a3 TGHS.

```
                                                                 signal peptide
TGPM   ----------------------------------------MRTSPLLPATTTLYLVLW-   18
TGLF   MSRALPITPPDQWHTWEESTECKAAAAVAAADDDAAAAAWDMTGSDVS---AVPQAALWQ  57
TGLP   ------------------------------------------------------------  0

                                     TG Type 1-1
TGPM   --------IGTISALEYSDSKTSLASACELLRQQAMEDGRLHVPQCSPHGAFRPLQCDAA  70
TGLF   REQAEEEEPGTRRQPQYSDSKTSRASSCELLRQQAMEDGRSHVPQCSPHGAFRPLQCDAA  117
TGLP   -----MKTALMKSSHEYSDSKTSRASSCELLRQQAMEDGRSHVPQCSPHGAFRPLQCDAA  55
                       :******* **:************* *******************
                                        TG Type 1-2
TGPM   GEPCWCVDAAGEELPGTRRAEGPPPSCLSFCQLHRQRVLLSGYLNGTRALHLPQCTQDGR  130
TGLF   GDPCWCVDAAGEELPGTRRAEGQPPSCLSFCQLHRQRVLLSGYLNGTRALHLPQCAQDGR  177
TGLP   GDPCWCVDAAGEELPGTRRAEGQPPSCLSFCQLHRQRVLLSGYLNGTRALHLPQCAQDGR  115
       *:******************** ********************************:****
                                              TG Type 1-3
TGPM   YRPAQVDSSGQGWCVDADGMEVYGTRQLGAPLSCPSPCQVSVRRAVRASTPGSPPPQCDR  190
TGLF   YRPAQVDSSGQGWCVDADGMEVYGTRQLGAPLSCPSPCQVSVRRAVRASTPGSPPPQCDR  237
TGLP   YRPAQVDSSGQGWCVDADGMEVYGTRQLGVPLSCPSPCQVSVRRAVRASTPGSPPPQCDR  175
       *****************************.******************************

TGPM   DEQRVLATQCQLIGTAGRSTLSLLDAFSMHPEAFRSLAEFRARLPGVHAYCYCADPQGRE  250
TGLF   DEQRILATQCQLIGTAGRSTLSLLDAFSMHPEAFRSLAEFRARLPGVHAYCYCADPQGRE  297
TGLP   DEQRILATQCQLIGTAGRSTLSLLDAFSMHPEAFRSLAEFRARLPGVHAYCYCADPQGRE  235
       ****:*******************************************************
                                                          TG Type 1-4
TGPM   LPETGDRGVRSFLFVAYFSGHGLNHEVVGSVLGKTRHEKFLKVQDRLPGHIVRPTVCEVE  310
TGLF   LPETGMELLPLGEEDDAAALTALAPSAQEAPVFRLLRRRFLALRLLLSGSFRCPTVCEVE  357
TGLP   LPETGMELLPLGEEDDAAALTALAPSAQEAPVFRLLRRRFLALRLLLSGSFRCPTVCEVE  295
       *****  .  :          :   .*   ..   :  :  :   :.:**  ::   *  *  :  *******
                                                             LINKER
TGPM   RSLAERLGGRGFAPVCSQDGSYIPTQCHGPTCWCVDTRGDEVFGSRVRDSVPNCGSSSSP  370
TGLF   RSLADRLGGRGFAPVCSQDGSYVPTQCHGPTCWCVDTRGDEVFGSRVRDSVPNCGSSSSP  417
TGLP   RSLADRLGGRGFAPVCSQDGSYVPTQCHGPTCWCVDTRGDEVFGSRVRDSVPNCGSSSSP  355
       ****:*****************:*************************************

TGPM   CQLERQLALSHLFLGPSAPFIDDLISALPRDGSVSFSESPSQEMQHELWLHWRALREQLL  430
TGLF   CQLERQRALSHLFLGPSAPFTDDLISALPRDGSASFAESPSQEKQHELWLHWRALREQLL  477
TGLP   CQLERQRALSHLFLGPSAPFTDDLISALPRDGSASFAESPSQEKQHELWLHWRALREQLL  415
       ****** ************* ************.**:****** ****************

TGPM   DGGLGEILSQASRLFPASGQASRLLGDLVGGIFHTQELALTAAGANVQPGRIAEMLFGDR  490
TGLF   DGGLSEILSQASRLFPASGQGSRLLGDLMGGIFPTQELALTAAGANVQPGRIAEMLFGGG  537
TGLP   DGGLGEILSQASRLFPASGQGSRLLGDLMGGMFPTQELALTAAGANVQPGRIAEMLFGGG  475
       ****.***************.*******:**:* ************************.

TGPM   FLKNLKFFNFTGAVGGRGTFNFSKVFGQVGLTGMYDGANFEELARLFAPGEDSYLTRGSS  550
TGLF   FLKNLKFFNFTGAVGGRGTFNFSKVFGQVGLTGMYDGANFEELARLFAPGEDSYLTRGSS  597
TGLP   FLKNLKFFNFTGAVGGRGTFNFSKVFGQVGLTGMYDGANFEELARLFAPGEDSYLTRGSS  535
       ************************************************************

TGPM   NFSRESFNFDQSIDDPFARATNLERNRNVLGFVVTLLEDARFYGLVRDITPLIGLSGGDV  610
TGLF   NFSRESFNFDQSIDDPFARATNLERNRNVLGFVVTLLEDPRFYGLVRDIAPLIGLSGGDV  657
TGLP   NFSRESFNFDQSIDDPFARATNLERNRNVLGFVVTLLEDPRFYGLVRDIAPLIGLSGGDV  595
       *************************************** *********:**********
                            TG Type 1-5
TGPM   GLNALTFLRMAEEDRPTQNAADAADGPPASRPFVARCGEDGGAYEPVQCHGAACWCVDAH  670
TGLF   GMNALTFLRVAEEDRQTQNAADADDGPRASRPFVARCGEDGGAYEPVQCRGAACWCVDAH  717
TGLP   GLNALTFLRVAEEDRPTQNAADAYDGTRASRPFVARCGEDGGAYEPVQCRGAACWCVDAH  655
       *:*******:***** ******* **  ********************:**********
                         TG Type 1-6
TGPM   GREIAGTRAVGGAPPRCPSACEGERARALLARRGRPVGTPLFVPECDGAGDYRPVQCSGD  730
TGLF   GREVAGTRAVGGVPPRCPSACEGERARALLARRGRPVGTPLFVPECDGAGDYRPVQCSGD  777
TGLP   GREVAGTHAVGGVPPRCPSACEGERARALLARRGRPVGTPLFVPECDGAGDYRPVQCSGD  715
       ***:***:****.***********************************************
```

```
                                    TG Type 1-7
TGPM    RCFCVGAGGAEVPGTGRPLGDPVTCPTPCQAAAASELLQQLRHLGPVLRGDAPSSLPPLY    790
TGLF    RCFCVGAGGAEVPGTWRPLGDPVTCPTPCQVAAAGELLQQLRHLGPVLRGDAPSSPPPIY    837
TGLP    RCFCVGAGGAEVPGTWRPLGDPVTCPTPCQVAAAGELLQQLRHLGPVLRGDAPSSPPPIY    775
        *************** **************.***.******************** **:*

TGPM    VPRCDARGGWRPVQCDGAGQQIREFVDAWTKDEGNASKATLSDLRELLARVRGAGGGGGG    850
TGLF    VPRCDTRGGWRPVQCDGAGQQIREFVDAWTKDERDASKATLSDLRALLARARGAG-----    892
TGLP    VPRCDARGGWRPVQCDGAGQQIREFVDAWTKDEGDASKATLSDLRALLARARGAG-----    830
        *****:*************************** :********** ****.****

TGPM    GGKGLRSFLSFLYDSGRQDMFPELSLYPSLRGPPEVLNFSGSSGRFLRNPEVVWKILTEN    910
TGLF    -GEGLRSFLSFLYDSGRQDLFPELSLYPSLQGPLEVLNLSGSSGRFLRNPEVVWKILTEN    951
TGLP    -GDGLRSFLSFLYDSGRQDLFPELSLYPSLQGPIEVLNLSGSSSRFLRNPEVVWKILTEN    889
         *.****************:**********:** ****:****.****************
                                                        TG Type 1-8
TGPM    ASFVYTGDYAEFSGRYADFESRLCWCVDAAGHEIDGTRTVSPGQLPKCPGACHLAAADVN    970
TGLF    ASFVYTGDYAEFSGGYGDFESRLCWCVDAAGHEIDGTRTASPGQLPKCPGACHLAAADVS    1011
TGLP    ASFVYTGDYAEFSGGYGDFESRLCWCVDAAGHEIDGTRTASPGQLPKCPGACHLAAADVS    949
        ************** *.**********************.*******************.

TGPM    RYLQQADLLIGSAGASSAGEGVAFGRGLAFTEDELLGSPLGLGDVKGEVAAVLAGGTGYA    1030
TGLF    RYLQQADLLIGSAGASSAGEGVAFGRGLAFTEDELLGSPLGLGDVKGEVAAILAGGTGYA    1071
TGLP    RYLQQADLLIGSAGASSAGEGVAFGRGLAFTEDELLGSPLGLGDVKGEVAAILAGGTGYA    1009
        ***************************************************:********

TGPM    VRLAAQAAMHFHWRRRFFGPGTIGEGVFNGFDPYMPQCTEPGGWEPAQCVVRAGAGFCWC    1090
TGLF    VRLAAQAAMHFHWRRRFFGLGTIGEGVFNGFDPYMPQCTEPGAWEPAQCDK--SSGFCWC    1129
TGLP    VRLAAQAAMHFHWRRRFFGLGTIGEGVFNGFDPYMPQCTEPGAWEPAQCDK--SSGFCWC    1067
        ******************* **********************.******    .:*****
                            TG Type 1-9
TGPM    VDAAGEFVAGSLVARPRRRPQCATPCQRARAEALLTGWKSLGSVENGSSIEVLTKHTPAC    1150
TGLF    VDAAGEFVAGSLVARPRRRPQCATPCQRARAEALLTGWKSLGSVENGSSIEVLTKHTPAC    1189
TGLP    VDAAGEFVAGSLVARPRRRPQCATPCQRARAEALLTGWKSLGSVENGSSIEVLTKHTPAC    1127
        ************************************************************
                                                      TG Type 1-10
TGPM    TPSRKYKKC------HYLLRPNWWPSPSYSYT---PTHPSPTTNAESPSLCELRRRQALL    1201
TGLF    TPAGGQFEARQRDSESGLEQRSWCVDPASGQQAEPLGRDDPSGDIRCPSLCELRRRQALL    1249
TGLP    TPAGGQFEARQRDSESGLEQRSWCVDPASGQQAEPLGRDDPSGDIRCPSLCELRRRQALL    1187
        **:    :.        * : .*  .*: .       : .*: : ..*************
                                                        HINGE
TGPM    RNAGLGSVPECDSRGDYAARQCADGTCWCAGAGGEEIPDTRRHAGEPGVVCHEPRCELPQ    1261
TGLF    RDAGLGSVPECDSRGDYAARQCATGTCWCASAGGEEIPGTRRHAGEPGVVCHEPRCQLRQ    1309
TGLP    RDAGLGSVPECDSRGDYAARQCAEGTCWCASAGGEEIPGTRRHAGEPGVVCHEPRCQLPQ    1247
        *:********************* ******.*******.*****************:* *

TGPM    DERRAGRWALVCEDGAAATAAAGLQRCLLACLRGFAHTLDGAGHDGGAPREFQCNATSGE    1321
TGLF    DERRVGGWALVCEDGTAAAAAAGLQRCSLACLRGFARTLDGASHGGGAPQEFQCNATSGE    1369
TGLP    DERRVGGWALVCEDGTNA-AAAGLQRCSLACLRGFARTLDGASHGGGAPQEFQCNATSGE    1306
        ****.* ********: * ******** ********:*****.*.****:**********

TGPM    WIGSSPQPDACQEIHPWQAVQLSISFTLRFPEEKMCNPEYEGLLESFNSFLLADLTARGF    1381
TGLF    WIGSSPQPDACQEIHPWQAVQLSISFTLRFPEEKVCNPEYEGLLESFNSFLLADLTARGF    1429
TGLP    WIGSSPQPDACQEIHPWQAVQLSISFTLRFPEEKVCNPEYEGLLESFNSFLLADLTARGF    1366
        **********************************:*************************

TGPM    CHLSNGV------GSDAVAVCDAASVRVRCQDALRLLVNVTWRSSLSRLPVQSYPDLHNI    1435
TGLF    CHLSNGAQSAPGGGSGAVAVCDAASVRVRCQDALRLLVNVTWRSSLSRLPVQSYPDLHNI    1489
TGLP    CHLSNGAQSAPGGGSGAVAVCDAASVRVRCQDALRLLVNVTWRSSLSRLPVQSYPDLHNI    1426
        ******.      **.********************************************

TGPM    ERAFVAHAQRFAQLLDDRGYRLALNGRDYSSEGAARFRKDELYDTSPAFRVACEPGYVRI    1495
TGLF    ERAFVAHAQRFAQLLDDRGYRLALNGRDYSSEGAARFRKDELYDTSPAFRMACKPGYVRI    1549
TGLP    ERAFVAHAQRFAQLLDDRGYRLALNGRDYSSEGAARFRKDELYDTSPAFRMACEPGYVRI    1486
        **************************************************:**:******
```

```
      TG Type 2-1       TG Type 2-2       TG Type 2-3        Spacer 1 TG Type 1-11
TGPM  AQGCAACPRGSLWRDGGCALCPADHYQDEAGQTACTPCPADTGTLTAGAAAAFQCRTPCE  1555
TGLF  AQGCAACPRGSLWRDGVCAPCPTDHYQDEPGQTACTPCPADSGTLTTGAAAASQCRTPCE  1609
TGLP  AQGCAACPRGSLWRDGVCAPCPTDHYQDEPGQTACTPCPADSGTLTTGAAAASQCRTPCE  1546
      **************** ** **:****** ***********:****:***** *******
                                                  Spacer 2
TGPM  REALRRGQLGAGERLELYCDPEGLYEGEKPAKGGQDEVRRMFEEVPSEDLVLGAVDAVAL  1615
TGLF  REALRRGQLGAGERVELYCDPEGLYEGEKPAKGGQDEVRRMFEEVPSEDLVLGAVDAVAL  1669
TGLP  REALRRGQLGAGERVELYCDPEGLYEGEKPAKGGQDEVRRMFEEVPSEDLVLGAVDAVAL  1606
      **************:*********************************************
                     TG Type 3-a1
TGPM  RIRTETGPVQEVEPRCLQECLKEERCDFFVFSTQGDVSSCHFYNGSRESYQCEKAPEEKS  1675
TGLF  RIRTETGPVQEVEPRCLQECLKEERCDFLVFSTQGDVSSCHFYNGSRESYQCETAPEAPG  1729
TGLP  RIRTETGPVQEVEPRCLQECLKEERCDFLVFSTQGDVSSCHFYNGSRESYQCETAPEAPG  1666
      ****************************:************************.***   .

TGPM  ALGISLCVVFIAIKCFVRGVGFHVSSASGRFYFHAGHELPSWDGQLYRPTGFGNAASGAY  1735
TGLF  FLGDPDSSWVERLSCRPRVS-L-SPGSVFTVYRKKGHELPSWDGQLYRPTGFGNAASGAY  1787
TGLP  FLGDPDSSWVERLSCRPRVS-L-SPGSVFTVYRKKGHELPSWDGQLYRPTGFGNAASGAY  1724
       **   .  .  :.*  *   :    .:   .* : *************************
                        TG Type 3-b1
TGPM  RTLALPAGTTTLPDAHLYCRSACSAQACCDGFLLKELP--PGVLVCALLSGPTVLTCRAP  1793
TGLF  RTLALPAGTTTLPDAHLYCRSACSAQACCDGFLLKELPLDNGVLVCALLSGPTVLTCRAP  1847
TGLP  RTLALPAGTTTLPDAHLYCRSACSAQACCDGFLLKELPLDNGVLVCALLSGPTVLTCRAP  1784
      **************************************   ******************

TGPM  WWPRTADEYGDGECSGLRVHKPSRTFGFSLGGVHYNASNPPSAVASTAAAAAAASQVIYL  1853
TGLF  GWPRTADEYGDGECSGLRVHKPSRTFGFSLGGVHYNASNPPSAVASTAAAATAATAAAAQ  1907
TGLP  GWPRTADEYGDGECSGLRVHKPSRTFGFSLGGVHYNASNPPSAVASTAAAATAATAAAAQ  1844
       ***************************************************:**:  .

TGPM  -----WAGSEAGSRDVCLHTKAYTPQPGEPAGAADVSGRFSTLRPEEVTLQPEMRVAQQN  1908
TGLF  VPAAALCGAPMCSPGLCRESFAV---TASASTAPDVSGRFSTLRPEEVTLQPEMRVAQQN  1964
TGLP  VPAAALCGAPMCSPGLCRESFAV---TACASAAPDVSGRFSTLRPEEVTLQPEMRVAQQN  1901
            .*:    *  .:*  :  *     .    :  * **************************
                     TG Type 3-a2
TGPM  FWLFRRAFSPPQQATSWCLRRCRRDALCKAAAVGDGPAGWLECLLYPDTQSCGTSQELLE  1968
TGLF  FWLFRRAFSP-QQATSWCLRRCRGDALCKAAAVGDGPAGWLECLLYPDTQSCGTSQQLLD  2023
TGLP  FWLFRRAFSP-QQATSWCLRRCRGDALCKAATVGDGPAGWLECLLYPDTQSCGTSQQLLD  1960
      ********** ************ *******:***********************:**:

TGPM  RPARAPSCASLLPRIDGTLYRKRDGAEGPVRRLYRRTAFRSEIGVLLRSVANLTGMALSD  2028
TGLF  WPARAPSCASLLPRLDGTLYRKRDGAEGPVRRLYRRTAFRSEVGVLLRSVANLTGMALFD  2083
TGLP  WPARAPSCASLLPRLDGTLYRKRDGAEGPVRRLYRRTAFRSEVGVLLRSVANLTGMALFD  2020
       *************:***************************:*************** *
        TG Type 3-b2
TGPM  GFSRCERTCDEDPCCEGFAFLRRALSSAGEALLQCATHTGPGVQLCGRESPMGATFGCPP  2088
TGLF  GFSRCERTCDEDPCCEGFAFLRSSLS--REALLQCATHTGPGVQLCGRESPVGATFGCPP  2141
TGLP  GFSRCERTCDEDPCCEGFAFLRSSLS--REALLQCATHTGPGVQLCGRESPVGATFGCPP  2078
      **********************  :**   ***************************:********

TGPM  PGVDTASSLFGWYRAQARSSPQTAQLCPAVTLPPAPTKGAADGFQSLDVSTAAVDPTISA  2148
TGLF  PGVDTASSLFGWYRAQARSSPQAAQLCPAVSLPPAPTKGAADGFHCLDVSTAAVDPTISA  2201
TGLP  PGVDTASSLFGWYRAQARSSPQAAQLCPAVSLPPAPTKGAADGFHCLDVSTAAVDPTISA  2138
      **********************:*******:*************:.***************
                              TG Type 3-a3
TGPM  FDVVVLLGPGAGGSPRAAPPSDEWCLAVCQRSPWCASVGVRHLGDSTRCVLHAETLAC-G  2207
TGLF  FDVVVLLGPGAGGSPQAAPPSDEWCLAACQQSPWCASVGVRHLGERTRCVLHAETLACSG  2261
TGLP  FDVVVLLGPGAGGSPQAAPPSDEWCLAACQQSPWCASVGVRHLGERTRCVLHAETLACSG  2198
      ***************:***********.**:***************:  ************ *
                              Spacer 3                   ChEL
TGPM  YGARGGRHCRLGVLEPPARLYRKKAPSPGGGQGPVRPVVPLGSGQLQGASRTVEVDSQVK  2267
TGLF  HGARGGRHCHLGVLEPPARLYRKK----GGGQGLVRPTVPLGSGQLQGASRTVEVDSQVK  2317
TGLP  HGARGGRHCHLGVLEPPARLYRKK----GGGQGLVRPTVPLGSGQLQGASRTVEVDSQVK  2254
      :********:**************    ***** ***.***********************
```

```
TGPM      VIDVFLGVPYAAPPLDANRFSGPQPPRPLAVNETWDADRYKTRPDCLQPDGQRGTSLSED      2327
TGLF      VVDVFLGVPYAAPPLDANRFSGPQAPRPLAVNVTWDADRYK--PDCLQPDGQRGTSLSED      2375
TGLP      VVDVFLGVPYAAPPLDANRFSGPQAPRPLAVNVTWDADRYK--PDCLQPDGQRGTSLSED      2312
          *:********************** ******* ********  *****************

TGPM      CLYLNIFVPKVKPHNASVLVFFHGGDNSFGGSAQGPLDPSYLAALGDIIVVTANYRLGLF      2387
TGLF      CLYLNIFVPKVKPHNASVLVFFPGGDNSFGGSAQGPLDPSYLAALGDIIVVTANYRLGLF      2435
TGLP      CLYLNIFVPKVKPHNASVLVFFPGGDNSFGGSAQGPLDPSYLAALGDIIVVTANYRLGLF      2372
          ********************** *************************************

TGPM      GFLSTGDEAAAGNWGLLDQQAALRWVRDHAALFGGSAGAVTVAGDRSSADNVGLHLVAPG      2447
TGLF      GFLSTGDEAAAGNWGLLDQQAALRWVRDHAALFGGSAGAVTVAGDRSSADNVGLHLVAPG      2495
TGLP      GFLSTGDEAAAGNWGLLDQQAALRWVRDHAALFGGSAGAVTVAGDRSSADNVGLHLVAPG      2432
          ************************************************************

TGPM      SRGLFQRAILMGGSVLSPSAVQLDSAT--ARSQAASLAQLVGCGHSSSEYLVRCLRDRSA      2505
TGLF      SRGLFQRAILMGGSVLSPSAVQLDSAAAAAARSQAAALAQLVGCGHSSSEYLVRCLRSRSA      2555
TGLP      SRGLFQRAILMGGSVLSPSAVQLDSAAAAAARSQAAALAQLVGCGHSSSEYLVRCLRSRSA      2492
          **************************:  ******:********************.***

TGPM      LALNAAQAQLLSGEYLQRWAAVVDGHFLRDIPAEAAARGALADVDIVLGATEDDGALGRS      2565
TGLF      LALNAAQAQLLSGEYLQRWAAVVDGHFLRDIPAEAAARGALADVDIVLGATEDDGALGRS      2615
TGLP      LALNAAQAQLLSGEYLQRWAAVVDGHFLRDIPAEAAARGALADVDIVLGATEDDGALGRS      2552
          ************************************************************

TGPM      RLPQVFPRFTSMSYSSAGFDVALQKVLRGAGASSFARDAVRWFYKRGDGANPPPLADDES      2625
TGLF      RLPQVFPRFTTMSYSSAGFDVALQKVLRGAGDSSFARDAVRWFYKRGDGADPPPLAADES      2675
TGLP      RLPQVFPRFTTMSYSSAGFDVALQKVLRGAGDSSFARDAVRWFYKRGDGADPPPLAADES      2612
          **********:******************** ******************:***** ***

TGPM      WLLSNVSRDYDVICPAVHMAEMWAARGKANAFLYYVPTQQSWNTLEWEPHSDTQLAFGVP      2685
TGLF      WLLSNVSRDYDVVCPAVHMAEMWASRGKANAFLYYVPTQQSWNTLEWEPHSDTQLAFGVP      2735
TGLP      WLLSNVSRDYDVVCPAVHMAEMWASRGKANAFLYYVPTQQSWNTLEWEPHSDTQLAFGVP      2672
          ************:***********:***********************************

TGPM      LHPERKGRSCVGEQQLSRHFIGYLANFVKSGDPSFPNRHARGAGSRLLPAWPRFSLNEAG      2745
TGLF      LHPERKGRSCVAEQQLSRHFIGYLANFVKSGDPSFPNRHARGAGSRLLPAWPRFSLNEAG      2795
TGLP      LHPERKGRSCVAEQQLSRHFIGYLANFVKSGDPSFPNRHARGAGSRLLPAWPRFSLNEAG      2732
          ***********.************************************************

TGPM      GFYKEVRAGMRNQRGLKMRECSFWKDYVQPLMAATGGIGEVERQWREDYNAWRQEALLEW      2805
TGLF      GLYKEVRAGMRNHRGLKMRECSFWKDYVQPLMAATGGIGEVERQWREDYHAWRQEALLEW      2855
TGLP      GLYKEVRAGMRNHRGLKMRECSFWKDYVQPLMAATGGIGEVERQWREDYHAWRQEALLEW      2792
          *:**********:************************************:**********

TGPM      RVQMSNFRAAVTGSMATERPAAAPYV      2831
TGLF      RVQMSNFRAAVTGSTATERPAAAPYV      2881
TGLP      RVQMSNFRAAVTGSTATERPAAAPYV      2818
          ************** ***********
```

**Supplementary Figure S7. Comparative alignment of full-length thyroglobulin sequence from *Petromyzon marinus* (TGPM, 2831 amino acids), *Lampetra fluviatilis* (TGLF, 2881 amino acids) and *Lampetra planeri* (TGLP, 2818 amino acids).** The alignment was performed using the Clustal Omega program. Amino acids are represented using single-letter codes. The figure displays the linker, hinge, and ChEL domain, along with eleven TG type 1 modules, three TG type 2 modules, five TG type 3 modules, and spacers 1, 2, and 3. Cysteine residues are highlighted in yellow. N-terminal hormonogenic acceptor and donor sites, and C-terminal hormonogenic sites are marked in red. The signal peptide of TGPM is boxed.

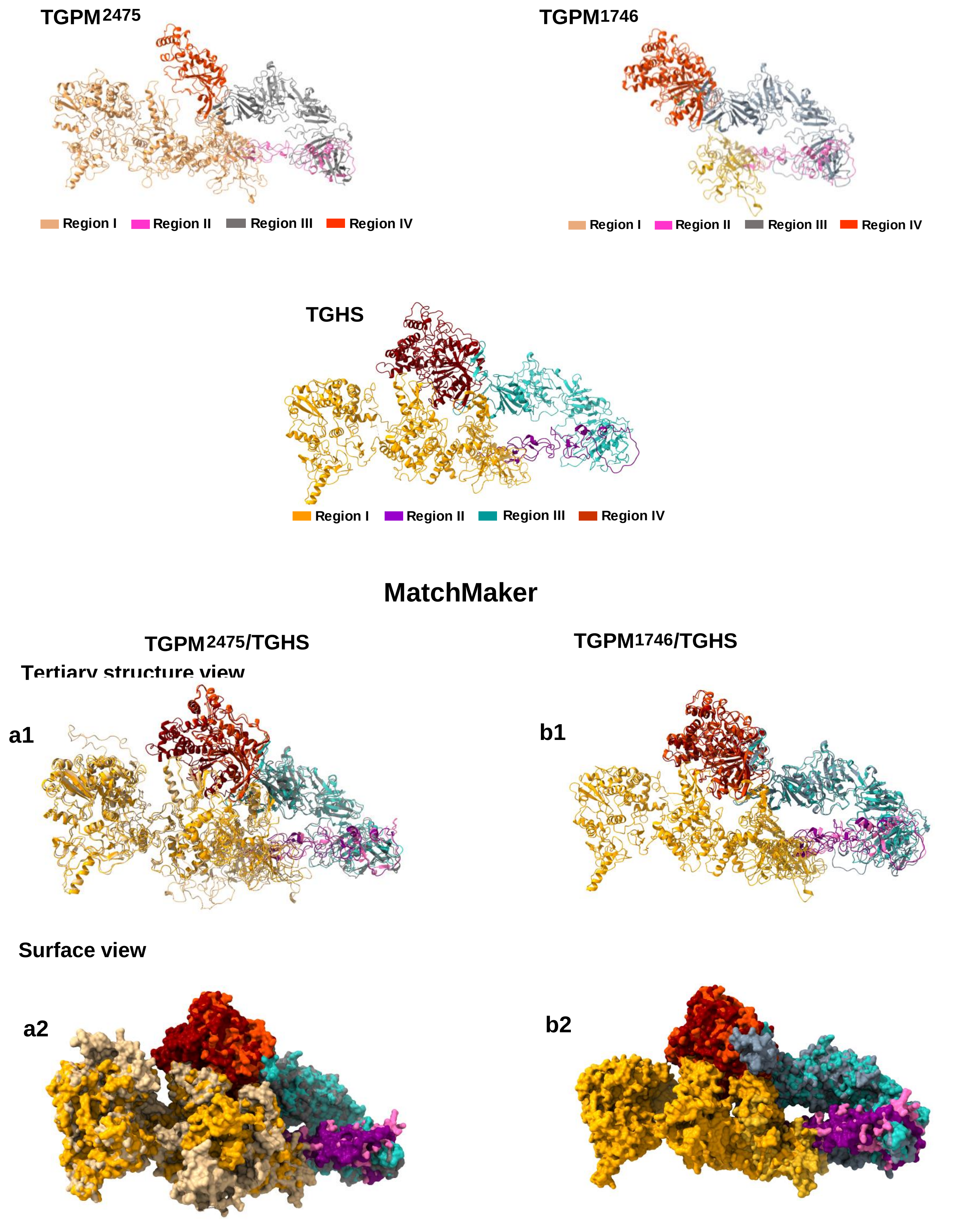


**Supplementary Figure S8. Three-dimensional atomic structure alignment of thyroglobulins from *Petromyzon marinus* (TGPM$^{2475}$ and TGPM$^{1747}$) and *Homo sapiens* (TGHS).** The top panel shows the homology model of TGPM$^{2475}$ and TGPM$^{1747}$. The middle panel shows the TGHS model. The bottom panel illustrates the structural superposition of thyroglobulin monomers (TGPM$^{2475}$ with TGHS and TGPM$^{1747}$ with TGHS) across the four canonical regions (I-IV) of the classical model, performed using the Matchmaker command in UCSF ChimeraX. a1 and b1 panels display the superposition of monomers without surface rendering, while the a2 and b2 include surface visualization.

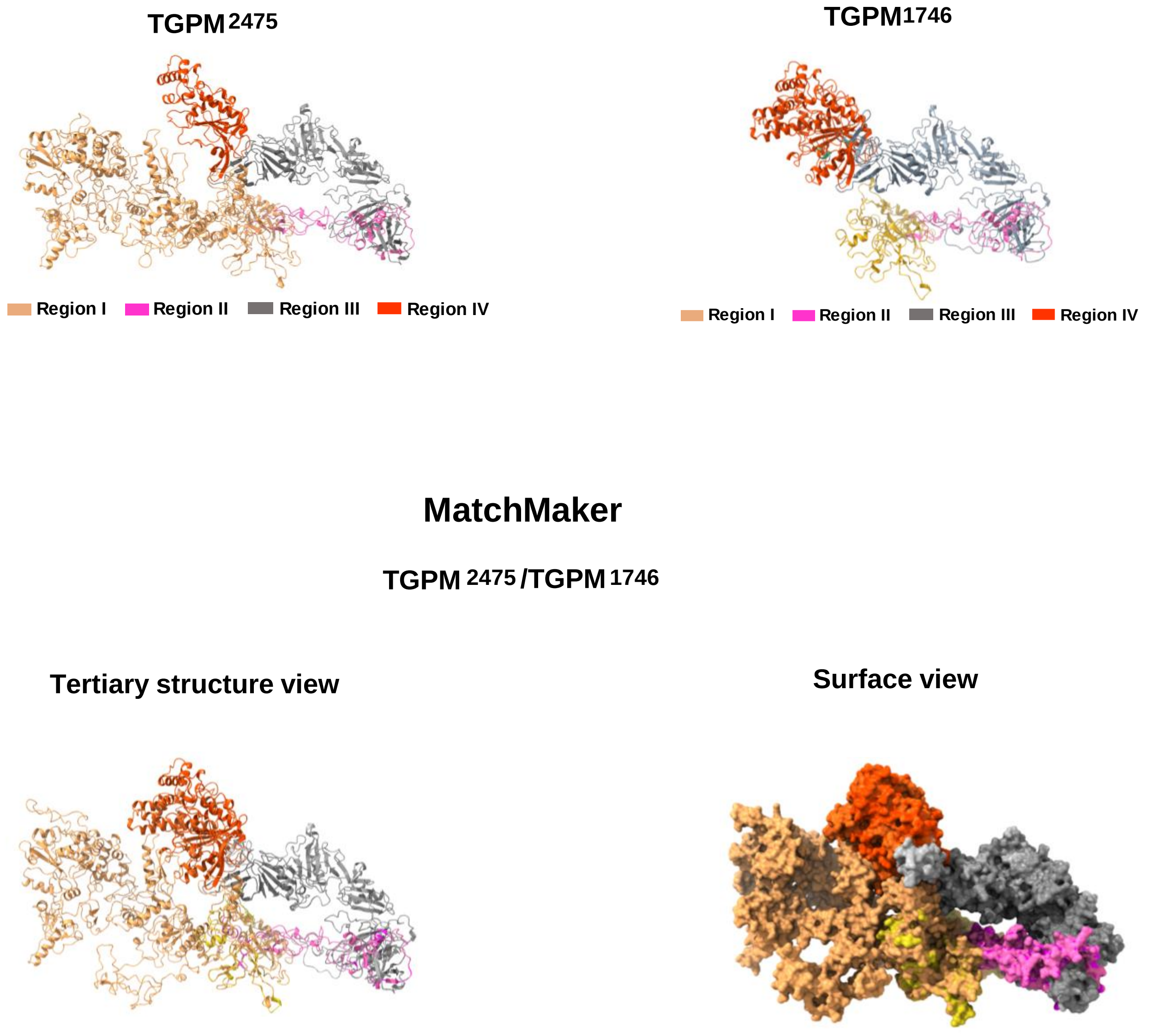


**Supplementary Figure S9. Three-dimensional atomic structure alignment of thyroglobulins from *Petromyzon marinus* (TGPM$^{2475}$ and TGPM$^{1747}$).** The top panel shows the homology model of TGPM$^{2475}$ and TGPM$^{1747}$. The bottom panel illustrates the structural superposition of thyroglobulin monomers (TGPM$^{2475}$ with TGPM$^{1747}$) across the four canonical regions (I-IV) of the classical model, performed using the Matchmaker command in UCSF ChimeraX. The left panel displays the superposition of monomers without surface rendering, while the right panel includes surface visualization.

signal peptide

```
Homo_sapiens                     -----------------------------------------------------MALVLEI  7
Pan_troglodytes                  -----------------------------------------------------MALVLEI  7
Gorilla_gorilla                  -----------------------------------------------------MALVLEI  7
Pongo_pygmaeus                   -----------------------------------------------------MALVLGI  7
Macaca_mulatta                   -----------------------------------------------------MALVLEI  7
Macaca_fascicularis              -----------------------------------------------------MALVLEI  7
Rattus_norvegicus                ----------------------------------------------------MMTLVLWV  8
Mus_musculus                     ----------------------------------------------------MTALVLWV  8
Cavia_porcelus                   -----------------------------------------------------MALVLWA  7
Canis_lupus_familiaris           -----------------------------------------------------MALALWV  7
Panthera_leo                     -----------------------------------------------------MVLAPWV  7
Bos_taurus                       -----------------------------------------------------MALALWV  7
Trichechus_manatus_latirostris   -----------------------------------------------------MALVLWV  7
Columbia_livia                   -----------------------------------------M--E-V--PQRMVGSVPFC  14
Taeniopygia_guttata              -----------------------------------------M--G-L--SHCKMGSVPFC  14
Larus_michahellis                -----------------------------------------M--E-L--SPRKMGSVPFC  14
Gallus_gallus                    -----------------------------------------------------MGSIPAC  7
Struthio_Camelus                 ------------------------------------------------------------  0
Chelonia_mydas                   -----------------------------------------------------MGPVLLS  7
Alligator_mississippiensis       ------------------------------------------------------------  0
Crotalus_tigris                  -----------------------------------------------------MRALLFI  7
Python_bivittatus                ------------------------------------------------------------  0
Eublepharis_macularius           -----------------------------------------------------MGLFLLN  7
Xenopus_tropicalis               -----------------------------------------M--APV--LLSTM-PYRFI  14
Aquarana_Catesbeiana             -----------------------------------------------------M-ARLSI  6
Carassius_auratus                -----------------------------------------------MQNMEEMKLLCVT  13
Cyprinus_carpio                  -----------------------------------------------MQNMDEKKLLCVT  13
Danio_rerio                      -----------------------------------------------MQNMEEMKLLCLT  13
Astyanax_mexicanus               ---------------------------------------------------MNMHMFWAL  9
Clupea_harengus                  MTLHFH------LGAYKG---------V---PQRLP----AQ--GELLLCSDRMGLVCLT  36
Oryzias_latipes                  ---------------------------------------------------MMAWLLFIA  9
Xiphophorus_maculatus            ---------------------------------------------------MMAGLLCTV  9
Stegastes_partitus               ----------------------------------------------------MAWLLQIT  8
Cynoglossus_semilaevis           ----------------------------------------------------MAWLLSII  8
Lepisosteus_oculatus             -----------------------------------------------------MGYLGSA  7
Lampetra_fluviatilis             MSRALPITPPDQWHTWEESTECKAAAAVAAADDDAAAAAWDMTGSDVS---AVPQAALWQ  57
Lampetra_planeri                 ------------------------------------------------------------  0
Petromyzon_marinus               -----------------------------------------MRTSPLLPATTTLYLVLW-  18
```

$T_4$-forming acceptor site

TG type 1-1

```
Homo_sapiens                     FTLLASICWVSANI----FEYQ-VDAQPLRPCELQRETAFLKQADYVPQCAEDGSFQTVQ  62
Pan_troglodytes                  FTLLASICWVSANI----FEYQ-VDAQPLRPCELQRETAFLKQADYVPQCAEDGSFQTVQ  62
Gorilla_gorilla                  FTLLASICWVSANI----FEYQ-VDAQPLRPCELQRETAFLKQADYVPQCAEDGSFQTVQ  62
Pongo_pygmaeus                   FSLLASVCWVSANI----FEYQ-VDAQPLRPCELQRETAFLKQADYVPQCAEDGSFQTVQ  62
Macaca_mulatta                   FSLLASVCWVSANI----FEYQ-VDAQPLRPCELQRERAFLKQADYVPQCAEDGSFQPVQ  62
Macaca_fascicularis              FSLLASVCWVSANI----FEYQ-VDAQPLRPCELQRERAFLKQADYVPQCAEDGSFQPVQ  62
Rattus_norvegicus                STLLSSVCLVAANI----FEYQ-VDAQPLRPCELQREKAFLKQDEYVPQCSEDGSFQTVQ  63
Mus_musculus                     STLLSSVCLVAANI----FEYQ-VDAQPLRPCELQREKAFLKQAEYVPQCSEDGSFQTVQ  63
Cavia_porcelus                   FGLLGSACLVSANI----FEYQ-VDAQPLRPCELQRERAFLKRMEYIPQCSEDGSFQTIQ  62
Canis_lupus_familiaris           FSLLGSACVVSANI----FEYQ-VDAQPLRPCELQRERAFLKGAEYVPQCAEDGSYQTVQ  62
Panthera_leo                     LSLLGSACLVSANI----FEYQ-VDAQPLRPCELQRERAFLKRADYVPQCTEDGSYQTVQ  62
Bos_taurus                       FGLLDLICLASANI----FEYQ-VDAQPLRPCELQRERAFLKREDYVPQCAEDGSFQTVQ  62
Trichechus_manatus_latirostris   FGLLGSASSAWANI----FEYQ-VDMQPLRPCELQRERAFLKREDYVPQCSEDGSFQAVQ  62
Columbia_livia                   TLLCCLISIAAANI----FEYQ-TDAQPLRPCELQREKAFSGGETYVPQCSEDGQFRTVQ  69
Taeniopygia_guttata              TL-CCFISIAAANI----FEYQ-TDSQPLRPCELQREEAFAAGEAYVPQCTEDGQFRTVQ  68
Larus_michahellis                TLLCCFISVAAANI----FEYQ-TDSQPLRPCELQREKAFSGGETYVPQCSEDGQFRTVQ  69
Gallus_gallus                    TLLS-LISIAAANI----FEYQ-TDSQPLRPCELQREKAFTGGETYVPQCSEDGQFRAVQ  61
Struthio_Camelus                 -----------MRQ----KEYQ-TDSQPLRPCELQREKAFSEGETYVPQCSEDGQFRTVQ  44
Chelonia_mydas                   TL-FFFISTASANI----FEYQ-AESQPLRPCELQREKAFLEGEVYVPQCLEDGQFRTVQ  61
Alligator_mississippiensis       ---------MEDLF----REYQ-AESQPLRPCELRREKAFLEGEDHVPQCSEDGQFRTVQ  46
Crotalus_tigris                  TCLC-FTDKALTSIFVPNVEYE-AETQPLRPCELQREKAFHDGEDYIPQCSEDGQFRKKQ  65
Python_bivittatus                ------------------LEYQ-AESQPLRPCELQREKAFRDGEDYVPQCSEDGQFRKIQ  41
Eublepharis_macularius           TFLC-FVNIAAASI----FEYQ-AETQPLRPCELQREKAFQRGEDYVPQCSEDGQFRNIQ  61
Xenopus_tropicalis               YTIFCVFRIIAAIE----TEYQ-LDSQPLRPCELARERALSNRESYVPQCSSEGLYRNIQ  69
Aquarana_Catesbeiana             ITLTCIIGTASATV----AEYE-LESQPLLPCEIAREKSLSVGGDYIPQCSVEGQYRNVQ  61
Carassius_auratus                LTLLSIFTLTCGKI----SEYQ-LETESLSQCEQLRAVSSAQGHEHIPQCSEDGRFRHVQ  68
Cyprinus_carpio                  FTLLGIFSLTCGKI----SEYQ-LETESLSQCEQLRAVSSTQGHEHIPQCSEDGRFRHVQ  68
Danio_rerio                      STLICIFSLTCGKI----SEYQ-LETESLSQCEQLRSVSAEQERQHVPQCFEDGRFRHVQ  68
Astyanax_mexicanus               AFLLISCCITDGKI----SEYQ-LESESRSPCELLRSVGSEQQNEHIPQCSEDGRYRYVQ  64
Clupea_harengus                  ILLAATLMLSEGKI----SEYQ-LETESLSQCELLRGVGPTQQRDHVPQCSEDGRFRHVQ  91
Oryzias_latipes                  GILVHFPDPLLGNA----SEYQ-LESDTQTQCQLLRGVAAAKQPGDVPHCFEDGSFRPVQ  64
Xiphophorus_maculatus            WILLCGAALLDGRA----SEYQ-LESEALSRCEILRSVAVGTKQEEVPHCLEDGSFRSVQ  64
Stegastes_partitus               CILVCCPALLQGKA----SEYQ-LESETLSQCEALRGVAVAKQQGHIPHCTEDGRFRPVQ  63
Cynoglossus_semilaevis           TC-LGGLSLLHAKP----SEYQ-LESEPLSRCESLRGVAVANKKVVVPQCSEDGSFRPVQ  62
Lepisosteus_oculatus             FISLCCFSLSLSKR----SEYQ-LESQSLSQCELLRDEGSVQGGEFTPQCLDDGRFRNIQ  62
Lampetra_fluviatilis             REQAEEEE--PGTR--RQPQYSDSKTSRASSCELLRQQAMEDGRSHVPQCSPHGAFRPLQ  113
Lampetra_planeri                 -----MKT--ALMK--SSHEYSDSKTSRASSCELLRQQAMEDGRSHVPQCSPHGAFRPLQ  51
Petromyzon_marinus               ----------IGTI--SALEYSDSKTSLASACELLRQQAMEDGRLHVPQCSPHGAFRPLQ  66
```

```
                                                                  TG type 1-2
Homo_sapiens                      CQNDGRSCWCVGANGSEVLGSRQ-PGRPVACLSFCQLQKQQILLSGYINSTDTSYLPQCQ   121
Pan_troglodytes                   CQNDGRSCWCVGADGSEVLGSRQ-PGRPVACLSFCQLQKQQILLSGYINSTDTSYLPQCQ   121
Gorilla_gorilla                   CQNDGHSCWCVGADGSEVLGSRQ-PGRPVACLSFCQLQKQQILLSGYINSTDTSYLPQCQ   121
Pongo_pygmaeus                    CQNDGRSCWCVGADGSEVLGSRQ-PGRPVACLSFCQLQKQQILLSGYINSTDTSYLPQCQ   121
Macaca_mulatta                    CQNDGRSCWCVGADGSEVLGSRQ-PGRPVACLSFCQLQKQQILLSGYINSTDTSYLPQCQ   121
Macaca_fascicularis               CQNDGRSCWCVGADGSEVLGSRQ-PGRPVACLSFCQLQKQQILLSGYINSTDTSYLPQCQ   121
Rattus_norvegicus                 CQNDGQSCWCVDSDGTEVPGSRQ-LGRPTACLSFCQLHKQRILLSSYINSTDALYLPQCQ   122
Mus_musculus                      CQNDGQSCWCVDSDGREVPGSRQ-LGRPTVCLSFCQLHKQRILLGSYINSTDALYLPQCQ   122
Cavia_porcelus                    CRNDGRSCWCVDADGREVPGSRQ-SRRPAACLSFCQLHRQQILLNGYLNSTATSYLPQCQ   121
Canis_lupus_familiaris            CQNDGRTCWCVGADGVEVPGSRQ-PARPAACLSFCQLQRQQILLSGYLNSTATSYLPQCQ   121
Panthera_leo                      CEGDGGSCWCVGADGKEVPGSRQ-PGRPVACLSFCQLQRQQILLRGYVNSTATSYLPQCQ   121
Bos_taurus                        CGKDGASCWCVDADGREVPGSRQ-PGRPAACLSFCQLQKQQILLSSYINSTATSYLPQCQ   121
Trichechus_manatus_latirostris    CRKDGRACWCVDADGREVPGSRQ-PGRLVSCQSFCQLQKQQILVSGYLNSTATSYLPQCQ   121
Columbia_livia                    CSRDGLSCWCVDEDGIEVPGSKQ-NGYPISCLSFCQLQKQRILVSRYINSSSVSYIPQCL   128
Taeniopygia_guttata               CSRNGLSCWCVDENGVEVPGSKQ-NGYPISCLSFCQLQRQRILVSRYINSSSISYIPQCL   127
Larus_michahellis                 CSRNGLSCWCVDENGIEVPGSKQ-NGYPVSCLSFCQLQKQRILVSRYINSSSISYIPQCL   128
Gallus_gallus                     CSRNGLSCWCVDENGIEVPGSKQ-NGYPISCLSFCQLQKQRVLVSGYINSSSISYIPQCL   120
Struthio_Camelus                  CSRDGLSCWCVDENGIEVPGSKQ-NGYPLSCLSFCQLQKQRILVSRYINSSSFSYVPQCL   103
Chelonia_mydas                    CNTNGLSCWCVDANGIEVPGTKQ-TGVPIACLSFCQLQKQQILVSHYINSSSTSYVPECL   120
Alligator_mississippiensis        CSKNNLSCWCVDDKGAEVPGSKQ-NGVPISCLSFCQLQKQQVLVSRYINSSTISYIPQCL   105
Crotalus_tigris                   CDRKGLSCWCVDEKGIEVTGTKR-SGSSLSCLSFCQLQKQQILVSRYINSSETAYIPQCL   124
Python_bivittatus                 CNKNGVSCWCVDEKGIEVTGSKQ-SGSSIACLSFCQLQKQRILVSRYINSSATAYIPQCL   100
Eublepharis_macularius            WNKLQFSCWCVDEKGTELPGSKQ-NGSSIACLSFCQLQKQQILISRYINSSATPYIPQCL   120
Xenopus_tropicalis                CSGDGQTCWCVNANGAELAGSRQ-TDAPPVCLSFCQLAKQRILVSGYINSTSTSHIPQCK   128
Aquarana_Catesbeiana              CNRRGTSCWCVNSDGTEVAGSRQ-AASPPICLSFCQLKKQEILVSGYINSSSTSYIPQCK   120
Carassius_auratus                 CNHGGRECWCVNAEGLEIPGSRQ-NASTVHCLTSCQLQRQQALQTEG-----ITLVPACL   122
Cyprinus_carpio                   CNRGGGECWCVNVEGLEIPGSRQ-NASTVYCLTSCQLQRQQALQTEG-----ITLVPSCL   122
Danio_rerio                       CNRGGGECWCVNAEGFEIPGSRQ-NGSSVYCLTTCQLQRQRALQTED-----ATLVPVCL   122
Astyanax_mexicanus                CNRG-GECWCVNSDGTEIPGSRQ-NDTVVTCLTPCQLQRQRLLLSGD-----AGVVPRCL   117
Clupea_harengus                   CSAG-GECWCVDAEGAELPGSRQ-NGSAIHCLTSCQMHRQRVLQRGD----ETGLLPVCS   145
Oryzias_latipes                   CSGRSQECWCVDADGREIAGTRT-NNSALSCPSVCQLQT----------------RLRCS   107
Xiphophorus_maculatus             CGGRSQECWCVDSEGQEVTGTRT-NNSIPHCPTPCQLQS----------------LLRCA   107
Stegastes_partitus                CSGQSQECWCVNAEGQEVAGTRT-NSSAPHCPTACQLQS----------------VLRCS   106
Cynoglossus_semilaevis            CSGRGHECWCVDAAGQEIVGTLT-NNSALHCPSPCQLQS----------------ELRCS   105
Lepisosteus_oculatus              CRRDGLMCWCVTANGTEVPGTKR-NGSTIHCLSSCELHRQQVLRSGE-----AVFLPQCS   116
Lampetra_fluviatilis              CDAAGDPCWCVDAAGEELPGTRRAEGQPPSCLSFCQLHRQRVLLSGYLNGTRALHLPQCA   173
Lampetra_planeri                  CDAAGDPCWCVDAAGEELPGTRRAEGQPPSCLSFCQLHRQRVLLSGYLNGTRALHLPQCA   111
Petromyzon_marinus                CDAAGEPCWCVDAAGEELPGTRRAEGPPPSCLSFCQLHRQRVLLSGYLNGTRALHLPQCT   126

                                         T4-forming donor site              TG type 1-3
Homo_sapiens                      DSGDYAPVQCDVQQVQCWCVDAEGMEVYGTRQLGRPKRCPRSCEIRNRRLLHGVGD---K   178
Pan_troglodytes                   DSGDYTPVQCDVQQVQCWCVDAEGMEVYGTRQLGRPKRCPRSCEIRNRRLLHGVGD---K   178
Gorilla_gorilla                   DSGDYTPVQCDVQQVQCWCVDAEGMEVYGTRQLGRPKRCPRSCEIRNRRLLHGVGD---K   178
Pongo_pygmaeus                    DSGDYTPVQCDVQQVHCWCVDAEGMEVYGTRQLGRPKRCPRSCEIRNRRLLHGVGD---K   178
Macaca_mulatta                    DSGDYMPVQCDVQQVQCWCVDAEGMEVYGTRQPGRPKRCPRSCEIRNRRLLHGVGD---K   178
Macaca_fascicularis               DSGDYMPVQCDVQQVQCWCVDAEGMEVYGTRQPGRPKRCPRSCEIRNRRLLHGVGD---K   178
Rattus_norvegicus                 DSGNYAPVQCDLQQVQCWCVDTEGMEVYGTRQQGRPTRCPRSCEIRSRRLLHGVGD---K   179
Mus_musculus                      DSGNYAPVQCDLQRVQCWCVDTEGMEVYGTRQQGRPTRCPRSCEIRNRRLLHGVGD---R   179
Cavia_porcelus                    DSGDYASVQCDLWQTECWCVDTEGMEVYGTRQLGRPTRCPRSCEIRNRRILHGVGD---K   178
Canis_lupus_familiaris            ESGGYAPVQCDLWQQQCWCVDAEGMEVYGTRQLGRPARCPRSCDIRNRRLLHGVGD---K   178
Panthera_leo                      DSGDYAPVQCDPWRGQCWCVDAEGMEVYGTRQPGRPTRCPRSCEIRNRRLLHGVGD---R   178
Bos_taurus                        DSGDYSPVQCDLRRRQCWCVDAEGMEVYGTRQQGRPARCPRSCEIRNRRLLHGVGD---R   178
Trichechus_manatus_latirostris    DSGGYAPVQCDVWQLQCWCVDAEGMEVYGTRQRGRPTRCPRSCEIRNRRILHGVGD---K   178
Columbia_livia                    DSGAFDEVQCDMEMGQCWCVDPEGMEIYGTRQRGKPRQCPGSCEIRDRRILHGFGE---K   185
Taeniopygia_guttata               DSGAFDEVQCDMELGQCWCVDPEGMEIYGTRQRGKPAQCPGNCEIRDRRILHGFGE---K   184
Larus_michahellis                 DSGAFDAVQCDTELGQCWCVDPEGMEIYGTRQRGKPTRCPGNCEIRDRRILHGFGE---K   185
Gallus_gallus                     DSGAFDVMQCDVELGQCWCVDQEGMEIYGTRQRGKPTQCPGTCEIRDRRILHGVGE---K   177
Struthio_Camelus                  DSGMFDTVQCDVELGQCWCVDPEGMEIYGTRQRGKPTQCPGNCEIRDRRILHGVGE---K   160
Chelonia_mydas                    NSGEFDPVQCDVGPGQCWCVDSEGMEIYGTRQTGKPMQCPGSCEIRDRRILHGVGE---R   177
Alligator_mississippiensis        DSGEFAPVQCDVGLGQCWCVDSEGMEIYGTRQTGKPTQCPGSCEIRDRRILHGVGD---R   162
Crotalus_tigris                   DSGDFDPIQCDLALVQCWCVDGEGMEIYGTRQKGKPRQCPGRCEIRDRRILHGVGE---K   181
Python_bivittatus                 DSGDFDPIQCDLALEQCWCVDGEGMEIYGTRQKGKPKQCPGQCDIRDRRILHGVGE---K   157
Eublepharis_macularius            DSGDFDPIQCDVGLRQCWCVDTEGMEIYGTRQRGKPTRCPGKCEIRDRRILHGVGE---K   177
Xenopus_tropicalis                DSGEFEPVQCHRESGQCWCVDSEGMEIYGTRQTGKPSQCPRSCQVRGRRILHGFGE---K   185
Aquarana_Catesbeiana              NSGEYEDVQCDRQLGECWCVDNEGMEIYGTRQIGQPTRCPKRCAVRDRRILHDVGE---K   177
Carassius_auratus                 DSGEFEPVQCDASRGQCWCVDQEGMEIYGTRQNGNPSKCPGTCEVRQRRLLHGIGE---P   179
Cyprinus_carpio                   DSGEFEPVQCDASRGQCWCVDQEGMEIYGTRQNGKLSKCPGTCEVRQRRLLHGIGE---P   179
Danio_rerio                       DSGEYQQVQCDASRSQCWCVDLEGMEIYGTRQNGKPSKCPGTCEVRERRLLHGIGE---R   179
Astyanax_mexicanus                DSGEYEPVQCDGALGQCWCVDMEGMEIYGTRQNGRPSQCPSVCETRQRRLLHGVGE---S   174
Clupea_harengus                   NSGEYQAVQCQPAAGQCWCVDQEGMEIYGTRQNGRPARCPGGCEIRERRLLHGVSE---P   202
Oryzias_latipes                   PSGLFEAIQCDSSRGQCWCVDQDGMELYGTRQNGRPQRCPGSCQVRARRLIHSSG-----   162
Xiphophorus_maculatus             PSGLFEPVQCD-SRGRCWCVDQDGMELYGTRQAGRPRRCPSSCEVRSRRLLHSSA----S   162
Stegastes_partitus                ASGLFEQVQCESSRGQCWCVDQDGMELYGTRQSGRPHRCPGSCEVRKRRLLHTSTSPSPP   166
Cynoglossus_semilaevis            PSGQFESIQCDTNRGHCWCVDHDGMELYGTRQNGRPQQCPGSCEVRGRRLLHGAS--SSM   163
Lepisosteus_oculatus              DSREYEPVQCDSTRSQCWCVDAEGMEIYGTRQNGQPVRCPGACEVRDRRLLHGVGD---S   173
Lampetra_fluviatilis              QDGRYRPAQVD-SSGQGWCVDADGMEVYGTRQLGAPLSCPSPCQVSVRRAVRASTP--GS   230
Lampetra_planeri                  QDGRYRPAQVD-SSGQGWCVDADGMEVYGTRQLGVPLSCPSPCQVSVRRAVRASTP--GS   168
Petromyzon_marinus                QDGRYRPAQVD-SSGQGWCVDADGMEVYGTRQLGAPLSCPSPCQVSVRRAVRASTP--GS   183
```

```
Homo_sapiens                      SPPQCSAEG-EFMPVQCKFVNTTDMMIFDLVHSYNRFPDAFVTFSSFQRRFPEVSGYCHC   237
Pan_troglodytes                   SPPQCSAEG-EFMPVQCKFVNTTDMMIFDLVHSYNRFPDAFVTFSSFQRRFPEVSGYCHC   237
Gorilla_gorilla                   SPPQCSAEG-EFMPVQCKFVNTTDMMIFDLVHSYNRFPDAFVTFSSFQRRFPEVSGYCHC   237
Pongo_pygmaeus                    SPPQCSAEG-EFMPVQCKFVNTTDMMIFDLVHSYNRFPDAFVTFSSFQRRFPEVSGYCHC   237
Macaca_mulatta                    SPPQCSAEG-EFMPVQCKFVNTTDMMIFDLVHSYNRFPDAFVTFNSFQRRFPEVSGYCHC   237
Macaca_fascicularis               SPPQCSAEG-EFMPVQCKFVNTTDMMIFDLVHSYNRFPDAFVTFNSFQRRFPEVSGYCHC   237
Rattus_norvegicus                 SPPQCDADG-EFMPVQCKFVNTTDMMIFDLIHNYNRFPDAFVTFSAFRNRFPEVSGYCYC   238
Mus_musculus                      SPPQCTADG-EFMPVQCKFVNTTDMMIFDLIHNYNRFPDAFVTFSSFRGRFPEVSGYCYC   238
Cavia_porcelus                    SPPQCSPEG-EFLPVQCKFVNTTDMMIFDLIHSYNRFPDAFVTFSSFKSRFPEVSGYCHC   237
Canis_lupus_familiaris            SPPQCSPDG-AFMPVQCKFVNTTDMMIFDLVHSYNRFPDAFVTFSAFRSKFPDVSGYCHC   237
Panthera_leo                      SPPQCSPDG-RFLPVQCKFVNTTDMMIFDLVHSYNRFPDAFVTFSAFRNKFPEVSGYCHC   237
Bos_taurus                        SPPQCSPDG-AFRPVQCKLVNTTDMMIFDLVHSYSRFPDAFVTFSSFRSRFPEVSGYCYC   237
Trichechus_manatus_latirostris    SPPQCTAEG-EFMPVQCKFVNTTDMMIFDLIQSYNRFPDAFMTFRTFQSRFPKVSGYCHC   237
Columbia_livia                    SPPQCSADG-EFLPVQCKFVNTTDMMFFDLVHSYNRLPEAFQTFSSLRKTFPEVSGYCYC   244
Taeniopygia_guttata               SPPQCSADG-EFLPVQCKFVNTTDMSVFDLVHSYNRLPRAFQKFSSVREAFPEISGYCYC   243
Larus_michahellis                 SPPQCSADG-EFLPVQCKFVNTTDMMFFDLVHSYNRLPRAFQTFSSLRETFPEVSGYCYC   244
Gallus_gallus                     SPPQCSEDG-EFLPVQCKFVNTTDMMVFDLLHNYNRLPAAFQTFSSFRETFPEVSGYCYC   236
Struthio_Camelus                  SPPQCSPDG-EFFPVQCKFVNTTDMMFFDLVHNYNRLPEAFQTFSSLRRTFPEVSGYCYC   219
Chelonia_mydas                    SPPQCSADG-EFLPVQCKFVNMTDMMVFDLVHSYNRFPDAFQTFSSFRSMFPEVSGYCYC   236
Alligator_mississippiensis        SPPQCSADG-EFLPVQCKFVNMTNMMVFDLVHSYNRFPEAFQTFSSFRSMFPGVSGYCYC   221
Crotalus_tigris                   SPPQCSTDG-EFLPVQCKFVNTTDRMVFDLIHHFNRLPKAFLTFSSFRSLFPEINGYCYC   240
Python_bivittatus                 TPPQCTTDG-EFLPVQCKFVNTTDRMVFDLIHHFNRFPRAFLTFSSFRNLFSEISGYCYC   216
Eublepharis_macularius            SPPQCSADG-EFLPVQCKFVNRTNMMVFDLLHHFNRFPEAFLTFSSFRTSFPEISGYCYC   236
Xenopus_tropicalis                TPPHCSADG-TFQPIQCKLLNTTDKMAVDVVSSFSRFPETFESFSSLRESFPEISGYCYC   244
Aquarana_Catesbeiana              VPPHCSEDG-GFQPVQCKLVNTTDQMVVDLVTTFSRSPDSFKSFSSLRESFPEISGYCYC   236
Carassius_auratus                 SPPQCSDDG-GFLPVQCKLVNTTDRMIFDLLHTFNRDPEVFQTFSAFRKAYPDLSSYCFC   238
Cyprinus_carpio                   SPPQCSDDG-GFLPVQCKFVNTTDRMIFDLLHTFNRDPELFQTFSAFRKAYPELSSYCFC   238
Danio_rerio                       SPPQCSDDG-GFLPVQCKFVNTTDRMIFDLLHTFNRQPEVFQTFSGFRKAYPELSSYCFC   238
Astyanax_mexicanus                SPPQCSDNG-RFLPVQCKFINTTNMMVFDLLHAFNRRPEVFQTFSGFRKAYPELSSYCYC   233
Clupea_harengus                   SPPQCSDDG-VFLPVQCKFINTTDSMVFDLLHTFNRHPELFQTFSGFRNTFPEISSYCSC   261
Oryzias_latipes                   SPPQCAEDG-SFLPVQCKFINMTDRMELDLLHAFNRFPEAFQTFSNFRKVFPMVSSYCFC   221
Xiphophorus_maculatus             SPPQCAADG-SFLPVQCKFINTTDRTELDLLLAFSSSPEAFDTFRNFRKFFPLVSSYCFC   221
Stegastes_partitus                SPPQCDEDG-TFLPVQCKFINTTNRAELDLLHAFNRFPEAFETFSSFRKFFPLVSSYCFC   225
Cynoglossus_semilaevis            SPPQCSRDG-RFLPVQCKFINMTDKSELDLLQAFSRFPESFSTFSSFRKTFPMVSSYCYC   222
Lepisosteus_oculatus              SPPQCTASG-DFLPVQCKFINTTDMTVFDLLHTFNRFPGVFQTFTGFRKMFPEISGYCFC   232
Lampetra_fluviatilis              PPPQCDRDEQRILATQCQLIGTAGRSTLSLLDAFSMHPEAFRSLAEFRARLPGVHAYCYC   290
Lampetra_planeri                  PPPQCDRDEQRILATQCQLIGTAGRSTLSLLDAFSMHPEAFRSLAEFRARLPGVHAYCYC   228
Petromyzon_marinus                PPPQCDRDEQRVLATQCQLIGTAGRSTLSLLDAFSMHPEAFRSLAEFRARLPGVHAYCYC   243
```

```
Homo_sapiens                      ADSQGRELAETGLELLLDEIYDTIFAGLDLPSTFTETTLYRILQRRFLAVQSVISGRFRC   297
Pan_troglodytes                   ADSQGRELAETGLELLLDEIYDTIFAGLDLPSTFTETTLYRILQRRFLAVQSVISGRFRC   297
Gorilla_gorilla                   ADSQGRELAETGLELLLDEIYDTIFAGLDLPSTFTETTLYRILQRRFLAVQSVISGRFRC   297
Pongo_pygmaeus                    ADSQGRELAETGLELLLDEIYDTIFAGLDLPSTFTETTLYRILQRRFLAVQSLISGRFRC   297
Macaca_mulatta                    ADSQGRELAETGLELLLDEIYDTIFAGLDLPSTFTETTLYRILQRRFLAVQSVISGRFRC   297
Macaca_fascicularis               ADSQGRELAETGLELLLDEIYDTIFAGLDLPSTFTETTLYRILQRRFLAVQSVISGRFRC   297
Rattus_norvegicus                 ADSQGRELAETGLELLLDEIYDTIFAGLDQASTFTQSTMYRILQRRFLAIQLVISGRFRC   298
Mus_musculus                      ADSQGRELAETGLELLLDEIYDTIFAGLDQASTFTQSTMYRILQRRFLAIQLVISGRFRC   298
Cavia_porcelus                    ADSQGRELAETGLELLLDEIYDTIFAGLDLTSTFTESTLYRILQRRFLAVQLVVSGRFRC   297
Canis_lupus_familiaris            ADSLGRELAGTGLELLLDEIYDTIFAGLDLASTFAETTLYRILQRRFLAVQLVLSGRFRC   297
Panthera_leo                      ADSQGRELAETGLELLLDEIYDTIFAGLDLASTFTETTLYRILQRRFLAVQLLISGRFRC   297
Bos_taurus                        ADSQGRELAETGLELLLDEIYDTIFAGLDLASTFAETTLYRILQRRFLAVQLVISGRFRC   297
Trichechus_manatus_latirostris    ADSQGRELAETGLELLLDEIYDTIFAGLDLASTFSETTLYRILQRRFLAVQLVISGRFRC   297
Columbia_livia                    ADSLGRELADTGLELLLDEVYDTVFSALEPARTFMETSIYRILHRRFLGVQLATTGKFRC   304
Taeniopygia_guttata               VDSLGRELADTGLELLLDEVYDTIFSEPEPARTFTESSIHRILQRRFLGVQLATSGKFRC   303
Larus_michahellis                 TDSLGRELADTGLELLLDEVYDTVFSALEPARTFTETSVYRILQRRFLGVQLATSGKFRC   304
Gallus_gallus                     VDSLGRELADTGLELLLDEVYDTVFSVLEPARTFTETSVYRILQRRFLGVQLATSGRFRC   296
Struthio_Camelus                  ADSLGRELADTGLELLLDEVYDTIFSALEPARTFTETSIYRILQRRFLGVQLATSGKFRC   279
Chelonia_mydas                    ADSLGRELAETGLELLLEEVYDTVFTALQPAHTFTETSIYRILQRRFLGVQLATSGRFRC   296
Alligator_mississippiensis        ADSLGRELAETGLELLLEEVYDTIFSALEPALTFAETTMYRILQRRFLGVQLAALGRFRC   281
Crotalus_tigris                   ADSLGRELEDTGLEMLLENVYDTVFASLEPVPTFAESSMYRIRQRRFLGVQLATTGRFRC   300
Python_bivittatus                 ADSLGRELEDTGLEMLLENVYDTVFASLEPVPTFAESSMYRILQRRFLGIQLATTGRFRC   276
Eublepharis_macularius            ADSLGRELEETGLEMLLDNVYDTVLASMEPAQTFHETSMYRILQRRFLGVRLATSGRFRC   296
Xenopus_tropicalis                ADRLGRELAGTGLELLLDEVYDTVFSGLAPSRTFTETTVYRILQRRFLGVRLMTCGRFRC   304
Aquarana_Catesbeiana              ADLLGRELAGTGVELLLDEVYDTVFSTFTRSRTFTETAIYRILQRRFLGVQLITSGKFRC   296
Carassius_auratus                 ADSRGREMPSTGVELLLDEVYDTAFSDQTAGSTFAQSNMYRILQRRYLAVQLAVSGRFRC   298
Cyprinus_carpio                   ADSRGREMPSTGVELLLDEVYDTAFSDQTAGSTFAQSNMYRILQRRYLAVQLAVNGRFRC   298
Danio_rerio                       ADSRGREMPSTGVELLLDEVYDTAFSDQTAGSSFAESNMYRILNRRYLAVQLAINGKFRC   298
Astyanax_mexicanus                ADTRGRELPSTGLELLLDEVYDTAFSGLDVGRSFSQTNMYRILQRRFFAVQLALTGRFRC   293
Clupea_harengus                   ADSRGRELPSTGLELLLDEVYDTAFSDTGAGRSFTQSNMYRILQRRFLAVQLAMTGRFRC   321
Oryzias_latipes                   SNSQGRELENTGVELLFSEIPESAFSGLSSGQSFSQTNIYKVLQRRMLGIQLAVSGHFRC   281
Xiphophorus_maculatus             SDGRGRELQGTGVELLMSEVYDSAFSGSRSARSFAQTNIYRVLQRRMLGVRLALTGHFRC   281
Stegastes_partitus                TDSRGREMENTGVELLLSEVYDSAFSGLSSGRSFAQTNIYRVLQRRMLGLRLALTGHFRC   285
Cynoglossus_semilaevis            SDSRGRELDNTGLELLLSGVDDSAFLGLGSVHSFSQTNVYQVLQRRMLGVRLALTGFFRC   282
Lepisosteus_oculatus              ADSRGRELANTGLELLLDEVYDTAFSGLEPGPSFTKSNMYRILQRRFLGIQLAMSGRFRC   292
Lampetra_fluviatilis              ADPQGRELPETGMELLPLGEEDDAAALTALAPSAQEAPVFRLLRRRFLALRLLLSGSFRC   350
Lampetra_planeri                  ADPQGRELPETGMELLPLGEEDDAAALTALAPSAQEAPVFRLLRRRFLALRLLLSGSFRC   288
Petromyzon_marinus                ADPQGRELPETGDRGVRSFLFVAYFSGHGLNHEVVGSVLGKTRHEKFLKVQDRLPGHIVR   303
```

TG type 1-4

```
Homo_sapiens                     PTKCEVERFTATSFG-HPYVPSCRRNGDYQAVQCQTEGPCWCVDAQGKEMHGTRQ-QGEP     355
Pan_troglodytes                  PTKCEVERFTATSFG-HPYVPSCRRNGDYQAVQCQTEGPCWCVDAQGKEMHGTRQ-QGEP     355
Gorilla_gorilla                  PTKCEVERFTATSFG-HPYVPSCRRNGDYQAVQCQTEGPCWCVDAQGKEMHGTRQ-QGEP     355
Pongo_pygmaeus                   PTKCEVERFTATSFG-HPYVPSCHQNGDYQAVQCQMEGPCWCVDAQGKEMHGTRQ-QGEP     355
Macaca_mulatta                   PTKCEVERFTATSFG-HPYVPSCRRNGDYQAVQCQTEGPCWCVDVQGKEIHGTRQ-QGEP     355
Macaca_fascicularis              PTKCEVERFTATSFG-HPYVPSCRRNGDYQAVQCQTEGPCWCVDVQGKEIHGTRQ-QGEP     355
Rattus_norvegicus                PTKCEVEQFTATSFG-HPYIPSCHRDGHYQTVQCQMERMCWCVDAQGIEIPGTRQ-QGQP     356
Mus_musculus                     PTKCEVEQFAATRFG-HSYIPRCHRDGHYQTVQCQTEGMCWCVDAQGREVPGTRQ-QGQP     356
Cavia_porcelus                   PTKCEVERFAAASFD-HPYIPSCHRNGDYQAVQCQTQGPCWCVDAQGKEIPGTQQWAGEP     356
Canis_lupus_familiaris           PTRCEVEQFTASSFG-HPYTPSCRRDGGFQAVQCQRGGPCWCVDARGKEIPGTRR-PGER     355
Panthera_leo                     PTKCEMERFTATSFG-HPYVPSCRRSGDYEAVQCQRAGPCWCVDAQGEEIHGTRR-QGER     355
Bos_taurus                       PTKCEVERFAATSFR-HPYVPSCHPDGEYQAAQCQQGGPCWCVDSRGQEIPGTRQ-RGEP     355
Trichechus_manatus_latirostris   PTKCEVEQFTATSFG-HPYVPTCSRNGDYQPVQCQREGPCWCVDARGKEIQGTRQ-QGGL     355
Columbia_livia                   PSKCEVERYTALRYQ-HAYVPACDADGGYTPVQCHQGGQCWCVDTKGQEVQGTKR-RGQL     362
Taeniopygia_guttata              PSRCEAERAAAQRFQ-QPLVPSCAADGGHTSLQCHPWGQCWCVDTQGHEVPGTRR-RGQP     361
Larus_michahellis                PSKCEVERYAALRYQ-HAYVPSCDADGGYAPVQCQQGGQCWCVDTKGQEVQGTKR-RGQP     362
Gallus_gallus                    PSKCEVERYAAQRYQ-HAYIPSCDTDGGYTAVQCQQGGQCWCVDTKGQEVQGTKR-RGQP     354
Struthio_Camelus                 PSKCEVERYAAIRYQ-HAYMPSCDTEGGYTPVQCQQGGQCWCVDSKGQEIQGTKR-QGQL     337
Chelonia_mydas                   PSKCELERFAAIRFR-HAYSPSCEDNGDYAPVQCQEGPCWCVDPEGREIQGTRR-QGQL     354
Alligator_mississippiensis       PSKCEVERSTAVRFG-HTYKPSCDDHGDYNPLQCQQDGQCWCVGNKGQEFQGTRR-QGQL     339
Crotalus_tigris                  PSKCEIEYFAATRFE-EDYKPLCEEIGAYKPTQCQLNGECWCVDSRGQEIPGTRR-RGQQ     358
Python_bivittatus                PSKCEIEYSTATRFE-EDYKPLCEDSGAYKPTQCQLNGECWCVDTRGQEIPGTRR-KGQQ     334
Eublepharis_macularius           PSKCETERFSAVRFQ-QSYRPSCDESGAFESTQCLPDGQCWCVDSKGKEILGTRS-HGQR     354
Xenopus_tropicalis               PSVCEVQRFTASQMG-DIYVPSCDDNGDFNPIQCQTGQQCWCVDSKGREIHGTRR-LGDS     362
Aquarana_Catesbeiana             PSACEIQRFTSSQMG-DIYVPTCDEDGDFVPVQCQANGQCWCADLTGREIYGSRL-VGKP     354
Carassius_auratus                PTPCESERAAATRAG-NGFVPSCTDNGAYVSTQCQLGGQCWCVDADGKEIFGTRQ-YGVP     356
Cyprinus_carpio                  PTPCESERAAATQAG-NSFVPSCIGNGAYVSTQCQSSGQCWCVDADGKEIFGTRQ-YGVP     356
Danio_rerio                      PSPCESERAAASQAG-NSFVPSCTDNGAYVPTQCQSGGQCWCVDTYGKEIFGTRQ-NGVP     356
Astyanax_mexicanus               PTPCESERSASSQAG-NVYVPSCDARGQYLPSQCQAGGQCWCVDSDGKEIFGTRQ-RGVP     351
Clupea_harengus                  PTPCESERSAASKAG-NVFVPSCDASGAYVPTQCQTGGQCWCVGADGKEVFGTRK-QGQP     379
Oryzias_latipes                  PSPCEEERRAALETS-SVFVPTCERGGSYSFTQCQQGGQCWCVDPSGREVPGTRQ-LGGP     339
Xiphophorus_maculatus            PTPCEEERRAALDSS-SVFVPSCEAGGRFSPKQCQQGALCWCVDPTGRQLTGPRR-PGDG     339
Stegastes_partitus               PSPCEEERRAA--AS-SVFVPSCEADGSFTSRQCQQGGQCWCVDPTGRELPGTRQ-QGDS     341
Cynoglossus_semilaevis           PTSCEEERRAATAAA-NLYVPSCERDGAYTSRQCQQGVQCWCVDATGREIPGTRQ-LGDS     340
Lepisosteus_oculatus             PTQCEREWMSATRAG-NVHVPSCDERGHYDPVQCQAGGHCWCVDSAGKEILGTRMQGGIP     351
Lampetra_fluviatilis             PTVCEVERSLADRLGGRGFAPVCSQDGSYVPTQCH-GPTCWCVDTRGDEVFGSRV-RDSV     408
Lampetra_planeri                 PTVCEVERSLADRLGGRGFAPVCSQDGSYVPTQCH-GPTCWCVDTRGDEVFGSRV-RDSV     346
Petromyzon_marinus               PTVCEVERSLAERLGGRGFAPVCSQDGSYIPTQCH-GPTCWCVDTRGDEVFGSRV-RDSV     361
```

LINKER

```
Homo_sapiens                     PSC-AEGQSCASERQQALSRLYFGTSGYFSQHDLFSSPEKRWASPRVARFATSC------     408
Pan_troglodytes                  PSC-AEGQSCASERQQALSRLYFGTSGYFSQHDLFSSPEKRWASPRVARFATSC------     408
Gorilla_gorilla                  PSC-AEGQSCAAKRQQALSRLYFGTSGYFSQHDLFSSPEKRWASPRVARFATSC------     408
Pongo_pygmaeus                   PSC-AEGQSCASKRQQALSRLYFGTSGYFSQHDLFSSPEKRWASPRVARFATSC------     408
Macaca_mulatta                   PSC-AEGRSCASKRQQALSRLYFGTSGYFSQHDLFSSPEKRWASPRVARFATSC------     408
Macaca_fascicularis              PSC-AEGRSCASKRQQALSRLYFGTSGYFSQHDLFSSPEKRWASPRVARFATSC------     408
Rattus_norvegicus                LFC-AKDQSCASERQQALSRLYFETPGYFSPQDLLS-SEDRLVPVSGARLDISC------     408
Mus_musculus                     PSC-AADQSCALERQQALSRFYFETPDYFSPQDLLS-SEDRLAPVSGVRSDTSC------     408
Cavia_porcelus                   PSC-AGDQSCSSERQQALSRLYFGPSGYFSQSNLFPAPEEEQASLGGTRPATAC------     409
Canis_lupus_familiaris           PSC-AETQLCASERRQALSRLFFGPSGYFSQHSLFLAPDGRQVSQRVARFATSC------     408
Panthera_leo                     PSC-AEDQSCASERRQALSRLHFGPSGYFSQHGLFLAPEGRPVSQRVARFATSC------     408
Bos_taurus                       PSC-AEDQSCPSERRRAFSRLRFGPSGYFSRRSLLLAPEEGPVSQRFARFTASC------     408
Trichechus_manatus_latirostris   PSC-AEDRSCTSERQQALSRLYFGPSGHFSQHNLFFSPEERQASQRVARFSSSC------     408
Columbia_livia                   PAC-GEEQGCISERRRALSRIFHGPAGYFSQHNLFSTPDMQ--SEKISGFSRPC------     413
Taeniopygia_guttata              PAC-GEEQSCISERRRALSRLLYGPAGYFSQSSLFSTPDKQ--SDKTDGFSRPC------     412
Larus_michahellis                PAC-GEELACISERRRALSRIFYGPAGYFSQADLFSTPDMQ--LEKIAGFSKPC------     413
Gallus_gallus                    PAC-GEEQICISERQRALSRIFYGPAGYFSQSDLFSIPDVQ--SEKRAGFSRPC------     405
Struthio_Camelus                 PAC-GEEQVCLSQRRQALSRIFYGPAGYFSQHSLFSTPDVQ--SEKIADFSRPC------     388
Chelonia_mydas                   PTC-GEDRACASERQQALSRLFYGPVGHFSQHSLFDTPDMK--SENIDRFSSAC------     405
Alligator_mississippiensis       PVC-GEEQTCISERRQALSKLFYGPVGHFSQHSLFNTQEMK--AEEIGRFSKSC------     390
Crotalus_tigris                  PIC-GKEPNCVSERLVSLSRLFYGPIGHFSHNNLFSATP-E--PQKLVGSLRFC------     408
Python_bivittatus                PIC-GKEPTCVSERLVSLSRLFYGPVGHFSQDNLFSATQ-E--PQKIVGSSRFC------     384
Eublepharis_macularius           PAC-GDEHACASERQRALSALFYGPVGHFSQHHLFSAAE-E--PQKTAKFSRSC------     404
Xenopus_tropicalis               PSC-NM-DDCPTKRRQALSSLFNGPTGHFGKHSLFVTQEEELGSQIN---AKFC------     411
Aquarana_Catesbeiana             VKC-DIYQDCPVKRRQALSTLFYGPTGHFVQQSLFLTQAEELGTKIG---FNYC------     404
Carassius_auratus                NCS-TGVKDCLSERRQALSRLFYGPAGD-----LFSISKDT--L----SFPGTC------     398
Cyprinus_carpio                  NCS-TGVKDCLSERRQALSRLFYGPAGDFSKSKVFSISKET--L----NFVGTC------     403
Danio_rerio                      KCS-TGVKDCLSERRQALSKLFYGPAGGFSNSNVFSSSKDT--V----SFLGTC------     403
Astyanax_mexicanus               DCG-TKGKDCYSERRLALSRLFYGPAGHLSQNNAFSSPQDT--TKKDASPFSPC------     402
Clupea_harengus                  DCS-AGLRDCHSERRQALSRLLSGPVAV--------RPADT--QQEGVTPVSVC------     422
Oryzias_latipes                  LACGSGPADCPSQRRRALSRLLSGPISPPLK-------------SSSRRLAASC------     380
Xiphophorus_maculatus            LVCGSGPADCPARRRLALSRLFSGPVRPPRA-------------S--SGSAVSC------     378
Stegastes_partitus               LVCSSGPVDCLSQRRLALSRLFSGPAEPPLE-------------TASGRSPASC------     382
Cynoglossus_semilaevis           LVCSPD-QNCFSRRRLALSQLFSGPVAPSPS--------------LSSSPPASC------     379
Lepisosteus_oculatus             DCS-DGIKDCPSERRQALSRLFYGPSGYFSQHNVFTVKQAD--GKLRASFLRLC------     402
Lampetra_fluviatilis             PNCGSSSSPCQLERQRALSHLFLGPSAPFTDDLISALPRDGSA-----SFAESPSQEKQH     463
Lampetra_planeri                 PNCGSSSSPCQLERQRALSHLFLGPSAPFTDDLISALPRDGSA-----SFAESPSQEKQH     401
Petromyzon_marinus               PNCGSSSSPCQLERQLALSHLFLGPSAPFIDDLISALPRDGSV-----SFSESPSQEMQH     416
```

| | | |
|---|---|---|
| *Homo_sapiens* | -----PPTIKELFVDSGLLRPMVEGQS-QQF-SVSENLLKEAIRAIFPSRGLARLALQFT | 461 |
| *Pan_troglodytes* | -----PPTIKELFVDSGLLRPMVEGQS-QQF-SVSENLLKEAIRAIFPSRGLARLALQFT | 461 |
| *Gorilla_gorilla* | -----PPTIKELFVDSGLLRPMVEGQS-QQF-SVSENLLKEAIRAIFPSRGLARLALQFT | 461 |
| *Pongo_pygmaeus* | -----PPTIKELFVDSGLLHPMVEGQS-QQF-SVSESLLKEAIRAIFPSRGLARLALQFT | 461 |
| *Macaca_mulatta* | -----PPTIKELFVDSGLLRPMVEGQS-QQF-SVSESLLKEAIRAIFPSRGLARLALQFT | 461 |
| *Macaca_fascicularis* | -----PPTIKELFVDSGLLRPMVEGQS-QQF-SVSESLLKEAIRAIFPSRGLARLALQFT | 461 |
| *Rattus_norvegicus* | -----PPRIKELFVDSGLLRSIAVERY-QQL-SESRSLLREAIRAIFPSRELAGLALQFT | 461 |
| *Mus_musculus* | -----PPRIKELFVDSGLLRSIAVEHY-QRL-SESRSLLREAIRAVFPSRELAGLALQFT | 461 |
| *Cavia_porcelus* | -----PSQIKQLFVDSGLLHLMTKGQD-PQA-SAFQSLLGEATRAIFPSRELARLALQFT | 462 |
| *Canis_lupus_familiaris* | -----PPLIKELFVDSGILRPMVEGQD-KQF-SASETLIREAIGAIFPSRELARLALQFT | 461 |
| *Panthera_leo* | -----PPLVKELFVDSGILHPMGEGQG-KQF-SASETLLREAIGAIFPSRELARLALQFT | 461 |
| *Bos_taurus* | -----PPSIKELFLDSGIFQPMLQGRD-TRF-VAPES-LKEAIRGLFPSRELARLALQFT | 460 |
| *Trichechus_manatus_latirostris* | -----PPLFRELFVDSGLLRPMVEGQD-KQF-PASQSLLKEAIRSIFPSRELARLALQFT | 461 |
| *Columbia_livia* | -----PPYFKELFLDSGLSSPVVQSPYVSQI-PGLEITLSEAITGMFPSRELAEVVLQFT | 467 |
| *Taeniopygia_guttata* | -----PPSFKELFLDSGLSSPLAQSPFASQT-PELETILSEAITGMFPSRELARVALQFT | 466 |
| *Larus_michahellis* | -----PPSFKELFLDSGLSSPVAQSPYASQI-PGLETTLSEAITGMFPSRELAQVALQFS | 467 |
| *Gallus_gallus* | -----PPSFKELFLDSGLSSPIMQSPYASQI-PGLESILSEAISGMFPSRELAQVALQFT | 459 |
| *Struthio_Camelus* | -----PPSFKALFLDSGLSSPITQSQYASQI-LQLESTLSEAVTGMFPSRELAQVALQFT | 442 |
| *Chelonia_mydas* | -----PPSFKELFVDSGLLSPITEKQDASQL-FALESILSDVVRGMFPSRQLAQVALQFT | 459 |
| *Alligator_mississippiensis* | -----PPSFKGLFVDSGLLSPLTERPGLNQQ-LELESILSEAVRGMFPSRELAQVALQFT | 444 |
| *Crotalus_tigris* | -----PFSFRDFFVGSTVLSSTSGKTSAVQL-LTFETLLGEAVGGMFPSRELAQIALRFT | 462 |
| *Python_bivittatus* | -----PLSFKDFFVGSRVLSPISGKNSASQL-LTFETLLGEAVGGMFPSRNLAQVALQFT | 438 |
| *Eublepharis_macularius* | -----PPSLKELFVDSGLLSSVIDTPDASQL-LALESILSEAIRGLFPSRELAQVALQFT | 458 |
| *Xenopus_tropicalis* | -----SSYLTELFGKSGLLFPIMQKVR-----FQLGSFVRDMVAGLFPSKELMQIALNFV | 461 |
| *Aquarana_Catesbeiana* | -----PSYIVETFSDSGLLFELATSVG-----LPLKSFIVEMIKGIFESKEQIQLALKFT | 454 |
| *Carassius_auratus* | -----SPEFQELLANSGLLQSLPELER-----PKVGDILSEVLQGMFPSGALALKALAHT | 448 |
| *Cyprinus_carpio* | -----SPEFQELLANSGLLQSLPELEH-----PNVGDMLAEVLQGMFPSGALALKALAHA | 453 |
| *Danio_rerio* | -----SPEFQELLANSGLLQSLPELER-----PKVTNILADVLQGMFPSGALALKALALA | 453 |
| *Astyanax_mexicanus* | -----SPEFQELLANSGLARSLPESER-----SDIGEIMAELVQGMFPTGALALKALSLT | 452 |
| *Clupea_harengus* | -----SSEFQELLAKSGLLRSLPESDR-----SDIGDILAELVHGMFPTGALALKALTLT | 472 |
| *Oryzias_latipes* | -----SSLLQPLRD----LL--PEE---ADE-TNFLFRLVEVLDGLFPSVGGALQALN-G | 424 |
| *Xiphophorus_maculatus* | -----VSLLRPLRV----L----GY---SDP-TSFLARLVEVLQGLFPSVGGALEALT-R | 420 |
| *Stegastes_partitus* | -----LSLLRPLRN----LL--PAE---MDP-TSFLSRLVQVLQGLLPSVGGALQALT-R | 426 |
| *Cynoglossus_semilaevis* | -----LSLLRPLQP----LL--PTQ---ENP-TSHLSLMLEVFHGLFPSVGEAFKALV-S | 423 |
| *Lepisosteus_oculatus* | -----NPEVRELFVKSGLLLTLPELTN-----SSFGTVVGEIFQGIFPSREMALKALRFA | 452 |
| *Lampetra_fluviatilis* | ELWLHWRALREQLLDGGLSEILSQASRLFPASGQGSRLLGDLMGGIFPTQELALTAAGAN | 523 |
| *Lampetra_planeri* | ELWLHWRALREQLLDGGLGEILSQASRLFPASGQGSRLLGDLMGGMFPTQELALTAAGAN | 461 |
| *Petromyzon_marinus* | ELWLHWRALREQLLDGGLGEILSQASRLFPASGQASRLLGDLVGGIFHTQELALTAAGAN | 476 |

| | | |
|---|---|---|
| *Homo_sapiens* | TNPKRLQQNLFGGKFLVNVGQFNLSGALGTRGTFNFSQFFQQLGLASFLNGGRQEDLAKP | 521 |
| *Pan_troglodytes* | TNPKRLQQNLFGGKFLVNVGQFNLSGALGTRGTFNFSQFFQQLGLASFLNGGRQEDLAKP | 521 |
| *Gorilla_gorilla* | TNPKRLQQNLFGGKFLVNVGQFNLSGALGTRGTFNFSQFFQQLGLASFLNGGRQEDLAKP | 521 |
| *Pongo_pygmaeus* | TNPKRLQQNLFGGKFLVNVGQFNLSGALGTRGTFNFSQFFQQLGLASFLNGGRQEDLAKP | 521 |
| *Macaca_mulatta* | TNPKRLQQNLFGGKFLVNLGQFNLSGALGTRGTFNFSQFFQQLGLASFSNGGRLEDLAKP | 521 |
| *Macaca_fascicularis* | TNPKRLQQNLFGGKFLVNLGQFNLSGALGTRGTFNFSQFFQQLGLASFSNGGRLEDLAKP | 521 |
| *Rattus_norvegicus* | TNPKRLQQNLFGGTFLVNAAQLNLSGALGTRSTFNFSQFFQQFGLPGFLVRDRATDLAKL | 521 |
| *Mus_musculus* | TNPKRLQQNLFGGTFLANAAQFNLSGALGTRSTFNFSQFFQQFGLPGFLNRDRVTTLAKL | 521 |
| *Cavia_porcelus* | TNPKRLQQNLFGGTFLVNLGQFNLSGALGMRGTFNFSQFFQQLGLSGFLNRGGAVEPVKL | 522 |
| *Canis_lupus_familiaris* | TDPTRLQQNLFGGKFLVNVGRFNLSGAIGTRGTFNFSQFFQQFGLPGFQNSETD-LA-QP | 519 |
| *Panthera_leo* | VDPKRLQQNLFGGKFLVNVGQFNLSGALGTKGTFNFSQFFQQFGLPGFQNGGTD-LAAKA | 520 |
| *Bos_taurus* | TNAKRLQQNLFGGRFLVKVGQFNLSGALGTRGTFNFSHFFQQLGLPGFQDGRALADLAKP | 520 |
| *Trichechus_manatus_latirostris* | ADPKRLQQNLFGGNFLANVGQFNVSGAIGTRGTFNFSQFFQQLGLPGFKNGEGFVDVAKQ | 521 |
| *Columbia_livia* | ANPKRFQENLFGGRFLKNLIQFNFTGALGTSGKYSIGQFFLQEDVVDGDNIQSLLELA-- | 525 |
| *Taeniopygia_guttata* | ANPKRFQENLFGGRFLKNLIQFNFTGVLGTNGKYSIGQFFPQEDLSEMGNGQGPLESAEA | 526 |
| *Larus_michahellis* | TNPKRFQENLFGGRFLKNLIQFNFTGALGTSGKYSIRQFFPQEDVAERDNNSSLLESAEA | 527 |
| *Gallus_gallus* | ANPKRFQENLFGGKFLKNMIQFNFTGALGTSGKYSIGQFFTQEDATGRNGSQGLLQSAAA | 519 |
| *Struthio_Camelus* | SNPKRFQENLFGGRFLRNLIQFNFTGALGTSGKYSIGQFFPQESATERDNGQGLLESAGA | 502 |
| *Chelonia_mydas* | TNPKRFQENLFGGKFLKNLIQFNFTGALGTRGKFSIGQFFQQGSLMAMDNGGGFAEHPLG | 519 |
| *Alligator_mississippiensis* | SNPKRFQENLFGGKFLKNLIQFNFTGALGTRGKFNIGQFSQQGGM---DSGESVAKLAEE | 501 |
| *Crotalus_tigris* | SNPKRFQENLFGGMFLKNLVQFNFTGALGPNSKFNLDKYFQQVGMR-----SGFSELAGQ | 517 |
| *Python_bivittatus* | NSPKRFQENLFGGMFLKNLVQFNFTGALGPNSKFNLDKYFQQVGMR-----SGFSELVHQ | 493 |
| *Eublepharis_macularius* | SNHKRFQENLFGGKFLKNLVQFNFTGALGTNGKFNLEQFFQQIGRS-----DGFAEPTRQ | 513 |
| *Xenopus_tropicalis* | RNPKNFQENLFGGKYLKNLGSFNFSGAVGTNSKFNFSGFFQQIGLTGMYSGGNFRELAKL | 521 |
| *Aquarana_Catesbeiana* | SNPIRFQQNLFGGKYLKNLGSFNFTGAIGAKNKFSFSDFFKQIGLTGMYSGGNFKELAKL | 514 |
| *Carassius_auratus* | TNPKRLQENLFGGKFLKNAGNFNFSSTVGNSGTLNFSQVFRQVGLM--QAVTDLMQLAKT | 506 |
| *Cyprinus_carpio* | PNPKRLQENLFGGKFLKNAGNFNFTGTVGNSGTLSFSKIFSQVGLM--QAVTDLMQLAKT | 511 |
| *Danio_rerio* | QNPKRLQENLFGGKFLKNAGNFNFSGTVGDSGTLSFRQIFSQVGLT--QVVTDLMQLAKT | 511 |
| *Astyanax_mexicanus* | SNPKRLQENLFGGKFLKNAGNFNFTGTVGSRGTVSFSQAFLQVGLS--QDAKDLTKLAKI | 510 |
| *Clupea_harengus* | PNPKRLQENLFGGKFLKNAGNFNFSGAVGSRGTLSFSQTFSQVGVT--LSGAQLVQLAKI | 530 |
| *Oryzias_latipes* | SSPRFLQENLFGGDFLKSAGALNFSGEVGDQGTISPDCLSPQ------------------ | 466 |
| *Xiphophorus_maculatus* | SPARRLQENLFGGKFLQNAAAFNYTGVLGPRGAVGLDQLSAQ------------------ | 462 |
| *Stegastes_partitus* | SSPQRLQENLFGGKFLKSAAAFNFSGSVGTEGALGLGRLSSQ------------------ | 468 |
| *Cynoglossus_semilaevis* | VSPHLLQEELFGGEFLTSVATFNLSGAVGARGAVELL----------------------- | 460 |
| *Lepisosteus_oculatus* | TNPKRLQENLFGGKFLKNIRYYNFTGAVGPSGTFNFNETFQQVGLK--QFSGDFVQLAKI | 510 |
| *Lampetra_fluviatilis* | VQPGRIAEMLFGGGFLKNLKFFNFTGAVGGRGTFNFSKVFGQVGLTGMYDGANFEELARL | 583 |
| *Lampetra_planeri* | VQPGRIAEMLFGGGFLKNLKFFNFTGAVGGRGTFNFSKVFGQVGLTGMYDGANFEELARL | 521 |
| *Petromyzon_marinus* | VQPGRIAEMLFGDRFLKNLKFFNFTGAVGGRGTFNFSKVFGQVGLTGMYDGANFEELARL | 536 |

```
Homo_sapiens                     LSVGLDSNSSTGTPEAAKKDGTMNKPTVGSFGFEINLQENQNALKFLASLLELPEFLLFL   581
Pan_troglodytes                  LSVGLDSNSSTGTPEAAKKDGTMNKPTVGSFGFEINLQENQNALKFLASLLELPEFLLFL   581
Gorilla_gorilla                  FSVGLDSNSSTGTPEAAKKDGTMNKPTVGSFGFEINLQENQNALKFLASLLELPEFLLFL   581
Pongo_pygmaeus                   LSVGLDSNSSTGTAEAAKKDGTMNKPTVGSFGFEINLRENQNALKFLASLLELPEFLLFL   581
Macaca_mulatta                   VSVGLDSNSSTGTPEAAKKDVAMNKPIVGGFGFEINLQENQNALKFLASLLELPEFLLFL   581
Macaca_fascicularis              VSVGLDSNSSTGTPEAAKKDVAMNKPIVGSFGFEINLQENQNALKFLASLLELPEFLLFL   581
Rattus_norvegicus                LPVSLDSSPTPVPLRVPEKRVAMNKSVVGTFGFKVNLQENQDALKFLVSLMELPEFLVFL   581
Mus_musculus                     LPVRLDSSSTPETLRVSEKTVAMNKRVVGNFGFKVNLQENQDALKFLVSLLELPEFLVFL   581
Cavia_porcelus                   PSVVLDVNSA-------TKSLSMNKTIVGSFGFQINVQENQNALKFIVSLLELPEFLLFL   575
Canis_lupus_familiaris           LSVGLDSQPATAPPEASKEGTAMNVPVVGSFGFQINLQENQNALTFLASLLELPEFLLFL   579
Panthera_leo                     LSLGLDSNPAAEPPEASVKGAARNRPMVGSFGFEINLQENQNALTFLASLLELPEFPLFL   580
Bos_taurus                       LSVGLNSNPASEAPKASKIDVALRKPVVGSFGFEVNLQENQNALQFLSSFLELPEFLLFL   580
Trichechus_manatus_latirostris   LSVRPDANLATGTPEASEK--AMNLPLVGSFGFEINLRENQNALKFLASLLELPEFLVFL   579
Columbia_livia                   -------------EVSKESSILSKPLVGNFGRTVTLQDNQNKMKFLSSVLELPAFFTFL   571
Taeniopygia_guttata              FSL-----------EASKGSSILSKPLVSSFGRTVTLQDNQNLMKFLSYVLELPEFFTFL   575
Larus_michahellis                LSL-----------EVSQGRSILNKPLVGSFGRTITLQGNQNMMKFLSSVLELPEFFTFL   576
Gallus_gallus                    FSL-----------EVSKGSSILSQALVGSFGRTVTLQDNQNIMKFLSSVLELPEFFTFL   568
Struthio_Camelus                 FSL-----------EVSN-SSVFSQPLMGSFGRTITLQDNQNRIKFLAAVLELPAFFTFL   550
Chelonia_mydas                   LSL-----------ETSKESFNLSQPLVDSFGRTVSLQNNQNGVEFLASLLELPEFFAFL   568
Alligator_mississippiensis       PSL-----------ETSEESFITSKPLVDSFGRTVSLQDNQNAMKFLASVLELPEFFTFL   550
Crotalus_tigris                  LLQ-----------GSSQEKFNRSQSRVDSFGRKVSLEDNQKAIQFLASLLETPEFFSFL   566
Python_bivittatus                LFQ-----------EDPQQKFNLSQSLVDSFGRMVNLQDNQNAIKFLASLLEAPEFFTFL   542
Eublepharis_macularius           IAS-----------K--VPNFSLSQPLINSFGQTVNLQDNQNAVKFLASVLEAPEFFTFL   560
Xenopus_tropicalis               FSAAEDSYLTKDSLNISKPVFNLNQPIQGDFGRIVNLRENQDLVGVFASVLELEEFALFL   581
Aquarana_Catesbeiana             FSSEEDSYLSRD-ANISKPFFNLNQTILASFGRTVNLQENQKRVGIFLSILERKEFAALL   573
Carassius_auratus                VTD-----------DS--ASLKLDQEISDAFGRSVNLKNNQDMIKLVSMALENEQFLTVL   553
Cyprinus_carpio                  VTD-----------DS--ASLKLDQEISDAFGRSVNLKNNRDMIKLVSMALENEQFLTVL   558
Danio_rerio                      FTD-----------SS--ENLNLDQEISDAFGRSVNLKNNREMIKLVSMALENTHFLSTL   558
Astyanax_mexicanus               FSV-----------DS--TGPNLDREVSDAYGRSVNLQRNRDLIKLIGRALENEQFFTTL   557
Clupea_harengus                  FSP-----------GQVMGGETLDREITDAFGRSVSLKNNQDLIQLLGATLESEPFLATL   579
Oryziaz_latipes                  ---------------------------------NPRLHENQNLVQFVSKALEDPAFFSVL   493
Xiphophorus_maculatus            ---------------------------------S--LQENRDLVRSVSRALEDPAFLSAL   487
Stegastes_partitus               ---------------------------------SAVLQKNRDIVQSISMALEDAAFLSAL   495
Cynoglossus_semilaevis           -----------------------------------YSQKNGDLVLSVSRALEDPAFLSSV   485
Lepisosteus_oculatus             FSS-----------ETDSKPLYLDQKIIDSFGRNINLKDNQNLVELVGMILENEQFVTTL   559
Lampetra_fluviatilis             FAPGEDSYLTRGSSNFSRESFNFDQSIDDPFARATNLERNRNVLGFVVTLLEDPRFYGLV   643
Lampetra_planeri                 FAPGEDSYLTRGSSNFSRESFNFDQSIDDPFARATNLERNRNVLGFVVTLLEDPRFYGLV   581
Petromyzon_marinus               FAPGEDSYLTRGSSNFSRESFNFDQSIDDPFARATNLERNRNVLGFVVTLLEDARFYGLV   596
```

TG type 1-5

```
Homo_sapiens                     QHAISVPE-DVARDLGDVMETVLSSQT------------CEQTPERLFVPSCTTE-GSYE   627
Pan_troglodytes                  QHAISVPE-DVARDLGEVMETVLSSQT------------CEQTPERLFVPSCTTE-GSYE   627
Gorilla_gorilla                  QLAISVPE-DVARDLGDVMETVFSSQT------------CEQTPERLFVPSCTTE-GSYE   627
Pongo_pygmaeus                   QHAISVPE-DVARDLGDVMEMVLSSQT------------CEQTPERLFVPSCTTE-GSYE   627
Macaca_mulatta                   QHAISVPE-DVARDLGDVMETVLRSQT------------CEQTPERLFVPSCTTE-GSYE   627
Macaca_fascicularis              QHAISVPE-DVARDLGDVMETVLRSQT------------CEQTPERLFVPSCTTE-GSYE   627
Rattus_norvegicus                QRAVSVPE-DRARDLGDVMEMVFSAQA------------CKQTSGRFFVPSCTAE-GSYE   627
Mus_musculus                     QRAVSVPE-DIARDLGDVMEMVFSAQA------------CKQMPGKFFVPSCTAG-GSYE   627
Cavia_porcelus                   QRALSVPE-DVARDVGDVMGMVLSSQA------------CGQSPGRLFVPSCTEE-GGYE   621
Canis_lupus_familiaris           QHAISVPE-DIAKDLGDVMEMVLSSQD------------CEQTPGNLFVPQCSAE-GDYE   625
Panthera_leo                     QHAISVPE-DIAKDLGDVMQTVLSSQG------------CGQTPGSLFVPLCSAE-GGYE   626
Bos_taurus                       QHAISVPE-DIARDLGDVMEMVFSSQG------------CGQAPGSLFVPACTAE-GSYE   626
Trichechus_manatus_latirostris   QHAISVPE-DVARDLGDTMEMVLNSQT------------CEPPSTKLFVPSCTAE-GNYE   625
Columbia_livia                   QQVISVPR-SIAEDLGEVVKIALGSKD------------CGEEPRDLFVPTCTKE-GRYE   617
Taeniopygia_guttata              QQVISVPE-SVAEDLGEVVKLALGSGG------------CGEEPRDPFVPTCTKE-GRYE   621
Larus_michahellis                QQVISVPK-SVAEDLGEVVKLALGSEG------------CDELPKGLFVPTCTKE-GRYE   622
Gallus_gallus                    QQVISVPR-SIAEDLGEVVKLALGSRD------------CGEEPRDLFVPTCTKE-GRFE   614
Struthio_Camelus                 QQVISVPK-SIAENLGEVVKLALGSQD------------CGEEARDLFVPTCTKE-GRYE   596
Chelonia_mydas                   QHVISVPE-NIAEDLGEVVKMALKSKD------------CTEQPMDLFVPTCTKE-GRYE   614
Alligator_mississippiensis       QHVIFVPE-DISEDLGEVVRIVLRSKD------------CTEESSNLFVPTCTKE-GRYE   596
Crotalus_tigris                  QHVISVPE-SVTRDLGKVVRILLKARD------------CTEEDKDALVPDCTVD-GNYK   612
Python_bivittatus                QHVISVPE-SVTKDLGNVVRILLKSKD------------CTEETNDIFVPDCTKD-GKYK   588
Eublepharis_macularius           RHIISIPE-SIAEDLGDIVKIVLRSQD------------CKEQAKNLFVPECTEE-GKYK   606
Xenopus_tropicalis               RNVISVPT-NVAQSAAEAVQVVVQSND------------CEKRSSEIYIPVCKED-GRYN   627
Aquarana_Catesbeiana             RDITSMPP-SIAEDISEAMKIIMASKD------------CRNRK-EALVPTCTKE-GRYE   618
Carassius_auratus                REAIKQLKAKESSQLGQLFHAIFQKSDV----------CKSVTSTSTLYVPQCLED-GRYQ   603
Cyprinus_carpio                  REAIKQLKAEESSQLGQLFHAIFQKSDV----------CKSVTSTSTLYVPQCTED-GRYQ   608
Danio_rerio                      RVAIKQLKAEETTQLGSLFRDIFQKSDV----------CKPMSSSSAPYLPQCTED-GLYQ   608
Astyanax_mexicanus               REAITLLKAEDSDQLGPLFQAVFQNEDV----------CSSPASPSSLYLPQCTED-GQYQ   607
Clupea_harengus                  REAITLLKAEDSTQLGALFRAVFQGSKP----------D-MCQSPSAAYVPQCDEQ-GQFE   628
Oryziaz_latipes                  KNTLMSS--SSSASLYQVLAPLVTSCSE----------E-EEDEAADIFIPRCTPG-GAFQ   540
Xiphophorus_maculatus            RQTLMGLSGAPSGSLQQVLDEVLRSCSE----------E-EDD-SSSLFVPRCSPS-GGYQ   535
Stegastes_partitus               QNTLTGLSSSQSASLQKVLTPLLRSCSN----------D-QED-EGSIFVPSCTSS-GGFQ   543
Cynoglossus_semilaevis           RQALTGLS--SSLRLQEVLTPLLHYCTA----------E-EQKYSEAVVPSCTSS-GAFQ   532
Lepisosteus_oculatus             RDIISFLRVEDSEDFSSFFQALLQNTEP----------EVCERDSSVLFVPQCTKR-GQYE   609
Lampetra_fluviatilis             RDIAPLIG-LSGGDVGMNALTFLRVAEEDRQTQNAADADDGPRASRPFVARCGEDGGAYE   702
Lampetra_planeri                 RDIAPLIG-LSGGDVGLNALTFLRVAEEDRPTQNAADAYDGTRASRPFVARCGEDGGAYE   640
Petromyzon_marinus               RDITPLIG-LSGGDVGLNALTFLRMAEEDRPTQNAADAADGPPASRPFVARCGEDGGAYE   655
```

```
                                                                     TG type 1-6
Homo_sapiens                      DVQCFSGECWCVNSWGKELPGSRVRG-GQPRCPTDCEKQRARMQSLMGSQPAGSTLFVPA   686
Pan_troglodytes                   DVQCFAGECWCVNSWGKELPGSRVRG-GQPRCPTDCEKQRARMQSLMGSQPAGSTLFVPA   686
Gorilla_gorilla                   DVQCFAGECWCVNSWGKQLPGSRVRG-GQPRCPTDCEKQRARMQSLMGSQPAGSTLFVPA   686
Pongo_pygmaeus                    DVQCFAGDCWCVNSWGKELPGSRVRG-GQPRCPTDCEKQRARMQSLMGSQPAGSSLFVPA   686
Macaca_mulatta                    DVQCFAGECWCVDSWGKELPSSRVRG-GQPRCPTDCEKQRARMQSLMGSQPAGSSLFVPA   686
Macaca_fascicularis               DVQCFAGECWCVDSWGKELPSSRVRG-GQPRCPTDCEKQRARMQSLMGSQPAGSSLFVPA   686
Rattus_norvegicus                 DIQCYAGECWCVNSQGKEVEGSRVSG-GHPRCPTKCEKQRAQMQNLAGAQPAGSSFFVPT   686
Mus_musculus                      DIQCYAGECWCVDSRGKELDGSRVRG-GRPRCPTKCEKQRAQMQSLASAQPAGSSFFVPT   686
Cavia_porcelus                    DVQCFAGDCWCVDFWGKEIPNSRVRG-GQPRCPTECEKQRARMQSLIRNQPAGASVFVPS   680
Canis_lupus_familiaris            DVQCFAGECWCVDSRGKELAGSRVRG-GRPRCPTECEKQRALMRSLSGSLPAGASLFVPS   684
Panthera_leo                      DVQCFAGDCWCVDSRGKELSGSRVRG-GRPRCPTECEKQRARMQTLEGSQPAGSSLLVPS   685
Bos_taurus                        EVQCFAGDCWCVDAQGRELAGSRVRG-GRPRCPTECEKQRARMQSLLGSQPAGSSLFVPA   685
Trichechus_manatus_latirostris    DVQCSAGECWCVDSRGKELLGSRVRG-RRPRCPTECEKQRAQMQNLMGSQPAGSSLSVPS   684
Columbia_livia                    EVQCYAGECWCLDTSGKEVPGSRVPG-KRPRCPTDCEKQRRNLQNLKQTLPAGSDLFIPS   676
Taeniopygia_guttata               EVQCFAGECWCLDTSGKEVPGSRRRD-KRPKCPTECEKQRRNLKNLKESLPAGSDLFIPS   680
Larus_michahellis                 QVQCYAGECWCLDTSGKEVPGSRVQG-KRPRCPTDCEKQRRNLQNLKQTLPAGSDLFIPS   681
Gallus_gallus                     EVQCYAGECWCLDTSGKEVPGSRVQG-ERPRCPSDCEKQRRNLQNLKQSLSAGSDLFIPS   673
Struthio_Camelus                  EIQCYAGECWCLDTQGKEVPGSRVRG-KHPKCPTDCEKQRSSLENLQRSQPAGSDLFIPS   655
Chelonia_mydas                    EIQCYAATCWCVDSRGKEVPGSRVLG-KRPRCPTACEKQRESLQSLKQSQPAGSDLFVPS   673
Alligator_mississippiensis        EIQCYAAECWCVDSQGREVPGSRVQG-KRPRCATTCEKQRSRLQRLKQSQPAGSDPFIPT   655
Crotalus_tigris                   EIQCNRGECWCVDLRGKEIPGSRVRG-TNPRCPTVCEKQRASLQFLIRDQPAGSEMFVPS   671
Python_bivittatus                 EIQCNRGECWCVDLRGKEIPGSRVQG-THPRCPTMCEKQRASLQLLIRDQPAGSELFVPS   647
Eublepharis_macularius            DIQCYAGDCWCVDSNGKEISGSRVHG-THPKCPTLCEKQRDSLQQLKRIQPAGSDLFVPS   665
Xenopus_tropicalis                DIQCSKTECWCVDDQGREIDRTRTQG-KHPRCPTKCEKERYRQTVVRESLPPGAELFIPA   686
Aquarana_Catesbeiana              EIQCDNSECWCVDDKGLEIPGSRTTD-KKPKCPSKCEKERESQILLRKSQPAGSDLFIPA   677
Carassius_auratus                 EVQCQDSECWCVDSRGLEITGSRITG-SRPRCSSQCEKERQMAIAVKASSSAGSEVFIPK   662
Cyprinus_carpio                   EVQCQGSECWCVDSRGLEITGSRTTG-SRPRCSSQCEKERQMAIAVKASSSAGSEVFIPK   667
Danio_rerio                       DVQCQGSECWCVDSRGLEVPGSRTTG-SRPRCPSQCEKERQMAIAVKASSSAGSEVFIPK   667
Astyanax_mexicanus                AVQCQGSECWCADSQGQEVMGSRSIG-YRPRCPSRCEKERMMAIAVKASSSAGSEVFIPK   666
Clupea_harengus                   EVQCTGSECWCVDSEGREVKGSRTVG-SRPRCPSQCETERDAVVTAKASMSAGAQIFIPK   687
Oryzias_latipes                   EVQCRRSQCWCVDPQGQEVEGSRVAG-RPSRCPSLCERARVNALKAKANMAAGAKIHIPV   599
Xiphophorus_maculatus             PVQCRDAECWCVDPQGQEAPGSRTAG-RPARCPTSCETARATAMKVKSNMAAGAEIHIPA   594
Stegastes_partitus                DVQCGGADCWCVDSVGQEVAGSRTAG-HRPRCPTRCERARMTALKVRSNLAAGAEIHIPA   602
Cynoglossus_semilaevis            EVQCHSGECWCVDSRGQEVTGSRTSG-QRPRCPSRCEKEQAAALRVRGSLAAGAELHIPA   591
Lepisosteus_oculatus              EVQCLGSTCWCVDPQGKEIPGSRSEG-LKPKCPSKCEKERQSAILVKASQSAGSELFIPK   668
Lampetra_fluviatilis              PVQCRGAACWCVDAHGREVAGTRAVGGVPPRCPSACEGERARALLARRGRPVGTPLFVPE   762
Lampetra_planeri                  PVQCRGAACWCVDAHGREVAGTHAVGGVPPRCPSACEGERARALLARRGRPVGTPLFVPE   700
Petromyzon_marinus                PVQCHGAACWCVDAHGREIAGTRAVGGAPPRCPSACEGERARALLARRGRPVGTPLFVPE   715

                                                                             TG type 1-7
Homo_sapiens                      CTSEGHFLPVQCFNSECYCVDAEGQAIPGTRSAIGKPKKCPTPCQLQSEQAFLRTVQALL   746
Pan_troglodytes                   CTSEGHFLPVQCFNSECYCVDAEGQAIPGTRSATGKPKKCPTPCQLQAEQAFLRTVQALL   746
Gorilla_gorilla                   CTSEGHFLPVQCFNSECYCVDAEGQAIPGTRSAIGKPKKCPTPCQLQAEQAFLRTVQALL   746
Pongo_pygmaeus                    CTSEGHFLPVQCFNSECYCVDAEGQAIPGTRSAIGKPKKCPTPCQLQAEQALLRTVQALL   746
Macaca_mulatta                    CTSEGYFLPVQCFNSECYCVDAEGQAIPGTRSAMGKPKKCPTPCQLQAEQAFLRTVQALL   746
Macaca_fascicularis               CTSEGYFLPVQCFNSECYCVDAEGQAIPGTRSAMGKPKKCPTPCQLQAEQAFLRTVQALL   746
Rattus_norvegicus                 CTSEGYFLPVQCFNSECYCVDAEGQVIPGTQSTIGEPKLCPSVCQLQAEQAFLGVVGVLL   746
Mus_musculus                      CTREGYFLPVQCFNSECYCVDTEGQVIPGTQSTVGEAKQCPSVCQLQAEQAFLGVVGVLL   746
Cavia_porcelus                    CSTEGHFLPVQCFNSECYCVDSEGRAIPGTQSPLGEPKQCPSLCQLRAEQAFLGMVRTLL   740
Canis_lupus_familiaris            CTSEGHFLPVQRFNSECYCVDTNGQAIPGTRSLPGEPKQCPTPCQLQAEQAFLGIVQVLG   744
Panthera_leo                      CTSEGHFLPVQCFGSECYCVDPKGQPIPGTRSVLGEPKQCPTACQLQAEQAFLRTVRALG   745
Bos_taurus                        CTSKGNFLPVQCFNSECYCVDTEGQPIPGTRSALGEPKKCPSPCQLQAERAFLGTVRTLV   745
Trichechus_manatus_latirostris    CTSEGHFLPVQCSNSECYCVDTEGQAIPGTQSITGEPKQCPTPCQLQAEQAFLGTVQALL   744
Columbia_livia                    CTEDGDFLPLQCYGTNCFCVDLNGKTIPGISGKAGKPMQCPSACQVRAGQEFLKAVKVML   736
Taeniopygia_guttata               CTEDGDFLPLQCYGTNCFCVDLNGKTIPGIRGEAGNPMKCPSACQVAAGQEFLRAVKLLL   740
Larus_michahellis                 CTEDGDFLPLQCYGTNCFCVDLDGKTIPGIRGKAGKPMQCPSACQVTAGQEYLKAVKLLL   741
Gallus_gallus                     CTKNGDFLPLQCYGTNCFCVDLNGKTIPGIRGNIGKPIQCPSACQITAGQEFLKTLKLLL   733
Struthio_Camelus                  CTKDGDFLPLQCYGTNCFCVDLKGKTIPRIRGDTGEPVQCPSTCQVTAGQEFLKTVKLLL   715
Chelonia_mydas                    CTKEGGFLPVQCHGTNCICVDLEGRAIPGTRKAGEPMQCPSACQVIAAQVFLRTVQLLL   733
Alligator_mississippiensis        CTEEGDFLPVQCHGPDCFCVDLDGRTIPGTKRKAGKPMQCPSPCQLTAGQVFLETVQLLL   715
Crotalus_tigris                   CTQEGKFLPVQCHGKNCFCVNSDGIPVPGISTNSGDPIQCPSDCHLAAGQAFLRSAQQFL   731
Python_bivittatus                 CTEEGKFLPVQCHGKNCFCVSSEGIPVPGIGSNSGEPIQCPSDCHLAGGQAFLRSVQQFL   707
Eublepharis_macularius            CTPEGKFLTVQCHGRNCFCVNSEGRTIPGMA-NTIQPIQCPSDCQLAGGEAFLSTVRQLL   724
Xenopus_tropicalis                CDLEGHFLTVQCTGKLCFCVDLEGRIIPGTQKVSGDDIQCPSLCQLAAGNAFLQSTMSFL   746
Aquarana_Catesbeiana              CDQNGSYRAVQCTGKHCFCVNLEGRNIPGTQKLSGENVQCPSYCQLAASNAFLQAANSLF   737
Carassius_auratus                 CETDGAYVALQCLGKSCFCVDRSGTKLSI--QSSGSSVQCPTSCQATATQQFLSTVHSVF   720
Cyprinus_carpio                   CETDGAYVALQCLGKSCFCVDHSGTKLSL--QSSGSSVQCPTSCQATATQQFLSTVHSVF   725
Danio_rerio                       CETDGAYVARQCLGKSCFCVDRSGTKLGI--QSSGSSLQCPTSCQAIAIQQFQSTVRSLL   725
Astyanax_mexicanus                CEEDGEYVSLQCLGKSCFCLDQRGVRYST--LGSGSSLQCPTDCQITASQYFMNTVNSLL   724
Clupea_harengus                   CEESGAYVQRQCQGQSCFCVDRSGSRLST--SAVGTPRQCPTDCQTSAAEAFLQTVRSAL   745
Oryzias_latipes                   CSEDGKFLPLQCVGSRCFCVDAEGRTVTSSQ-TAGGALRCPESV----------------   642
Xiphophorus_maculatus             CSEDGDFLPLQCIADRCFCVDGKGRKMAAGP--TGGAVTCPENK----------------   636
Stegastes_partitus                CSEGGDFLPLQCVGSQCFCVDAEGKTMIAGP--TGGAVTCSEKT----------------   644
Cynoglossus_semilaevis            CSENGDFLPLQCVGSRCFCVDAEGKRTAE---PSGNALTCAEMK----------------   632
Lepisosteus_oculatus              CEADGSFMPLQCSGRTCSCMDSAGMKITR--TTLGQPIQCPSVCQLAGAQKFLSTVQSLL   726
Lampetra_fluviatilis              CDGAGDYRPVQCSGDRCFCVGAGGAEVPGTWRPLGDPVTCPTPCQVAAAGELLQQLRHLG   822
Lampetra_planeri                  CDGAGDYRPVQCSGDRCFCVGAGGAEVPGTWRPLGDPVTCPTPCQVAAAGELLQQLRHLG   760
Petromyzon_marinus                CDGAGDYRPVQCSGDRCFCVGAGGAEVPGTGRPLGDPVTCPTPCQAAAASELLQQLRHLG   775
```

```
Homo_sapiens                    S--NSSMLPTLSDTYIPQCSTDGQWRQVQCNGPPEQVFELYQRWEAQNK-GQDLTPAKLL    803
Pan_troglodytes                 S--NSSMLPTLSDTYIPQCSTDGQWRQVQCDGPPEQVFELYQRWEAQNK---DLTPAKLL    801
Gorilla_gorilla                 S--NSSMLPTLSDTYIPQCSTDGQWRQVQCDGPPEQVFELYQRWEAQNK-GQDLTPAKLL    803
Pongo_pygmaeus                  S--NSSMLPTLSDTYIPQCSPDGQWRQVQCDGPPEQVFEWYQRWEAQNK-GQDLTPAKLL    803
Macaca_mulatta                  P--NSSMLPTLSNTYIPQCSADGQWRQVQCDGPPEQVFEWYQRWEAQNK-GQELMPAELL    803
Macaca_fascicularis             S--NSSMLPTLSDTYIPQCSADGQWRQVQCDGPPEQVFEWYQRWEAQNK-GQELMPAELL    803
Rattus_norvegicus               S--NSSMVPPISSVYIPQCSTSGQWMPVQCDGPHEQVFEWYERWNTQNSDGQELTTATLL    804
Mus_musculus                    S--NSSMVPSISNVYIPQCSASGQWRHVQCDGPHEQVFEWYERWKTQNGDGQELTPAALL    804
Cavia_porcelus                  S--NSSVPPTLSSIYIPQCSVSGQWQRVQCDGPPEQVFEWYDRWRAQNPEAKDLSPASLL    798
Canis_lupus_familiaris          S--DSSMLPTFSSSYIPQCSTAGQWRPVQCDGPPEQAFEWHERWEAQNNSGQELTPAELL    802
Panthera_leo                    S--DSSLLPTFASIYIPQCGTGGQWRPVQCDGPPEQAFEWYERWAAQNNGGQELTPAELL    803
Bos_taurus                      S--NPSTLPALSSIYIPQCSASGQWSPVQCDGPPEQAFEWYERWEAQNSAGQALTPAELL    803
Trichechus_manatus_latirostris  S--NSSLPPTLSSVYVPQCSTNGQWRRVQCDGPPEQAFEWYQRWVTQNKSGQELTPAELL    802
Columbia_livia                  S--DPGAQSQLSSVYVPQCDAAGGWRQVQCSGPPEQAFEWYGRWTAENNEGEALPVADLL    794
Taeniopygia_guttata             S--DPSAVPELSSLHLPQCDAGGGWRQVQCSGPPEQAFQWYERWIAENNQGKPLPVPELV    798
Larus_michahellis               S--GPSGMSQLSSVYLPQCDADGAWRQVQCGGPPEQAFEWYERWIAENNEGKTLPIADLL    799
Gallus_gallus                   S--DPSSLTQLSSIYLPQCDTDGAWRQVQCSGPPEQAFEWYERWITENNEGKTLPVANLL    791
Struthio_Camelus                S--DPSALSQLSSVYIPQCDADGAWRQVQCSGPPEQAFEWYQRWIEENNEGKIQPIANLL    773
Chelonia_mydas                  S--DPHALSQLSSVYIPQCSTNGNWRQVQCNGPPEQAFEWYQRWITQNNEGKTLPVVDLL    791
Alligator_mississippiensis      S--DTSALSQISRVYIPQCSTDGNWRPVQCSGPPEQAFEWYQRWISENNEGKTLPVTDLF    773
Crotalus_tigris                 L--NPTALTQLSNVYVPQCSLDGQWRPIQCNGPPEQAFEWYQGWNTQNNNNNVLAIPDLA    789
Python_bivittatus               L--SPTAVTQLSSIYVPQCSLDGQWRPIQCNGPPEQAFEWYQRWITQNSNGNALPFTDLA    765
Eublepharis_macularius          L--DPTPLRQLSSVYIPQCSRDGQWRRVQCDGPPEQAFEWYQRWITQNNNGTPLSFGDLM    782
Xenopus_tropicalis              A---EPELVQLSDVYVPQCAHDGKWKPVQCNGPAEQVFELYELWTKQNKNI---TLSETF    800
Aquarana_Catesbeiana            S--GLEELPELSKVYIPQCTLNGEWSTVQCSGPTEHAFEFYERWKTLNANN---SLLETL    792
Carassius_auratus               S--DTSSITQLSEVYIPRCASDGSWHQIQCDGPPEQATEFYREWVRIINSGQDLPVSEVL    778
Cyprinus_carpio                 S--DPSSVTQLSEVYIPRCAYDGSWHQIQCDGPPEQAIEFYREWVQIMNSGQDLPVSEVL    783
Danio_rerio                     S--DPPSVTQLSEIYIPRCSYDGSWHQIQCDGPPEQAIEFYREWVRIINLGQDLPASELL    783
Astyanax_mexicanus              A--SSSSVSQLSDVYVPRCASDGSWHQIQCDGPPEQAFDFYQEWIQINNDGKELPVSDLL    782
Clupea_harengus                 S--DPSSVSRLADVYVPRCTPDGRWHQVQCDGPPEQAFDFHQEWAQLNNAGKDLPVSELL    803
Oryzias_latipes                 ------------------------------------------------------------    642
Xiphophorus_maculatus           ------------------------------------------------------------    636
Stegastes_partitus              ------------------------------------------------------------    644
Cynoglossus_semilaevis          ------------------------------------------------------------    632
Lepisosteus_oculatus            S--SPTTAPQLSDVYIPQCNADGRWREVQCDGPPEQVFEFYQEWIKQNNAGEELSVLDVV    784
Lampetra_fluviatilis            PVLRGDAPSSPPPIYVPRCDTRGGWRPVQCDGAGQQIREFVDAWTKDERDASKATLSDLR    882
Lampetra_planeri                PVLRGDAPSSPPPIYVPRCDARGGWRPVQCDGAGQQIREFVDAWTKDEGDASKATLSDLR    820
Petromyzon_marinus              PVLRGDAPSSLPPLYVPRCDARGGWRPVQCDGAGQQIREFVDAWTKDEGNASKATLSDLR    835
```

```
Homo_sapiens                    VKIMSYREA------ASGNFSLFIQSLYEAGQQDVFPVLSQYPSLQDVPLAALEGKRP-Q    856
Pan_troglodytes                 VKIMSYREA------ASGNFSLFIQSLYEAGQQDVFPVLSQYPSLQDVPLAALEGKRP-Q    854
Gorilla_gorilla                 VKIMSYREA------ASGNFSLFIQSLYEAGQQDIFPVLSQYPSLQDVPLAALEGKRP-Q    856
Pongo_pygmaeus                  VKIMSYREA------ASGNFSFFIQSLYEAGQQGVFPVLSQYPSLQDVPLAALEGKRP-Q    856
Macaca_mulatta                  VKIMSYREA------ASGNFGLFIQSLYEAGQQGVFPVLSQYPSLQDVPLAALEGNRS-Q    856
Macaca_fascicularis             VKIMSYREA------ASGNFGLFIQSLYEAGQQGVFPVLSQYPSLQDVPLAALEGNRS-Q    856
Rattus_norvegicus               MKLMSYREV------ASTNFSLFLQSLYDAGQQSIFPVLAQYPSLQDVPQVVLEGATI-Q    857
Mus_musculus                    MKIVSYREV------ASRNFSLFLQSLYDAGQQRIFPALAQYPSLQDVPQVVLEGATT-P    857
Cavia_porcelus                  MTLISYKRE------ASGNFHLFVQTLYEAGQQGIFPVLAPFASLQEVPPAVLEGNLT-L    851
Canis_lupus_familiaris          MKIMSYREA------ASGSFRLFIQSLYAAGQQGIFPGLARYPSFEDVPLAVMEGNRT-Q    855
Panthera_leo                    MKIMSYKEA------ASRSFRLFIQSLYEAGQQGIFPGLARYPSFPDVPLAVMEGNLT-Q    856
Bos_taurus                      MKIMSYREA------ASRNFRLFIQNLYEAGQQGIFPGLARYSSFQDVPVSVLEGNQT-Q    856
Trichechus_manatus_latirostris  KRIMSYREV------ASKSFRLFIQSLYEAGQQGVFPGLARYSAFQDVPPVVLEGNLT-Q    855
Columbia_livia                  NIITGYKEA------SSKGFSAFIKALYEAGHQNVFPVFTTYSSFTDLPPEVLEGNVT-Y    847
Taeniopygia_guttata             NIIAGYKEA------SSGDFSAFVKALYEAGHQNVFPTFSMYSSFTDLPPQVLKGNLT-S    851
Larus_michahellis               SIITGYKEV------SSKGFSAFIKALFEAGHQNIFPAFTTYSSFTDLPPEVLEGNVT-Y    852
Gallus_gallus                   NIIIGYKNA------SSKGFSDFIKALYEVGHQNVFPAFTTYSSFADLPPEVLEGNVT-S    844
Struthio_Camelus                NVINGYKEA------SSKGFSAFIKALYEAGHQHIFPAFTKYSSFADLPPEVLEGNVT-N    826
Chelonia_mydas                  NILLEYKES------SSQGFAAFIKGLYVAGHQNIFPALAKYSSFAALPPGVLEGNVT-V    844
Alligator_mississippiensis      NILTEYKET------SSQRFAAFVKNLYEAGHQNIFPAFTKYPSFNALPVEVLDGSIT-T    826
Crotalus_tigris                 NILLDYKAR------SQQSFEDFIKILYTAGHQNVFPQFFKYSSFEAVPQDLLKGNT-LQ    842
Python_bivittatus               NILLDYKAR------SQKSFEDFVKILYEAGHQNIFPEFSKYSSFETVPQDLLEGNT-LQ    818
Eublepharis_macularius          AILLNYKER------SQPSFEAFIKALYEAGHQNVFPEFSRFPSFNDVPHDIREGNITVA    836
Xenopus_tropicalis              NIIQRYKRT------SAQSFPTFVKELYNNGHQKVFPIFSSYNTFNNVPAELLSGDLTS-    853
Aquarana_Catesbeiana            GLLLTYKES------TSQKFSNFLQKLYDQGHQNVFPMLSQYVSFSDLPEDVLSGNLTTV    846
Carassius_auratus               GILRAFARNTE----AMASFRVFVSELFKAGQHRVFPILARFGKFSDVPSDILDGNSKAI    834
Cyprinus_carpio                 GILRAFARNTE----AMASFRVFVSELFKGGQHRVFPVLARFGKFSDIPSDILDGNIKAI    839
Danio_rerio                     GILQTYARNTE----AMASFRAFVSELFKIGHHRVFPVLARFQKFSDIPSDLLDGNSEAV    839
Astyanax_mexicanus              GIPKDYAKNTE----AMASFRGFLSAVFQARHQKVFPALAKYEAFSELPSEILDGNTQAV    838
Clupea_harengus                 AILKNYGKNPA----AMSSFGGFLSALFEAGHQRVFPVLRRFATFTDLPQDVVQGQMEAV    859
Oryzias_latipes                 ------------------------------------------------------------    642
Xiphophorus_maculatus           ------------------------------------------------------------    636
Stegastes_partitus              ------------------------------------------------------------    644
Cynoglossus_semilaevis          ------------------------------------------------------------    632
Lepisosteus_oculatus            NIMKGYQKLPE----ALASFRGFVKELYSAGHQKVFPVLSRYASFSDVPEQILEGSRNSI    840
Lampetra_fluviatilis            ALLARARGAG------GEGLRSFLSFLYDSGRQDLFPELSLYPSLQGPLEVLNL----SG    932
Lampetra_planeri                ALLARARGAG------GDGLRSFLSFLYDSGRQDLFPELSLYPSLQGPIEVLNL----SG    870
Petromyzon_marinus              ELLARVRGAGGGGGGGKGLRSFLSFLYDSGRQDMFPELSLYPSLRGPPEVLNF----SG    891
```

```
Homo_sapiens                       PRENILLEPYLFWQILNGQLS-QYPGSYSDFSTPLAHFDLRNCWCVDEAGQELEGMRS-E    914
Pan_troglodytes                    PRENILLEPYLFWQILNGQLS-QYPGSYSDFSTPLAHFDLRNCWCVDEAGQELEGTRA-E    912
Gorilla_gorilla                    PRENILLEPYLFWQILNGQLS-QYPGSYSDFSTPLAHFDLRNCWCVDEAGQELEGTRA-E    914
Pongo_pygmaeus                     PRENVLLEPYLFWQILNGQLS-QYPGSYSDFSTLLAHFDLRNCWCVDEAGQELEGTRA-E    914
Macaca_mulatta                     SRENVLLDPYLFWQILNGQLS-RYPGPYSDFSTPLAHFDLRNCWCVDEAGQELEGTRA-E    914
Macaca_fascicularis                SRENVLLDPYLFWQILNGQLS-RYPGPYSDFSTPLAHFDLRNCWCVDEAGRELEGMRA-E    914
Rattus_norvegicus                  PGENIFLDPYIFWQILNGQLS-QYPGPYSDFSMPLEHFNLRSCWCVDEAGQELDGTRT-R    915
Mus_musculus                       PGENIFLDPYIFWQILNGQLS-QYPGPYSDFNMPLEHFNLRSCWCVDEAGQKLDGTQT-K    915
Cavia_porcelus                     STENLFLEPYFFWQMLNGQLS-RYPGPYSDFSKPLAHTDLRSCWCVDAAGQELEGTRA-R    909
Canis_lupus_familiaris             PGGNILLEPYLFWQILNGQLS-RYPGAYSDFSAPLAHFDLRSCWCVSEAGRELEGTRT-E    913
Panthera_leo                       PGGNILLDPYVFWQILNGQLS-RYPGPYSDFSTPQAHLDLRSCWCVNEAGQELESTRT-E    914
Bos_taurus                         PGGNVFLEPYLFWQILNGQLD-RYPGPYSDFSAPLAHFDLRSCWCVDEAGQKLEGTRN-E    914
Trichechus_manatus_latirostris     PAGNILLEPYLFWQILNGQLS-RYPGPYSDFSTPLAHFDLRSCWCVDRAGQELEGTWA-E    913
Columbia_livia                     ASENILLDPYTFWQLLNEQLT-YYPGPYTDFSAPLNHFELRDCWCVDNKGEELQGTKT-E    905
Taeniopygia_guttata                ASENILLDPYTFWQLLTDQLS-YYPGPYTDFSAPLGHFELRDCWCVDSKGGELEGTKA-G    909
Larus_michahellis                  ASENILLDPYTFWQLLNEQLT-YYPGPYTDFSAPLSHFELRNCWCVDSKGEELQGTKA-E    910
Gallus_gallus                      ASENILLDPYTFWQLLNEQFT-YYPGAYTDFSAPLSHFELRDCWCVNSNGEELQGTRA-E    902
Struthio_Camelus                   ASENILLDPYTFWQLLNGQLT-YYPGAYTDFSAPLSHFELRDCWCVDSKGKELQGTKA-K    884
Chelonia_mydas                     ASENILLDPYTFWQLLHGQVT-HYPGSYTDFNALLGHFELRNCWCVDSKGEELQGTKA-E    902
Alligator_mississippiensis         ESENILLDPYTFWQLLQGQLN-YYPGSYADFSALLGHFELRNCWCVDDKGEELQGTKT-E    884
Crotalus_tigris                    ISENALLDPFIFWQLLQGRLN-HYPGSYSDFSIPLGHFDLRNCWCVDDKG-RLQGRQA-S    899
Python_bivittatus                  SSENVLLDPFTFWQLLQGHFN-QYPGSYSDFSIPLGHFDLRNCWCVDKKG-RLQGSQA-N    875
Eublepharis_macularius             SSENVLLEPFTFWQLLQGQLI-HYPGSYSDFSHPLGHFDLRRCWCVDEQG-QMQGSKA-E    893
Xenopus_tropicalis                 PSDNILLNPFVFWRLLNGSLT-FYPGPYTAFSQPLSHFELRNCWCVDLEGQKLEQKEV-S    911
Aquarana_Catesbeiana               SSDNLLFNPYIFWRLLTGSLS-DYPGSYSDFSSPLGHIEQRSCWCVDPDGQKIPGMET-V    904
Carassius_auratus                  YGPSVFLNPLSLWRLIRGEDA-GYPGPLSDFSLPLGSFHLRQCWCVNPQGDMLADSKA-P    892
Cyprinus_carpio                    YGPSVFLNPLSLWRLIRGENS-SYPGPLSDFSLPLGSFHLRQCWCVNPQGDMVTDSKA-P    897
Danio_rerio                        YGPSVYLNPLSLWRLIRLDDS-GYPGLLSDFSVPLGSFDLRQCWCVDLEGDMLAGSKA-P    897
Astyanax_mexicanus                 SGPSVFLNPLYLWKLLRGDSS-QYPGPLADFSAPLNHFDLRQCWCVDQTGGMIAGSKA-P    896
Clupea_harengus                    FGPSVFLNPLSMWTLLKGGAS-RYPGPLSDFSLPLGHLHLRQCWCVSPAGDTVPDTKA-P    917
Oryzias_latipes                    ------------------------------------------------------------    642
Xiphophorus_maculatus              ------------------------------------------------------------    636
Stegastes_partitus                 ------------------------------------------------------------    644
Cynoglossus_semilaevis             ------------------------------------------------------------    632
Lepisosteus_oculatus               SGSSVLLNPLTLWQLLHGNST-QYPGQYSDFSSPLGHFELRRCWCVNKNGEMIMDTKV-G    898
Lampetra_fluviatilis               SSGRFLRNPEVVWKILTENASFVYTGDYAEFSGGYGDFESRLCWCVDAAGHEIDGTRTAS    992
Lampetra_planeri                   SSSRFLRNPEVVWKILTENASFVYTGDYAEFSGGYGDFESRLCWCVDAAGHEIDGTRTAS    930
Petromyzon_marinus                 SSGRFLRNPEVVWKILTENASFVYTGDYAEFSGRYADFESRLCWCVDAAGHEIDGTRTVS    951
```

TG type 1-8

```
Homo_sapiens                       PSKLPTCPGSCEEAKLRVLQFIRETEEIVSASNSSRFPLGESFLVAKGIRLRNEDL-GLP    973
Pan_troglodytes                    PSKLPTCPGSCEEAKLRVLQFIRETEEIVSASNSSRFPLGESFLVAKGIRLRNEDL-GLP    971
Gorilla_gorilla                    PSKLPTCPGSCEEAKLRVLQFIRETEEIVSASNSSRFPLGESFLVAKGIRLRNEDL-GLP    973
Pongo_pygmaeus                     PSKLPTCPGSCEEAKLRVLQFIRETEDIVSASNSSRFPPGESFLVAKGIRLRNEDL-GLP    973
Macaca_mulatta                     PSKLPTCPGSCEEAKLLVLQFIRETEEIVSASNSSRFPLGESFLVAKGIRLRNEDL-GLP    973
Macaca_fascicularis                PSKLPTCPGSCEEAKLLVLQFIRETEEIVSASNSSRFPLGESFLVAKGIRLRNEDL-GLP    973
Rattus_norvegicus                  AGEIPACPGPCEEVKFRVLKFIKETEEIVSASNASSFPLGESFLVAKGIQLTSEEL-GLP    974
Mus_musculus                       PGEIPACPGPCEEVKLRVLKFIKETEEIVSASNASSFPLGESFLVAKGIQLTSEEL-DLP    974
Cavia_porcelus                     PNQVPACPGSCEEVKLDVLQFIKETEEIVSAFNTSRFLVGESFLIAKGIRLTSEEL-RLP    968
Canis_lupus_familiaris             PSKVPACPGSCEEVKLRVLQFLKETEEIVLASNSSWFPLGESFLAAKGIWLTDEEL-SLP    972
Panthera_leo                       PSKVPACPGSCEEVKSRVLQFIKETEDIVLASNSSWFPLGESFLAAKGILLTDEEL-SLP    973
Bos_taurus                         PNKVPACPGSCEEVKLRVLQFIREAEEIVTYSNSSRFPLGESFLAAKGIRLTDEEL-AFP    973
Trichechus_manatus_latirostris     PSKVPACPGSCEEAKLRVLQFIKEAEEIVLVSNSSQFPLGESFLVAKGIRLSNKEL-SLP    972
Columbia_livia                     VNQVPACPGICEGVKQKAMKFMEEAEQLILASNSSHFPFGESYLMAKGIQLTDNDLLRSA    965
Taeniopygia_guttata                VNQVPACPGICEGVKQEAMKFMEEAEQLILASNESQFAFGESFLLAKGIELRDRDLLRSA    969
Larus_michahellis                  VNQVPACPGMCEGIKQEAMKFMEEAEQLILASNSSHFPFGESFLMAKGIQLTDNDLLRSA    970
Gallus_gallus                      VSQVPTCPGMCQSVKQEAMKFIKETEQLIRDSNTSHFPFGESFLMAKGIQLMDSDLLRST    962
Struthio_Camelus                   VNQVPACPGICENVKQEALKFVEEAEQLILTSNSSHFPFGESFVMAKGIQLSNSDLLRSA    944
Chelonia_mydas                     ANKIPACPGACENVKKEAMQFIDEAEQLILASNGSHIPLGQSFLMAKGLRLTDNDLLHFA    962
Alligator_mississippiensis         VNKVPACPGACESMKKEAMQLIDEAEQLIVASNGSHFPFGESFLMAKGIQLMDHDLLHFT    944
Crotalus_tigris                    VNQVPTCLGTCEAAKREAVGFVNKVEQLVQESNSSHFPFGQSFLMSQGSVLRGRELWSN-    958
Python_bivittatus                  VNQVPICLGTCEAAKQEAVQFVNNVEQLIQESNSSHFPFGQSFLMTQVSMLTGEEPWSN-    934
Eublepharis_macularius             VNEFPGCPRACAAVQQEAMRFSEKVEQLIRESSSSHFPFGQSFLMAEGLTLTEDEL-LD-    951
Xenopus_tropicalis                 KNEVPQCPTSCELAKLRAMKFIKEAEDLSAISNISHFPWGLSFLIANGIELTERELLHPE    971
Aquarana_Catesbeiana               SKKVPKCPGMCELARMKSQRFIEEAQKIIGASNVTHFPLGLSFLLANGIQLSGRDLMYTE    964
Carassius_auratus                  VGQIPKCPGPCSMVQNQVSEFLKQAEELISASNDSHVPVGYGFLLAESVYLSPEELEQ--    950
Cyprinus_carpio                    VGQIPKCPGPCSMVQNQVSEFLRQAEELISASNDSHVPVGYGFLLAESVYLSPEELEQ--    955
Danio_rerio                        VGQIPKCPGPCSVVQSQVSEFLKQAEELISASNSTHVPVGYGFLLAESVYLSPEELEQ--    955
Astyanax_mexicanus                 VKQIPKCPGPCSLIKGQVDEFLAKAERQISLSNSSYIPVGYSFLLAESVNLISEAMQQ--    954
Clupea_harengus                    PNQVPKCPGTCALAEQEVTRFLATAEEFISVSNSSHVPVGYSFLLAEAVRLSPQELLLR-    976
Oryzias_latipes                    T--PKSSEGRCSDALAEVAAFRQEVNRIISLSNSSHIPLGYGFLLAEGLRLTPEELQV--    698
Xiphophorus_maculatus              TLELHSSAGRCSKALAEVTAFRQEVDSLVFLSNSTHFPVGYGFLLAEGLSLSQSEEELR-    695
Stegastes_partitus                 AQKLRSSAGQCSQALAEVTAFRQEVKSIIAVSNSSHLPLGYGFLLAEGLRLTPEELQIS-    703
Cynoglossus_semilaevis             ---PQRVSGACSGLLSKVTTFREEVSSLVALSRSSHLALGLGVLLAEGLSLTPEELQLS-    688
Lepisosteus_oculatus               INEIPTCPGPCSVVNQQVARFLQEAEDIITDSNNSRIPFGFSFLQAKGLQLTERELLINP    958
Lampetra_fluviatilis               PGQLPKCPGACHLAAADVSRYLQQADLLIGSAGASS--AGEGVAFGRGLAFTEDELLGSP    1050
Lampetra_planeri                   PGQLPKCPGACHLAAADVSRYLQQADLLIGSAGASS--AGEGVAFGRGLAFTEDELLGSP    988
Petromyzon_marinus                 PGQLPKCPGACHLAAADVNRYLQQADLLIGSAGASS--AGEGVAFGRGLAFTEDELLGSP    1009
```

| | | |
|---|---|---|
| *Homo_sapiens* | PLFP-PREAFAEQFLRGSDYAIRLAAQSTLSFYQRRRFSPDD-SAGASALLRSGPYMPQC | 1031 |
| *Pan_troglodytes* | PLFP-PREAFAEQFLRGSDYAIRLAAQSTLSFYQRRRFSPDD-SAGASALLRSGPYVPQC | 1029 |
| *Gorilla_gorilla* | PLFP-PREAFAEQFLRGSDYAIRLAAQSTLSFYQRRRFPPDD-SAGASALLRLGPYVPQC | 1031 |
| *Pongo_pygmaeus* | PLFP-PREAFAEQFLRGSDYAIRLAAQSTLSFYQRRRFSLDD-SAGASALLRSGPYMPQC | 1031 |
| *Macaca_mulatta* | PLFP-PREALAEQFLRGSDYAIRLAAQSTLSFYQRRRFSLDD-SAGASALLQLGPYVPQC | 1031 |
| *Macaca_fascicularis* | PLFP-PREALAEQFLRGSDYAIRLAAQSTLSFYQRRRFSLDD-SAGASALLQLGPYVPQC | 1031 |
| *Rattus_norvegicus* | PLYP-SREAFSEKFLRGSEYAIRLAAQSTLTFYQKLRASLGE-SNGTASLLWSGPYMPQC | 1032 |
| *Mus_musculus* | PQFP-SRDAFSEKFLRGGEYAIRLAAQSTLTFYQSLRASLGK-SDGAASLLWSGPYMPQC | 1032 |
| *Cavia_porcelus* | PHFP-SQEAFFEKFLRGRDYAIRLAAQSTSNLYQRLRTSQDD-WSETAVPLRSGPYVPQC | 1026 |
| *Canis_lupus_familiaris* | RLSP-SRETFSEKFLSGGDYALRLAAQSTLDFYQRRGFLLGD-STRTSALLRPVPYVPQC | 1030 |
| *Panthera_leo* | QLSP-SRATFSEKFLSGGDYAIRLAAQSTLDFYQRRAFLRGDDSTRASALLRPGPYVPQC | 1032 |
| *Bos_taurus* | PLSP-SRETFLEKFLSGSDYAIRLAAQSTFDFYQRRLVTLAE-SPRAPSPVWSSAYLPQC | 1031 |
| *Trichechus_manatus_latirostris* | WPFS-PWETFSEKFLMGSDYALRLAAQSTFTFYQRRGSSLGD-SAGAAALLSNGPYVPQC | 1030 |
| *Columbia_livia* | RPYE-SELVVSEELLMGSDYALQLAAWSVLHFYWRSHFTSKR-SAGEATQLGFLPYIPQC | 1023 |
| *Taeniopygia_guttata* | RPDG-LQAAIPQELLSGRDSALQLAAWSVLRFYWQSYFTSKS-SAGEATQLGFFPYIPQC | 1027 |
| *Larus_michahellis* | RPYE-SEVVVSEKLLSGSDYALQLAAQSVLHFYWRSRFTSKR-SAGETTRLGFLPYIPQC | 1028 |
| *Gallus_gallus* | RPYE-SEMMVSEKLLSGSDYALQLAALSVLNFYWQNHFTLKG-FAGEATQLGFHPYIPQC | 1020 |
| *Struthio_Camelus* | WPYE-SVILVSERMLSGSDYALQLAAQSVLHFYWRSHFTLKG-SAGEATQLGFHPYIPQC | 1002 |
| *Chelonia_mydas* | ETFQ-SGIAFSEKLLAGSDYAVRLAAQSTLHFYWRSHFTSKG-SAGQATLLGFHPYIPQC | 1020 |
| *Alligator_mississippiensis* | QSFQ-SGTMFSEKILTGSDYAIRLAAQSTLHFYWRNRFISRS-SAGEAMLLGFHPYIPQC | 1002 |
| *Crotalus_tigris* | -FSQ-LESTFAEILLAGSDYAIQLAAQSTLQFYRKRLFASLD-SAGEALRLAFLPYIPQC | 1015 |
| *Python_bivittatus* | -FSQ-LEVTFSETLLAGSDYAIRLAAQSTLRFYRKRFLASPD-SAREAVRLAFQPYIPQC | 991 |
| *Eublepharis_macularius* | -YSQ-PGITFSEGLLTGGEYAIRLAAQSTLHFYWRNLFASRG-SAGEATHLGLQPYVPQC | 1008 |
| *Xenopus_tropicalis* | GFFR-SGEPFFERFRDRGDYAVHLAAQSTLRFHWQQRSSLER-SSGEVSRVAYRPYVPQC | 1029 |
| *Aquarana_Catesbeiana* | E-YK-SGIALSESFLKKDMYAIQLAAYSTLKFFQESR----L-GSGETLQLSYAPYIPQC | 1017 |
| *Carassius_auratus* | -MRT-SKIPVTQTLLSNTNSALRLAAHSTLHFYWQSRLMADD-KDRQSLILGYQPYIPQC | 1007 |
| *Cyprinus_carpio* | -MRA-SKIPITQTLLSNTNSALRLAAHSTLHFYWQSRLMAND-KDRQSLMLGYQPYMPQC | 1012 |
| *Danio_rerio* | -TRS-SMIPVTQTLLSNTDLALRLAAHSTLHFYWQSRLMASD-RDRQSLMLGYQPYIPQC | 1012 |
| *Astyanax_mexicanus* | -TFS-TGFQVSEDFLSNTDSALRLAAHSTLHFYWQSRLLASE-TDRESLRLGYQPYSPQC | 1011 |
| *Clupea_harengus* | -AVP-AGAQLSDALLSHSSSSLRLAAHSTLHFYWQSRLMSSG-VDKESLQLGYQPYQPHC | 1033 |
| *Oryzias_latipes* | -IQS-EELQLSEELLSGSRAALPLAAFSTLQMLLPAG------------RRSYEPFLPQC | 744 |
| *Xiphophorus_maculatus* | LSQS-EELKVSERFLSRSRSALRLAASSTLLMLLPPH------------RRSYQLFTPQC | 742 |
| *Stegastes_partitus* | -QSE-EELRVSDRLLSRSKAALRLAAFSTVQMLLPPR------------RRSYQPFTPQC | 749 |
| *Cynoglossus_semilaevis* | -QPE-EALHIGEELLSGTRAALRLAAFSTVQMYLHPL------------GRSYQPYTPQC | 734 |
| *Lepisosteus_oculatus* | DSFE-SSLNFNEKLLSNSDSALRLAVHTTLQFYWRTHFSSAL-NQRDGFLLGYQPYRPQC | 1016 |
| *Lampetra_fluviatilis* | LGLGDVKGEVAAILAGGTGYAVRLAAQAAMHFHWRRRFFGLG-TIGEGVFNGFDPYMPQC | 1109 |
| *Lampetra_planeri* | LGLGDVKGEVAAILAGGTGYAVRLAAQAAMHFHWRRRFFGLG-TIGEGVFNGFDPYMPQC | 1047 |
| *Petromyzon_marinus* | LGLGDVKGEVAAVLAGGTGYAVRLAAQAAMHFHWRRRFFGPG-TIGEGVFNGFDPYMPQC | 1068 |

TG type 1-9

| | | |
|---|---|---|
| *Homo_sapiens* | DAFGSWEPVQCHA--GTGHCWCVDEKGGFIPGSLTARSLQIPQCPTTCEKSRTSGLLSSW | 1089 |
| *Pan_troglodytes* | DAFGSWEPVQCHA--GTGHCWCVDEKGGFIPASLTARSLQIPQCPTTCEKSRTSGLLSSW | 1087 |
| *Gorilla_gorilla* | DVFGSWEPVQCHA--GTGHCWCVDEKGGLIPASLTARSLQIPQCPTTCEKSRTSGLLSSW | 1089 |
| *Pongo_pygmaeus* | DAFGSWEPVQCHT--GTGHCWCVDEKGGFIPASLTARSLQIPQCQTTCEKSRTSGLLSSW | 1089 |
| *Macaca_mulatta* | DAFGSWEPVQCHT--GTGHCWCVDEKGGFIPASLTARSLQIPQCPTTCEKSRTSGLLSSW | 1089 |
| *Macaca_fascicularis* | DAFGSWEPVQCHT--GTGHCWCVDEKGGFIPASLTARSLQIPQCPTTCEKSRTSGLLSSW | 1089 |
| *Rattus_norvegicus* | NTIGGWEPVQCHP--GTGQCWCVDGWGELIPGSLMARSSQMPQCPTSCELSRANGLISAW | 1090 |
| *Mus_musculus* | NMIGGWEPVQCHA--GTGQCWCVDGRGEFIPGSLMSRSSQMPQCPTNCELSRASGLISAW | 1090 |
| *Cavia_porcelus* | DASGNWEPRQCHM--GTGYCWCMDAKGEFIPNSLTARSPQVLQCPSPCEQSRANGLLSGW | 1084 |
| *Canis_lupus_familiaris* | DVWGGWEPVQCHA--RTGYCWCVDGKGEYVPASLTARSPRILRCPTACEKSRASGLLSSW | 1088 |
| *Panthera_leo* | DAWGRWEPVQCHA--RTGYCWCVDRRGEYVPASLTARSSQIPRCPTACEASRTSGLLSSW | 1090 |
| *Bos_taurus* | DAFGGWEPVQCHA--ATGHCWCVDGKGEYVPTSLTARSRQIPQCPTSCERLRASGLLSSW | 1089 |
| *Trichechus_manatus_latirostris* | DAFGSWAPVQCYN--RTGHCWCVDGKGEYIPASLAARSPQTPQCPTTCEESRASGLISSW | 1088 |
| *Columbia_livia* | DGLGNWEPAQCYE--STGHCWCVDERGRYVMDSLVSRSAELPKCQTSCQRSRTNALISSW | 1081 |
| *Taeniopygia_guttata* | DGLGNWEPVQCYE--STGHCWCVDERGRYIMDSLVSRSSELPKCRTSCQRSRANALISSW | 1085 |
| *Larus_michahellis* | DGLGNWEPTQCYE--STGHCWCVDERGRYVMDSLVSRSAELPKCQTSCQRSRTNALISSW | 1086 |
| *Gallus_gallus* | DGLGNWEPTQCYE--STGHCWCVDERGRYVTDSLVSRSAESPKCQTSCQRSRANALISSW | 1078 |
| *Struthio_Camelus* | DGLGNWEPTQCYE--STGHCWCVDERGRYVTDSLISRSVELPNCQTSCQRSRANALISSW | 1060 |
| *Chelonia_mydas* | DGLGNWEPVQCYK--STGHCWCVDERGRYITNSLIIRSAHFPKCQTSCQRSRANALISSW | 1078 |
| *Alligator_mississippiensis* | DGLGNWEPVQCYE--STGHCWCVDKRGRYVTGSLRARSAQLPTCQTSCQRSRANALISSW | 1060 |
| *Crotalus_tigris* | DGQGNWEPIQCYE--STGHCWCVDETGRYVSDSLMTRSTQLPQCQTSCQRSQVNAEIASW | 1073 |
| *Python_bivittatus* | DGEGNWEPIQCYE--SSGHCWCVDEMGRYVSDSLVTRSTQLPQWV--------------- | 1034 |
| *Eublepharis_macularius* | DGLGNWEPVQCYD--SSGHCWCVDERGRYVLDSLVTRSAQLPQCRTPCQRSQTNALISSW | 1066 |
| *Xenopus_tropicalis* | DGLGNWEPVQYYG--STGHNWCVDADGNYIAGSLEGRTSRPRQCQTRCQQDQTNMVVSSW | 1087 |
| *Aquarana_Catesbeiana* | DGLGNWNPMQFYQ--GTGHYWCVDNEGSYLEGSLVSRTTSSPPKCQTQCRRAETKALISNW | 1075 |
| *Carassius_auratus* | DAYGQWLPNQCYQ--STGYCWCVDEEGRYITDSLMFRSAPSRQCQTMCQRIQSNFLLSDW | 1065 |
| *Cyprinus_carpio* | DAYGQWLPNQCYQ--STGHCWCVDEEGRYITGSLTSRSAPSRQCQTTCQRLQSNFLLSDW | 1070 |
| *Danio_rerio* | DAYGQWLPNQCYQ--STGLCWCVDEEGQYIADSLTSRSSLPQMCQTLCQRLQSHTLLSDW | 1070 |
| *Astyanax_mexicanus* | DAYGQWLPTQCYP--STGRCWCVDEEGGYITGSLTDRTVYLPQCQTPCQRSQAQYSMSVW | 1069 |
| *Clupea_harengus* | DSQGQWLPQQCHP--STGQCWCVDEEGRYITGSLTRHSTQPPQCPSRCQRAQSLSDLSDW | 1091 |
| *Oryzias_latipes* | DADGGWLHTQCSH--STGQCWCVDENGEYVSNSLKSRSLGLPKCPSRCQRAEAHFLLSDW | 802 |
| *Xiphophorus_maculatus* | DAAGNWRLTQCYH--STGQCWCVDEEGEFIPGSLR-RSLRLPRCRSRCQRAEAHSLLSDW | 799 |
| *Stegastes_partitus* | DADGRWMVTQCYH--STGQCWCVDEDGEYIPDSLTSRSLRLPRCLTRCQRAEAHSLLSGW | 807 |
| *Cynoglossus_semilaevis* | DADGNWLHTQCHH--STGQCWCVDDDGDYMANSLTSRSVKLPKCPTNCQRAQAHTLLSGW | 792 |
| *Lepisosteus_oculatus* | DAHGQWQPSQCYF--STGQCWCVDEEGSYIPGSLTSRSIQLPQCGTLCQRSHTRAVVSNW | 1074 |
| *Lampetra_fluviatilis* | TEPGAWEPAQCDK--SSGFCWCVDAAGEFVAGSLVARPRRRPQCATPCQRARAEALLTGW | 1167 |
| *Lampetra_planeri* | TEPGAWEPAQCDK--SSGFCWCVDAAGEFVAGSLVARPRRRPQCATPCQRARAEALLTGW | 1105 |
| *Petromyzon_marinus* | TEPGGWEPAQCVVRAGAGFCWCVDAAGEFVAGSLVARPRRRPQCATPCQRARAEALLTGW | 1128 |

```
Homo_sapiens                     KQA-RSQENPSP---KDLFVPACLETGEYARLQASGA------GTWCVDPASGEELR-PG   1138
Pan_troglodytes                  KQA-RSQENPSP---RDLFVPACLETGEYARLQASGA------GTWCVDPASGEELR-PG   1136
Gorilla_gorilla                  KQA-RSQENPSP---KDLFVPACLETGEYARLQASGA------GTWCVDPASGEELR-PG   1138
Pongo_pygmaeus                   KQA-RSQENPSP---KDLFVPACLETGEYARLQASDA------GTWCVDPASGEELL-PG   1138
Macaca_mulatta                   KQA-RSQGNSSP---KDLFIPACLETGEYARLQASEA------GTWCVDPASGEELL-PG   1138
Macaca_fascicularis              KQA-RSQGNPSP---KDLFIPACLETGEYARLQASEA------GTWCVDPASGEELL-PG   1138
Rattus_norvegicus                KQA-GHQRNPGP---GDLFTPVCLQTGEYVRQQTSGT------GAWCVDPSSGEGVP-TN   1139
Mus_musculus                     KQA-GPQRNPGP---GDLFIPVCLQTGEYVRKQTSGT------GTWCVDPASGEGMP-VN   1139
Cavia_porcelus                   KQA-GSQGDPSP---EGLFIPTCLETGEFARMQVSGA------GAWCVDPVSGEGMQ-PS   1133
Canis_lupus_familiaris           KQA-GSQGNPSP---KDLFIPTCLETGEFARRQESEG------GTWCVDPGSGAGRP-PG   1137
Panthera_leo                     KQA-GSQGNPSP---KDLFIPTCLETGEFARLQESEA------GAWCVDPASGAGLP-LA   1139
Bos_taurus                       KQA-GVQAEPSP---KDLFIPTCLETGEFARLQASEA------GTWCVDPASGEGVP-PG   1138
Trichechus_manatus_latirostris   KQA-VSQGNPSP---KDLFIPTCLQTGEYARLQVSKA------GTWCVDPASGDREL-RG   1137
Columbia_livia                   RQS-GSKLDAST---TDLFIPYCLETGEYAVLQRSNT------DTWCINPVSGEVFQ-RG   1130
Taeniopygia_guttata              RQS-SAKLDTSA---ADLFIPNCLETGEYAVLQRSDT------DIWCVDPVSGEIFQ-RG   1134
Larus_michahellis                RQS-GSKLGAST---ADLFVPYCLETGEYAVLQRSDA------DIWCVDPVSGEVLQ-RG   1135
Gallus_gallus                    RQS-GSKLEAAA---TDLFVPYCLETGEYAMLQRSYT------NTWCVDPESGEVFQ-RG   1127
Struthio_Camelus                 RQS-GSKLNATT---MDLFIPYCLETGEYAVLQRSYA------DVWCVDPASGEVFQ-RA   1109
Chelonia_mydas                   KQS-GSKLSATA---ADLFIPSCLETGEYTMLQKSDT------DTWCVDPTSGAIIQ-HS   1127
Alligator_mississippiensis       KQS-GAPCNTAS---ADLFIPSCLETGEYTLLQKLDS------DAQCVDPMSGDVLQ-RS   1109
Crotalus_tigris                  RMN-GQAQNAIS---AALFTPSCLENGDYANVQKLGS------RGWCVNPVSGEVTQ-ES   1122
Python_bivittatus                ----------------------DKNGDYANIQKLDS------QAWCLNPVSGEVMK-ES   1064
Eublepharis_macularius           RQN-ASKRDMTS---ADLFTPSCLETGAYAMLQKSET------GAWCVDPASGEVLQ-GS   1115
Xenopus_tropicalis               LPQTWSPQS-GP---VDMVVPGCMENGQFSALQTSES------QFWCVAPSSGQVIQ-HG   1136
Aquarana_Catesbeiana             LPRKSSTVSTET---YEGFKPNCTEAGQYVPLQRSDT------DSWCVSPVSGEAIP-KD   1125
Carassius_auratus                TQT-SSN----I---TYTYSPSCEEDGEFSVLQKPGTV---RLQGLCVSPITGQVIQ-PA   1113
Cyprinus_carpio                  TQT-SSN----I---TYTYSPSCEEDGEFSVLQKPSSV---HSQGLCVSPITGQVIQ-PA   1118
Danio_rerio                      RQT-SSN----I---TYTYRPSCEEDGEFSVLQKATSG---HMQGFCVSPITGRVIQ-PA   1118
Astyanax_mexicanus               TKS-TPD----I---TFSYSPSCEADGEFSVLQRQDSS-----VASCVSPVTGKMIQ-PA   1115
Clupea_harengus                  VKP-TSD----I---TSYYSPSCDEKGQFSVLQTPGTQ---GSSAWCVDPATGQTIQ-PA   1139
Oryzias_latipes                  MKG----SDVS----SSGYRPQCEQDGRFSVLQTGGA------AGWCVHPQTGEPIQ-AA   847
Xiphophorus_maculatus            MKA----SDITT---ASPYHPQCEQDGRFSVLQTGGA------AGWCVNPLTGETLQ-SA   845
Stegastes_partitus               MKG----SDITT---MSAYLPQCEQDGQFSVLQKGGA------AGWCVNPQTGETIQ-AA   853
Cynoglossus_semilaevis           RKS----SDVST---T--VPPQCDENGRYSLLQTGGA------GGSCVNPLTGEEMQ-TA   836
Lepisosteus_oculatus             KLS-ATDPSAMA---TTVINPSCQKNGEFTILQNGDRE---NGSVWCVNPLTGKPIQ-PA   1126
Lampetra_fluviatilis             KSL-GSVENGSSIEVLTKHTPACTPAGGQFEARQRDSESGLEQRSWCVDPASGQQAEPLG   1226
Lampetra_planeri                 KSL-GSVENGSSIEVLTKHTPACTPAGGQFEARQRDSESGLEQRSWCVDPASGQQAEPLG   1164
Petromyzon_marinus               KSL-GSVENGSSIEVLTKHTPACTPSRKYKKC------HYLLRPNWWPSPSYSYT---PT   1178
```

TG type 1-10

```
Homo_sapiens                     ---SSSSAQCPSLCNVLKSG-VLSRRVSPGYVPACRAEDGGFSPVQCDQAQGSCWCVMD-   1193
Pan_troglodytes                  ---SNSSAQCPSLCNVLKSG-VLSRRVSPGYVPACRAEDGGFSPVQCDQAQGSCWCVMD-   1191
Gorilla_gorilla                  ---LNSSAQCPSLCNVLKSG-VLSRRVSPGYVPACRAEDGGFSPVQCDQAQGSCWCVMD-   1193
Pongo_pygmaeus                   ---SNSSAQCPSLCNVLKSG-VLSRRVGPGYVPACRAEDGGFSLVQCDQAQGSCWCVMD-   1193
Macaca_mulatta                   ---SNSSAQCPSLCNVLKSG-VLSRRVSSGYVPACREEDGGFSPVQCDQAQGSCWCVTD-   1193
Macaca_fascicularis              ---SNSSAQCPSLCNVLKSG-VLSRRVGSGYVPACREEDGGFSPVQCDQAQGSCWCVTD-   1193
Rattus_norvegicus                ---TNSSAQCPGLCDALKSR-VLSRKVGLGYTPVCEALDGGFSPVQCDLAQGSCWCVLA-   1194
Mus_musculus                     ---TNGSAQCPGLCDVLKSR-ALSRKVGLGYSPVCEALDGAFSPVQCDLAQGSCWCVLG-   1194
Cavia_porcelus                   ---TNGSTQCPGLCSVLRNR-VLSRTVSPGYTPTCGTEDGAFSLVQCDPAQGSCWCVLD-   1188
Canis_lupus_familiaris           ---TDSSAPCPSLCEELSSG-TLSRRASSGYSPACRAEDGGFSPVQCDLAQGSCWCVLD-   1192
Panthera_leo                     ---TNSSAPCPRLCDVLPSG-VLPRRASSGSTPACSVEDGRSSPVRCDPAQGSCWCVLA-   1194
Bos_taurus                       ---TNSSAQCPSLCEVLQSG-VPSRRTSPGYSPACRAEDGGFSPVQCDPAQGSCWCVLG-   1193
Trichechus_manatus_latirostris   ---TDRSAQCPSLCEALRSG-VLERRAGLGYIPACGAGDRRFSPVQCDPAGDSCWCVLG-   1192
Columbia_livia                   SKDSDGNPECPSFCNMLKSK-TSLREAGKGYIPQCEGDDGSFSPVQCSQDQESCWCVFD-   1188
Taeniopygia_guttata              SKGLDGNPECPSLCNMLKSK-AVSREAARGDIPQCEGDRGSFSPVQCSQEQESCWCVFG-   1192
Larus_michahellis                SKDLDGNPECPSFCNMLKSK-TSLREAGKGYIPQCEGDNGTFSPMQCSQDEGSCWCVFD-   1193
Gallus_gallus                    NKDSDGNPECPTFCNMLKLK-VGLREVGKGYVPKCEEANGIFSPVQCTQDEETCWCVFD-   1185
Struthio_Camelus                 SEDSDGSPECPSFCNMLKSR-TSLREAGKGYIPQCEGEDGIFSPMQCSQDQESCWCVFD-   1167
Chelonia_mydas                   SKDSDGSPQCPGFCSMLKSK-VRSREIGKGYIPQCEGDNGNFSPVQCDQDQESCWCVFE-   1185
Alligator_mississippiensis       SQDSDGNPQCPALCSMLKPK-MSSREIGKGYIPQCEGNNGSFSPVQCSQDEKSCWCVFE-   1167
Crotalus_tigris                  KADSNGEVPCPSYCNLLKSK-ALVREIGRGYIPECESDPQGFSPVQCSEDQASCWCVFD-   1180
Python_bivittatus                EIDSNGEVRCPSYCSLLKSK-ALVREMGKGYIPRCDSDREGFSSTQCSEDQESCWCVFD-   1122
Eublepharis_macularius           SLDSKGRVHCPGYCSMLKSK-ALLWELGRGYIPQCEGSSGIFSPMQCDDNKESCWCVFE-   1173
Xenopus_tropicalis               KTDIPSNNECPGYCSVLKSE-VQGRNVGFPFVPECG-ENGDFSSTQCDQDGTMCFCVFP-   1193
Aquarana_Catesbeiana             VNAL-KDINCPRPCAILTEDNSQGEGIGVA----C---SGDGTREQCDPDSNVCYCLFP-   1176
Carassius_auratus                ITSPSGELQCPGWCSLQKTL-ALHREIGVGYEPQCVQDGQRFSPLQCDL--SYCWCVSQ-   1169
Cyprinus_carpio                  ITSPSGELQCPGWCSLQKTL-ALQREIGVGYEPQCVQDGQRFSPLQCDL--SYCWCVSQ-   1174
Danio_rerio                      ITNSSGELQCPGWCSLQKTL-ALQREVALGYEPECVQDGQHFSPLQCDL--SDCWCVSD-   1174
Astyanax_mexicanus               ILSPTGDLKCPGWCALQKKL-VAEREMGVGYEPQCVEGGERFSPLLCDA--SYCWCVSE-   1171
Clupea_harengus                  LQGPDGEYKCPGWCALQRAR-ALSREVGVGFEPVCEQEGAAFSALQCDQ--ADCWCVSED   1196
Oryzias_latipes                  GRGATGQLKCPSWCELQRRQ--------------CG-PDGSFAPLQCDV--TSCWCVTE-   889
Xiphophorus_maculatus            SRSAAGELTCPSWCELQRLR--------------CR-PDGSFDPLQCDV--TSCWCVSE-   887
Stegastes_partitus               TPNTAGQLTCPSLCELQGLQ--------------CR-PDGSFVPLQCDT--ATCWCVSE-   895
Cynoglossus_semilaevis           TRNSAEQLTCPTWCQLQGGQ--------------CD-SDGLFIPLQCDE--TSCWCVSV-   878
Lepisosteus_oculatus             AQNKAGSPQCPGWCEMLKSQ-VLRRDIGTGYVPECQEEGQLFSSVQCDQ--SSCWCVFQ-   1182
Lampetra_fluviatilis             RDDPSGDIRCPSLCELRRRQ-ALLRDAGLGSVPECD-SRGDYAARQCAT--GTCWCASA-   1281
Lampetra_planeri                 RDDPSGDIRCPSLCELRRRQ-ALLRDAGLGSVPECD-SRGDYAARQCAE--GTCWCASA-   1219
Petromyzon_marinus               HPSPTTNAESPSLCELRRRQ-ALLRNAGLGSVPECD-SRGDYAARQCAD--GTCWCAGA-   1233
```

HINGE

```
Homo_sapiens                        SGEEVPGTRVT----GGQPACESPRC-PLPFNA-SEVVGGTILCETISG-PTGSAMQQCQ     1246
Pan_troglodytes                     SGEEVPGTRVA----GGQPACESPRC-PLPFNA-SEVVGGTILCETTSG-PTGAAIQQCQ     1244
Gorilla_gorilla                     SGEEVPGTRVA----GGQPACESPRC-PLPFNA-SEVVGGTILCETTSG-PTGSAIQQCQ     1246
Pongo_pygmaeus                      SGEEVPGTRVA----GSQPACESPRC-PLPFNV-SEVVGGTILCETTSG-PIGAAIQQCQ     1246
Macaca_mulatta                      SGEEVPETRVA----GSQPACESPRC-PLPFNT-LEVVGGTILCETASG-PTGAAIQQCQ     1246
Macaca_fascicularis                 SGEEVPETRVA----GSQPACESPRC-PLPFNT-LEVVGGTILCETASG-PTGAAIQQCQ     1246
Rattus_norvegicus                   SGEEVPGTRVV----GTQPACESPQC-PLPFSG-SDVTDGVVFCETASS-SGVTTVQQCQ     1247
Mus_musculus                        SGEEVPGTRVV----GTQPACESPQC-PLPFSG-SDVADGVIFCETASS-SGVTTVQQCQ     1247
Cavia_porcelus                      SGEEVPGTHVV----GSQPACERPQC-PLPFLSLSDMPGGVILCKRASG-LGEATVQQCQ     1242
Canis_lupus_familiaris              SGEEVPGTRVA----GSQLACESPRC-PLPFNT-TDVDGGVIVCERASG-PGGAPVQRCQ     1245
Panthera_leo                        TGEEVPGTREA----GRQPACESPQC-PLPFNT-SDVAGGAILCERAAG-PGRSSIQRCQ     1247
Bos_taurus                          SGEEVPGTRVA----GSQPACESPQC-PLPFSV-ADVAGGAILCERASG-LGAAAGQRCQ     1246
Trichechus_manatus_latirostris      SGEEVPGTRVA----GSQTACENPQC-PLPFSV-ADVVGGAVLCEEASG-QGAATIQQCL     1245
Columbia_livia                      NGEEVPGTRIS----GGRPACASPQC-ALPFGA-SSVPNGAVFCENVSG-QA-SSVQQCW     1240
Taeniopygia_guttata                 NGEEVPGTRVN----GARPECASPQC-ALPFGA-SAVANGAVLCETISG-QA-PGIQQCQ     1244
Larus_michahellis                   NGEEVPGTRVS----GGRPACASPQC-ALPFGT-SSVLNGAVFCENISE-QA-SSVQQCW     1245
Gallus_gallus                       NGEEVPGTRVS----GGRPACESPQC-AVPFGT-SSLLNGAVFCDSVSE-QM-LGVQQCR     1237
Struthio_Camelus                    NGEEVPGTRVS----GGRPACESPQC-ALPFGA-SSILNGAIFCEKVSG-QT-SSVQQCQ     1219
Chelonia_mydas                      NGEEVPGTRVR----GERPACESPQC-LLPFNV-SYVVNGALFCANISD-QN-QSSQKCQ     1237
Alligator_mississippiensis          SGEEVPGSRVS----GERPACESPQC-VLPFNV-SHVVNGVIFCETVSD-QN-RNSQQCQ     1219
Crotalus_tigris                     NGQEAPGTRVN----GQKSACERPRC-PLPYNA-SSLTNGGVFCNI-NT-AN-TKTQQCQ     1231
Python_bivittatus                   NGQEAPGTRVN----GQRPACERPQC-PLPYNA-SYLSNGGVFCNT-PT-AT-SKTQQCQ     1173
Eublepharis_macularius              NGGEAPGTRDS----NRGPACERPRC-HLPFDA-PGIINGGVFCNNVSS-AT-SKTQECR     1225
Xenopus_tropicalis                  DGEEAFGTRQNVSEMGKPPTCKSPIC-PLAFSA-QDIRHGAVFCENILE-AG-ITFQKCQ     1249
Aquarana_Catesbeiana                TGEEATGTRVPMTER-TGFLCQAPVC-PLPFGI-KEIKHGSIFCSELVD-SG-KKVQKCQ     1231
Carassius_auratus                   SGKELPATFTNR-STGQTPACDIPEC-PLPFED---ISHGAVLCSTVIV-AG-QQTQRCE     1222
Cyprinus_carpio                     SGKELPVTRTNR-STGQTPACDIPEC-PLPFED---ISHGAVLCSTVIV-AG-QQTQSCE     1227
Danio_rerio                         SGKELPMTRSPR-STGQTPACNIPEC-PLPFGD---ISHGAVLCSSTNV-PS-QQMQRCE     1227
Astyanax_mexicanus                  SGQELPATRTTR-SLGKTPSCDYPQC-PLPFG----VSHGAAVCGNESV-AG-EQRQRCR     1223
Clupea_harengus                     SGLELPDTRTPR-RTGQTPSCDRPQC-PLAFGP-ASVSHGSMVCH-----PD-QRGQRCE     1247
Oryzias_latipes                     YGQEVVGTRTPQ-LTGVTPSCDRPLC-PA-----PTITHGALVCRSAAN-----GPQSCD     937
Xiphophorus_maculatus               DGQEVAGTRSPR-QTGRTPSCDRPLC-PA-----PNITHGALVCRPAAD-----GQQSCD     935
Stegastes_partitus                  DGQEVGGTRTPR-QAGRSPSCDRPLC-PA-----PVITHGALLCRPVVD-----GRQSCD     943
Cynoglossus_semilaevis              DGQEVGGTRTPQ-ETQLKPSCDRPLCRPS-----PGINHGALLCHTLSD-----GRQTCE     927
Lepisosteus_oculatus                NGQEAPGTRLRQ-VSGQTPKCDTPQC-PLPFGV-PGINNGAVFCKDILE-NG-QRRQQCQ     1237
Lampetra_fluviatilis                GGEEIPGTRRHAG--EPGVVCHEPRC-QLRQDE-RRVGGWALVCEDGTAAAAAAGLQRCS     1337
Lampetra_planeri                    GGEEIPGTRRHAG--EPGVVCHEPRC-QLPQDE-RRVGGWALVCEDGTNA-AAAGLQRCS     1274
Petromyzon_marinus                  GGEEIPDTRRHAG--EPGVVCHEPRC-ELPQDE-RRAGRWALVCEDGAAATAAAGLQRCL     1289


Homo_sapiens                        LLCRQGSWSVFPP--------GPLICSLESGRWES-QLPQPRACQRPQLWQTIQTQGHFQ     1297
Pan_troglodytes                     LLCRQGSWSVFPP--------GPLICSLESGRWES-QPPQPRACQRPQLWQTMQTQGHFQ     1295
Gorilla_gorilla                     LLCRQGSWSVFPP--------GPLICSLESGRWES-QPPQPWACQRPQLWQTIQTQGHFQ     1297
Pongo_pygmaeus                      LLCRQGSRSVFPP--------GPLICSLESGRWES-QLPQPRACQRPQLWQTIQTQGHFQ     1297
Macaca_mulatta                      LLCRQGSRSVFPR--------GPLICSLESRRWES-QLPQPRACQRPQLWQTIQTQGHFQ     1297
Macaca_fascicularis                 LLCRQGSRSVFPR--------GPLICSLESRRWES-QLPQPRACQRPQLWQTIQTQGHFQ     1297
Rattus_norvegicus                   LFCRQGLWNVFSP--------GPLICNLESQRWVT--LPLPRACQRPQLWQTMQTQAHFQ     1297
Mus_musculus                        LLCRQGLRSAFSP--------GPLICSLESQHWVT--LPPPRACQRPQLWQTMQTQAHFQ     1297
Cavia_porcelus                      LFCRQGYRSAFPS--------GLLVCNLESKRWVS-QPPQPLACQRPQLWQTVQTGAHFQ     1293
Canis_lupus_familiaris              LLCRRGYRSAFLP--------GPLVCSLEEGRWLS-QPPQPQACQRPHLWQTFQTQGQFQ     1296
Panthera_leo                        LRCLRGYWSAFPP--------GPLLCGLEEGRWVS-QPPQPHACQRPQLWQTVQTRARVL     1298
Bos_taurus                          LRCSQGYRSAFPP--------EPLLCSVQRRRWES-RPPQPRACQRPQFWQTLQTQAQFQ     1297
Trichechus_manatus_latirostris      LRCRQGYRSAFPP--------RPLVCRVESRSWEA-EPPQPRACQRLQPWQTVQTRAQLQ     1296
Columbia_livia                      LVCRQGFYSSSSS--------TSFQCDVQRRRWVS-AAPLYQACQKLQLFQTVQAQTHFQ     1291
Taeniopygia_guttata                 LVCRQGFHSAVPS--------SPSQCDARQRRWVS-AAPLPQACQKLQLFQTVQAQTHFQ     1295
Larus_michahellis                   LVCRQGFHSSSPS--------TSFQCDVQQRRWVS-AVPLYQACQKPQSFQTVQAQTHFQ     1296
Gallus_gallus                       LVCRQGYNSAASN--------TLFQCDVQKRLWIS-AAPLYHACQKLQLFQTVQAQTHFQ     1288
Struthio_Camelus                    LVCRQGFRSAPSS--------TSFQCDVQQRQWVL-AAPLPQACQKLQLFQTVQAQTHFQ     1270
Chelonia_mydas                      LICRQGFHSAFSN--------VTFLCDRESRRWIS-EPPLSQSCQKLQLFQNVQAQTHFE     1288
Alligator_mississippiensis          LVCRRGFQSAFSS--------KRFLCDVESRRWIS-DPPLSQTCQKLQLFQTVQAQTQFQ     1270
Crotalus_tigris                     MICYPGYQDVFSQ-------GDSLLCDTESLLWLT-SPPHSHACQRLQPFQSVQIQTQFQ     1283
Python_bivittatus                   VICSPGYQDVFSQ-------SDPLLCDTESLRWLA-TPPHSQACQRIQPFQSVQIQTQFQ     1225
Eublepharis_macularius              VICYPGFYPTFSR-------GEMLLCDVETGLWEG-DPPHSQTCQKLQPFQSVQTQSRFQ     1277
Xenopus_tropicalis                  LVCRKGYQNVLLQ--------DTFTCNTDTRLWGV-QVPHPGSCQKIQSFQSIETQAQFQ     1300
Aquarana_Catesbeiana                LVCQKGYTNLIST--------STFTCDPNSYLWIP-QPPHSQPCQRLEPFSMIQTRAWFQ     1282
Carassius_auratus                   LFCDQGYVNTLPV--------TSFLCDPQGKNWLD-DAPQSYACQKPQGLQTVQVSVQLK     1273
Cyprinus_carpio                     LFCDQGYVNTLPV--------ASFMCDPQAKNWLD-DAPLSYACQKPQVLQTVQVSLQLK     1278
Danio_rerio                         LYCDQSYVNTLPV--------ASFLCDPQAKNWLE-DAPLSNACQKPQVLQTVQVSVQLQ     1278
Astyanax_mexicanus                  VYCQQGYLNTLQV--------DSFSCDPVTKTWIS-DAPLSYACQRVQPLQTVRVSSVLQ     1274
Clupea_harengus                     MLCHQGYVSVFPA--------ATFLCDPKSRMWLS-DAPLANSCQRSQVFQTVSSGAALQ     1298
Oryzias_latipes                     LICNYGYENSLPV--------SSFLCETESGRWSGDIKPLGGACQISQPLQTLSLSQLWS     989
Xiphophorus_maculatus               LVCHHGYQNSLPV--------SSFLCGVESRQWEEEDRPLSGACQIAQPLQSFSSSQIWS     987
Stegastes_partitus                  LVCHHGYQNSLRV--------SSFQCETASQRWGEDDAPLSGACQISQPLQSVSSSQLWL     995
Cynoglossus_semilaevis              LICHHGYHNTLPI--------SSFLCDTESQQWVGDFTPVGGACQISQPLQVLSHSQLFS     979
Lepisosteus_oculatus                LTCHQGYQNTLPE--------SRFLCDVVTSNWLS-AQPLPDACQKPQVFQAVEAHTTFQ     1288
Lampetra_fluviatilis                LACLRGFARTLDGASHGGGAPQEFQCNATSGEWIG-SSPQPDACQEIHPWQAVQLSISFT     1396
Lampetra_planeri                    LACLRGFARTLDGASHGGGAPQEFQCNATSGEWIG-SSPQPDACQEIHPWQAVQLSISFT     1333
Petromyzon_marinus                  LACLRGFAHTLDGAGHDGGAPREFQCNATSGEWIG-SSPQPDACQEIHPWQAVQLSISFT     1348
```

```
Homo_sapiens                        LQLPPGKMCSADYADLLQTFQVFILDELTARGFCQIQVKTFGTL----VSIPVCNNSSVQ     1353
Pan_troglodytes                     LQLPPGKMCSADYASLLQTFQVFILDELTARGFCQIQVKTFGTL----VSIPVCNNSSVQ     1351
Gorilla_gorilla                     LQLPPGKMCSADYTGLLQTFQVFILDELTARGFCQIQVKTFGTL----VSIPVCNNSSVQ     1353
Pongo_pygmaeus                      LQLPPGKMCSADYAGLLQTFQVFILDELTARGFCQIQVKTFGTP----VSIPVCNNSSVQ     1353
Macaca_mulatta                      LQLPPGKMCSADYAGLLQAFQVFILDELTARGFCQIQVKTFGTL----VSVPVCDNSSVQ     1353
Macaca_fascicularis                 LQLPPGKMCSADYAGLLQAFQVFILDELTARGFCQIQVKTFGTP----VSVPVCDNSSVQ     1353
Rattus_norvegicus                   LLLPPGKMCSIDYSGLLQAFQVFILDELITRGFCQIQVKTFGTL----VSRTVCDNSSIQ     1353
Mus_musculus                        LLLPPGKMCSVDYSGLLQAFQVFILDELIARGFCQIQVKTFGTL----VSSTVCDNSSIQ     1353
Cavia_porcelus                      ILLPPGKRCSADYSGLLQAFRVFILDELMARGFCHVQVPTAGTL----VSHPVCDNSSVQ     1349
Canis_lupus_familiaris              LQLPPGKMCSADYAGLLPAFQVFLADELEARGFCQIQAKTLGTP----ISIPVCDSSTVQ     1352
Panthera_leo                        LRLPPGKMCSADYAGLLTAFRVFISDELVARGFCQIQAKAFGSP----VSIPVCDGSTVQ     1354
Bos_taurus                          LLLPLGKVCSADYSGLLLAFQVFLLDELTARGFCQIQVKTAGTP----VSIPVCDDSSVK     1353
Trichechus_manatus_latirostris      LLLPPGKTCSADYSGLMQAFQVLVLNELTARGFCQIQVKTFGIP----VSIPVCNDSTVQ     1352
Columbia_livia                      LLLPPERTCSSDYSGLLQAFQILISDELKARGLCHLQVNAFGDT----GSVSVCDDSTVY     1347
Taeniopygia_guttata                 LRLPPEKTCSSDYSGLLQAFQIFILDELKARGLCHLQVNAFGKT----GSVSMCDDSTVY     1351
Larus_michahellis                   LLLPPEKTCSSDYSGLLQAFQIFILDELKARGLCHLQVNAFGNT----GSLSVCDDSTVY     1352
Gallus_gallus                       LLLPPQKTCSSDYSGLLQAFQIFILDELKARGFCHLQVNAFGNT----GSVSVCDDSSVY     1344
Struthio_Camelus                    LLLPPQKTCSSDYSGLLQAFQIFILDEMKARGFCHLQVNAFGNT----RSVSVCDDSTVY     1326
Chelonia_mydas                      LLLPSEKSCSSDYAGLLQAFQIFILDALKALGFCHIQVNTFGNP----GSVSVCDDSAVL     1344
Alligator_mississippiensis          LLLPSGKACSSDYSGLLQAFQIFILDELKARGLCHIQVNAFGNS----RPVSLCDDSTVL     1326
Crotalus_tigris                     LILPPEKTCSPDYSGLLETFQIFILDELKSRGFCHIQVNTLGNL----VSVPVCGDSAVL     1339
Python_bivittatus                   LILPPKKTCSPDYSGLLETFQIFILDELKARGLCNIQVNTLENL----VSIPVCGDSAVL     1281
Eublepharis_macularius              LLLPPEKACSPDYSGLLEAFQIFIQDEMEARGLCHIQVNALEHL----VSVPVCGDSAVR     1333
Xenopus_tropicalis                  LLLPVGKKCMAEYSGLLQAFRTFILDDLRARGLCQIQVNYFGSR--GAGAVSVCDESTVF     1358
Aquarana_Catesbeiana                LLLPAGKMCTDDYAGLLEAFRTFILDDLRARGLCHIQVTTLGGR----GTVPLCDDSTIH     1338
Carassius_auratus                   LSLAEGQQSC--NSQSFEFQAALLRDMRATGLCSLQLTSSGQT----SSVAVCDESSVS     1326
Cyprinus_carpio                     LSLAEGQQSC---SSQSVELQAALLRDMRATGLCSLQLTLSGQT----SSVAVCDESSVS     1331
Danio_rerio                         LSLNEGQQSC---STQSVDLQAALLRDMRATGLCSLQLTSSGQT----SSVSVCDESSVS     1331
Astyanax_mexicanus                  LTLAAGQQGC---SGQRSALQASLLHDLRAAGLCSLQLS-SGQY----SSFSVCDNSSVL     1326
Clupea_harengus                     LSLKAGQRSC---DAQRSELQSSLLRDMRATGLCSVQLSDASRT----SGVSVCDESSVS     1351
Oryzias_latipes                     LA-SC---------SQADQTRSLLFNMMTSRGLCSAQLGSSGR------SVSLCDDSSVS     1033
Xiphophorus_maculatus               LSVSC---------SRISSLSSLLFSLLASRGLCSAQLPS-GR------SVALCGDSAVG     1031
Stegastes_partitus                  LPSSC---------SQISTLQSQLFNMMTSRGLCSAQLPVLGR------SVSLCDESSVS     1040
Cynoglossus_semilaevis              LSSPC---------SQISTLQPLLVRSLTSLGLCSMQLPVSTS------SVLVCDQSSVH     1024
Lepisosteus_oculatus                LSLAEGKNTC---RAVRSPFQSSLLQHLRGRGLCSLQMNSFGKS----VSVSVCDESTVT     1341
Lampetra_fluviatilis                LRFPEEKVCNPEYEGLLESFNSFLLADLTARGFCHLSNGAQSAPGGGSGAVAVCDAASVR     1456
Lampetra_planeri                    LRFPEEKVCNPEYEGLLESFNSFLLADLTARGFCHLSNGAQSAPGGGSGAVAVCDAASVR     1393
Petromyzon_marinus                  LRFPEEKMCNPEYEGLLESFNSFLLADLTARGFCHLSNGV------GSDAVAVCDAASVR     1402
```

```
Homo_sapiens                        VGCLTRERLGVNVTWKSRLEDIPVASLPDLHDIERALVGKDLLGRFTDLIQSGSFQLHLD     1413
Pan_troglodytes                     VGCLTSERLGVNVTWKSRLEDIPVASLPDLHDIERALVGKDLLGRFTDLIQSGSFQLHLD     1411
Gorilla_gorilla                     VGCLTRERLGVNVTWKSRLEDIPVASLPDLHDIERALVGKDLLGRFTDLIQSGSFQLHLD     1413
Pongo_pygmaeus                      VGCLSRERLGVNVTWKSRLEDIPVASLPDLHDIERALVGKDLLGRFTDLIQSGSFQLHLD     1413
Macaca_mulatta                      VGCLTREYLGVNVTWKSRLEDIPVASLPDLHDIERALVGKDLLGRFTDLIQSGSFQLHLD     1413
Macaca_fascicularis                 VGCLTREYLGVNVTWKSRLEDIPVASLPDLHDIERALVGKDLLGRFTDLIQSGSFQLHLD     1413
Rattus_norvegicus                   VGCLTAERLGVNATWKLQLEDISVGSLPNLHSIERALMGQDLLGRFANLIQSGKFQLHLD     1413
Mus_musculus                        VGCLTAERLGVNVTWKLQLEDISVGSLPDLYSIERAVTGQDLLGRFADLIQSGRFQLHLD     1413
Cavia_porcelus                      VGCLTAEHLEVNVTWKSQLKDIPAASLPDVHDIERALVGKDLLGHFTALVQSGTFQLHLD     1409
Canis_lupus_familiaris              VECVTGERLGVNVTWKLHLEDVPPASLPDLHDIEEALVGKDLIGRFTDLIQSGGFQLHLD     1412
Panthera_leo                        VECVSGERLGVNVTWKLRLQDVAPASLPDVRDIEEALVGKDLIGRFADLIQSGEFQLYLD     1414
Bos_taurus                          VECLSRERLGVNITWKLQLVDAPPASLPDLQDVEEALAGKYLAGRFADLIQSGTFQLHLD     1413
Trichechus_manatus_latirostris      VECLTVERLGVNVTWKSRLEDIPVASLPDLHDIEKALVGKDLVGRFADLIRSGEFQLHLD     1412
Columbia_livia                      VECLSVDRLGVNVTWRTQLENIPAASHPDLHDIENAIVGENLIGAFMNEIKGGYFLLHLD     1407
Taeniopygia_guttata                 VQCLGVDRLGVNITWRTQLENIPTASLPDLHDIENAIVGENLIGAFIREIKDGVFLLHLD     1411
Larus_michahellis                   VECLSVDRLGVNVTWRTQLENIPATSLPDLHDVENAIVGEKLIGAFIKEIEDGGFLLHLD     1412
Gallus_gallus                       VECLSVDRLGVNVTWRALLENIPAVSLPDLHDIENALVGEKLIGAFIKEIEDGRFLLYLD     1404
Struthio_Camelus                    VECLSTDRLGVNVTWKAPLGNIPATSLPDLHDIENAIVGENLIGAFIKQIKEGDFLLYLD     1386
Chelonia_mydas                      VECLSVDRLGVNVTWKVQLEGIPAEGLPDLHDIEKAIVGENLIGRFVALVESGGFVLHLD     1404
Alligator_mississippiensis          VNCLTVDRLGVNITWKAQLEDIPAASLPDLHDIENTIVGDNLIGRFVKLIKSGGFLLHLD     1386
Crotalus_tigris                     VQCAAADRLGVNVTWQAMLKNVPVTALPDLYDIEKAMVGENLLGRFEELIKTGGFILHLD     1399
Python_bivittatus                   VQCMAADRLGVNITWQAMLRDVPAASLPDLYDIEKAMAGENLLGRFEELIKSGGFVLHLD     1341
Eublepharis_macularius              VECLAVNRLGVNVTWRALLEDVPAASLPDLHNIEKAMVGD-LIGRFETLVRSGGFVFHLD     1392
Xenopus_tropicalis                  VECLAVDRLGVNVTWRTQLDDFPESFMPSLHDIEDALIGDNLVGRFASVISSGNYSLTLD     1418
Aquarana_Catesbeiana                VECLTSTRMGVNVTWTAQIYKIPESLLPAFLDIENALAVENVVGRLLSVIGSGSYSLDLD     1398
Carassius_auratus                   LECLNDKEVTAHITFRARLSDLPITSLPDLHDIDVALSEERLLNGVEELIQSGSYRSIFL     1386
Cyprinus_carpio                     LECLNDKEVTAHITFRARLSDLPITSLPDLHDIDMALSEERLLNGVKELIQSGSYRSIFL     1391
Danio_rerio                         LECSNDKEVTARITLMARLSDLPLAALPDLHDIDTALSEERLLNGLKKLILSGSYQSIFL     1391
Astyanax_mexicanus                  LECESEQSLTATITLKARLSDLPISALPDLYDIDSVFSSEKILKGVMEIIRSGSYRSVYV     1386
Clupea_harengus                     LECVTEDRLTAHFTLTARLSALPSSALPDLHDVDLALSSERFLAGLLDLIGDERYQSLMS     1411
Oryzias_latipes                     LQCDGAPT--LKVSWSAALADLPAAYLPDLHDVALFLNETRLLDGVRGLLLDLGSAMTPE     1091
Xiphophorus_maculatus               LWCDAGDSVRLTLRWTADLSDLTTSDLPHLHHMAIFLNESRLLEGVQGILGNLRSLLTSD     1091
Stegastes_partitus                  LRCSGDDSLRLTLTWTAAMSDLPTTDLPDLHDVGLFLNVSRLLEGVQGILGNIRSTLTSE     1100
Cynoglossus_semilaevis              LQCDSDQSLKLTVTWRVAFSDVPTSDLPHLHDIVLFLNESRLLEGVQQLLGNIKSTL---     1081
Lepisosteus_oculatus                LECLSAKRMRANITWKAQLGDIPTEALPDLHDIDNAFVSRNLVKGIIDLIGSGVFWPVLD     1401
Lampetra_fluviatilis                VRCQDALRLLVNVTWRSSLSRLPVQSYPDLHNIERAFVAH--AQRFAQLLDDRGYRLALN     1514
Lampetra_planeri                    VRCQDALRLLVNVTWRSSLSRLPVQSYPDLHNIERAFVAH--AQRFAQLLDDRGYRLALN     1451
Petromyzon_marinus                  VRCQDALRLLVNVTWRSSLSRLPVQSYPDLHNIERAFVAH--AQRFAQLLDDRGYRLALN     1460
```

```
                                                                       TG type 2-1
Homo_sapiens                        SKTFPAE-TIRFLQGD-HFGTSPRTWFGCSEGFYQVLT--SEASQDGLGCVKCPEGSYSQ   1469
Pan_troglodytes                     SKTFPAETTIRFLQGD-HFGTSPRTWFGCSEGFYQVLT--SEASQDGLGCVKCPEGSYSQ   1468
Gorilla_gorilla                     SKTFPAETTIRFLQGD-HFGTSPRTWFGCSEGFYQVLT--SEASQDGLGCVKCPEGSYSQ   1470
Pongo_pygmaeus                      SKTFPADTTIRFLQGD-HFGTSPRTWFGCSEGFYQVLT--SEASQDGLGCVKCPEGSYSE   1470
Macaca_mulatta                      SKTFPADTTIRFLQGD-HFGTSPRTWFGCLEGFYQVLT--SEASQDGLGCVKCPEGSYSQ   1470
Macaca_fascicularis                 SKTFPADTTIRFLQGD-HFGTSPRTWFGCLEGFYQVLT--SEASQDGLGCVKCPEGSYSQ   1470
Rattus_norvegicus                   SKTFSADTILYFLNGD-RFVTSPMTQLGCLEGFYRVST----TSQDPLGCVKCPEGSFSQ   1468
Mus_musculus                        SKTFSADTTLYFLNGD-SFVTSPRTQLGCMEGFYRVPT----TRQDALGCVKCPEGSFSQ   1468
Cavia_porcelus                      SKTFLADTSIRFLQGDSSFGASPRTWFGCMEAFYQVLAT-SEASRDPLGCVMCPEGSYSL   1468
Canis_lupus_familiaris              SKTFPADTSIYFLQGD-RFGTSPRAWFGCLEGFHQVLAP-SNAPQDPWGCVKCPEGSYFQ   1470
Panthera_leo                        SKTFPADPSIYFLQED-GFGTSPRTWFGCLEGFHQVLAT-GSAPRDPPGCVKCPEGSYFQ   1472
Bos_taurus                          SKTFSADTSIRFLQGD-RFGTSPRTQFGCLEGFGRVVAA-SDASQDALGCVKCPEGSYFQ   1471
Trichechus_manatus_latirostris      SKTFPADTSIHFLQGD-RFGTSPRAWFGCLEGFSRVSAS-SKTSQESLGCVKCPEGSYFQ   1470
Columbia_livia                      SKQFLADSVVDFPRDE-EFDLSPTVKLGCSNGFRRTSSP-GKVAPNSQGCVVCPPGSHFQ   1465
Taeniopygia_guttata                 SKQFIADSVIDFPRDE-EFDLSPRVQLGCRSGFRRISSP-GKAGPNSQGCVVCPPGSYFQ   1469
Larus_michahellis                   SKQFLADSFVDFPWDE-EFDLSPSVQLGCRSGFRRISSP-GKTVPNSQGCVVCPPGSHFQ   1470
Gallus_gallus                       SKQFLADSFIDFPQDE-EFDLSPRVQLGCRSGFQRISSP-GKIVPNAQGCVVCPPGTNFQ   1462
Struthio_Camelus                    SKQFVADSFIDFPRDE-EFDLSPRVRLGCRSGFQRTSSP-GRTVPNSQGCVICPPGSHFE   1444
Chelonia_mydas                      SKRFPADTSIHFPRDE-DFDLSPSVQLGCRSGFRRSSSL-GRAIPNSQGCVVCPSGSSFQ   1462
Alligator_mississippiensis          SKQFPAATSISFLRDE-DFDLSPSVQLGCRSGFRRSSSP-GTAISNSQGCVVCPPGSYFQ   1444
Crotalus_tigris                     NKQFQADTSIRSPGRG-DSGMSPKVSLGCKSGFYKVLTT-GEVIPNSQGCALCPPGSSFQ   1457
Python_bivittatus                   NKQFQADTSIRFPGRG-DSGTSPKVYLGCKSGFQTVLTT-GEVVPNSQGCAVCPPGSSFQ   1399
Eublepharis_macularius              SKRFQADTSITFLRDV-DFNISSRVHVGCKSGFRKALTA-EEVAPNSRGCVVCPPGAYFQ   1450
Xenopus_tropicalis                  SKEFVADRSVLFPHME-KLGSFPQTSLGCAQGFLKSLNS---KTGQFNGCVICPAGSFAQ   1474
Aquarana_Catesbeiana                SKLFVADKSDFLPKDS-----STQVRLSCAKGFQKLLISRADEIGNVAGCGICPVGSYSE   1453
Carassius_auratus                   YER-----SLA---------LSIPPSFSCTDGYKQL--------PQSTGCVVCPAGSFSS   1424
Cyprinus_carpio                     YDR-----SLA---------LSIPPSFSCMDGYKQL--------PQSTGCVVCPAGSFSS   1429
Danio_rerio                         SDH-----SLA---------LSSPPSFSCMTGYSQL--------PQSTGCVLCPAGSFFS   1429
Astyanax_mexicanus                  SEP-----TVA---------ISSPASFSCSVGYQQL--------PDSAGCVVCPAGSFSS   1424
Clupea_harengus                     AKR-----AEA---------ILSETSFSCAAGYQMI--------PDAKGCVVCPAGTFSS   1449
Oryzias_latipes                     P----------------KLVSMTTPSFGCYRGYRLD--------SDGNGCVVCPAGSFSQ   1127
Xiphophorus_maculatus               P----------------KLVLVALPTFGCSHGYRLS--------SDGAGCVICPAGSFHQ   1127
Stegastes_partitus                  P----------------KLVSVTMPSFSCSRGYRLS--------NDGEGCVVCPAGSFSR   1136
Cynoglossus_semilaevis              -------------------VSTTPPIFGCSHGYHLT--------NNSEGCLVCPAGTFAT   1114
Lepisosteus_oculatus                SEPFPPDVSYN---------FVPEVQFGCTQGYRPL--------LGVRGCAICPTGTFVS   1444
Lampetra_fluviatilis                GRDYSSEGAARFRKDE-LYDTSPAFRMACKPGYVRI----------AQGCAACPRGSLWR   1563
Lampetra_planeri                    GRDYSSEGAARFRKDE-LYDTSPAFRMACEPGYVRI----------AQGCAACPRGSLWR   1500
Petromyzon_marinus                  GRDYSSEGAARFRKDE-LYDTSPAFRVACEPGYVRI----------AQGCAACPRGSLWR   1509

                                        TG type 2-2      TG type 2-3     Spacer 1   TG type 1-11
Homo_sapiens                        DEECIPCPVGFYQEQAGSLACVPCPVGRTTISAGAFSQTHCVTDCQRNEAG---------   1520
Pan_troglodytes                     DEECIPCPVGFYQEQAGSLACVPCPVGRMTISAGAFSQTHCVTDCQRNEAG---------   1519
Gorilla_gorilla                     DEECIPCPVGFYQEQAGSLACVPCPVGRMTISAGAFSQTHCVTDCQRNEAG---------   1521
Pongo_pygmaeus                      DEECIPCPVGFYQEQAGSLACVPCPVGRMTISAGAFSQTHCVTDCQRNEAG---------   1521
Macaca_mulatta                      DEECIPCPVGFYQEQAGSLACVPCPAGRMTISAGAFSQTHCVTDCQGNEAG---------   1521
Macaca_fascicularis                 DEECIPCPVGFYQEQAGSLACVPCPAGRMTISAGAFSQTHCVTDCQRNEAG---------   1521
Rattus_norvegicus                   DGKCTPCPAGTYQGQAGSSACIPCPRGRTTTITGAFSKTHCVTDCQRDEAG---------   1519
Mus_musculus                        DGRCTPCPAGTYQEQAGSSACIPCPRGRTTITTGAFSKTHCVTDCQKNEAG---------   1519
Cavia_porcelus                      DGQCVPCPVGFYQEEAGSVACVPCPAGRTTAAIGAFSQRHCVTDCQRNEAG---------   1519
Canis_lupus_familiaris              KEICIPCPVGFYQEQAGSMDCVPCPVGRTTISPGAFSHTHCVTDCQRSEVG---------   1521
Panthera_leo                        EEICIPCPVGFYQEQAGSVACVPCPRGRTTVSAGAFSHTHCVTDCQKSEAG---------   1523
Bos_taurus                          DEQCIPCPAGFYQEQAGSLACVPCPEGRTTVYAGAFSQTHCVTDCQKNEVG---------   1522
Trichechus_manatus_latirostris      DEQCVPCAVGFYQERAGGSACVPCPVGRTTMSAGAFSQTHCVTDCQRNEAG---------   1521
Columbia_livia                      NDECVPCPIGYYQYQTGRSSCIKCPVGKTTNSYGAFSAEHCITYCQSNEQG---------   1516
Taeniopygia_guttata                 DDECIPCPLGYYQYQTGHSFCIKCPVGKTTNSYGAISADHCVTPCQGGEQG---------   1520
Larus_michahellis                   NDECIPCPLGYYQYQTGRSSCIKCPVGKTTNSYGAFSADHCITYCQTNEQG---------   1521
Gallus_gallus                       NGECIPCPLGYYQYQTGRLSCIKCPVGKTTISYGAFSADDCITYCQGNEQG---------   1513
Struthio_Camelus                    NEECVPCPLGYYQYQTGSSSCIKCPVGKTTNSYGAFSADHCITYCQSNEQG---------   1495
Chelonia_mydas                      TDECIPCPPGFYQLQTGSSSCIRCPVGKTTISAGAFRADHCITDCERNRQG---------   1513
Alligator_mississippiensis          NGECTPCPHGSYQSQTGSPSCIKCPLGKTTVSTGAFTADHCITDCQQNRQS---------   1495
Crotalus_tigris                     NEKCIPCPSGFYQDQAGSNFCVKCPPGKFTISTGAFKEDHCVTDCQRNEQA---------   1508
Python_bivittatus                   NESCIPCPSGFYQDRAGSNFCVKCPLGKFTVSAGAFRADHCITDCQRNEQI---------   1450
Eublepharis_macularius              TGHCIPCPSGFYQEQTGSLSCNKCPLGKTTVSIGAFKAEHCVTACQMDERD---------   1501
Xenopus_tropicalis                  NEVCKLCPLGFYQVEPGTASCTKCPLGTTTVNGGAYNRSHCVTSCQRNSIG---------   1525
Aquarana_Catesbeiana                DGECVPCPHGFYQEKSGSSQCLKCPLGTTTAYAGAYKPNQCVTHCQANTRG---------   1504
Carassius_auratus                   GGVCTLCPRDTFQVEEGQAFCSPCPSGTSTVAQGAFSASHCLTECQQS--K---------   1473
Cyprinus_carpio                     GGVCTLCPRDTFQVEEGQVFCSPCPSGTSTEAQGAFSASHCLTECQQS--K---------   1478
Danio_rerio                         GGVCTPCPRDTFQEDEGQVFCSPCPAGTSTVAQGAFSASHCLTECKKS--K---------   1478
Astyanax_mexicanus                  GGRCDVCPRGSYQAEAGQDFCSPCPSGTSTAALGASSPSHCLSECQQS--K---------   1473
Clupea_harengus                     GAVCSACPTGTYQERAGQSVCRRCPMGTSTVAQGAFSATQCVTSCQQS--G---------   1498
Oryzias_latipes                     EGACLLCPEGTYQEEKGRGFCKSCPKGSSP--VGASSVTQCETRCERR--G---------   1174
Xiphophorus_maculatus               QGACLLCPQGTYQELEGRDFCTRCVRGSSP--AGASAANQCETDCERR--G---------   1174
Stegastes_partitus                  EGVCLLCPQGTYQDEEGRDFCNRCPRGSSP--AGASSIDLCVTECRGR--G---------   1183
Cynoglossus_semilaevis              EGACLLCPPGTYQDEEGRSQCSKCPTGSSP--AGATSVNQCVTECQKR--G---------   1161
Lepisosteus_oculatus                GAVCTPCPHGSYQDQEGSHFCKQCPQGWSTGSTGAFKATHCLTDCQNNPLG---------   1495
Lampetra_fluviatilis                DGVCAPCPTDHYQDEPGQTACTPCPADSGTLTTGAAAASQCRTPCEREALRRGQLGAGER   1623
Lampetra_planeri                    DGVCAPCPTDHYQDEPGQTACTPCPADSGTLTTGAAAASQCRTPCEREALRRGQLGAGER   1560
Petromyzon_marinus                  DGGCALCPADHYQDEAGQTACTPCPADTGTLTAGAAAAFQCRTPCEREALRRGQLGAGER   1569
```

Spacer 2

```
Homo_sapiens                       --LQCDQNGQYRASQKDRGSGKAFCVDGEGRRLPWWETEAPLEDSQCLMMQKFEKVPESK   1578
Pan_troglodytes                    --LQCDQNGQYRASQKDRGSGKAFCVDGEGRRLPWWETEAPLEDSQCLMMQKFEKVPESK   1577
Gorilla_gorilla                    --LQCDQNGQYRASQKDRGSGKAFCVDGEGRRLPWWETEAPLEDSQCLMMQKFEKVPESK   1579
Pongo_pygmaeus                     --LQCDQNGQYRASQRDGGSGKAFCVDGEGRRLPWWETEAPLEDSQCLMMQKFEKAPESK   1579
Macaca_mulatta                     --LQCDQNGEYRASQRDRGSGKAFCVDGEGWRLPWSETEAPLEDSQCLMMQKFEKAAESK   1579
Macaca_fascicularis                --LQCDQNGEYRASQRDRGSGKAFCVDGEGWRLPWSETEGPLEDSQCLMMQKFEKAAESK   1579
Rattus_norvegicus                  --LQCDQNGQYQANQKDMDSGEVFCVDSEGQRLQWLQTEAGLSESQCLMMRKFEKAPESK   1577
Mus_musculus                       --LQCDQNGQYQASQKNRDSGEVFCVDSEGRKLQWLQTEAGLSESQCLMIRKFDKAPESK   1577
Cavia_porcelus                     --LQCDQDGQYQASQKDDASGTAFCMDSEGHRLLWSQTEAPLSDSQCLMMQKFEKAPEWK   1577
Canis_lupus_familiaris             --LRCDQDGQYRASQRDGAGGKAFCVDGEGRRLPWSETEAPLTDSQCQMMQKFEKAPDSK   1579
Panthera_leo                       --LQCDQDGQYRASQRDRGSGRAFCVDSEGRRLLWLETEAPLSESQCLMMQKFEKAPESK   1581
Bos_taurus                         --LQCDQDSQYRASQRDRTSGKAFCVDGEGRRLPWTEAEAPLVDAQCLVMRKFEKLPESK   1580
Trichechus_manatus_latirostris     --LQCDQDGQYRARQTDRASGKAVCVDREGQRLPGSETEALLTESQCLMMRKFEKLSESK   1579
Columbia_livia                     --LQCDEDGQYRPSQQDPDTKKSFCVDSSGAALDWKETDDLLTDYQCLVLRKFERVPSSR   1574
Taeniopygia_guttata                --LQCDGEGQYRPAQQDNATQKSLCVDSFGAALDWTETEDLLTNEQCLVLRKFERVPASS   1578
Larus_michahellis                  --LQCDENGQYRPSQQDPDTKKSFCIDSFGAALDWTETDDLLIDNQCLVLRKFERVPASR   1579
Gallus_gallus                      --LQCDGDGQYRPSQQDPDTEKSFCVDDFGAALDWTETDDLLTDYECLVLRKFEKVPAST   1571
Struthio_Camelus                   --LQCDEDGQYRPSQQDLDTKKWFCVDSFGTALGWTEADDLLTDYQCLVLRKFDRVPASK   1553
Chelonia_mydas                     --LQCSEDGQYRPSQADPITKKSFCADSLGMTLDWTETDHPLTDSQCLVLRKFERVPESK   1571
Alligator_mississippiensis         --LQCDEEGQYRPSQQDPITKKSFCVDNLGTRLGWTETDDLLTDSQCLVLRKFERVPTSK   1553
Crotalus_tigris                    --ERCDNNGQYRPSYQDPSTQKSFCVDSNGKRLDWTETGGPLADSQCSVLRHFEKVPASN   1566
Python_bivittatus                  --ERCDENGQYRPSYKDSSTQKSFCADIKGKRLDWTEADGPLTDSQCSVLRRFEKVPASN   1508
Eublepharis_macularius             --LQCDEDGQYIPYFEDPATNKSFCVDSLGKILEWTETDGPITVSWCQVLGRFERVPATK   1559
Xenopus_tropicalis                 --LLCDANGQYQPSQKDVATNRYFCVDVFGKRLIWSETDTELTDAQCLLLRKFELVPEDK   1583
Aquarana_Catesbeiana               --LRCDDKGQFLPLQRDEPTEKYFCVDNSGKKLTWTETDQELQNDQCLLLQKFDLVPEDG   1562
Carassius_auratus                  --RSCTKKGEFLSAQKDPMSGRWFCVSSQGEELSWTSSDDPVTVEECKVMEKFAVVASSS   1531
Cyprinus_carpio                    --LSCTKKGEFLSAQKDLMSGRWLCVSSQGEELSWTSSDAPMTVEECKVMEKFEVVAPSS   1536
Danio_rerio                        --LSCTTKGDFLSAQKDPVSGKWRCVSAQGEELTWTSSDGLLTVEECKVMEKFEVVSSSS   1536
Astyanax_mexicanus                 --LSCTDRGEFLSAQKHFQSGKWLCITSKGEELTWTSADEAVSAEQCKVLEKFEAVPVSS   1531
Clupea_harengus                    --MRCTERGDFLSTQQDILSGRWQCVTLSGQPLPWTSAEEPITEQDCAALQNFDAVPDSQ   1556
Oryzias_latipes                    --LRCSQTGDFLPAQPDLLTSRWSCFSKDGVELEWTSSSKMLTDEDCSVLTRFQEVPGSE   1232
Xiphophorus_maculatus              --LRCTQRGDFLPAQPNFLSGGWRCVSSEGAELEWTSREERLADEECSVLSLFQAVPGSD   1232
Stegastes_partitus                 --LRCSQEGDFLPAQPDLQSGRWRCFSSEGMELEWTSSDATLTDAECSVFRKFEAVPRSK   1241
Cynoglossus_semilaevis             --MRCSERGDFLSAQPGL-SGRWRCLSSEGVELEWTSSDKALTDEECTIFISVQTVSGSD   1218
Lepisosteus_oculatus               --LECTEEGAYREAQRDAASEKWFCVTDRGEKLNWTVTMETLTVSQCKVMQKFEDVPKTQ   1553
Lampetra_fluviatilis               VELYCDPEGLY-------------------------EGEKPAKGGQDEVRRMFEEVPSED   1658
Lampetra_planeri                   VELYCDPEGLY-------------------------EGEKPAKGGQDEVRRMFEEVPSED   1595
Petromyzon_marinus                 LELYCDPEGLY-------------------------EGEKPAKGGQDEVRRMFEEVPSED   1604
```

TG type 3a-1

```
Homo_sapiens                       VIFDANAPVAVRSKVPD--SEFPVMQCLTDCTEDEACSFFTVSTTEPEISCDFYAWTSDN   1636
Pan_troglodytes                    VIFDANAPVAVRSKVPD--SEFPVMQCLTDCTEDEACSFFTVSTMEPEISCDFYAWTSDN   1635
Gorilla_gorilla                    VIFDANAPVAVRSKVPD--SEFPVMKCLTGCSEPRSSWA-----TEPD---------PDN   1623
Pongo_pygmaeus                     VIFDANAPVAVRSKVPD--SEFPVMQCLRDCAEDEACSFFTVSTTEPEISCDFYAWTSDN   1637
Macaca_mulatta                     VIFDANAPVAVRSKVPD--SEFPVMQCLTDCAEDEACSFLTVSMTEPEVSCDFYAWTSDN   1637
Macaca_fascicularis                VIFDANAPVAVRSKVPD--SEFPVMQCLTDCAEDEACSFLTVSMTEPEISCDFYAWTSDN   1637
Rattus_norvegicus                  VIFDASSPVIVKSRVPS--ANSPLVQCLADCADDEACSFVTVSSMSSEVSCDLYSWTRDN   1635
Mus_musculus                       VIFDANSPVIVKSSVPS--ADSPLVQCLTDCANDEACSFLTVSTMESEVSCDFYSWTRDN   1635
Cavia_porcelus                     VNFSAAAPGMVRSKVPG--SDSPLMQCLADCAEDQGCSFFTMFTTESEVACDFYNWTSDN   1635
Canis_lupus_familiaris             VVFSANVTAVGRSKVSG--SGSPLLQCLTDCALDETCSFLAVSTAGSEVSCDFYGWTSDS   1637
Panthera_leo                       VTFRADVTVAGRSRVPG--SESPLTQCLTDCALDEACSFLAVSTAGSEVLCDFYAWTNNH   1639
Bos_taurus                         VIFSADVAVMVRSEVPG--SESSLMQCLADCALDEACGFLTVSTAGSEVSCDFYAWASDS   1638
Trichechus_manatus_latirostris     VIFSADIPFVISSKVPG--SESPLMQCLADCADDEVCIFLTVSTGGPEVSCEFYAGTNDS   1637
Columbia_livia                     LIYGSEEAQILQSQTFQGDLQSPAFQCISGCEREDTCGFVTLTPSAAEILCELYSAAEVN   1634
Taeniopygia_guttata                LIYGSEEAQILQSQSFEGNLQSTYFQCISGCEREDACGFAAVTPAGTEVLCELYSAAEAN   1638
Larus_michahellis                  LIYGLEEAQILQSQSFGGDLQSSAFQCISGCEREEACGFATVTPAGAGVLCELYSAAEVN   1639
Gallus_gallus                      FVYGSDNVQILKSQSFGRDLQSLTFQCISDCDKEEACSFAAVTPSGAEILCDLYSAAEVN   1631
Struthio_Camelus                   LIHGSEENEILQSQSFGGDLQSPVLQCISDCEKEEACGFATVTSAGTQILCELYSAAEIN   1613
Chelonia_mydas                     LIYGAEDARILRSETIQGDSQSALLQCISDCETDESCGFITVSAPGSEVLCELHSAAETN   1631
Alligator_mississippiensis         IIYSMEDAEVFRSKTVQGDLQSPFWQCISDCVQEESCSFVTVSTTGSKVLCELYRAVETN   1613
Crotalus_tigris                    LIFTTKESETIGTKTLQGTPMNTLQQCVAECEEDESCGFLTLSAMGSETLCELYSARESN   1626
Python_bivittatus                  LMFSTEESETIGTKAFQGDPMSTLQQCIAECEKDESCGFLTMSAMGSETLCELYSARESN   1568
Eublepharis_macularius             LSILTDELETIRSEIFRGEPDSALWPCAVECEEDESCGFFALSAAETGMLCELYGADESN   1619
Xenopus_tropicalis                 ILFIGEDSGVTQPKAN-G-QIKSLLDCVTDCGTEESCDYISVSTNGSGLICEQYSANESN   1641
Aquarana_Catesbeiana               LIVDDKNMGIAQSTSVKT-LRKTFLECIDDCAKDESCDYVTVSGNGTGVTCEQYNGEGSN   1621
Carassius_auratus                  LLLKSESAEILSSESSTQPKEKQLRKCVLDCALEDSCQHLAIFRDKEKTFCELYSTSVDN   1591
Cyprinus_carpio                    LLLKSESAEILSSDSSTQPKEKQLMKCVLDCALEDSCQHVAIFSDKEKTYCELYSTSADN   1596
Danio_rerio                        LLLKSESADTLSSDSSSQPKETQLRKCVLDCAQDDSCLYVAVFSDEEKTHCEMYSTSADN   1596
Astyanax_mexicanus                 LLLDSDDAVVLWSDSSSQPLETQLRKCVFECAEDDSCQHVAVFRDGAQTLCDLYSTSEEN   1591
Clupea_harengus                    LVFDAQDSILLRSETVDLPLENQLRKCVIDCSLDKTCQHVAAYSAEGRPHCDLYSTATAN   1616
Oryzias_latipes                    LMAAAEDVDILLTLT------SDLLTCIDACAAEPSCHHVALFSR--QTQCELYSTDPIN   1284
Xiphophorus_maculatus              LMVGTGDTDVLRTIT------SDLSACLHACAAEPSCHHVALFDR--RTRCDLYSS--LN   1282
Stegastes_partitus                 LTVGAEAAEVLQTMT------SDLTTCIQACAAEPSCHHVALFNR--QTQCELYSTNTVN   1293
Cynoglossus_semilaevis             LILGTEDVELLQTIT------SDLKGCIQACAAEPSCHHVALFDS--QTQCELYSTQTLN   1270
Lepisosteus_oculatus               WILDANDAVIMSSKTSNESLESYLKTCIAECAADESCLHLAVFTEGSTAHCDLYSTSNAN   1613
Lampetra_fluviatilis               LVLGAVDAVALRIRTETGPVQEVEPRCLQECLKEERCDFLVFSTQGDVSSCHFYNGSRES   1718
Lampetra_planeri                   LVLGAVDAVALRIRTETGPVQEVEPRCLQECLKEERCDFLVFSTQGDVSSCHFYNGSRES   1655
Petromyzon_marinus                 LVLGAVDAVALRIRTETGPVQEVEPRCLQECLKEERCDFFVFSTQGDVSSCHFYNGSRES   1664
```

```
Homo_sapiens                          VACMTSDQKRDALGNSKATSFGSLRCQVKVRSHG-Q-DSPAVYLKKGQGSTTT-LQKRFE   1693
Pan_troglodytes                       VACMTSDQKQDALGNSKATSFGSLRCQVKVRSRG-Q-DSPAVYLKKGQGSTTT-LQKSFE   1692
Gorilla_gorilla                       NAASFHLQKQDALGNSKATSFGSLRCQVKVRSRG-Q-DSPAVYLKKGQGSTTT-LQKSFE   1680
Pongo_pygmaeus                        VDCMTSDQKQDALGNSKATSFGSLHCQVKVRSRG-Q-DSPAVYLKKGQGSTTT-LQKSFE   1694
Macaca_mulatta                        VACMTSDQKQDALGNSKATSFGSLRCQVKVRSRG-Q-DSPAVYLKKGQGSTTT-LQKSFE   1694
Macaca_fascicularis                   VACMTSDQKQDALGNSKATSFGSLRCQVKVRSRG-Q-DSPAVYLKKGQGSTTT-LQKSFE   1694
Rattus_norvegicus                     FACVTSDQEEDAVDSLKETSFGSLRCQVKVRNSG-K-DSLAVYVKKGHEFTAS-GQKSFE   1692
Mus_musculus                          FACVTSDQEQDAMGSLKATSFGSLRCQVKVRNSG-K-DSLAVYVKKGYESTAA-GQKSFE   1692
Cavia_porcelus                        FACTTSAQEQDALRSLQVTGFGSLMCQVKVRNRN-Q-KSLAVYLKKGQEFTAA-GQKTFE   1692
Canis_lupus_familiaris                IACTTSAQHQDTLGNSEATSFGSLRCQMTVRSGA-Q-DSLAVYLKKGQEFTTT-SQKSFE   1694
Panthera_leo                          MACTTSAQHQDTLGNSGATGFGSLRCQVTVRNGT-P-DSPAVYLKKGQEFTTT-SRKSFK   1696
Bos_taurus                            IACTTSGRSEDALGTSQATSFGSLQCQVKVRSRE-G-DPLAVYLKKGQEFTIT-GQKRFE   1695
Trichechus_manatus_latirostris        IVCTASDQEENALGNVMATSFGSLRCQVKVRRQD-Q-DSLAVYLKQGQQLTTM-SQKSFE   1694
Columbia_livia                        FNCTTSGLVQGVLGNPTATSIGRLSCLLHVRNPE-G-DAVMVYLKRGQEFNSS-GLKTFE   1691
Taeniopygia_guttata                   FNCTTSGLVQGVLGNPAATSISGLSCLLRVRNPE-G-DALMAFLKRGLEFLSS-GPKAFE   1695
Larus_michahellis                     FNCTTSGLVQGVFGNPAATSIDRLSCLLHVRNPE-G-DAVTAYLKRGQEFNSS-GLKTFE   1696
Gallus_gallus                         FNCTISGLVQGALGNPAATSIGRLSCLLYIRNPE-G-DAVMVYLKKAQEFNSS-GLKTFE   1688
Struthio_Camelus                      FNCTASELVQGALGNPAATSVGSLSCLLDIRNQE-G-DAVMVYLKKGQEFNSS-GLKTFE   1670
Chelonia_mydas                        FNCTTSGLMKGVMGNSDATSIANLSCLLKVRNPE-K-DEVTVYLKKGQEFTIS-GLKIFE   1688
Alligator_mississippiensis            FNCTTSGLMQGALGNSAASSIAHLSCLLKIRNQE-K-DEVAIYLKKGQEFTTS-GLKTFE   1670
Crotalus_tigris                       YNCSTSGLVHGAWDNSLTTNIASLKCLIKVRSPD-KKDSLNVYLKKGQEFTTA-GVKRFE   1684
Python_bivittatus                     YNCSTSGLVHGVWDNSLTTSIASLSCLIKVRSPD-KKNSLSVYLKKGQEFTTA-GVKTFE   1626
Eublepharis_macularius                YNCSASGLAEGVWDNSATTRIAHLSCLIKVRSHD-Q-DAVGVYLKQGQEFTTA-GVKTFE   1676
Xenopus_tropicalis                    IVCTDVQQKESVLGNTDSVKVENPKCQMKIRQRN-T-GSHTVYGKKGYTFLRS-GQALFE   1698
Aquarana_Catesbeiana                  IICSTSNQVQTALGNSAATSMERLRCQHKVRLSS-I-EDISVYRKKGEEFPST-GHRDFK   1678
Carassius_auratus                     VECRTSEQSKGFLGNDGAEMFQTLNCVLKIK-AD-E-PGLIVLRKKGHEFSTA-GPKSFE   1647
Cyprinus_carpio                       VECRTSEQSKGFLGNDGAETFQTLNCVLKIK-GD-E-PNLMVLRKKGHEFSTA-GLKSFE   1652
Danio_rerio                           VECRTTEPSKGFLGNDGAEAFQTLICVLKIK-GD-E-PDLTVLRKKGHEFSTA-GLKRFE   1652
Astyanax_mexicanus                    VDCRTSQRSKGFLGNEGAESFQTLSCLLKVKIGD-K-PNLTVLRKKGHEFSTA-GPKKFE   1648
Clupea_harengus                       VECRTSEQTKGFLGNPVAELYQSLRCSLKLRVER-R-PNLTVLRKKGHEFSAVVRQKTFE   1674
Oryzias_latipes                       TVCNASQPSKGFLGNPEAERYDWLSCSLKVRGGA---SGLLVLRKKGEEFGLKLQQRAMV   1341
Xiphophorus_maculatus                 VVCSASQQARGFLGNPRAERFHKLRCSLSVRGGA---SDLLVLRKNGAEL----QQQSFV   1335
Stegastes_partitus                    TDCNTSQQSKGFLGNLQAERFDGLSCSVRVRGGA---SDLLVIRKKGAEFSSQQQQQSFV   1350
Cynoglossus_semilaevis                TQCVTSQQTKGFLDNPEAEVFDWVHCSVKVRGSA---SDLLVIRKKGEGS-----SSTFV   1322
Lepisosteus_oculatus                  IKCTTSEKATGFLGNSVTDVYESLTCLVKVQDGG-G-QALIVLRKKGHEFTTR-SQKMFE   1670
Lampetra_fluviatilis                  YQCETAPEAPGFLGDPDSSWVERLSCRPRVS-L-SPGSVFTVYRKKGHELPSW-DGQLYR   1775
Lampetra_planeri                      YQCETAPEAPGFLGDPDSSWVERLSCRPRVS-L-SPGSVFTVYRKKGHELPSW-DGQLYR   1712
Petromyzon_marinus                    YQCEKAPEEKSALGISLCVVFIAIKCFVRGVGFHVSSASGRFYFHAGHELPSW-DGQLYR   1723
```

TG type 3b-1

```
Homo_sapiens                          PTGFQNMLSGLYNPIVFSASGANLTDAHLFCLLACDRDLCCDGFVLTQV--QGGAIICGL   1751
Pan_troglodytes                       PTGFQNMLSGLYNPIVFSASGANLTDAHLFCLLACDRDLCCDGFVLTRV--QGGAIICGL   1750
Gorilla_gorilla                       PTGFQNMLSGLYNPIVFSASGANLTDAHLFCLLACDRDLCCDGFVLTQV--QGGAIICGL   1738
Pongo_pygmaeus                        PTGFQNMLSGLYNPIVFSASGANLTDAHLFCLLACDHDLCCDGFVLTQV--QGGDVICGL   1752
Macaca_mulatta                        PTGFQNMLSGLYNPIVFSASGANLTDAHLFCLLACDRDLCCDGFVLTQV--QGVAIICGL   1752
Macaca_fascicularis                   PTGFQNMLSGLYNPIVFSASGANLTDAHLFCLLACDRDLCCDGFVLTQV--QGVAIICGL   1752
Rattus_norvegicus                     PTGFQNVLSGLYSSVVFSALGTNLTDTHLFCLLACDQDSCCDGFIVTQV--KEGPTICGL   1750
Mus_musculus                          PTGFQNVLSGLYSPVVFSASGANLTDTHTYCLLACDNDSCCDGFIITQV--KGGPTICGL   1750
Cavia_porcelus                        PTGFQNTLSGLYDSVMFSASGANLTDAHLFCLLTCERDACCDGFILSQM--PGGPLICGL   1750
Canis_lupus_familiaris                QTGFQNMLSGLYRPVIFPASGADLTAAHLFCLLACDRDSCCDGFILAQL--QGGPVICGL   1752
Panthera_leo                          PTGFQNALSGTYGAVVFSASGADLTGAHLFCLLACDRDSCCDGFILAQV--QGGPLICGL   1754
Bos_taurus                            QTGFQSALSGMYSPVTFSASGASLAEVHLFCLLACDHDSCCDGFILVQV--QGGPLLCGL   1753
Trichechus_manatus_latirostris        QTDFQNTLSGMYSPFVFSASGANVTDAHLFCLLACDRDPCCDGFILAQV--QGGPIICGL   1752
Columbia_livia                        RTSFQNVLSGVYHSVVLSASSASLTDAQLFCRQTCSRDSCCDGFILSQIALEGGLILCGL   1751
Taeniopygia_guttata                   RTGFQNVLSGVFRSLVLPAAGTSPTDAGLLCRQECSRDSCCDGFILSQLLLDGGTILCSL   1755
Larus_michahellis                     RTGFQNVLSGVYRSVVLSAASTSLTDAQLFCRQICSRDSCCDGFILSQIALDGGTILCSL   1756
Gallus_gallus                         RTSFQNVLSGVYRSMALSAAGTSLTDAQLFCRQTCSRDSCCDGFILSQIALDGGTILCSL   1748
Struthio_Camelus                      RTNFQNVFSGIYRSLVLSAASTSLTDAQLFCRQTCSRDSCCDGFILSQIALDGGTVLCSL   1730
Chelonia_mydas                        LTGFQNVLSGVYSSVVLSAAGTSLTDVHLFCRQACSQDPCCDGFILSHVVLDGGTILCGL   1748
Alligator_mississippiensis            MTDFQNVFSGIYSSMVLSAASTSLTDVHLLCRQACSQDPCCDGFILSQMTLHGGTILCGL   1730
Crotalus_tigris                       KTDFQDVVSGVYSISVVSASGTSLTDVHLFCRQACHQDACCDGFILSQLILDGGSIACSL   1744
Python_bivittatus                     KTDFQDVVSGVYSIRVVSAAGTSLTDVHLFCRQVCHQDACCDGFILSQLILDGGSIACSL   1686
Eublepharis_macularius                MTDFQDVVSGVYRNLVLSAAETSLTDVHLFCRQACSQDACCDGFILSQVILNGGSILCGL   1736
Xenopus_tropicalis                    RTGFGNIISGAYKTLVFSADGASLSEAHLFCHQVCRKDPCCDGFILSQIILNKGTILCGL   1758
Aquarana_Catesbeiana                  RTDFGNTVSGVYNATLFPGGGASLTDGYYVCRQMCEQDACCTGFILSKIILNKGTIVCGL   1738
Carassius_auratus                     KLSFRKTGSGVYRTLVFDAHGTTLADVHRFCVNSCSRESCCDGFILNQNVLDGGSIMCGL   1707
Cyprinus_carpio                       RLSFRKTGSGVYRTLVFDAHGTTLADVHRFCVNSCSRENCCDGFILNQNVLDGGSIMCGL   1712
Danio_rerio                           RLSFRKAGSGVYRTLVFDARGSALADVHRFCVDSCGRETCCDGFILNQNVLNGGSIMCGF   1712
Astyanax_mexicanus                    RLTFHKASSGVYRTLVFEASGASLADVHRFCVDTCRQESCCDGFILNQNVLNGGSLMCGL   1708
Clupea_harengus                       RLAFRKAGSGVYRTLVFAASGAELADAHRFCADICSQEACCDGFILNKNILDGGSLMCGL   1734
Oryzias_latipes                       KTRMTKALSGVFRTQVFSSRRTSLSDAHRFCQDGCSRHECCDGFVLNQNTLEGGSVLCGW   1401
Xiphophorus_maculatus                 RMRMRKAVSGVFRTQVFSSRNTSLSDARRFCQDGCSRDACCHGFILNQNVLDGGSLLCGW   1395
Stegastes_partitus                    RTRMMKALSGVFRTQVFSSRQTSLSDAHRFCQDGCSRDACCDGFILNQNSLDGGTLLCGW   1410
Cynoglossus_semilaevis                RSPMMKAVSGVFKTQMFSSTRTTLTDAYRFCQDGCSQDSCCEGYVLNQIIVNGGFHLCGW   1382
Lepisosteus_oculatus                  RQDFHKAGSGVYQTSVFEARGVSLTDVHHFCQEACNQETCCDGFILNQNVLNGGTIMCGL   1730
Lampetra_fluviatilis                  PTGFGNAASGAYRTLALPAGTTTLPDAHLYCRSACSAQACCDGFLLKELPLDNGVLVCAL   1835
Lampetra_planeri                      PTGFGNAASGAYRTLALPAGTTTLPDAHLYCRSACSAQACCDGFLLKELPLDNGVLVCAL   1772
Petromyzon_marinus                    PTGFGNAASGAYRTLALPAGTTTLPDAHLYCRSACSAQACCDGFLLKELP--PGVLVCAL   1781
```

```
Homo_sapiens                        LSSPSVLLCNVKDWMDPSEAW-ANATC-PGVTYDQESHQVILRLGDQEFIKSLTP-----    1804
Pan_troglodytes                     LSSPSVLLCNVKDWMDPSEAW-ANATC-PGVTYDQESHQVILRLGGQEFIKSLTP-----    1803
Gorilla_gorilla                     LSSPSVLLCNVKDWMDPSEAW-ANATC-PGVTYDQESHQVILHLGGQEFIKSLTP-----    1791
Pongo_pygmaeus                      LSSPSVLLCNVKDWMDPSEAR-ANATC-PGVTYDQESHQVTLHLGGQEFIKSLTP-----    1805
Macaca_mulatta                      LSSPNVLLCNVKDWMDPSEAR-ANATC-PGVTYDQESRQVTLHLGGQEFIKSLTP-----    1805
Macaca_fascicularis                 LSSPNVLLCNVKDWMDPSEAR-ANATC-PGVTYDQESRQVTLHLGGQEFIKSLTP-----    1805
Rattus_norvegicus                   LSAPDILVCHINDWRDASDTQ-ANGTC-AGVTYDQGSRQMTMSLGGQEFLQGLTL-----    1803
Mus_musculus                        LSSPDILLCHINDWRDTSATQ-ANATC-AGVTYDQGSRQMTLSLGGQEFLQGLAL-----    1803
Cavia_porcelus                      LSSPDVLVCNDDDWKDSSKVQ-ANTTC-PGVTYEQGSHQVTFLLGGQEFVKSLAP-----    1803
Canis_lupus_familiaris              LSSPDVLLCHVKDWRDPTEAQ-ANATC-PGVTYDRGSRQATLRLGGQEFR-VPVA-----    1804
Panthera_leo                        LSSPDVLLCNVKDWRDPTEAR-ANATC-PGVTYDQGSRPAALRLGGQEFE-IPAA-----    1806
Bos_taurus                          LSSPDVLLCHVRDWRDPAEAQ-ANASC-PGVTYDQDSRQVTLRLGGQEIR-GLTP-----    1805
Trichechus_manatus_latirostris      LSSPSILLCSVKDWRDAPEAQATNTTC-PGVTYDQGSRQVTLRLGGQEFIQSLTP-----    1806
Columbia_livia                      ISYPDVLICNANDWSPKVTSV-FDGIC-GDVSYDEKEMMFFFTLGGQVFSGTSKL-----    1804
Taeniopygia_guttata                 LSSPDVLICNAKGWSPAPASA-MGEMC-KGVSYDEKEKRFSFTLGGQVFSGTSQL-----    1808
Larus_michahellis                   MSSPDVLICNANDWSPTPTSG-MDGIC-KGVSYDEKEKMFSFTLGGQVFSGTSKV-----    1809
Gallus_gallus                       MTSPDVLICNANDWSPTQTSV-INEIC-RGLSYDEGGKIFSFTLGGQVFSGTSEL-----    1801
Struthio_Camelus                    MSYPDVLLCNANDWSPTAKSV-RDGIC-KGVSYDEKEKMFSFMLGGQIFSGSFKL-----    1783
Chelonia_mydas                      MSYPDVLICHRNDWNRNSKPG---GIC-KGVNYNEKKQQFLFSLGGQEFAGTSEL-----    1799
Alligator_mississippiensis          MSYPDVLICNTNDWSQTSIPG-TDGIC-KGVNCDEKDKKFSFSLGGQVFTGTCEL-----    1783
Crotalus_tigris                     LSSPSALICNLHDWDGSSTRR-QGDIC-QRTKYSEGKSQYTIRIGDQVLAGASEL-----    1797
Python_bivittatus                   LSSPGALICNLNDWDGSSTRQ-QGDIC-QRTKYKEGRSQYTVRIGDQVLAGASEL-----    1739
Eublepharis_macularius              MSYPDVFICNNKDWDGSTTHG-REDRC-QREKYDEGRKEYTFRLGGQVLTGTSEV-----    1789
Xenopus_tropicalis                  LSSPDVLLCNVNDWSGTSLLG-GEGVC-KGVRSNKEQKMFSFFLGGQEFTGSYSMLSESI    1816
Aquarana_Catesbeiana                LRSPNVLLCNKNDWSSTSRLG-GEGIC-KGVASNKQKKMFSFFLGGQEFTGSYSLLSQSV    1796
Carassius_auratus                   LSAPSVLQCSEADWDARG-VASSSRICGAGIQYSKQLKRFTFNFGGQNFTINDAALPAS-    1765
Cyprinus_carpio                     LSAPSVLQCSEADWDARG-VASSSRICGAGVQYSKQLKRFTFNFGGQNFTITDAALPAS-    1770
Danio_rerio                         LTAPTVLQCSESDWDVKS-LSSSSRICGAGVQYSKQLKRFTFSFGGQNFTITDAALPAS-    1770
Astyanax_mexicanus                  LSFPSVLQCSEGDWDVSS-VASSNRICGAGLKYNKQQKQFTFEFGGQNFTITDSALPVS-    1766
Clupea_harengus                     LSHPDVLLCSEADWDVAG-LGQSSRICGAGVMYSKQNKRFTFNFGGQIFNITDTALPAS-    1792
Oryzias_latipes                     LRSPSVLSCQDGDWDVSSQGP-ANRTCGAGLSYNEEQQSFLFDFGGQAFTILDYEVPAD-    1459
Xiphophorus_maculatus               LRAPSVLMCEDEDWDVIGQGP-ANRICGAGLSYSEQQRSFELDFGGEKFTITDGALPGD-    1453
Stegastes_partitus                  LRAPSVLMCGEQDWDVIGQGA-ANRICGAGLTYNKEQRSFVFDFGGQKFTITDSALPAD-    1468
Cynoglossus_semilaevis              LSSPSVLMCGKGDWDVIGHGT-ANRICGAGLTYNEQQKSFLFDFGGQTFNITDSSLPSD-    1440
Lepisosteus_oculatus                LSYPDVFLCRDQDYDDAAAQTGGKRVCGAGVKYNKLRKQFTFSFGGQNFTLNDMALPAT-    1789
Lampetra_fluviatilis                LSGPTVLTCRAPGWPRTADEY-GDGEC-SGLRVHKPSRTFGFSLGGVHYNASNPPSAVA-    1892
Lampetra_planeri                    LSGPTVLTCRAPGWPRTADEY-GDGEC-SGLRVHKPSRTFGFSLGGVHYNASNPPSAVA-    1829
Petromyzon_marinus                  LSGPTVLTCRAPWWPRTADEY-GDGEC-SGLRVHKPSRTFGFSLGGVHYNASNPPSAVA-    1838

Homo_sapiens                        -------------LEGTQDTFTNFQQV-----YLWKDSDMGSRPESMG-CRKDTVPRPAS    1845
Pan_troglodytes                     -------------LEGTQDTFTNFQQV-----YLWKDSDMGSRPESMG-CRKDTVPRPAS    1844
Gorilla_gorilla                     -------------LEGTQDTFTNFQQV-----YLWKDSDMGSRPESVG-CRKDTVPRPAS    1832
Pongo_pygmaeus                      -------------LEGTQDTFTNFQQV-----YLWKDSDMGSRPESMG-CRKDTVPRPAS    1846
Macaca_mulatta                      -------------LEGTQDTFTNFQQV-----YLWKDSDMGSRPESMG-CRKDTVPRPAS    1846
Macaca_fascicularis                 -------------LEGTQDTFTNFQQV-----YLWKDSDMGSRPESMG-CRKDTVPRPAS    1846
Rattus_norvegicus                   -------------LEGTQDSFISFQQV-----YLWKDSDIGSRPESMG-CGRGMVPKSEA    1844
Mus_musculus                        -------------LEGTQDSFTSFQQV-----YLWKDSDMGSRPESMG-CERGMVPRSDF    1844
Cavia_porcelus                      -------------LERAPRTVIGFQQV-----YLWKDSDMASRPESLG-CRKDVVPRSES    1844
Canis_lupus_familiaris              -------------LEGTSGTVTSFQRV-----YLWKESDMGSRSESMG-CRRDMEPRPAS    1845
Panthera_leo                        -------------LEGTQDAVTSFQQV-----YLWKDSDMGSRSESMG-CRKDTEPGPTS    1847
Bos_taurus                          -------------LEGTQDTLTSFQQV-----YLWKDSDMGSRSESMG-CRRDTEPRPAS    1846
Trichechus_manatus_latirostris      -------------LDRTQDTFSSFQRV-----YLWKDSDMGSRSESVG-CRRHMAPRPAS    1847
Columbia_livia                      -------------VQEAERNFTTFQQV-----YLWRGSDMVTRTRTSE-CDAAALKTEND    1845
Taeniopygia_guttata                 -------------MEETERNFTTFQEI-----YLRRDSDMVTRMKTSS-CEAAALKTKDD    1849
Larus_michahellis                   -------------AEEAERNFTTFQQV-----YLWRGSDMVTRTKTSA-CNAAALKTEND    1850
Gallus_gallus                       -------------SEEAERNFTTFQQV-----YLWRGSDMVTRPETSE-CDSTALKTEDN    1842
Struthio_Camelus                    -------------AEERERNFTTFQHV-----YLWRGSDMVTRLKTSA-CDAAAFKMEND    1824
Chelonia_mydas                      -------------SEEMEGAFSSFQRV-----YLWRGSDMITRTKSSA-CDAAVLQTQTE    1840
Alligator_mississippiensis          -------------PEGPEGFFTSFQRV-----YLWRGSNMITRRPSSA-CGAAVLQTQTG    1824
Crotalus_tigris                     -------------AEDTEKFFTSFQRV-----YFWKDSDMMTRTGPTM-CDATLVPPQSK    1838
Python_bivittatus                   -------------ADNAENPFTSFQRV-----YLWRDSTMMTRNGWTM-CDATVVPTQTK    1780
Eublepharis_macularius              -------------TKEKDNSFVSFQRV-----YLWRDSDMLTRTRSAG-CDVAAFQIQTE    1830
Xenopus_tropicalis                  GKVEYTTELTAEVKEEIQQLFVTFQRV-----FLKRDGSGN--G-LSD-CSAGSVQELNN    1867
Aquarana_Catesbeiana                GMVEYSTELTKDIKDEIQALFTGFQSV-----FLRTDVRTV--TNEAE-CP--QRASQNN    1846
Carassius_auratus                   ----------SKNKTGYQETLFSFQRI-----YLWKESDMNTRPKTPSACIGSAVLEDSK    1810
Cyprinus_carpio                     ----------SKNKTGYQETLFSFQRI-----YLWKESDMNTRPKTPSACSGSAVVEDSK    1815
Danio_rerio                         ----------SKNKTGYQETLISFQRV-----YLWKESDMNTRAKTLSACSGSAVGEDSR    1815
Astyanax_mexicanus                  ----------SKNRTDYEATIIGFQRV-----YLWKESDMSTRPKTDGSCGRLALLGTPA    1811
Clupea_harengus                     ----------SKNKTDYQASIIGFQRV-----YIWKESDMSTRPKTPA-CASSPSQTATG    1836
Oryzias_latipes                     ----------GKNKIDHQPSIVSFQAV-----YLSSG--------S--LCAAD----LS     1490
Xiphophorus_maculatus               ----------SRTRKDYQATIISFQAI-----YLNTAAAAGGSGSA--SCGAAE----VP    1492
Stegastes_partitus                  ----------SKNKKDYQASIISFQSI-----YLNSDSAVSSSGSS--SCAAAE----LS    1507
Cynoglossus_semilaevis              ----------SKN-KDYQSSIVSFQGI-----YLNTGSPVSGSP-L--SCAAAE----PS    1477
Lepisosteus_oculatus                ----------SKNKTDYQATLISFQRV-----YLWKDSDMNTRPKSAPGCASIV-PPEGK    1833
Lampetra_fluviatilis                --------------STAAAATAATAAAAQVPAAALCGAPMCSPGLCRE--SFAV---TAS    1933
Lampetra_planeri                    --------------STAAAATAATAAAAQVPAAALCGAPMCSPGLCRE--SFAV---TAC    1870
Petromyzon_marinus                  --------------STAAAAAAASQVIYL-----WAGSEAGSRDVCLH--TKAYTPQPGE    1877
```

TG type 3a-2

```
Homo_sapiens                        PTEAGLTTELFSPVDLNQVIVNGNQSLSSQKHWLFKHLFSA-QQANLWCLSRCVQEHSFC      1904
Pan_troglodytes                     PTEAGLTTELFSPVDLNQVIVNGNQSLSSQKHWLFKHLFSA-QQANLWCLSRCVQEHSFC      1903
Gorilla_gorilla                     PTEAGLTTELFSPVDLNQVIVNGNQSLSSQKHWLFKHLFSA-QQANLWCLSRCVQEHSFC      1891
Pongo_pygmaeus                      PTETGLTTELFSPVDLNQVIVNGNQSLPSQKHWLFKHLFSA-QQANLWCLSRCVQEHSFC      1905
Macaca_mulatta                      PTETGLTTDLFSPVDLDQVIVNGNRSLPSQKHWLFKHLFSA-QQANLWCLSRCVQEHSFC      1905
Macaca_fascicularis                 PTETGLTTDLFSPVDLDQVIVNGNRSLPSQKHWLFKHLFSA-QQANLWCLSRCVQEHSFC      1905
Rattus_norvegicus                   PEGADMATELFSPVDITQVIVNTSHSLPSQQYWLSTHLFSA-EQANLWCLSRCAQEPVFC      1903
Mus_musculus                        P--GDMATELFSPVDITQVIVNTSHSLPSQQYWLFTHLFSA-EQANLWCLSRCAQEPIFC      1901
Cavia_porcelus                      SAETGSTAELFSPVDLSQVIVSVNRSLPSQRHWLFKHRFSA-QQAKLWCLSRCSQEPSFC      1903
Canis_lupus_familiaris              PTETDLTTELFSPVDLDQVIVNGSQSLPSQQHWLFKHLFSP-QQANLWCLSRCVQEPSFC      1904
Panthera_leo                        PTEPDLTTELFFPVDLNQVVVNGSLSLPNQQHWLFKHLFSP-QQANLWCLSRCIQEPSFC      1906
Bos_taurus                          PSETDLTTGLFSPVDLIQVIVDGNVSLPSQQHWLFKHLFSL-QQANLWCLSRCAGEPSFC      1905
Trichechus_manatus_latirostris      PAETDLVAELFSPVDLNQIIVSGNRSLPSQQHWLFKHLFSP-QQANLWCLSRCVQEASFC      1906
Columbia_livia                      LTLSDSTKDLFYLMDNSQIQSDQNYSLPYQQYWVFRQKYSA-EEAVLWCLTRCTQEDEFC      1904
Taeniopygia_guttata                 LVLSDSTKDLFYLMDNSQIQSDQNYSLPYQQYWVFRQKYSD-EEALLWCLTRCAQEEEFC      1908
Larus_michahellis                   LTLSDSTKDLFYLMDNSQIQSDQNSSLPYQQYWVFRQKYSA-EEAVLWCLTRCAQEDEFC      1909
Gallus_gallus                       IMLSGSTKDLFYLMDDSQLRSDPNSSIPYQQYWVFRQKYSA-EEAMLWCLTRCAQEDEFC      1901
Struthio_Camelus                    LELADSTKDLFYLMDNGLIRSDQNSSLPYQQYWVFRQKYSA-EKAMLWCLTRCAQE-EFC      1882
Chelonia_mydas                      LMLPDSVMELFSLMDNSLIQLDQSRSLPSQQYWLFKHKYSA-EQATLWCLTRCAQEDAFC      1899
Alligator_mississippiensis          LMLSESAGELFSVMDNNLIQIDRSRSLPAQEYWLFKHKFSL-KQATRWCLTRCTQEGAFC      1883
Crotalus_tigris                     SRLSASTKKLFSIVERNQISTELNRPLPSQKYWLFKQRYSA-EEALLWCLTRCV-EDEFC      1896
Python_bivittatus                   SLLSASTKELFSIVENNQILTEPNRSLPSQKYWLFKHRYSA-EEVLVWCLARCV-EDEFC      1838
Eublepharis_macularius              SELTASALELFSLMESSQIQIDQHRSLPSQEYWIFRHKYSA-EQARLWCLTRCA-EDEFC      1888
Xenopus_tropicalis                  SSISDSVLDLFLPVESNTVTTDPHLIISSQQYGVSRLRYSS-DQALRWCLTRCKEEGSWC      1926
Aquarana_Catesbeiana                ANITESVRDLFLPVNDNHVVINQDIDLPHTEYWISKDRYTP-EQAQNWCLSRCKEEESWC      1905
Carassius_auratus                   STLSDSIKEIFGVLEDSDVQVDPERELPSQLYWIFKHQYTF-QEAQLWCLKRCG-EEQLC      1868
Cyprinus_carpio                     IILSDSIKETFGVLENGDVKVDPERELPSQLYWIFKHQYTF-QEAQLWCLKRCG-EEQLC      1873
Danio_rerio                         SALSDSVKEAFDVLDSGDVNVDPERELPNQLYWIFKHQYSF-QEAQLWCLKRCE-EEELC      1873
Astyanax_mexicanus                  STVSDSVKEMFDILSGGDISINSDKKTPTQQYWLFKHQFTA-EEARLWCLKRCM-EEELC      1869
Clupea_harengus                     ELLSDAVLRSFGSLEASEVQVDPKKAIPSQQYWIFKHQFTP-EQAKLWCLKRCD-EEELC      1894
Oryzias_latipes                     APLDESLQQKFEALSKDDVVVDPQRKLPVLSFWLNKNNYNS-QQALLWCLTRCY-EEQQC      1548
Xiphophorus_maculatus               APLNASVQLRFQSLSSADVLVDPQRKQPVLSFWLSKNDYSS-QNALLWCLTRCD-EEERC      1550
Stegastes_partitus                  PPLDDSVQLKFESLSADDVLVDPQRKLPVLSFLLNKNNYNS-QQALLWCLTRCD-AEPQC      1565
Cynoglossus_semilaevis              SPLDGTVHKKFRSVPEDDVVVDPHKKLSALRFLLNKKSFNS-QHALLWCLTRCD-AESLC      1535
Lepisosteus_oculatus                ILLSDTVQEAFTPVESSNVLVNPTQDIGHQQYWIFKHEFSA-QQAQLWCLKRCE-EEEFC      1891
Lampetra_fluviatilis                ASTAPDVSGRFSTLRPEEVTLQPEMRVAQQNFWLFRRAFSP-QQATSWCLRRCR-GDALC      1991
Lampetra_planeri                    ASAAPDVSGRFSTLRPEEVTLQPEMRVAQQNFWLFRRAFSP-QQATSWCLRRCR-GDALC      1928
Petromyzon_marinus                  PAGAADVSGRFSTLRPEEVTLQPEMRVAQQNFWLFRRAFSPPQQATSWCLRRCR-RDALC      1936

Homo_sapiens                        QLAEITESA-SLYFTCTLYPEAQVCDDIMESNA-----QGCRLILPQMPKALFRKKVILE      1958
Pan_troglodytes                     QLAEITESA-SLYFTCTLYPEAQVCDDIMESNA-----QGCRLILPQMPKALFRKKVILE      1957
Gorilla_gorilla                     QLAEITESA-SLYFTCTLYPEAQVCDDIMESNA-----QGCRLVLPQMPKALFRKKVILE      1945
Pongo_pygmaeus                      QLAEITESA-SLYFTCTLYPEAQVCDDIMESNA-----QGCRLILPQMPKALFRKKVILE      1959
Macaca_mulatta                      QFAEITESA-SLYFTCTLYPEAQVCDDILESNA-----QGCRLILPQRPKALFQKKVILE      1959
Macaca_fascicularis                 QFAEITESA-SLYFTCTLYPEAQVCDDILESNA-----QGCRLILPQRPKALFQKKVILE      1959
Rattus_norvegicus                   QLADIMESS-SLYFTCSLYPEAQVCDNDVESNA-----KNCSQILPRQPTALFQRKVVLN      1957
Mus_musculus                        QLADITKSS-SLYFTCFLYPEAQVCDNVMESNA-----KNCSQILPHQPTALFRRKVVLN      1955
Cavia_porcelus                      QLAEVTDSA-PSYFTCSLYPDARVCDDVLEPNA-----QGCSLVLPQQPGTLLQKKVVLN      1957
Canis_lupus_familiaris              QLVEITDSA-PLYFTCALYPEARVCDSVMESSP-----KGCRLILPHRPETLFRKKVILR      1958
Panthera_leo                        QLVEITDST-PLYFTCTLYPEAQVCDDVMEPNP-----KGCRLFLPHQPKTLFRKKVILR      1960
Bos_taurus                          QLAEVTDSE-PLYFTCTLYPEAQVCDDILESSP-----KGCRLILPRRPSALYRKKVVLQ      1959
Trichechus_manatus_latirostris      QLAEIIDST-PLYFTCTLYPEAQVCDDVMETSP-----KGCRVILPHRPKMLFQKKVILK      1960
Columbia_livia                      RMADIQSTA-DLYFVCTLYPEAQICDGSIDQIP-----ENCRTVLTWQPQTLYRKIVTLK      1958
Taeniopygia_guttata                 RMADLQSAT-DKYFVCTLYPEAQICDSNISQIP-----GNCATVLPWQPQTLYHKIVTLK      1962
Larus_michahellis                   RMADLQNAT-DVYFVCTLYPEAQICDGSINQMP-----ENCRTVLPRQPQTLYRKIVTLK      1963
Gallus_gallus                       QMADLRNTA-DTYSVCTLYPEAQICDGNIEQIL-----ENCHTVLPWQPQALYHKIVTLR      1955
Struthio_Camelus                    QIADLQNAT-DTYFVCTLYPEAQICDGNIDQIP-----ANCLTVLPWQPQMAYHKIVTLK      1936
Chelonia_mydas                      QLAALRNTT-STYFACILYPEAQICDNYINHIP-----QNCQTVLPQQLQTVYHKIVTLE      1953
Alligator_mississippiensis          QLVDLQNTT-GTYFVCTLYPEAQVCDNSINKIP-----DSCQTILPREPQIVHYKRVTLG      1937
Crotalus_tigris                     LLADIPNNTDTAFFPCTLYPAAQVCNTSINNIP-----NNCKIVLPQNPQLLYQKTVPLE      1951
Python_bivittatus                   LLADVPNNTDTAFFPCTLYPAAQVCNTFINNIP-----NNCKIVLPQKPRLLYQKIVSLE      1893
Eublepharis_macularius              HLADIPNVTTQTYFPCTLYPVAQVCDN-INNVP-----ENCRTVLPQKPNILYQKKVSLA      1942
Xenopus_tropicalis                  LLTDLQETT-EEFFTCTLYPDAWKCSNSPSLTP-----DDCKITLHNKPQRLYLKKDMLG      1980
Aquarana_Catesbeiana                RLVVLQDSM-KQFFTCIIYPDTWNCNNYTDLVP-----ATCDIVLNHKPQSLYRRKETLS      1959
Carassius_auratus                   HVSDIRDEG-PLYFACALYPDTRVCGAYDKPLR-----QACSLVMTQRPQNAYQKNVSLL      1922
Cyprinus_carpio                     HVSEIRDEG-PLYFACALYPDTRVCGAYDKPLR-----QACSLVMTKSLQTAYQKKVSLL      1927
Danio_rerio                         HVSDIRDEG-PLYFACVLYPDTRVCGAYDKPLR-----QTCSLVMTQSLQTAYQKKVSLT      1927
Astyanax_mexicanus                  HVADLRDEG-SVYSVCMLYPDTRVCGAYDKPLR-----QACSLVLPQDPQTAYRKKVDLS      1923
Clupea_harengus                     HVSDLRDEG-SVYFLCELYPDTRVCGAYDKPLR-----QACSLQLPRDPQTAHRKRVDLT      1948
Oryzias_latipes                     AVADLADADLAGFFSCSLYPNTRVCGAYDMVPM-----VPCPPLMDRLPQNAYSKKVELS      1603
Xiphophorus_maculatus               SVADLRDAESHRFFLCSLYPDTRVCGGYDRPIK-----RSCRPLLDRATNNTYSKKVDLN      1605
Stegastes_partitus                  SVADLSDA--AGFFSCSLFPDSRVCGAYDKPLR-----RPCRPLLDRTPNNTYSKKGNLS      1618
Cynoglossus_semilaevis              AVADLTEAETAGFYSCSLFPDTRVCGAYDKPLR-----AACRLLLNRAPNNTHIKRVELS      1590
Lepisosteus_oculatus                KVTDLQDND-TLHFTCTLYPDTQICGSYNKPIQ-----RSCTLVLPVEPQSVHRKKIELT      1945
Lampetra_fluviatilis                KAAAVGDGP-AGWLECLLYPDTQSCGTSQQLLDWPARAPSCASLLPRLDGTLYRKRDGAE      2050
Lampetra_planeri                    KAATVGDGP-AGWLECLLYPDTQSCGTSQQLLDWPARAPSCASLLPRLDGTLYRKRDGAE      1987
Petromyzon_marinus                  KAAAVGDGP-AGWLECLLYPDTQSCGTSQELLERPARAPSCASLLPRIDGTLYRKRDGAE      1995
```

TG type 3b-2

```
Homo_sapiens                          DKVKNFYTRLPFQKLMGISIRNKVPMSE-KSISNGFFECERRCDADPCCTGFGFLNVSQL    2017
Pan_troglodytes                       DKVKNFYTRLPFQKLMGISIRNKVPMSE-KSISNGFFECERRCDADPCCTGFGFLNVSQL    2016
Gorilla_gorilla                       DKVKNFYTRLPFQKLMGISIRNKVSMSE-KSISNGFFECERRCDADPCCTGFGFLNVSQL    2004
Pongo_pygmaeus                        DKVKNFYTRLPFQKLLGISIRNKVPMSE-KSISNGFFECERQCDADPCCTGFGFLNVSQL    2018
Macaca_mulatta                        DKVKNFYTRLPFQKLMGISIRNKVPMSE-KSISNGFFECERRCDADPCCTGFGFLNVSQL    2018
Macaca_fascicularis                   DKVKNFYTRLPFQKLMGISIRNKVPMSE-KSISNGFFECERRCDADPCCTGFGFLNVSQL    2018
Rattus_norvegicus                     DRVKNFYTRLPFQKLSGISIRDRIPMSE-KLISNGFFECERLCDRDPCCTGFGFLNVSQM    2016
Mus_musculus                          DRVKNFYTRLPFQKLTGISIRDKVPMSG-KLISNGFFECERLCDRDPCCTGFGFLNVSQL    2014
Cavia_porcelus                        SRVKNFYTRLPFQKLAGISIRNKVPMSE-KPISNGFFECERLCDLDLCCTGFGFLNVSQS    2016
Canis_lupus_familiaris                DKVKNFYTRLPFQKLMGISIRNKMPMSE-KSISNGFFECERLCDADPCCTGFGFLNVSQL    2017
Panthera_leo                          DKVKNFYTRLPFQKLTGISIRNKVPMSE-KSISNGFFECERLCDADPCCTGFGFLNVSQL    2019
Bos_taurus                            DRVKNFYNRLPFQKLTGISIRNKVPMSD-KSISSGFFECERLCDMDPCCTGFGFLNVSQL    2018
Trichechus_manatus_latirostris        DKVKNFYTRLPFQKLTGISIRSKVPMSE-KSISNGFFECERRCDADPCCTGFGFLNVSQS    2019
Columbia_livia                        SSVKSFYTRVPFQKVTAISVRNKTDMSR-KTVSDGFFECERWCDADPCCTGFSFFNDSQI    2017
Taeniopygia_guttata                   SSVKSFYTRVPFQKVTEISVRNKTDMSR-KAVSDGFFECERWCDADPCCTGFGFFNDSQL    2021
Larus_michahellis                     SSVKSFYTRIPFQTVTGISVRNKTDMSR-KTVSDGFFECERWCDADPCCTGFGFFNDSQL    2022
Gallus_gallus                         STVKSFYTRVPFRKATGISVRNVTDMSR-KAVSDGFFECERWCDADPCCTGFGFLNVTQL    2014
Struthio_Camelus                      SSVKSFYTRVPFQKVTGISVRNKTDMSR-KAVSDGFFECERWCDADPCCTGFGFFNDSQL    1995
Chelonia_mydas                        GTVKSFYTRLPFRKVTGISVRNKIDMSG-KAVSDGFFECERRCDADPCCTGFGLLNESQS    2012
Alligator_mississippiensis            ATVKNFYTRLPFRKVTGIAVRNKIDLSG-KAISDGFFECERWCDADPCCMGFGLLNGLQS    1996
Crotalus_tigris                       GSVKNFYTRLPFRKVSGVSVRNKMDMSG-KTVSNGFFDCERLCDADLCCSGFGLLNASQD    2010
Python_bivittatus                     GSVKNFYTRLPFRKMSGVSVRNKMDMSG-KTVSNGFFDCERLCDADPCCSGFGLLNASQG    1952
Eublepharis_macularius                GSVKNFYTRLPFGKVSGISVRNKTDVSG-KTVGDGFFDCERLCDSDPCCTGFGLLNASQA    2001
Xenopus_tropicalis                    NTVKNFYSVLPFRQITDVSVRRKIDMTG-KSISNGFFECELHCDADPCCKGFGYLQTTHG    2039
Aquarana_Catesbeiana                  NKVKNFYVPVPYRKLSEITVRNRITVTG-ESVSKGFFLCELHCDADPCCRGFGYIQRPEI    2018
Carassius_auratus                     GSVKSFYGRVPFKKMVSYSVRSRVSLSS-KSITEGFFECERRCDEDPCCRGIGYVRDSGV    1981
Cyprinus_carpio                       GSVKSFYSRVPFKKMVSYSVRSRVSLSS-KSIPEGFFECERRCDEDPCCRGIGYVQDSGV    1986
Danio_rerio                           GSVKSFYSRVPFKKMVSYSVRSRVSVSS-KPITEGFFECERRCDEDPCCRGIGYVKDSGV    1986
Astyanax_mexicanus                    GSVESFYNRVPFKKMVSYSVRSKVSVSS-KPITEGFLECERRCDEDPCCRGIGYIRDTGT    1982
Clupea_harengus                       GSVESFYTRVPFKKMVSYSVRSRVNLSA-KPITEGFLECERRCDEDPCCRGMGYVKDTAA    2007
Oryzias_latipes                       GPVKNFYQRISFQKSDSNGVRSRAVMSENTPLSQGFRDCERRCDEDPCCRGFGFVGYNKS    1663
Xiphophorus_maculatus                 GPVQSFYERVPFQKMVSYSVRNRATLGQNQLLPDGFRECERRCDEDPCCRGFGFVRETKS    1665
Stegastes_partitus                    GPVKSFYEKVSFQKMVSYSVRSRVSLKQNTPLPEGFMECERRCDEDPCCRGIGFVRDTKS    1678
Cynoglossus_semilaevis                GPVKSFYQRISFQKFDSYSVRSRVSLGENAPLTQGFRECERRCDEDPCCRGFGFVRDAKS    1650
Lepisosteus_oculatus                  GRVKNFYRRVPFTKLVAYSVRNRVNQTG-KPVTNGFNECELRCDEDPCCKGFGFVRDSQS    2004
Lampetra_fluviatilis                  GPVRRLYRRTAFRSEVGVLLRSVANLTG-MALFDGFSRCERTCDEDPCCEGFAFLRSSLS    2109
Lampetra_planeri                      GPVRRLYRRTAFRSEVGVLLRSVANLTG-MALFDGFSRCERTCDEDPCCEGFAFLRSSLS    2046
Petromyzon_marinus                    GPVRRLYRRTAFRSEIGVLLRSVANLTG-MALSDGFSRCERTCDEDPCCEGFAFLRRALS    2054

Homo_sapiens                          K-----------GGEVTCLTLNSLGIQMCSEENGGAWRILDCGSPDIEVHTYPFGWYQKP    2066
Pan_troglodytes                       K-----------GGKVTCLTLNSLGIQMCSEENGGAWRILDCGSPDIEVHTYPFGWYQKP    2065
Gorilla_gorilla                       K-----------GGEVTCLTLNSLGIQMCSEENGGAWRILDCGSPDIEVHTYPFGWYQKP    2053
Pongo_pygmaeus                        K-----------GGEVTCLTLSSLGIQMCSEENGGAWRILDCGSPDIEVHTYPFGWYQKP    2067
Macaca_mulatta                        K-----------GGEVTCLTLNSLGLQMCSEENGGAWRILDCGSPDIEVHTYPFGWYQKP    2067
Macaca_fascicularis                   K-----------GGEVTCLTLNSLGLQMCSEENGGAWRILDCGSPDIEVHTYPFGWYQKP    2067
Rattus_norvegicus                     Q-----------GGEMTCLTLNSMGIQTCSEENGATWRILDCGSEDTEVHTYPFGWYQKP    2065
Mus_musculus                          Q-----------GGEVTCLTLNSMGIQTCNEESGATWRILDCGSEDTEVHTYPFGWYQKP    2063
Cavia_porcelus                        K-----------GGELTCLTLSSMGVQTCSEESGGAWRILDCGSPDTEVRIYPFGWYQKP    2065
Canis_lupus_familiaris                T-----------GGEVTCLTLNSLGLQTCSEENMGAWRILDCGSPDTEVRTYPFGWYQKP    2066
Panthera_leo                          K-----------GGEVACLTLNSLGLQTCSEESGGTWRILDCGSPDTEVRTYPFGWYQKP    2068
Bos_taurus                            K-----------GGEVTCLTLNSLGLQTCSEEYGGVWRILDCGSPDTEVRTYPFGWYQKP    2067
Trichechus_manatus_latirostris        K-----------GGEVTCLTLSSLGLQTCSEENGGAWRILDCNSPDTEVRTYPFGWYQKP    2068
Columbia_livia                        P-----------GGKIVCLTLNSLGIQTCAEETRSAWQVSNCSSPDAEVRIHPFGWYQKP    2066
Taeniopygia_guttata                   S-----------GGKIVCVTLNSLGIQTCVEETRSSWHISDCSSPGAEVKIHPFGWYQKP    2070
Larus_michahellis                     S-----------GGKIVCLTLNSLGIQTCAEETRSVWQVSNCSLSDAEVRIHPFGWYQKP    2071
Gallus_gallus                         S-----------GGKIMCLTLNSLGIQTCSEES-GAWQISNCSLPDTEVRIHPFGWYQKP    2062
Struthio_Camelus                      S-----------GGKIMCLTLNSLGIQMCAEDTRSAWQVSDCGGPEAAVRIHPFGWYQKP    2044
Chelonia_mydas                        T-----------GGDVLCLTLNSLGIQTCTEETGSTWQISDCSSPD--VRIYPFGWYQKP    2059
Alligator_mississippiensis            T-----------GGKVLCLTLNGLGIQTCAEETRSAWQVSDCSSSDAEANTYPFGWYQKP    2045
Crotalus_tigris                       T-----------DSKILCLTLGSLGIQTCSDELRSDWQISPCTSLETEARVHPFGWYQKS    2059
Python_bivittatus                     T-----------GSKVLCLTLDSLGIQTCSNEFRSDWQIFNCTSLETEARVHPFGWYQKS    2001
Eublepharis_macularius                T-----------DGKVLCLIILNNLGIQTCSEELRNDWRVSDCAPLQAEARIHPFGWYQKG    2050
Xenopus_tropicalis                    P-----------GTETRCLLLSSMGIQSCMDTPGNQWRVLNCSSSIEGRGTHPFGWFQKP    2088
Aquarana_Catesbeiana                  Q-----------GLERICLPLASLGVQSCPEDTQSSLGISNCSSPDEGMKTSPFGWYQKP    2067
Carassius_auratus                     PSTQEPFHSDESRSDVLCLTLNSLGIQTCGENDDTNWRIQDCSPSKVQTEVYPFGWYEKP    2041
Cyprinus_carpio                       P-----------RSDVLCLTLNSLGIQTCGENDSTNWRIQDCSPSKVQTEVYPFGWYEKP    2035
Danio_rerio                           A-----------GSDVLCLTLNSLGIQTCGENDSTSWRIQDCSPSQVQSQLYPFGWYEKP    2035
Astyanax_mexicanus                    A-----------VADVSCLTLNSLGIQTCGEDDRTEWRVQDCSPSKVQTGLHPFGWYEKP    2031
Clupea_harengus                       AS---------SSGSELLCLTLNSLGIQTCGESDRTSWRVQDCSPSKVETQVYPLGWYEKP    2059
Oryzias_latipes                       PG-------------LVCLVLASLGVQTCSEKD-STWTTQDCSPTEAKTTPYPFGWYQKP    1709
Xiphophorus_maculatus                 RG-------------RLCLPLVSLGVLTCSEDDASTWTTQDCSPPEAETSPEPFGWYQKP    1712
Stegastes_partitus                    PG-----------GSDVVCLALISLGVQTCSESDSTTWKTQDCRPSAIKVSPEPFGWYQKP    1728
Cynoglossus_semilaevis                PG-----------GAKVVCVSLLSLGVLTCDEASESSWRRQDCAPPAVLSSPEPFGWFLKP    1700
Lepisosteus_oculatus                  P-----------GNEVLCLTLNSIGVQTCSDQEKTTWRVMDCSTSKAESNVLPFGFYKKP    2053
Lampetra_fluviatilis                  -----------REALLQCATHTGPGVQLCGRESP-VGATFGCPPPGVDTASSLFGWYRAQ    2157
Lampetra_planeri                      -----------REALLQCATHTGPGVQLCGRESP-VGATFGCPPPGVDTASSLFGWYRAQ    2094
Petromyzon_marinus                    SA---------GEALLQCATHTGPGVQLCGRESP-MGATFGCPPPGVDTASSLFGWYRAQ    2104
```

```
Homo_sapiens                      IAQ-NNAPSFCPLVVLPSLT--EKVSLDSWQSLALSSVVVDPSIRHFDVAHVSTAATS--   2121
Pan_troglodytes                   IAQ-NNAPSFCPLVVLPSLT--EKVSLDSWQSLALSSVVVDPSIRHFDVAHVSTAATS--   2120
Gorilla_gorilla                   IAQ-NNAPSFCPLVVLPSLT--EKVSLDSWQSLALSSVVVDPSIRHFDVAHVSTAATS--   2108
Pongo_pygmaeus                    IAQ-NNAPSFCPSVVLPSLT--EKVSLDSWQSLALSSVAVDPSIRHFDVAHVSTAATS--   2122
Macaca_mulatta                    IAQ-NSAPSFCPSVVLPSLT--EKVSLDSWQSLALSSVVVDPSIRHFDVAHVSTAATS--   2122
Macaca_fascicularis               IAQ-NSAPSFCPSVVLPSLT--EKVSLDSWQSLALSSVVVDPSIRHFDVAHVSTAATS--   2122
Rattus_norvegicus                 AVW-SDAPSFCPSAALQSLT-EEKVALDSWQTLALSSVIIDPSIKHFDVAHISISATR--   2121
Mus_musculus                      AVW-SDTPSFCPSAALQSLT-EEKVTSDSWQTLALSSVIVDPSIKHFDVAHISTAATS--   2119
Cavia_porcelus                    VAQ-SDVPSFCLPAVPPAFT--KKVALDSWQSLAVSSVAVDPSIRHFDVAHISTIA-T--   2119
Canis_lupus_familiaris            AAR-NDAPSFCPVVTPPSLP--EKVTLGSWQSLAPSAVVIDSSIRNFDVAHVSTATTN--   2121
Panthera_leo                      VAQ-NDAPSFCPLAVLPFLP--EKVALGSWQSLALSAAVVDPSIRNFDVAHVSTAATR--   2123
Bos_taurus                        VSP-SDAPSFCPSVALPALT--ENVALDSWQSLALSSVIVDPSIRNFDVAHISTAAVG--   2122
Trichechus_manatus_latirostris    DSQ-NDVPSFCPSVILPPLT--EKVPPDAWQSLTISSVIVDPSIRHFDVAHISTAAAS--   2123
Columbia_livia                    ADLRNTMPNLCPTVSLPLGP--ESDYLDKWQPLNVSSVLLDSSISNFEVVQVSRDISN--   2122
Taeniopygia_guttata               ADLKNTIPNLCPPVNVPFRP--ESEYMDAWQSLNVSSVLMDSSISHFEVVHVSRDISN--   2126
Larus_michahellis                 ADLKNTMPNLCPPVSLPLRP--EDEYSDIWQPLNMSFVLMDSSISNFEVVQVSRDISN--   2127
Gallus_gallus                     ADLTNTVPNLCPPVSLPFRP--ESEYLDTWQPLNVSSVLVDSTISNFEAVHVSREISN--   2118
Struthio_Camelus                  ADLENTMANLCPPASLPLRP--ESEYLNIWQPLDVSSVLMDSSISNFEAVQVSRDIAN--   2100
Chelonia_mydas                    ATMTKTIPNACPPVLLPRSP--ENVPLDSWLLLDASSVLTDPSILNFDIAQVSREFSD--   2115
Alligator_mississippiensis        ANLKRTIPSVCPPVLLPLQP--ETVSLDMWQLLDASSVLVDSSLVNFDVVQVSRDNSG--   2101
Crotalus_tigris                   ----SIVPRVCPPAPSLEKQ--ANVALDKWQRLDRTSTILDPSISKFDVIQVSRDASS--   2111
Python_bivittatus                 ----SIVPRVCPPVPLPEKQ--GKVTLDKWQRLDRSSILLDPSISKFDVIHVSRDPSS--   2053
Eublepharis_macularius            ADRMANVPSVCPAVHLPGTE--ENVSLDKWQSLDSSSFLVDSSISQFDIVHLSIEAAN--   2106
Xenopus_tropicalis                DNQRSPPSSLCPPVDVLKIK--Q-EDLGDWLLLDKSSAVIDPSLSAYDMVQISRESPD--   2143
Aquarana_Catesbeiana              DGEGSILPGLCPTADVLKTT--KPVLLDDWITLDPSAVTIDPSLSNYDEVLISRGASN--   2123
Carassius_auratus                 VNQWIKNPSVCPPFSLRSPS--KNADLKNWKQLDASSVKVDSSVSAFDIVHISRDIAD--   2097
Cyprinus_carpio                   VNQWIKNPNVCPPFSLRSPS--KNSDLKNWKQLDASSVKVDSSVSAFDIVHISRDIAE--   2091
Danio_rerio                       VNQWPNNPDVCPPFSLRSPS--KNADLQKWKKLDVASVYVDASVSAFDIVHISKDIAE--   2091
Astyanax_mexicanus                VNQWTKSPGICPSFKLRAPS--KNVNLKDWTLLNASSILVDSSVSAFDIVHLSKDIAE--   2087
Clupea_harengus                   VNQWTKSPRLCPTFGLRSPS--KNVTMLDWKLQDTSVVLVDPSLSMFDIVHISKDIAG--   2115
Oryzias_latipes                   VNQWSSSPALCPPLSLPETR--NNVSLDDWHLLPDSLVLIDPSLSTYDVIHVSRDIAT--   1765
Xiphophorus_maculatus             VNQWTSSPALCPPFSLRDRR--NNVSMDDWRLLSESSLLIDASLSTYDVIHVSRDIDT--   1768
Stegastes_partitus                VNQWSSSPALCPPFNLPPTK--SNVSMEEWRLLSESSITIDPSLSTYDVIHVSRDIAT--   1784
Cynoglossus_semilaevis            VQQWTSAPALCPEFGLT--A--PNVSLSKWRLLDESSVLVDPSVSTYDVIHISSDISG--   1754
Lepisosteus_oculatus              VNQWNHNPKACPAFHLPSPYQSQNAVLSAWEFLDFSSVAIDPSISEYDVIHLSKDITE--   2111
Lampetra_fluviatilis              ARSSPQAAQLCPAVSLPPAP--TKGAADGFHCLDVSTAAVDPTISAFDVVVLLGPGAGGS   2215
Lampetra_planeri                  ARSSPQAAQLCPAVSLPPAP--TKGAADGFHCLDVSTAAVDPTISAFDVVVLLGPGAGGS   2152
Petromyzon_marinus                ARSSPQTAQLCPAVTLPPAP--TKGAADGFQSLDVSTAAVDPTISAFDVVVLLGPGAGGS   2162
```

TG type 3a-3

```
Homo_sapiens                      --NFSAVRDLCLSECSQHEACLITTLQTQPGAVRCMFYADTQSCTHSLQ------GQNCR   2173
Pan_troglodytes                   --NFSAVRDLCLSECSQHEACLITTLQTQPGAVRCMFYADTQSCTHNLQ------GQNCR   2172
Gorilla_gorilla                   --NFSAVRDLCLSECSQHEACLITTLQTQPGAVRCMFYADTQSCTHSLQ------GQDCR   2160
Pongo_pygmaeus                    --NFSAVRDLCLSECSQHEACLITTLQTQPGAVRCMFYADTQSCTHSLQ------GQNCR   2174
Macaca_mulatta                    --NFSAVRDLCLSECSQHEDCLITTLQTQPGAVRCMFYADTHSCTYSLQ------GQNCR   2174
Macaca_fascicularis               --NFSAVRDLCLSECSQHEDCLITTLQTQPGAVRCMFYADTHSCTYSLQ------GQNCR   2174
Rattus_norvegicus                 --NFSLAQDFCLQECSRHQDCLVTTLQIQQGVVRCVFYPDIQSCEHSLR------SKTCW   2173
Mus_musculus                      --NFSMAQDFCLQQCSRHQDCLVTTLQIQPGVVRCVFYPDIQNCIHSLR------SHTCW   2171
Cavia_porcelus                    --NFSAARDFCLQECSHHQACITTTLQTWPGATRCVFYPDTQSCTHTLQ------GQSCQ   2171
Canis_lupus_familiaris            --DFSDARDFCLLECSRHQACLVTTLRAQPGAVRCVFYADAQICTHSLQ------AQNCQ   2173
Panthera_leo                      --DFSDARDFCLLECSRHQACLITTLQPRPGAVRCVFYADTQICTHSLQ------AQNCR   2175
Bos_taurus                        --NFSAARDRCLWECSRHQDCLVTTLQTQPGAVRCMFYADTQSCTHSLQ------AQNCR   2174
Trichechus_manatus_latirostris    --DFSAARDLCLLECSRHQACIVTALQTRPGAVRCMFYADTQSCTHSLQ------SQSCR   2175
Columbia_livia                    --DFSTAREICLSACSKNQSCTVVTLEIQPSAIRCLFYPDTQMCTHGLQ------RHSCW   2174
Taeniopygia_guttata               --DFSLARDFCLSACSKNQSCAVVTLEIQPSAIRCIFYPDTQMCAHDLQ------GHSCW   2178
Larus_michahellis                 --DFSTARDFCLSACSKNQSCTVVTLEIQPSAIRCLFYPDTQICKHGLQ------GHSCW   2179
Gallus_gallus                     --GFANAKDYCLSACSRNQSCAVVTLEIQASAIRCLLYPDTQICSHGLE------GHSCK   2170
Struthio_Camelus                  --DFSTARDFCLSACSKNQSCTVVTLEIQPSAIRCLFYPDTQICTHSLQ------GHSCR   2152
Chelonia_mydas                    --DFSTARDFCLSACSKNQSCTVITLEIQPSAIRCIFYPDTQICTHGLQ------GHSCR   2167
Alligator_mississippiensis        --NFTAARDICLSACSKSQSCIVVTLKIQLSTIRCIFYPDTKICTHGLQ------GHSCR   2153
Crotalus_tigris                   --EFAAIRDFCLSVCSKDNLCLVTTLEMLPSAARCMFYPETQSCTLSLQ------GHHCR   2163
Python_bivittatus                 --EFATIRDFCLSVCSKDNLCLVTTLEILSSAIRCIFYPETQSCTLSLQ------GHHCR   2105
Eublepharis_macularius            --DFASARDFCLSVCSKNKFCLVTTLEIRPSATRCMFYPETQSCRLSLQ------GHLCQ   2158
Xenopus_tropicalis                --NLQAAQNYCLAVCARAPSCVTATVALQEAAVRCLFYPDNQNCFYGLR------NHQCQ   2195
Aquarana_Catesbeiana              --QLSTVQSYCLSECARVPTCITTTINVLQAAIKCLFYPETQNCYYSLK------GHRCQ   2175
Carassius_auratus                 --DLDKVRDWCLSACEESESCSAVSVDSRESAMRCVMYPDTHTCLSTTS------GQRCF   2149
Cyprinus_carpio                   --DLDKVRDWCLSACEESESCSAVSVDSRESAMRCVMYPDTHTCLPTTS------GQRCF   2143
Danio_rerio                       --DLDKVKNWCLSACEDSESCSAVSIDSRESAVRCVMYPDTHTCLPTTS------GRRCL   2143
Astyanax_mexicanus                --DLDRVRDWCLAACEENDSCYAVSVDSKESAVRCVMYPDSHICLPTSR------GQHCV   2139
Clupea_harengus                   --DAERVRDWCLAACEGSVSCVAVSLESRESASRCVLYPDTHSCIPSED------TQDCR   2167
Oryzias_latipes                   --DKDRTRDWCLYACREAESCAAVSLSEVESATRCILYPDTTVCGLSSS----STTSSCQ   1819
Xiphophorus_maculatus             --DRERTRDWCLHACQEAESCVAVSLREAESAVRCILYPDTTTCGLSSAPDSISPASSCR   1826
Stegastes_partitus                --DGDKTRDWCLHACQEAESCAAVSLREEESAIRCILYPDTTTCGLSSAPGSSNPSYSCR   1842
Cynoglossus_semilaevis            --DQDKTRNWCLHACQEAESCVAVSLSEGQSGVRCVLYPDTTTCGLSSSPHSSQSTSSCR   1812
Lepisosteus_oculatus              --DFNKSRDWCLSACTENASCATVTVDITDSAVRCVFYPDTHRCSPSTL------GQECK   2163
Lampetra_fluviatilis              PQAAPPSDEWCLAACQQSPWCASVGVRHLGERTRCVLHAETLACSGHGA----RGGRHCH   2271
Lampetra_planeri                  PQAAPPSDEWCLAACQQSPWCASVGVRHLGERTRCVLHAETLACSGHGA----RGGRHCH   2208
Petromyzon_marinus                PRAAPPSDEWCLAVCQRSPWCASVGVRHLGDSTRCVLHAETLAC-GYGA----RGGRHCR   2217
```

Spacer 3 ChEL

```
Homo_sapiens                          LLLREEATHIYRKPGISLLS-YEASVPSVPISTHGRLLGRSQAIQV-GTSWKQVDQFLGV   2231
Pan_troglodytes                       LLLREEATHIYRKPGISLLS-YEASVHSVPISTHGRLLGRSQAIQV-GTSWKQVDQFLGV   2230
Gorilla_gorilla                       LLLREEATHIYRKPGISLLS-YEASVPSVPISTHGRLLGRSQAIQV-GTSWKQVDQFLGV   2218
Pongo_pygmaeus                        LLLREEATHIYRKPGISLLS-YEASVPSVPISTHGRLLGRSQAIHV-GTSWKRVDQFLGV   2232
Macaca_mulatta                        LLLREEATHIYRKPGISLLS-YEASVPSVLIVTHGRLLGRSQAIQV-GTSWKQVDQFLGV   2232
Macaca_fascicularis                   LLLREEATHIYRKPGISLLS-YEASVPSVLIVTHGRLLGRSQAIQV-GTSWKQVDQFLGV   2232
Rattus_norvegicus                     LLLHEEAAYIYRKSGAPLHQSDGISTPSVHIDSFGQLQGGSQVVKV-GTAWKQVYQFLGV   2232
Mus_musculus                          LLLHEEATYIYRKSGIPLVQSDVTSTPSVRIDSFGQLQGGSQVIKV-GTAWKQVYRFLGV   2230
Cavia_porcelus                        LLLREEAIHIYRKPGTTLLRSEGGSVPSVLIAPHGWLQGRSQAVQV-GTAWKQVDQFLGV   2230
Canis_lupus_familiaris                LLLREEATHIYRKLNIPLLS-FGTSVPSVTITPHGQLLGRSQAIQV-GTSWKQVDQFLGV   2231
Panthera_leo                          LLLREEATHIYRKLSNPLLS-SGTPAPSVTITPHGQLLGRSRAVQV-GTSWKQVDQFLGV   2233
Bos_taurus                            LLLHEEATYIYRKPNIPLPG-FGTSSPSVPIATHGQLLGRSQAIQV-GTSWKPVDQFLGV   2232
Trichechus_manatus_latirostris        LLLREEASQVYRKLGISPLS-FNASAPSVPIATHGRLLGRSQAIQV-GTAWKQVDQFLGV   2233
Columbia_livia                        VLLKEPATYVYRRQDLFLPISESDLTPSVYIPSHGDLMGKSQVIRV-GSEWRNISQFLGI   2233
Taeniopygia_guttata                   ILLKEPATYIYRRQDLLLPISESDSTPRVYIPSHGYLMGKSQVIHV-GSGWRNISQFLGI   2237
Larus_michahellis                     VLLKEPATYIYRRQDLFLPISESDLTPSVYIPSHGDLVGKSQVIRI-GSEWRNISQFLGI   2238
Gallus_gallus                         VLLKEPATYIYRRQDSILPTSESDLAPSVYIQSHGDLIGKSKAVRI-GSEWKNISQFLGI   2229
Struthio_Camelus                      ILLKESATYIYRRQDLFLPTSASDLTSSVYVPSHGDLIGKSLVIHI-GSEWRNISQFLGI   2211
Chelonia_mydas                        VLLKEPATYIYWRQDLFLPISAPDLTSSVYIPSHGVLIGRSQVIRV-GSEWRNVSQFFGI   2226
Alligator_mississippiensis            ILLKEPATYIYRRQDLFLPISEPGVT-SVNIPSQGVLIGRSRAIRI-GAEWRSVSWFLGI   2211
Crotalus_tigris                       ILLKEPAAYIYRKQDVFLPTAERNEDLSVFIPSHGTLLGISQVIQV-GSEWKSIGQFLGV   2222
Python_bivittatus                     ILLKEPAAYIYRKQDAFLPTAERNSDPSVHIPSQGTLLGISQVIQV-GSKWKSIGQFLGI   2164
Eublepharis_macularius                LLLKEPATYIYRKKDAVSSITENNGTVAVSISSQGVLYGASKVIQV-GSNWKSIRQFLGI   2217
Xenopus_tropicalis                    LLVKEPATFIFRRKVRPPPST------SVAI-PQGTLLGKSEAVVI-GSNIKNVVQFLGI   2247
Aquarana_Catesbeiana                  LLVRESATYTFRKRATARPLT------SVAI-PLGIIEGKSQAVLI-GSDVRTVNQFLGV   2227
Carassius_auratus                     LVTKEPARAVYVRTG--LR----PELTSVSIPDHGTLLGESEVKPITGSDSKRVTYFLGV   2203
Cyprinus_carpio                       LVTKEPAQAVYVRTG--LQ----PELTSVSIPDHGTLLGESEVKPITGSDSKRVTYFLGV   2197
Danio_rerio                           LVTKEPAQSVYIRTG--VQ----LEFTSVSIPDHGTLLGESEVKPITGSDSKRVTYFLGV   2197
Astyanax_mexicanus                    LLTKEPAQYVYLKTV--LK----PELTTVSVPGHGALLGESEVKAF-GVDSKPVTYFLGV   2192
Clupea_harengus                       LRIREPSSQVYLRTVSALR----QDVPSVVIPGHGQLLGQSAVTAV-GSNRKNVWQFLGI   2222
Oryzias_latipes                       LVIREPASQVYLRTERL------PSVTSVSIPGQGVLEGVAVETTV-GSDRRRVIQFLGV   1872
Xiphophorus_maculatus                 LVVKEPAPQVYLRTERS------PQAASVSVPGHGTLQGVAMETAV-GSDRRTVIRFLGV   1879
Stegastes_partitus                    TVLREPASQVYLRAERM------PSPTSVSIPGHGILQGVAVETVL-GSDRRTVVQFLGV   1895
Cynoglossus_semilaevis                LLLREPADTVYLRADRP------PVVTTVSIPGNGLLQGSTVETWL-GSEQRTVVQFLGV   1865
Lepisosteus_oculatus                  LLLREQGLLVLHKKN--FK----PRLTSVFIPDHGMVLGESQLKLL-GSDRKEVNHFLGV   2216
Lampetra_fluviatilis                  LGVLEPPARLYRKK----GGGQGLVRPTVPL-GSGQLQGASRTVEV-DSQVKVVDVFLGV   2325
Lampetra_planeri                      LGVLEPPARLYRKK----GGGQGLVRPTVPL-GSGQLQGASRTVEV-DSQVKVVDVFLGV   2262
Petromyzon_marinus                    LGVLEPPARLYRKKAPSPGGGQGPVRPVVPL-GSGQLQGASRTVEV-DSQVKVIDVFLGV   2275
```

```
Homo_sapiens                          PYAAPPLAERRFQAPE---PLNWTGSWDAS--KPRASCWQPGTRTSTSPGVSEDCLYLNV   2286
Pan_troglodytes                       PYAAPPLAERRFRAPE---PLNWTGSWDAS--KPRASCWQPGTRTSTSPGVSEDCLYLNV   2285
Gorilla_gorilla                       PYAAPPLAERRFQEPE---PLNWTGSWDAS--KPRASCWQPGTRTSTSPGVSEDCLYLNV   2273
Pongo_pygmaeus                        PYAAPPLAERRFQAPE---PLNWTGSWDAS--KPRASCWQPGTRTSTTPGVSEDCLYLNV   2287
Macaca_mulatta                        PYAAPPLAERRFRAPE---PLNWTGSWDAS--QPRASCWQPGTRTSTSPGVSEDCLYLNV   2287
Macaca_fascicularis                   PYAAPPLAERRFRAPE---PLNWTGSWDAS--KPRASCWQPGTRTSTSPGVSEDCLYLNV   2287
Rattus_norvegicus                     PYAAPPLAENRFQAPE---VLNWTGSWDAT--KLRSSCWQPGTRTPTPPQISEDCLYLNV   2287
Mus_musculus                          PYAAPPLADNRFRAPE---VLNWTGSWDAT--KPRASCWQPGTRTPTPPQINEDCLYLNV   2285
Cavia_porcelus                        PYASPPLAERRFRPPE---PLNWTGSWDAT--KPRASCWLPGVHTPTPPGVSEDCLYLNV   2285
Canis_lupus_familiaris                PYATPPLAESRFRAPE---PLNWTGSWDAT--KPRASCWQPGTRTLESPGVDEDCLYLNV   2286
Panthera_leo                          PYASPPLAESRFRAPE---PLNWTGPWDAT--QPRASCWQPGTRTPTSPGVSEDCLYLNV   2288
Bos_taurus                            PYAAPPLGEKRFRAPE---HLNWTGSWEAT--KPRARCWQPGIRTPTPPGVSEDCLYLNV   2287
Trichechus_manatus_latirostris        PYAAPPLAESRFRAPE---PLNWTGSWDAT--KPRASCWQPGARTLKPPGVSEDCLYLNV   2288
Columbia_livia                        PYAAPPLAERRFSPPE---PFAWVETWNAT--AARAACWQPGDGEAPPYSVSEDCLYLNI   2288
Taeniopygia_guttata                   PYAAPPLGERRFRPPE---PFAWLEAWNAT--AARAACWQPGDGEAPSQSVSEDCLYLHI   2292
Larus_michahellis                     PYAAPPLAERRFSPPE---PFAWVETWNAT--VARAACWQPGDGEAPPYSVSEDCLYLNV   2293
Gallus_gallus                         PYAAPPLAERRFNPPE---PFAWEKTWNAT--VARSTCWQPGDRETPSYSVSEDCLYLNV   2284
Struthio_Camelus                      PYAAPPLAERRFSPPE---PFAWVETWNAT--EARATCWQPGDGDPPSYSVSEDCLYLSV   2266
Chelonia_mydas                        PYAVPPIAENRFHPPE---PFTWLQSWNAT--MVRASCWQPGDGLHQSSPVSEDCLYLNV   2281
Alligator_mississippiensis            PYAAPPVAENRFRPPA---PFTWLESWNAT--MARAACWQPGDGAVQASTVSEDCLYLNV   2266
Crotalus_tigris                       PYAAPPVGENRFHPPQ---PFTWTDFWNAT--TIRANCWQPGDD--SFSSVSEDCLFLNI   2275
Python_bivittatus                     PYAAPPVGENRFHPPQ---PFTWTDIWNAT--TTRASCWQPGDDT-SLSSVSEDCLFLNI   2218
Eublepharis_macularius                PYAAPPLAENRFRPPQ---VFTWVESWNAT--TTRASCWQPGDDEVSPSSVSEDCLYLNI   2272
Xenopus_tropicalis                    PYAAPPVGVYRFSPPQ---PVNWTGEWNAT--YSRASCLQPGDGKAQYSSVGEDCLYLNV   2302
Aquarana_Catesbeiana                  PYAAPPTGENRFRPPQ---PYTWTGTWNAT--FVRSSCLQPGDGKAQYSSVSEDCLYLNV   2282
Carassius_auratus                     PYARPPVGELRFSPPL---PADWTGTWNAT--FSRSSCLQPGDP--GDSTSSEDCLYLNV   2256
Cyprinus_carpio                       PYARPPVGELRFSPPQ---PADWTGTWNAT--FSRSSCLQPGDP--IDSTSSEDCLYLNV   2250
Danio_rerio                           PYARPPIGDLRFSPPQ---PADWTGTWNAT--FFRSSCLQPGDL--TDSSSSEDCLYLNV   2250
Astyanax_mexicanus                    PYARPPVGDLRFSPPQ---PADWTGTWNAT--FARPSCLQPGDT--SDSRSSEDCLYLNI   2245
Clupea_harengus                       PYAQPPIKTLRFSPPQ---PADWTGSWDAT--VTRPSCIQPGVN----SGTSEDCLYLNV   2273
Oryzias_latipes                       PYARPPIGALRFEAAQ---PADWTGTWDAS--KPRPSCLQPGDG--ENSASSEDCLYLNV   1925
Xiphophorus_maculatus                 PFARPPIGALRFEAAQ---PADWTGTWDAT--KPRPSCVQPGDD--EDSASSEDCLYLNI   1932
Stegastes_partitus                    PYARPPIGSLRFEAAQ---TADWTGTWDAT--KPRPSCVQPGDV--ESAASSEDCLYLNI   1948
Cynoglossus_semilaevis                PYARPPTSSLRFQNAQ---PFNWTGTWDAT--KPRATCLQPGDT--TSDSSSEDCLYLNI   1918
Lepisosteus_oculatus                  PYALPPTGNNRFRPPQ---PPQWTDTWNAT--YTRPSCLQPGDD--QTLAADEDCLYLNI   2269
Lampetra_fluviatilis                  PYAAPPLDANRFSGPQAPRPLAVNVTWDADRYK--PDCLQPDGQ--RGTSLSEDCLYLNI   2381
Lampetra_planeri                      PYAAPPLDANRFSGPQAPRPLAVNVTWDADRYK--PDCLQPDGQ--RGTSLSEDCLYLNI   2318
Petromyzon_marinus                    PYAAPPLDANRFSGPQPPRPLAVNETWDADRYKTRPDCLQPDGQ--RGTSLSEDCLYLNI   2333
```

| | | |
|---|---|---|
| *Homo_sapiens* | FIPQNVAPNASVLVFFHNTMDREESEGWPAIDGSFLAAVGNLIVVTASYRVGVFGFLS-S | 2345 |
| *Pan_troglodytes* | FIPQNVAPNASVLVFFHNTMDGEESEGWPAIDGSFLAAVGNLIVVTASYRVGVFGFLS-S | 2344 |
| *Gorilla_gorilla* | FIPQNVAPNASVLVFFHNTMDGEESEGWPAIDGSFLAAVGNLIVVTASYRVGVFGFLS-S | 2332 |
| *Pongo_pygmaeus* | FIPQNVAPNASVLVFFHNTMDGEESEGWPAIDGSFLAAVGNLIVVTASYRVGVFGFLS-S | 2346 |
| *Macaca_mulatta* | FIPQNVAPNASVLVFFHNTMDREGSEGWPAIDGSFLAAVGNLIVVTASYRVGVFGFLS-S | 2346 |
| *Macaca_fascicularis* | FIPQNVAPNASVLVFFHNTMDREGSEGWPAIDGSFLAAVGNLIVVTASYRVGVFGFLS-S | 2346 |
| *Rattus_norvegicus* | FVPENLVSNASVLVFFHNTVEMEGSGGQLNIDGSILAAVGNLIVVTANYRLGVFGFLS-S | 2346 |
| *Mus_musculus* | FVPENLVSNASVLVFFHNTMEMEGSGGQLTIDGSILAAVGNFIVVTANYRLGVFGFLS-S | 2344 |
| *Cavia_porcelus* | FVPGNLASRAPVLVFFHNAAEEDVSKGQLAIDGSFLAAIGNLIVVTANYRTGVFGFLR-S | 2344 |
| *Canis_lupus_familiaris* | FVPQNVAPNASVLVFFHNTLEGRGSEGPLAIDGSYLAAIGNLIVVTAGYRVGIFGFLS-S | 2345 |
| *Panthera_leo* | FVPQNAAPNASVLVLFHNTVEGRGSEGQLAIDGSFLAAIGNLVVVTAGSRVGVFGFLS-S | 2347 |
| *Bos_taurus* | FVPQNMAPNASVLVFFHNAAEGKGSGDRPAVDGSFLAAVGNLIVVTASYRTGIFGFLS-S | 2346 |
| *Trichechus_manatus_latirostris* | FVPESVVPNASVLVFFHNTREGKEREGRLTIDGSFLAAVGNLIVVTASYRVGIFGFLS-S | 2347 |
| *Columbia_livia* | FVPATTVKNMSVLLFFHNGGSYSAEVGKTTIDGSYLAAISNIIVVTANYRVGVFGFLS-T | 2347 |
| *Taeniopygia_guttata* | FVPATTVKNMSVLVFFHNGGSYNAEEGKTTLDGSYLAAIGNIIVVTANYRVGVFGFLS-T | 2351 |
| *Larus_michahellis* | FVPATTVKNMSVLLFFHNGGSYNAEAGKTTIDGSYLAAVSNAIVVTANYRVGVFGFLS-T | 2352 |
| *Gallus_gallus* | FVPVTTVKNMSVLLFFHSGGSGSTEAGKTIIDGSYLAAVSNTIVVTADYRVGVFGFLS-V | 2343 |
| *Struthio_Camelus* | FVPITTTKNLSVLLFFHNGGSDNAEMGKATIDGSYLAAISNIIVVIANYRVGVFGFLS-T | 2325 |
| *Chelonia_mydas* | FAPASIGRNMAVLMFFHNGVSGEGEKGRTVVDGSYLASVSDVIVVTASYRVGVFGFLS-T | 2340 |
| *Alligator_mississippiensis* | FVPADTVRNTSVLLFFHNGGNDGTEKGNAAIDGSYLAAVSDVIVVIASYRVGIFGFLS-T | 2325 |
| *Crotalus_tigris* | FVPQNNAGNLPVLVFFRNSINADDQAKKAVLDGSYLAGVGNLIVVTANFRVGVFGFFS-G | 2334 |
| *Python_bivittatus* | FVPQNNAGNLPVLVFFHNSIDADDQAKKTVLDGSYLAGVGNLIVVTASYRVGVFGFFS-T | 2277 |
| *Eublepharis_macularius* | YVPDNNDGNLSVLVFFHNGGAGDSGQTRTLVDGSFLAGVGNIIVVTANYRVGVFGFLT-T | 2331 |
| *Xenopus_tropicalis* | FVPQHAGTNAAVLLYFHNSPSDYSEKGQTFIDGSYLAAIGNIVVVTAGYRVGVFGFLSNT | 2362 |
| *Aquarana_Catesbeiana* | FVPHITRQSTAVLLYFHNSPSDYSETGQTFIDGSYQAAIGDIIVVTAGYRVGVFGFLS-A | 2341 |
| *Carassius_auratus* | FVASSVGKNSPVLVFFHNS-------GSGLLDGSYLAAVGNIIVVTASFRVAAFGFLS-A | 2308 |
| *Cyprinus_carpio* | FVASSVGKNAPVLVFFHNS-------GSGLLDGSYLTAVGNIIVVTASFRVAAFGFLS-A | 2302 |
| *Danio_rerio* | FVASSVAKNAPVLVFFHNS-------GSDLLDGSYLAAVGNIIVVTASFRMAAFGFLS-A | 2302 |
| *Astyanax_mexicanus* | FVPSSVKRAAPVLVFFHNA-------GSGLLDGSYLAAVGNIIVVTVNFRVAAFGFLS-T | 2297 |
| *Clupea_harengus* | FAPAGL-RDAPVLVFFHNPSSAVSTDGPGLLDGSYLAAVGNIIVVTASFRVSAFGFLS-T | 2331 |
| *Oryzias_latipes* | FTPAARRGRVPVLVFFFNPSAD-E--TPGLLDGSALAAVGNIVVVTASYRTAALGFLS-T | 1981 |
| *Xiphophorus_maculatus* | FTLAAQRGRVPVLVYFFNPPAN-QNQNQNLLDGSALAALGNVVVVTASYRTAALGFLT-A | 1990 |
| *Stegastes_partitus* | FTPSALRGRVPVLVFFFNPSAS-Q--SPGLLDGSTLAAVGNIIVVTASYRTAALGFLS-T | 2004 |
| *Cynoglossus_semilaevis* | FSPAARRGNVPVLVFFFNPSAN-P--SPALLHGSTLAAVGNIVVVTASYRTAALGFLS-T | 1974 |
| *Lepisosteus_oculatus* | YVPRSIRDNSSVLIFFHNPVRDASNTGQDPLDGSYLAAVGDIIVVTVSFRVGAFGFLS-A | 2328 |
| *Lampetra_fluviatilis* | FVPKVKPHNASVLVFFPGGDNSFGGSAQGPLDPSYLAALGDIIVVTANYRLGLFGFLS-T | 2440 |
| *Lampetra_planeri* | FVPKVKPHNASVLVFFPGGDNSFGGSAQGPLDPSYLAALGDIIVVTANYRLGLFGFLS-T | 2377 |
| *Petromyzon_marinus* | FVPKVKPHNASVLVFFHGGDNSFGGSAQGPLDPSYLAALGDIIVVTANYRLGLFGFLS-T | 2392 |

| | | |
|---|---|---|
| *Homo_sapiens* | GSGEVSGNWGLLDQVAALTWVQTHIRGFGGDPRRVSLAADRGGADVASIHLLTARATNSQ | 2405 |
| *Pan_troglodytes* | GSGDVSGNWGLLDQVAALTWVQTHIRGFGGDPRRVSLAADRGGADVASIHLLTARATNSQ | 2404 |
| *Gorilla_gorilla* | GSGEVSGNWGLLDQVAALTWVQTHIRGFGGDPRRVSLAADHGGADVASIHLLTARATNSQ | 2392 |
| *Pongo_pygmaeus* | GSGEVSGNWGLLDQVAALTWVQTHIRGFGGDPRRVSLAADRGGADVASIHLLTARATNSQ | 2406 |
| *Macaca_mulatta* | GSGEVSGNWGLLDQVAALTWVQTHIRGFGGDPRRVSLAADRAGADVASIHLLMARATNSQ | 2406 |
| *Macaca_fascicularis* | GSGEVSGNWGLLDQVAALTWVQTHIRGFGGDPRRVSLAADRAGADVASIHLLMARATNSQ | 2406 |
| *Rattus_norvegicus* | GSDEVAGNWGLLDQVAALTWVQTHIGAFGGDPQRVTLAADRGGADVASIHLLITRPTRLQ | 2406 |
| *Mus_musculus* | GSDEVAGNWGLLDQVAALTWVQSHIGAFGGDPQRVTLAADRSGADVASIHLLISRPTRLQ | 2404 |
| *Cavia_porcelus* | GSSEVTGNWGLLDQVAALTWVQIHIGAFGGDPRQVSLAADRAGADVASIHLLTTRAAGPR | 2404 |
| *Canis_lupus_familiaris* | GSGELSGNWGLLDQLAALTWVQTHIGVFGGDPRRVTLAADRGGADVASIHLLTTRAADSR | 2405 |
| *Panthera_leo* | GSGEVSGNWGLLDQLAALTWVQTHIAVFGGDPRRVTLAADRGGADVAGIHLLTSRASDSR | 2407 |
| *Bos_taurus* | GSSELSGNWGLLDQVVALTWVQTHIQAFGGDPRRVTLAADRGGADIASIHLVTTRAANSR | 2406 |
| *Trichechus_manatus_latirostris* | GSSDVSGNWGLMDQVAALSWVQTHIREFGGDPQRVSLAADRGGADVASIQLLTTRAVNAR | 2407 |
| *Columbia_livia* | GSSEMSGNAGLLDQLAALTWVQQNIVSFGGDPSRVSLGADRGGADVTSIHLLT-EAADMG | 2406 |
| *Taeniopygia_guttata* | GSAEVSGNAGLWDQLAALRWVQQNIASFGGDPGRVSLGAARGGADIASVHLLT-DTAGAD | 2410 |
| *Larus_michahellis* | GSPEASGNAGLLDQLAALKWVQQNIASFGGDPSRVSLGADRGGADVTSIHLLT-EAASMD | 2411 |
| *Gallus_gallus* | GSAEASGNAGLLDQLAALKWVQQNIGSFGGDPSMVSVGADRGGADIASIHLLT-ETINMG | 2402 |
| *Struthio_Camelus* | GSPEARGNAGLLDQLAALKWVQQNIASFGGDPSQVSLGADRSGADITSLHLLT-ETADMD | 2384 |
| *Chelonia_mydas* | GSSVARGNWGLLDQVEALKWVQKNIASFGGDPSQISIAADRSGADITSIHLLA-GAADSN | 2399 |
| *Alligator_mississippiensis* | GSQVATGNWGLLDQVAALRWVKKNIASFGGDPRQISIAADRSGADITSIHLLA-KSVDSN | 2384 |
| *Crotalus_tigris* | GSNVASGNWGLWDQDMALEWIRRNIASFGGDKGQITIGASQRGADLISIHLLK-RTSNPI | 2393 |
| *Python_bivittatus* | GSSVANGNWGLLDQEMALKWIQKNIAMFGGDAGQITIGANERGADFISTHLLK-RALNPI | 2336 |
| *Eublepharis_macularius* | GSSVASGNWGLLDQEAALKWTRENIASFGGDPSHITIAADRSGADVASLHLVT-GAVDSG | 2390 |
| *Xenopus_tropicalis* | GETAPSGNWGLLDQVAALKWVQENIAYFGGDPTLISIAADRAGADISSIHTIV---PEGK | 2419 |
| *Aquarana_Catesbeiana* | GKTVPNGNWGLLDQIAALKWVQENIAYFGGDPKQICVGADGAGADISSLHLLS---AENK | 2398 |
| *Carassius_auratus* | DSSALPGNYGLQDQAAALGWVQKNIALFGGDPTKVTVGAERNGADIASLHLTS--PSASS | 2366 |
| *Cyprinus_carpio* | DSSALPGNYGLQDQAAALGWVQKNIALFGGDPTKVTVGAERNGADIASLHLTS--PSASS | 2360 |
| *Danio_rerio* | GSSALPGNYGLQDQAAALGWVQKNIALFGGDPTKVTVGAERNGADIASLHLTS--PSASS | 2360 |
| *Astyanax_mexicanus* | GSGTLPENFGLQDQAAALGWVQKNIALFGGDPTKVTVGADRNGADLASLHLSS--SSSSS | 2355 |
| *Clupea_harengus* | GSDSLPGNYGLLDQVAALQWVRKNIAQFGGDSRRVTLGAERNGADIASLHLTS--SSSSG | 2389 |
| *Oryzias_latipes* | GASGLRGNFGLTDQEAVLRWVNAYISLVGGDYNRVTVGAERGGADITSLHLLS---KPRP | 2038 |
| *Xiphophorus_maculatus* | G--GLHGNYGLSDQEAALRWVHAHISLVGGDSGSVTVGAERRGADISSLTLLS---SSPP | 2045 |
| *Stegastes_partitus* | GVSGLRGNYGLSDQEAVLRWVNAHISLVGGDNTKVTVGAERGGADITSLHLLS---SSPP | 2061 |
| *Cynoglossus_semilaevis* | GDAHLRGMYGLSDQEAVLRWVHAHIALMGGDNSRVTVGAERGGADITSLHLLS---SASS | 2031 |
| *Lepisosteus_oculatus* | GSEAAPGNYGLQDQVLALRWIQSNIGHFGGDPGKVTLGAERSNADIASLLLTS--AVAQG | 2386 |
| *Lampetra_fluviatilis* | GDEAAAGNWGLLDQQAALRWVRDHAALFGGSAGAVTVAGDRSSADNVGLHLVA--PGSRG | 2498 |
| *Lampetra_planeri* | GDEAAAGNWGLLDQQAALRWVRDHAALFGGSAGAVTVAGDRSSADNVGLHLVA--PGSRG | 2435 |
| *Petromyzon_marinus* | GDEAAAGNWGLLDQQAALRWVRDHAALFGGSAGAVTVAGDRSSADNVGLHLVA--PGSRG | 2450 |

| | | |
|---|---|---|
| *Homo_sapiens* | LFRRAVLMGGSALSPAAVISHE---RAQQQAIALAKEVSCPMSSSQ---EVVSCLRQKPA | 2459 |
| *Pan_troglodytes* | LFRRAVLMGGSALSPAAVISHE---RAQQQAIALAKEVSCPMSSSQ---EVVSCLRQKPA | 2458 |
| *Gorilla_gorilla* | LFRRAVLMGGSALSPAAVISHE---RAQQQAIALAKEVSCPTSSSQ---EVVSCLRQKPA | 2446 |
| *Pongo_pygmaeus* | LFRRAVLMGGSALSPAAVISHE---RAQQQAIALAKEVSCPMSSSQ---EVVSCLRQKPA | 2460 |
| *Macaca_mulatta* | LFRRAVLMGGSALSPAAIISHE---RAQQQAVALAKEVSCPVSSSQ---EVVSCLRQKPA | 2460 |
| *Macaca_fascicularis* | LFRRAVLMGGSALSPAAIISHE---RAQQQAVALAKEVSCPVSSSQ---EVVSCLRQKPA | 2460 |
| *Rattus_norvegicus* | LFRKALLMGGSALSPAAIISPD---RAQQQAAALAKEVGCPNSSVQ---EVVSCFRQKPA | 2460 |
| *Mus_musculus* | LFRKALLMGGSALSPAAIISPE---RAQQQAAALAKEVGCPTSSIQ---EVVSCLRQKPA | 2458 |
| *Cavia_porcelus* | LFRRVLLMGGSALSPAAIISPS---RAQQQTMALAEEVNCPTSSSS---EVESCLRQTPA | 2458 |
| *Canis_lupus_familiaris* | LFRRAILMGGSAFSPAVVISQG---RAQEQAAALAEEIGCPTSSTQ---ELVSCLRQKPA | 2459 |
| *Panthera_leo* | LFRRAVLMGGSAFSPAAVISQE---RARQQAAALAEEVGCPPSSIQ---DMVSCLRQKPA | 2461 |
| *Bos_taurus* | LFRRAVLMGGSALSPAAVIRPE---RARQQAAALAKEVGCPSSSVQ---EMVSCLRQEPA | 2460 |
| *Trichechus_manatus_latirostris* | LFRRAVLMGGSALSPAAIISQE---RAQQQATALAKEVDCPTSPSQ---DLVSCLRQKPA | 2461 |
| *Columbia_livia* | LFRSALLMGGSAFSPASIISKR---RAQIQAAVLADDVGCPSSTSE---EIVACLRQLPA | 2460 |
| *Taeniopygia_guttata* | LFSRLLLMGGSAFSPASIITTR---RAQTQAAALAEDVGCPSSTSE---EIVACLRQLPA | 2464 |
| *Larus_michahellis* | LFKRVLLMGGSAFSPASVISKR---RARTQAAVLAEDVECPSSTPE---EIVACLRQLPA | 2465 |
| *Gallus_gallus* | LFRRMLLMGGSAFSPASIITKR---RAQAQAAVLADDVGCPSSTSE---EIVSCLRQLPA | 2456 |
| *Struthio_Camelus* | LFKRVLLMGGSAFSPASIITEK---RAQTQVAILADDLGCPSSTSE---EIVACLRQMPA | 2438 |
| *Chelonia_mydas* | LFKRALLMGGSAFSPVSVISEK---KAQDQVAVLANELGCPSSSSE---EIVSCLRQLPA | 2453 |
| *Alligator_mississippiensis* | LFERAVLMGGSAFSPVSVISKK---TAQEQVRVLANELGCPSSSSK---ETITCLRQLPA | 2438 |
| *Crotalus_tigris* | LFRRALLMGGSAFSPAITLSKE---KAQVQAAVLAREVGCPSTRSP---ELASCLRQLPA | 2447 |
| *Python_bivittatus* | LFKRALLMGGSAFSPAVTLSKE---KARAQAAVLARELGCPSTSSA---ELVSCLRQLPA | 2390 |
| *Eublepharis_macularius* | LFKRALLMGGSAFSPASTISKE---KAREQAADLAAEFGCPSDNSE---ESLSCLRQLPA | 2444 |
| *Xenopus_tropicalis* | LFRRALLMGGSAFSPMMVTSEK---RAKEQVEFLAAEVECSSSN-K---DLLTCLRGVDA | 2472 |
| *Aquarana_Catesbeiana* | AIRRAVLMGGSAFSPMLTVSGR---RAQEQVKFLAEEVGCPATDDE---GTLACLGKVDA | 2452 |
| *Carassius_auratus* | LFRRALLMGGSVFSPAVVTSMS---KAQAQTASLARELGCSASDIS---QLLTCLRSKPA | 2420 |
| *Cyprinus_carpio* | LFRRALLMGGSVFSPAVVISTS---KAQAQTASLARELGCSASDIS---QLLTCLRSKPA | 2414 |
| *Danio_rerio* | LFSRALLMGGSVFSPAVVMSSS---KAQAQTSSLARELDCQAADPS---QLLDCLRSKPA | 2414 |
| *Astyanax_mexicanus* | LFRQALLMGGSTFSPAAVMSKR---KAQEQAASLSTELGCSSSEPS---QILTCLRGKTA | 2409 |
| *Clupea_harengus* | LFLRALLMGGSVFSPAAVQSRS---RAQEQSSSLGREMGCSSTDSA---QLLACLRQAPA | 2443 |
| *Oryzias_latipes* | LFQRMMLMGGSVFSPSLVQTSA---SARRRALDLAQELGCVTSDST-DEKLVSCLRAVPY | 2094 |
| *Xiphophorus_maculatus* | -FHRMILMGGSVFSPSLVQTAA---AGRRQAVDLARELGCVTSDPSDDGPMVACLRSTPV | 2101 |
| *Stegastes_partitus* | LFQRMMLMGGSVFSPVLVQTPS---TSRRQAVDLATELGCPTS---DDDKIAACLRAAPV | 2115 |
| *Cynoglossus_semilaevis* | LFHRMILMGGSLFSPSSVSSSS-----SSLVMDLVSELGCVSSDPS---VISSCLRALSA | 2083 |
| *Lepisosteus_oculatus* | LFHRVLLLGGTAFSPASVINSF---KAREKTTSLGTEVGCSSTDHD---QLISCLRTVPA | 2440 |
| *Lampetra_fluviatilis* | LFQRAILMGGSVLSPSAVQLDSAAAAARSQAAALAQLVGCGHSSSE---YLVRCLRSRSA | 2555 |
| *Lampetra_planeri* | LFQRAILMGGSVLSPSAVQLDSAAAAARSQAAALAQLVGCGHSSSE---YLVRCLRSRSA | 2492 |
| *Petromyzon_marinus* | LFQRAILMGGSVLSPSAVQLDSAT--ARSQAASLAQLVGCGHSSSE---YLVRCLRDRSA | 2505 |

| | | |
|---|---|---|
| *Homo_sapiens* | NVLNDAQTKLLAVSGPFHYWGPVIDGHFLREPPARALKRSLWVEVDLLIGSSQDDGLINR | 2519 |
| *Pan_troglodytes* | SVLNDAQTKLLAVSGPFHYWGPVIDGQFLREPPARALKRSLRVEVDLLIGSSQDDGLINR | 2518 |
| *Gorilla_gorilla* | SVLNDAQTKLLAVSGPFHYWGPVIDGQFLREPPARALKRSLRVEVDLLIGSSQEDGLINR | 2506 |
| *Pongo_pygmaeus* | NVLNDAQTKLLAVSGPFHYWGPVIDGQFLREPPARALKRSLWLEVDLLIGSSQDDGLINR | 2520 |
| *Macaca_mulatta* | NILNDAQTKLLAVSGPFHYWGPVIDGQFLREPPARALKRSLRAEVDLLIGSSQDDGLINR | 2520 |
| *Macaca_fascicularis* | NILNDAQTKLLAVSGPFHYWGPVIDGQFLREPPARALKRSLRAEVDLLIGSSQDDGLINR | 2520 |
| *Rattus_norvegicus* | NILNEAQTKLLAVSGPFHYWGPVVDGQYLRELPSRRLKRPLPVKVDLLIGGSQDDGLINR | 2520 |
| *Mus_musculus* | NILNDAQTKLLAVSGPFHYWGPVVDGQYLRELPSRRLKRPLPVKVDLLIGGSQDDGLINR | 2518 |
| *Cavia_porcelus* | NILNDAQTKLLAVSGPFHYWGPVVDGQYLLEDPARALRRPPLAKIDLLIGSSQDDGLINR | 2518 |
| *Canis_lupus_familiaris* | SILNDAQTKLLAVSGPFHYWGPVVDGQYLREAPARALQRTLRAKVDLLIGSSQDDGLINR | 2519 |
| *Panthera_leo* | SVLNDAQTKLLAVSGPFHYWGPVVDGQYLREAPARALQRSPRTKVDLLIGSSQDDGLINR | 2521 |
| *Bos_taurus* | RILNDAQTKLLAVSGPFHYWGPVVDGQYLRETPARVLQRAPRVKVDLLIGSSQDDGLINR | 2520 |
| *Trichechus_manatus_latirostris* | SVLNDAQTKLLAVSGPFHYWGPVVDGQYLREAPARALQRAPRAKVDLLIGSSQDDGLIDR | 2521 |
| *Columbia_livia* | RVLNDAQTKLLAISGPFQYWGPVMDGVYVQEPLAKALQRPRLRKVDLLIGSAQQDGLISR | 2520 |
| *Taeniopygia_guttata* | RVLNDAQTKLLAISGPFQYWGPVMDGIYLREPLAKALQRPRLRKVDLLIGSAQQDGLISR | 2524 |
| *Larus_michahellis* | RVLNDAQTKLLAISGPFQYWGPVMDGIYLREPLAKALQRPQLRKVDLLIGSAQQDGLIIR | 2525 |
| *Gallus_gallus* | RVLNDAQTKLLAISGPFQYWGPVVDEVYLQEPLAKALQRPQLRKVDLLIGSAQQDGLISR | 2516 |
| *Struthio_Camelus* | RVLNNAQTKLLAISGPFQYWGPVMDGIYLREPLAKALQRRQLRKVDLLIGSAQQDGLISR | 2498 |
| *Chelonia_mydas* | NTLNDAQTRLLAISGPFQYWGPVVDGIYLREPLAAAVQRSRLRKVDLLIGSAQQDGLISR | 2513 |
| *Alligator_mississippiensis* | SALNDAQTKLLAISGPFQYWGPVVDGIYLQEPLAAALQRTRLKKVDLLIGSAQQDGLISR | 2498 |
| *Crotalus_tigris* | KTLNDAQTKLLAISGPFQYWGPVVDGSYLQEPFPAALQRSRLSKMDILIGSAQQDGLISR | 2507 |
| *Python_bivittatus* | KALNDAQTKLLAISGPFQYWGPVVDGSYLQESFPAALQRSRPLKMDLLIGSAQQDGLIIR | 2450 |
| *Eublepharis_macularius* | KALNDAQTRRLAVSGPFQYWSPVVDGVYLQEPLTSAVQRSRAVKLDLLIGSSQEDGLISR | 2504 |
| *Xenopus_tropicalis* | GALNAAQTKLLAQRGPFQTWGPIVDKLYVKETPSRMLKQKKFQKIDLLIGSSELDGLISR | 2532 |
| *Aquarana_Catesbeiana* | IALNAAQTKLLAVRGPFQTWGPVIDGSYLRDSPTRLLQQKTAQNIDLFIGSTEHDGLISR | 2512 |
| *Carassius_auratus* | QTINAAQTKLLAVSGPLQAWSPVVDGTVVREKPSVALLSGNFHKAELLLGTSFEDGLISR | 2480 |
| *Cyprinus_carpio* | QSINAAQTKLLAVSGPLQAWSPVVDGTVVREKPSVALLSGHFHKAELLLGSSFEDGLISR | 2474 |
| *Danio_rerio* | HSINAAQTKLLAVSGPLQAWSPVVDGNVVREMPSVALQSGRFHKAEILLGSSFEDGLISR | 2474 |
| *Astyanax_mexicanus* | QSINAAQTKLLAVSGPLQAWSPVVDGAVVREDPTTALNSGRFYKVPLLLGSSAEDGLISR | 2469 |
| *Clupea_harengus* | EAINAAQTKLLAVSGPLQAWSPVVDGLSVKEDPASAFRGSRFHRGDLLMGSSAEDGLISR | 2503 |
| *Oryzias_latipes* | RDLNAAQTKLLVTRDPFQFWSPVHQAVSK---------SSPFHKVDLLLGTSEADGLISR | 2145 |
| *Xiphophorus_maculatus* | HKLNAAQTKLLAVSGPFQSWAPVRR--FE---------SESFHRVDLMLGTSEHDGLISR | 2150 |
| *Stegastes_partitus* | QQLNAAQTKLLAVRGPFQSWSPVRQAVFQ---------PASFHRVDLLLGTSEHDGLISR | 2166 |
| *Cynoglossus_semilaevis* | QTLNEVQTKLLAVSGPFHSWSPPRQSTLQ-------VPLSTLSRVDLLLGTSEQDGLIGR | 2136 |
| *Lepisosteus_oculatus* | LTLNAAQTKLLAVSGPFQAWAPVVDGISVQEPPSTAFHSSRFHNVDLMIGSSADDGLIGR | 2500 |
| *Lampetra_fluviatilis* | LALNAAQAQLLS-GEYLQRWAAVVDGHFLRDIPAEAAARGALADVDIVLGATEDDGALGR | 2614 |
| *Lampetra_planeri* | LALNAAQAQLLS-GEYLQRWAAVVDGHFLRDIPAEAAARGALADVDIVLGATEDDGALGR | 2551 |
| *Petromyzon_marinus* | LALNAAQAQLLS-GEYLQRWAAVVDGHFLRDIPAEAAARGALADVDIVLGATEDDGALGR | 2564 |

| | | |
|---|---|---|
| *Homo_sapiens* | AKAVKQFEESRGRTSSKTAFYQALQNSLGGEDSDARVEAAATWYYSLEHST--DD-YASF | 2576 |
| *Pan_troglodytes* | AKAVKQFEESQGRTSSKTAFYQALQNSLGGEDSDARVEAAATWYYSLEHST--DD-YASF | 2575 |
| *Gorilla_gorilla* | AKAVKQFEESQGRTSSKTAFYQALQNSLGGEDSDARVEAAATWYYSLEHST--DD-YASF | 2563 |
| *Pongo_pygmaeus* | AKAVKQFEESQGRTSSKTAFYQALQNSLGGEDSDARVEAAATWYYSLEHST--DD-YASF | 2577 |
| *Macaca_mulatta* | AKAVKQFEESQGRTSSKTAFYQALQNSLGGEDSDARVEAAATWYYSLEHST--DD-YASF | 2577 |
| *Macaca_fascicularis* | AKAVKQFEESQGRTSSKTAFYQALQNSLGGEDSDARVEAAATWYYSLEHST--DD-YASF | 2577 |
| *Rattus_norvegicus* | AKAVKQFEESQGRTNSKTAFYQALQNSLGGEDSDARILAAAIWYYSLEHST--DD-YASF | 2577 |
| *Mus_musculus* | AKAVKQFEESQGRTNSKTAFYQALQNSLGGEDSDARILAAAVWYYSLEHST--DD-YASF | 2575 |
| *Cavia_porcelus* | AKAVKRFEESQGRTESKTAFYQALQNSLGGEDSDASVLAAATWYYSLEHSQ--DD-YASF | 2575 |
| *Canis_lupus_familiaris* | AKAVKQFEESQGRTSSKTAFYQALQNSLGGEDADAGVRAAATWYYSLEHST--DD-YASF | 2576 |
| *Panthera_leo* | AKAVKQFEESQGRTSSKTAFYQALQNSLGGEDADAGVLAAATWYYSLEHST--DD-YASF | 2578 |
| *Bos_taurus* | AKAVKQFEESQGRTSSKTAFYQALQNSLGGEAADAGVQAAATWYYSLEHDS--DD-YASF | 2577 |
| *Trichechus_manatus_latirostris* | AKAVKQFEESQGRTSSKTAFYQALQNSLGGEDGDTRVEAAATWYYSLEHST--DD-YASF | 2578 |
| *Columbia_livia* | AKAIKKFEESQGRSNSKTAFYQALQNSLGGEDSNSFIEDAATWYYSLEHST--DD-YSSF | 2577 |
| *Taeniopygia_guttata* | AKAIKKFEESQGRANSKTAFYQALQNSLGGEESNSFIEDAATWYYSLEHST--DD-YSSF | 2581 |
| *Larus_michahellis* | AKAIKKFEERQGRANSKTAFYQALQNSLGGEDSNSFIEDAATWYYSLEHST--DD-YSSF | 2582 |
| *Gallus_gallus* | AKAIKKFEESQGRTNSKTTFYQALQNSLGGEDSNLFIENAATWYYSLEHST--DD-YSTF | 2573 |
| *Struthio_Camelus* | AKAIKKFEESQGRGNSKTAFYQALQNSLGGEESNVFLEDAATWYYSLEHST--DD-YSSF | 2555 |
| *Chelonia_mydas* | AKAIKKFEESQGRANSKTAFYQALQNSLGGEDSNSFIEDAAAWYYSLQHST--AD-YASF | 2570 |
| *Alligator_mississippiensis* | AKAIKKFEERKGRVSSKTAFYQALQNSLGGEDSNSFIEDAATWYYSLRHST--ED-YATF | 2555 |
| *Crotalus_tigris* | AKAIKRFEESQGRTNSKTAFYQALQNSLGGEESNLLIQDAATWFYSLEHSS--TE-YASF | 2564 |
| *Python_bivittatus* | AKAIKRFEESQGRANSKTAFYQALQNSLGGEESNLLIQDAVTWFYSLQHSS--TE-YASF | 2507 |
| *Eublepharis_macularius* | AKAVKKFEERQGRANSKTVFYQALQNSLGGEDSNPMVQDAASWFYSLQHSS--DD-YASF | 2561 |
| *Xenopus_tropicalis* | AKAIKRFEETQGRVD-KTAFYQALQNSLGGVEANSLLQEAAVWFYSLEHSS--DD-YTGF | 2588 |
| *Aquarana_Catesbeiana* | AKAIKRFEESQGRGDSKMSFYQALQNSLGGEEMNPLVQEAAVWFYSLQHST--DD-YSSF | 2569 |
| *Carassius_auratus* | AKDIKNFEQLQGRADSKTAFYAALSNSLGGDDANAFVKEAATWFYSMQHSPTPSG-YNVF | 2539 |
| *Cyprinus_carpio* | AKNIKNFEQLQGRADSKTAFYAALSNSLGGDDANAFVKEAATWFYSLQHSPTPSG-YNVF | 2533 |
| *Danio_rerio* | AKNIKNFEQLQGRADSKTAFYAALSNSLGGDDANAFVKEAATWFYSLQHSPTPSG-YNVF | 2533 |
| *Astyanax_mexicanus* | AKNIKNFELLQGRADSKTAFYEALSNSLGGDDANAFVKEAATWFYSLQHSPTPSG-YNVF | 2528 |
| *Clupea_harengus* | ARNIKRFEELQGRSDGKTAFYEALSNSLGGPEASAFVKEAASWFYSLHHTPSPAG-YNVF | 2562 |
| *Oryzias_latipes* | ARRIEEFEVLPGQVDDSMPFYQALSRVLGGANGNELLGEV---VSSLDHNPSAAG-YSLF | 2201 |
| *Xiphophorus_maculatus* | ARRIKNFEALQGRADGKTAFYEALSRSLGGASGNALLKDAAAWFYALDHEPSAAG-YNLF | 2209 |
| *Stegastes_partitus* | ARRIKDFEVLQGRADGKTAFYEALSRSLGGASGNQLLKDAAAWFYSLDHSPSAAG-YNLF | 2225 |
| *Cynoglossus_semilaevis* | ARRIKDFEALQGRVNGKTAFYEALSRSLGGETGNPLLKEAATWFYSLDHSPTPQG-YSLF | 2195 |
| *Lepisosteus_oculatus* | AKRIKKFEVLQGRASSKTAFYEALSNSLGGDDSNSFVKDAATWFYSLLHDPSPVG-YNVF | 2559 |
| *Lampetra_fluviatilis* | SRLPQVFPRFTTMSYSSAGFDVALQKVLRGAGDSSFARDAVRWFYKRGDGADPPPLAADE | 2674 |
| *Lampetra_planeri* | SRLPQVFPRFTTMSYSSAGFDVALQKVLRGAGDSSFARDAVRWFYKRGDGADPPPLAADE | 2611 |
| *Petromyzon_marinus* | SRLPQVFPRFTSMSYSSAGFDVALQKVLRGAGASSFARDAVRWFYKRGDGANPPPLADDE | 2624 |

| | | |
|---|---|---|
| *Homo_sapiens* | SRALENATRDYFIICPIIDMASAWAKRARGNVFMYHAPENYGH--GSLELLADVQFALGL | 2634 |
| *Pan_troglodytes* | SRALENATRDYFIICPIIDMASAWAKRARGNVFMYHAPESYGR--GSLELLADVQFAFGL | 2633 |
| *Gorilla_gorilla* | SRALENATRDYFIICPIIDMASAWAKRARGNVFMYHVPESYGH--GSLELLADVQFAFGL | 2621 |
| *Pongo_pygmaeus* | SRALENATRDYFIICPIIDMASAWAKRARGNVFMYHAPESYGH--GSLELLADVQFAFGL | 2635 |
| *Macaca_mulatta* | SRALENATRDYFIICPIIDMASAWAKRARGNVFMYHAPESYGR--GSLELLADVQFAFGL | 2635 |
| *Macaca_fascicularis* | SRALENATRDYFIICPIIDMASAWAKRARGNVFMYHAPESYGR--GSLELLADVQFAFGL | 2635 |
| *Rattus_norvegicus* | SRALENATRDYFIICPIVNMASLWARRTRGNVFMYHVPESYGH--GSLELLADVQYAFGL | 2635 |
| *Mus_musculus* | SRALENATRDYFIICPMVNMASLWARRTRGNVFMYHVPESYGH--GSLELLADVQYAFGL | 2633 |
| *Cavia_porcelus* | SRALENATRDYFIICPTIDMARRWAKRARGNVFMYHAPESYGH--GSLELLADVQYAFGL | 2633 |
| *Canis_lupus_familiaris* | SRALENATRDYFITCPVIDMASHWARRARGNVFMYHAPESYDH--RSLELLADVQYAFGL | 2634 |
| *Panthera_leo* | SRALENATRDYFITCPVIDMARHWARRARGNVFMYHAPKSYGH--SSLELLADVQYAFGL | 2636 |
| *Bos_taurus* | SRALEQATRDYFIICPVIDMASHWARTVRGNVFMYHAPESYSH--SSLELLTDVLYAFGL | 2635 |
| *Trichechus_manatus_latirostris* | SWALENATRDYFITCPGIDMASHWARQARGNVFMYHVPGGSGR--SSPELLADVQYAFGL | 2636 |
| *Columbia_livia* | SRALENATRDQFITCPIINMASYWAAASRGNVFMYHVPESSSQSSLGQELLLDIQYAFGL | 2637 |
| *Taeniopygia_guttata* | SRALENATRDQFITCPIINMASYWAAASRGNVFMYHVPESFSQSSSGQEFLLDVQYAFGL | 2641 |
| *Larus_michahellis* | SRALENATRDQFITCPIINMASHWAAASRGNVFMYHVPESSSQSSSGQELLLDVQYAFGL | 2642 |
| *Gallus_gallus* | SRALENATRDQFITCPIINMANHWAANSRGNVFMYHVPASSSESSSGLELLLDVQYAFGL | 2633 |
| *Struthio_Camelus* | SRALENATRDQFITCPIINMANHWAANSRGNVFMYHVPESSSQSSSALELLPDVQYAFGL | 2615 |
| *Chelonia_mydas* | SRALENATRDHFITCPIISMAKHWADNSRGNVFMYHVPVSSSQSSSGLELLPDVQYAFGL | 2630 |
| *Alligator_mississippiensis* | SRALENATRDRFINCPIISMASYWADNSRGNVFMYHVPESSSESSSSLELLPDVQYGFGL | 2615 |
| *Crotalus_tigris* | SRALENATRDHFIICPTIDMAKEWANNKRGNIFMYHVPDSSSLSSSSVEYMPDVQLAFGL | 2624 |
| *Python_bivittatus* | SRALENATRDHFIICPTIDMAKDWASNGRGNIFMYHVPESSSLSSSSLEYVPDVQLAFGL | 2567 |
| *Eublepharis_macularius* | SRALENATRDHFIICPTIEMARNWAGSSRGNVFMYHVPESSSSSSSQELSPDIQYFFGL | 2621 |
| *Xenopus_tropicalis* | SKALEKATRDHFIACPVIKMAKHWAENSKGNIYLYHVPDTLSQSSSGLDLPEDVIRAFGL | 2648 |
| *Aquarana_Catesbeiana* | SRALENSTRDHFIACPAVKMADYWSNNSEGHVFMYYVPETYSQSSSGLDLPEDVIYTFGI | 2629 |
| *Carassius_auratus* | SRALENATRDLFIICPAVDMAEFWAANTRSSVHMYHLPEDSAYNSADLSVPMDVQYLFGV | 2599 |
| *Cyprinus_carpio* | SRALENATRDLFIICPAVDMAEFWAANTRSSVHMYHLPEDTAYNSVDLSVPMDVQYLFGV | 2593 |
| *Danio_rerio* | SRALENATRDLFIICPTVDMAEFWAANTQTGVYMYHLPENAAYNSVDLSVPMDVQYLFGV | 2593 |
| *Astyanax_mexicanus* | SRALENATRDYFITCPSVNMASFWASHTRSSVYMYHLPQDMAHNSADLSAPLDIQYVFGV | 2588 |
| *Clupea_harengus* | SRALESATRDLFIICPSKKMAEFWAANTKSNVFMYHLPENMTHISAALPVPLDVQLVFGL | 2622 |
| *Oryzias_latipes* | SRTLDNATRDLFIICPALQMAANWAE-SRANVFMYHQPAADAADRGDSLVPWDVQLVFGA | 2260 |
| *Xiphophorus_maculatus* | SRALNNATRDLFIVCPTVQMALHWAA-SGTNVFQYHQPASSAQDRADVSVPLDVQFLFGI | 2268 |
| *Stegastes_partitus* | SRALNNATRDLFIICPTLQMANHWAK-SKANVFLFHQPDAGAHSRADVSVPLDVQFVFGT | 2284 |
| *Cynoglossus_semilaevis* | SRSLNNATRDLFITCPSLQMANFWAKNSQSNVFLYHLPASTMNDRADVFVPLDVQLVFGV | 2255 |
| *Lepisosteus_oculatus* | SRALENATRDLFIICPIVKMASFWAANTRSNVFMYHVPEDITQTSTDFAVPMDVQLVFGL | 2619 |
| *Lampetra_fluviatilis* | SWLLSNVSRDYDVVCPAVHMAEMWASRGKANAFLYYVPTQQSWNTLEWEPHSDTQLAFGV | 2734 |
| *Lampetra_planeri* | SWLLSNVSRDYDVVCPAVHMAEMWASRGKANAFLYYVPTQQSWNTLEWEPHSDTQLAFGV | 2671 |
| *Petromyzon_marinus* | SWLLSNVSRDYDVICPAVHMAEMWAARGKANAFLYYVPTQQSWNTLEWEPHSDTQLAFGV | 2684 |

```
Homo_sapiens                      PFYPAYEGQFSLEEKSLSLKIMQYFSHFIRSGNPNYPYEFSRK-VPTFATPWPDFVPRAG     2693
Pan_troglodytes                   PFYPAYEGQFSLEEKSLSLKIMQYFSHFIRSGNPNYPYEFSRK-VPTFATPWPDFVPRAG     2692
Gorilla_gorilla                   PFYPAYEGQFSLEEKSLSLKIMQYFSHFIRSGNPNYPYEFSRK-VPTFATPWPDFVPRAG     2680
Pongo_pygmaeus                    PFYPAYEGQFSLEEKSLSLKIMQYFSHFIRSGNPNYPYEFSRK-VPTFATPWPDFVPRAG     2694
Macaca_mulatta                    PFYPAYEGQFSLEEKSLSLKIMQYFSHFIRSGNPNYPYEFSRK-VPTFATPWPDFVPYAG     2694
Macaca_fascicularis               PFYPAYEGQFSLEEKSLSLKIMQYFSHFIRSGNPNYPYEFSRK-VPTFATPWPDFVPYAG     2694
Rattus_norvegicus                 PFYSAYQGYFSMEEQSLSLKVMQYFSNFIRSGNPNYPHEFSQK-AAEFATPWPDFVPGAG     2694
Mus_musculus                      PFYSAYQGQFSTEEQSLSLKVMQYFSNFIRSGNPNYPHEFSRK-AAEFATPWPDFIPGAG     2692
Cavia_porcelus                    PFYPAYKGQFSLEEKSLSLKIMQFWSNFIRSGNPNYPHEFSWK-APEFAALWPDFVPGAG     2692
Canis_lupus_familiaris            PFYPVYEGQFTLEEKSLSLKIMQFFSNFVRSGNPNYPHEFSRK-ASELAAPWPDFVPRTG     2693
Panthera_leo                      PFHPTYEGQFTLEEKSLSLKIMQFLSNFIRSGNPNYPHEFSRK-APEFATPWPDFVPRAG     2695
Bos_taurus                        PFYPAYEGQFTLEEKSLSLKIMQYFSNFIRSGNPNYPHEFSRR-APEFAAPWPDFVPRDG     2694
Trichechus_manatus_latirostris    PFYPTYEGQFTREEKSLSLKIMQYFSNFIRSGNPNYPHEFSRR-APDFATPWPDFLPSAG     2695
Columbia_livia                    PFYPKYEEQFTLEEKILSLQIMQYISNFVNSGNPNYPHSFSRR-MSGVMPPWPMYLPNDE     2696
Taeniopygia_guttata               PFYPKYEEQFTDEEKGLSLQIMQYISNFINSGNPNYLHSFSRR-MLGVMPAWPMYLSSDD     2700
Larus_michahellis                 PFYPKYEEQFTLEEKSLSLQIMQYISNFVNSGNPNYPHSFSRK-MSEVMPPWPMYLPNDG     2701
Gallus_gallus                     PFYPKYEEQFTLEEKSLSLQVMQYISNFVNSGNPNYPHSFSRK-MSRLLPPWPMFLPNDD     2692
Struthio_Camelus                  PFYPKYEEQFTLEEKSLSLQIMQYISNFVKAGNPNYPHSFSRR-MSGVMPPWPKFLPNDD     2674
Chelonia_mydas                    PFHPQYEEQFTLEEKSLSLKIMQYIGNFIKSGNPNYPHNFSRR-LTGAVSPWPMFLSHAN     2689
Alligator_mississippiensis        PFYPQYEGQFTLEEKSLSLKIMEYIANFVKSGDPNYPHSFSSK-LTRSVSIWPMFLPHAN     2674
Crotalus_tigris                   PFYAQYKNRYTLEEKTLSLAIMQYLSNFIKSGNPNNPYEFSRK-RIEGLPPWPMFRADAG     2683
Python_bivittatus                 PFYPQYKNRYMLEEKTLSLAIMQYLSNFIKSGNPNSPYEFSRK-RIEGVSPWPMFRADAG     2626
Eublepharis_macularius            PFYPQYGNRYSLEEKTLSLAVMQYLANFIRSGNPNRPYAFSRK-ATGHAVPWPMFRADTG     2680
Xenopus_tropicalis                PFHTNYKNQFSSEEQILSLKIMQYVSNFVRTGNPNFPYKFLRR-ISKELPAWPMYLAHSN     2707
Aquarana_Catesbeiana              PFHSLYRNQFSGEERKLSLQIMQYLANFVKTGNPNFPYTYSKK-RNVMLPPWPEYRTQSA     2688
Carassius_auratus                 PLASEMHDLFISKERTLALQIMNYMANFIKSGNPNLPLAASRTSFSKFLPPWPQFMAHQG     2659
Cyprinus_carpio                   HLASEMRDIFSSKERMLALQIMNYMANFIKSGNPNLPLAASRTSFSKFLPPWPQFMPHQG     2653
Danio_rerio                       PLAAEQRDLFSYKEKTFTLQIMNYMANFIKSGNPNLPLAASRTSFGKFLPPWPQFMPHVG     2653
Astyanax_mexicanus                PHSPAMRELFTYTERKLSLQVMSYVANFIKTGNPNLPVSVSTTSFGEVLPPWPQFLPHPN     2648
Clupea_harengus                   PHSTPTQQLFSTQERTLSKQMMTYMANFIKAGDPNTPLSLARVSFAKLLPAWPRVLAQPT     2682
Oryzias_latipes                   PL---YSQRFTSSDRRLSLAAMSYVSSFIRTGNPNPW----PVTAESVLPRWQPVTSSQA     2313
Xiphophorus_maculatus             PLQPMSSQHFTSSDRRLSLAVMTYASSFIRTGNPNPS----RVWAESVLPRWQPV-TAEA     2323
Stegastes_partitus                PHHPVSFQRFSSSDRRLSLAVMSYASTFVQTGNPNPS----RVG--TALPRWQSVLSSEA     2338
Cynoglossus_semilaevis            PHQPTSYQRFTSSNRRLSLAMMSYVASFVKTGNSNPS----KKWAESSLPRWRGVQSFED     2311
Lepisosteus_oculatus              PHHPNTRQLFTEEERSLSLKIMHYVVNFIKSGNPNHPYSFSKLTLSETLPPWPRFLAHPE     2679
Lampetra_fluviatilis              PLHPERKGRSCVAEQQLSRHFIGYLANFVKSGDPSFPNRHARGAGSRLLPAWPRFSLNEA     2794
Lampetra_planeri                  PLHPERKGRSCVAEQQLSRHFIGYLANFVKSGDPSFPNRHARGAGSRLLPAWPRFSLNEA     2731
Petromyzon_marinus                PLHPERKGRSCVGEQQLSRHFIGYLANFVKSGDPSFPNRHARGAGSRLLPAWPRFSLNEA     2744
```

```
Homo_sapiens                      GENYKEFSELLPNRQGLKKADCSFWSKYISSLKTSADGAKGGQ-SAES--------EEEE     2744
Pan_troglodytes                   GENYKEFSALLPNRQGLKKADCSFWSKYISSLKASADGAKGGK-SAES--------E-EE     2742
Gorilla_gorilla                   GENYKEFSALLPNRQGLKKADCSFWSKYISSLKASADGAKGGQ-SAES--------EEEE     2731
Pongo_pygmaeus                    GENYKEFSALLPSRQGLKKADCSFWSKYISSLKASADGAKGGQ-SAES--------EEEE     2745
Macaca_mulatta                    GENYKEFSALLPNRQGLKKADCSFWSKYISSLKASADGAKGGQ-SAES--------EEEE     2745
Macaca_fascicularis               GENYKEFSALLPNRQGLKKADCSFWSKYISSLKASADGAKGGQ-SAES--------EEEE     2745
Rattus_norvegicus                 GESYKELSAQLPNRQGLKKADCSFWSKYIQTLKD-ADGAKDGQ-LTKS--------GEED     2744
Mus_musculus                      GESYKELSAQLPNRQGLKQADCSFWSKYIQTLKD-ADGAKDAQ-LTKS--------EEED     2742
Cavia_porcelus                    GESYQEFSSMLPIRQGLKQADCSFWSKYIQVLKTPADAALDGQ-STES--------EDEG     2743
Canis_lupus_familiaris            GESYKEFSVLLPNRQGLKSADCSFWAKYIQSLKAVADEAKQEL-SAES--------EEDD     2744
Panthera_leo                      GETYKEFSDLLPNRQGLKSADCSFWSKYIQSLKATADEAKQGP-SAES--------DAED     2746
Bos_taurus                        AESYKELSVLLPNRQGLKKADCSFWSKYIQSLKASADETKDGP-SADS--------EEED     2745
Trichechus_manatus_latirostris    GESYKEFSSLLPNRQGLKKADCSFWSKYIQSLKASADGANDQQ-PALR--------EEEE     2746
Columbia_livia                    GDNYKEFTVSLPTLKGLKKADCSFWSDYIRRLKASTGSASNGETFAEN--------SPSE     2748
Taeniopygia_guttata               GDNYKEFTVSLPTLKGLKKADCSFWSDYIRRLKASTGSASGGEPGETP--------S---     2749
Larus_michahellis                 GDNYKEFTVSLPTLKGLKKADCSFWSDYIRRLRASTGSASNEEPSAEN--------SPTD     2753
Gallus_gallus                     GDNYKEFTISMPTLKGLKKADCSFWSDYIRRLKASMGSESNGEQSAEF--------NSTD     2744
Struthio_Camelus                  GDNYKEFTVSLPTLKGLKKGDCSFWSDYIRRLKASTGSASNGEKSAEN--------SPSE     2726
Chelonia_mydas                    GDNYKEFTVSLPNHKGLKRAECSFWSDYIKTLKVSTSRTAIGAQSMES--------SPSE     2741
Alligator_mississippiensis        SDNYKEFTVSLPNRKGLKKAECSFWSDYIETLKESTGKTANGQQSTES--------SLGQ     2726
Crotalus_tigris                   GDNYKEFTASLPNHKGLKKSECSFW-DYIKTVKTLTSCQRQ--QPALS--------RPTK     2732
Python_bivittatus                 GDNYKEFTSSLQNHKGLKKMECSFW-DYIKTVKALTSCRRE--QPVPS--------RPTK     2675
Eublepharis_macularius            GDNYKEFTASLKNRKELRKAECSFWSDYVKTLKASSGCRRD--QPITS--------KASD     2730
Xenopus_tropicalis                GTNYKEFSSSLTNNQGLKNSECSFWNDYIPTLKVSTGLKSDFLPTEKT--------PESK     2759
Aquarana_Catesbeiana              GRKFKEFTKRLPNGQDLKHNECSFWNDYIPALKLSTSVKFSSVSPTKS--------SKSV     2740
Carassius_auratus                 GRGYKELSFTLSNRKNLQSPQCSFWSQYVPTLSTSTAKFSCETSVGDSGGIQNPTQVPKS     2719
Cyprinus_carpio                   GRGYKELSFTLSNRKNLQSPQCSFWSQYVPTLITSTAKFSCEISVGDSGGIQNPTQVPKS     2713
Danio_rerio                       GRAYKELSSTLVNRKNLQRSQCSFWSQYVPALTSSTAKFSCGTSAE-ETGVQTPTQEPKP     2712
Astyanax_mexicanus                GQSYKELSPSLSNRKSLQSAQCSFWSQYVPALSASTGKLLCNSAVGETGASLRPTSEPKP     2708
Clupea_harengus                   GDNYKQLSGALSNGRELRRAECSFWDDYVPALKSSTANFSQDAVVAESTGAAAPTPESKL     2742
Oryzias_latipes                   PPSYLELSPALKPQQGLSQTFCSPWGQLLARQSSEAERRG--------------------     2353
Xiphophorus_maculatus             PPTYLELSPALQHQQGLRQNACSFWRELGTRLTSKTGEFGAEPVQ-----------IPD-     2371
Stegastes_partitus                PPTYLELSPALRHQEGLSQNACSFWEQLGTRLTSSTGELGAEPVQPVL--------TAE-     2389
Cynoglossus_semilaevis            SPTYLELSPSLMNRQGLRQRACSFWNHMRTKLSRENVALKVDSFRTDH------------     2359
Lepisosteus_oculatus              GDNYKEFATSLDNRKALRKMECSFWTDYIPTLTESTRQFSSGISNEEANTLAVPTVETKL     2739
Lampetra_fluviatilis              GGLYKEVRAGMRNHRGLKMRECSFWKDYVQPLMAATGGIGEVERQWREDYHAW---RQEA     2851
Lampetra_planeri                  GGLYKEVRAGMRNHRGLKMRECSFWKDYVQPLMAATGGIGEVERQWREDYHAW---RQEA     2788
Petromyzon_marinus                GGFYKEVRAGMRNQRGLKMRECSFWKDYVQPLMAATGGIGEVERQWREDYNAW---RQEA     2801
```

$T_3$-forming acceptor and donor site

```
Homo_sapiens                          LT-----AGSGLREDLLS--LQEPGSKTYSK--  2768
Pan_troglodytes                       LT-----AGSGLREDLLS--LQEPGSKSYSK--  2766
Gorilla_gorilla                       LT-----AGSGLREDLLS--LQEPGSKSYSK--  2755
Pongo_pygmaeus                        LT-----AGSGLREDLLS--LQEPGSKSYSK--  2769
Macaca_mulatta                        LT-----AGSGLREDLLS--LQEPGSKSYSK--  2769
Macaca_fascicularis                   LT-----AGSGLREDLLS--LQEPGSKSYSK--  2769
Rattus_norvegicus                     LE-----VGPGSEEDFSG--SLEPVPKSYSK--  2768
Mus_musculus                          LE-----VGPGLEEDLSG--SLEPVPKSYSK--  2766
Cavia_porcelus                        LL-----AGSELEPNFLD--LPEPGSKSYSK--  2767
Canis_lupus_familiaris                EP-----ADSGLIGE--------PGSKSYSK--  2762
Panthera_leo                          PP----------TGE--------PGSKSYSK--  2759
Bos_taurus                            QP-----AGSGLTEDLLG--LPELASKTYSK--  2769
Trichechus_manatus_latirostris        LL-----AGSGLREDLLD--LPDPGSKTYSK--  2770
Columbia_livia                        AP-----SEGFTSQE--S--QPLGDKVAYSR--  2770
Taeniopygia_guttata                   --------E-V--TSLES--QPLGDEVAYSR--  2767
Larus_michahellis                     AP-----SEGLTSRE--R--QPLGDKVAYSR--  2775
Gallus_gallus                         AP-----SKGLSSSE--S--QPVGDKVAYSR--  2766
Struthio_Camelus                      AP-----SARLNSQETLS--QPVGDGVAYSR--  2750
Chelonia_mydas                        EP-----GAAFNSEDPQS--QSVEDKAAYSK--  2765
Alligator_mississippiensis            ES-----SVRVNSQDTPS--QSGEDKVAYSK--  2750
Crotalus_tigris                       GL-----AVTPMSKTTTV--KLLEDKAAYSK--  2756
Python_bivittatus                     EL-----AMTPISKTTAV--KLLEDKEAYSK--  2699
Eublepharis_macularius                PL-----AGESVPKSTQT--KPPEEKVAYSK--  2754
Xenopus_tropicalis                    L----------LPSLTPA--SPEQDKEGYNRRK  2780
Aquarana_Catesbeiana                  S----------PSAVTQA--SLKQGKESYN---  2758
Carassius_auratus                     IP-----DPSFS--LTPS--KPKSEKDAYN---  2740
Cyprinus_carpio                       IP-----DPSFS--LTPS--KPKSEKDAYN---  2734
Danio_rerio                           LT-----PASFT--LTPS--RQKSEKDAYN---  2733
Astyanax_mexicanus                    LP-----QTDFSSSVTQS--KPKSEKDAYN---  2731
Clupea_harengus                       I-------HAFQDLVTQN--KPKSEKDAYN---  2763
Oryzias_latipes                       ---------------------------------  2353
Xiphophorus_maculatus                 L----------PLAAPPS--QSQTNKDAYN---  2389
Stegastes_partitus                    L----------PVAAPSS--QSQTEKDGYN---  2407
Cynoglossus_semilaevis                V----------PKSRSFV----SSRTTSY----  2374
Lepisosteus_oculatus                  I-------GLFGTTVTQS--KPKSEKDTYN---  2760
Lampetra_fluviatilis                  LLEWRVQMSNFRAAVTGSTATERPAAAPYV---  2881
Lampetra_planeri                      LLEWRVQMSNFRAAVTGSTATERPAAAPYV---  2818
Petromyzon_marinus                    LLEWRVQMSNFRAAVTGSMATERPAAAPYV---  2831
```

**Supplementary Figure S10. Comparative alignment of full-length thyroglobulin sequences from 38 species, generated using the Clustal Omega program.** Amino acids are represented using single-letter codes. Signal peptides (gray), cysteine residues (yellow), and tyrosine residues (green) are highlighted. The figure displays the linker, hinge, and ChEL domains, along with eleven TG type 1 modules, three TG type 2 modules, five TG type 3 modules, and spacers 1, 2, and 3.

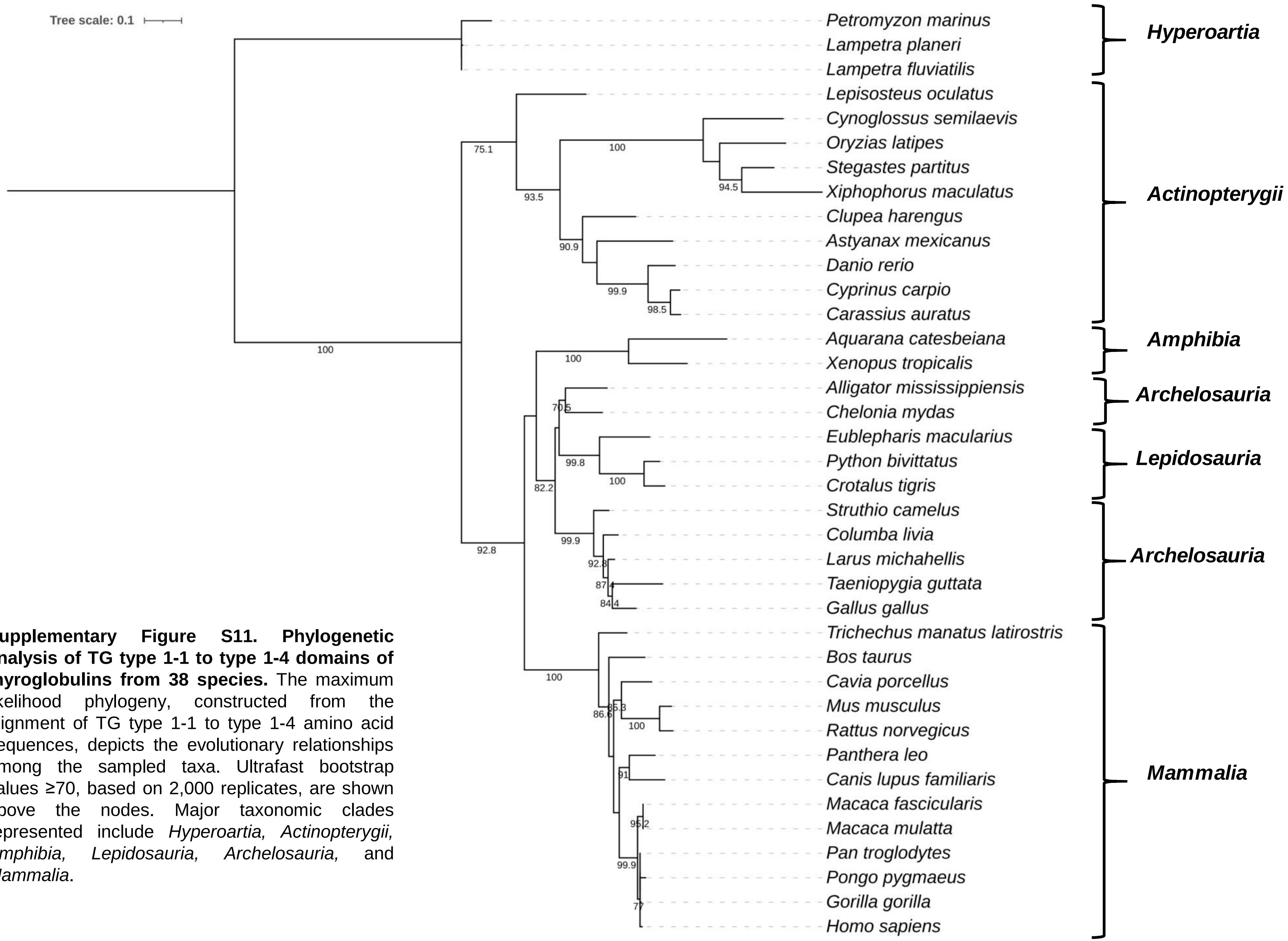


**Supplementary Figure S11. Phylogenetic analysis of TG type 1-1 to type 1-4 domains of thyroglobulins from 38 species.** The maximum likelihood phylogeny, constructed from the alignment of TG type 1-1 to type 1-4 amino acid sequences, depicts the evolutionary relationships among the sampled taxa. Ultrafast bootstrap values ≥70, based on 2,000 replicates, are shown above the nodes. Major taxonomic clades represented include *Hyperoartia, Actinopterygii, Amphibia, Lepidosauria, Archelosauria,* and *Mammalia*.

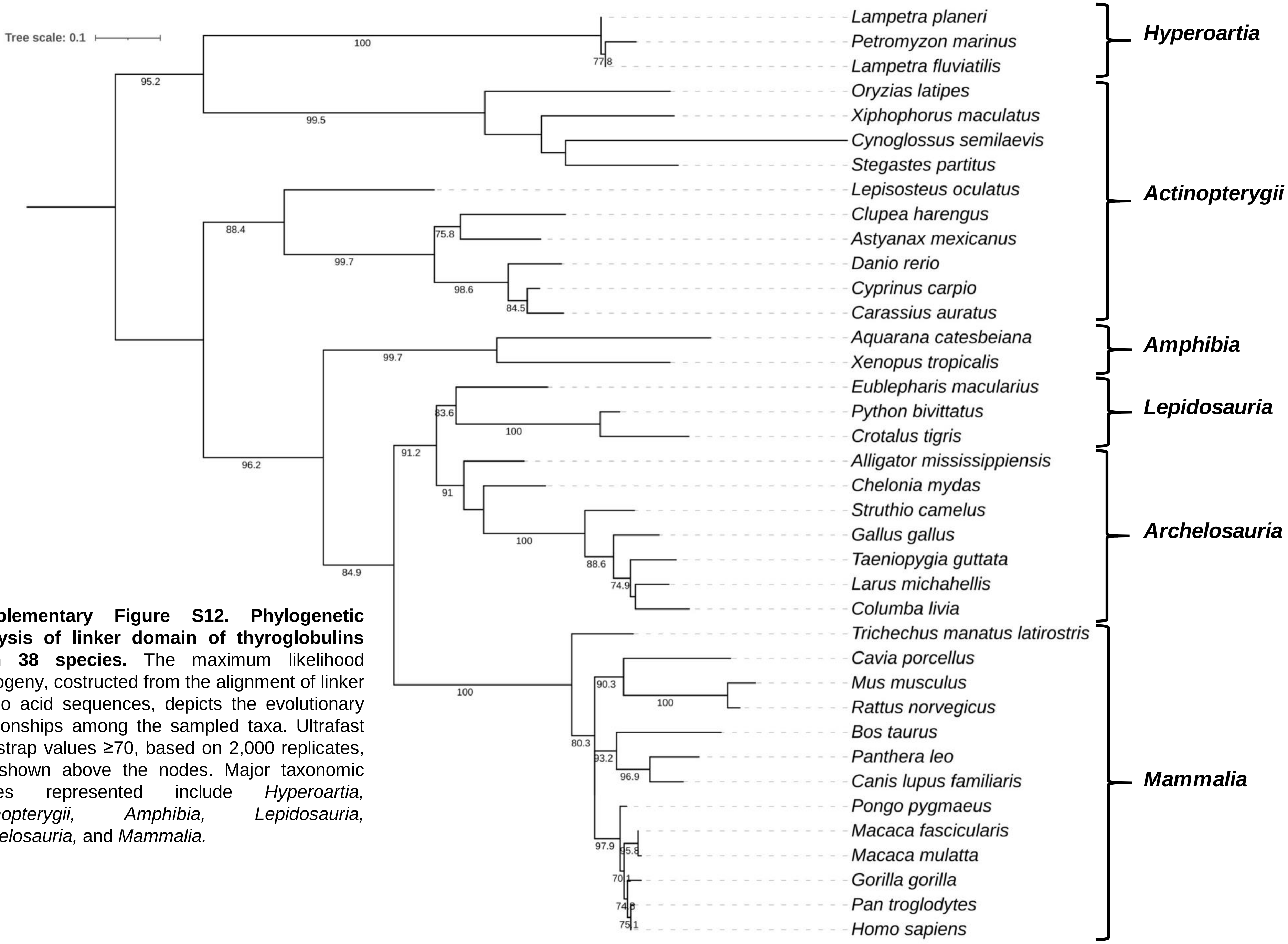


**Supplementary Figure S12. Phylogenetic analysis of linker domain of thyroglobulins from 38 species.** The maximum likelihood phylogeny, costructed from the alignment of linker amino acid sequences, depicts the evolutionary relationships among the sampled taxa. Ultrafast bootstrap values ≥70, based on 2,000 replicates, are shown above the nodes. Major taxonomic clades represented include *Hyperoartia, Actinopterygii, Amphibia, Lepidosauria, Archelosauria,* and *Mammalia.*

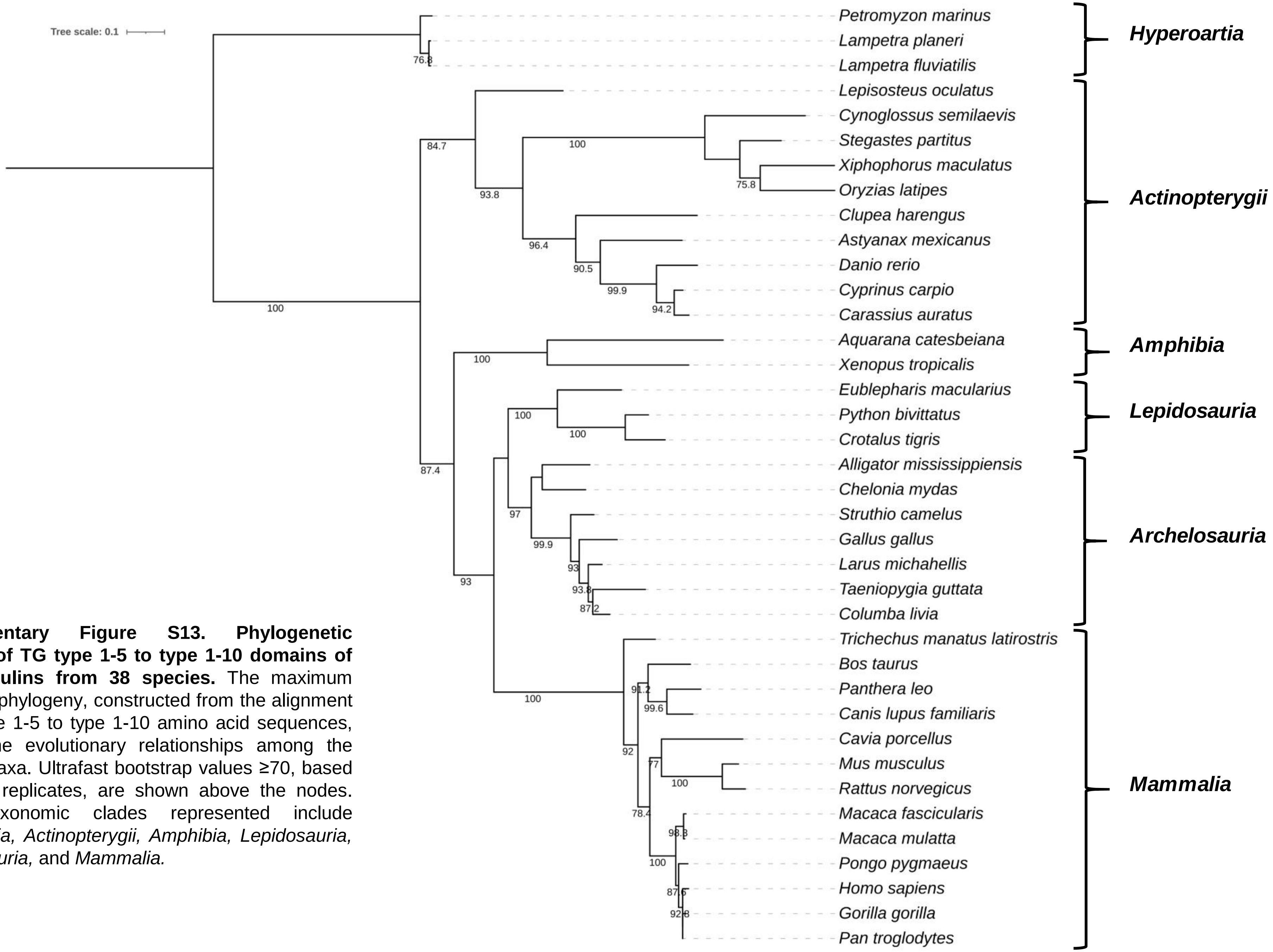


**Supplementary Figure S13. Phylogenetic analysis of TG type 1-5 to type 1-10 domains of thyroglobulins from 38 species.** The maximum likelihood phylogeny, constructed from the alignment of TG type 1-5 to type 1-10 amino acid sequences, depicts the evolutionary relationships among the sampled taxa. Ultrafast bootstrap values ≥70, based on 2,000 replicates, are shown above the nodes. Major taxonomic clades represented include *Hyperoartia, Actinopterygii, Amphibia, Lepidosauria, Archelosauria,* and *Mammalia.*

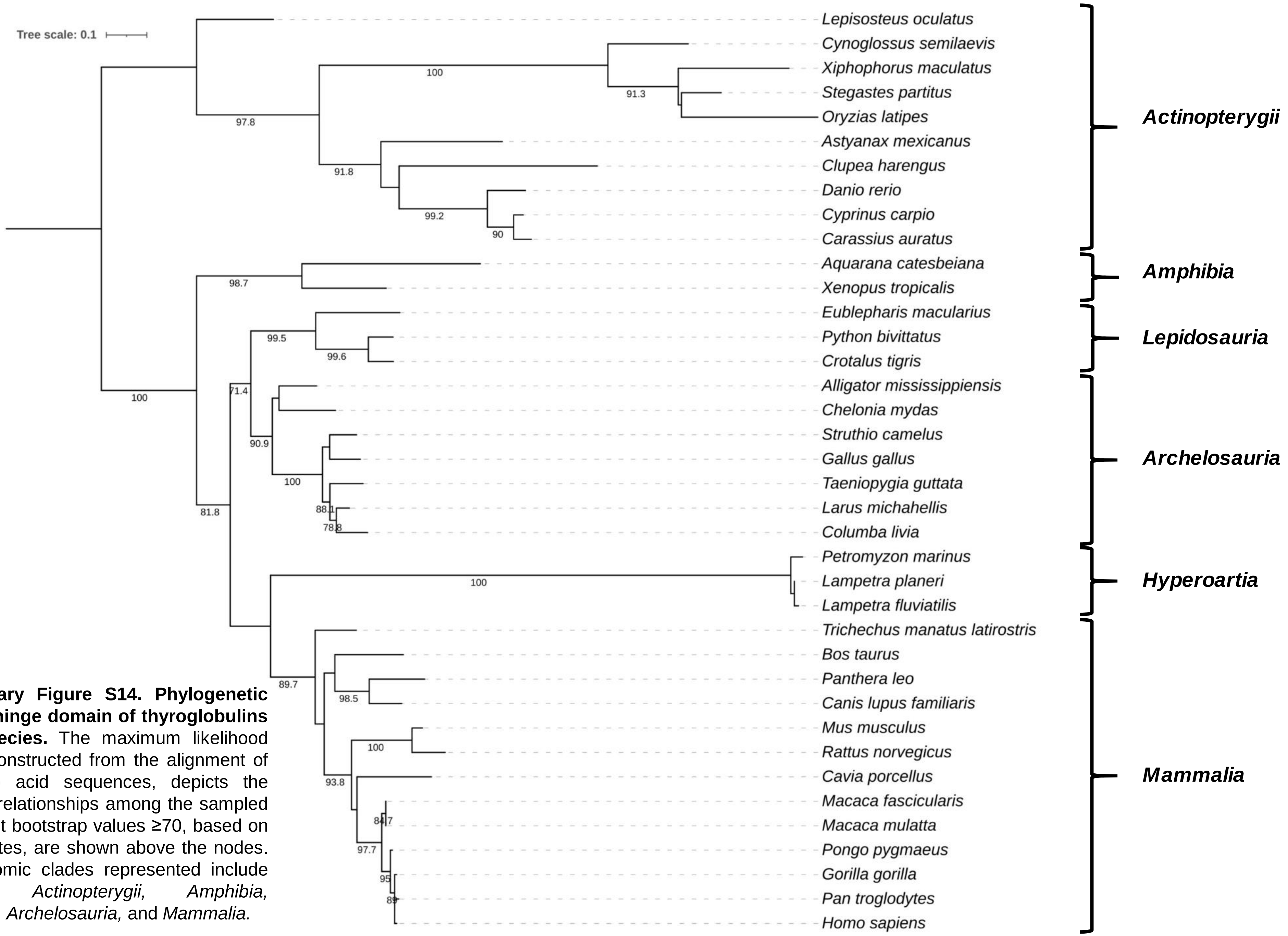


**Supplementary Figure S14. Phylogenetic analysis of hinge domain of thyroglobulins from 38 species.** The maximum likelihood phylogeny, constructed from the alignment of hinge amino acid sequences, depicts the evolutionary relationships among the sampled taxa. Ultrafast bootstrap values ≥70, based on 2,000 replicates, are shown above the nodes. Major taxonomic clades represented include *Hyperoartia, Actinopterygii, Amphibia, Lepidosauria, Archelosauria,* and *Mammalia.*

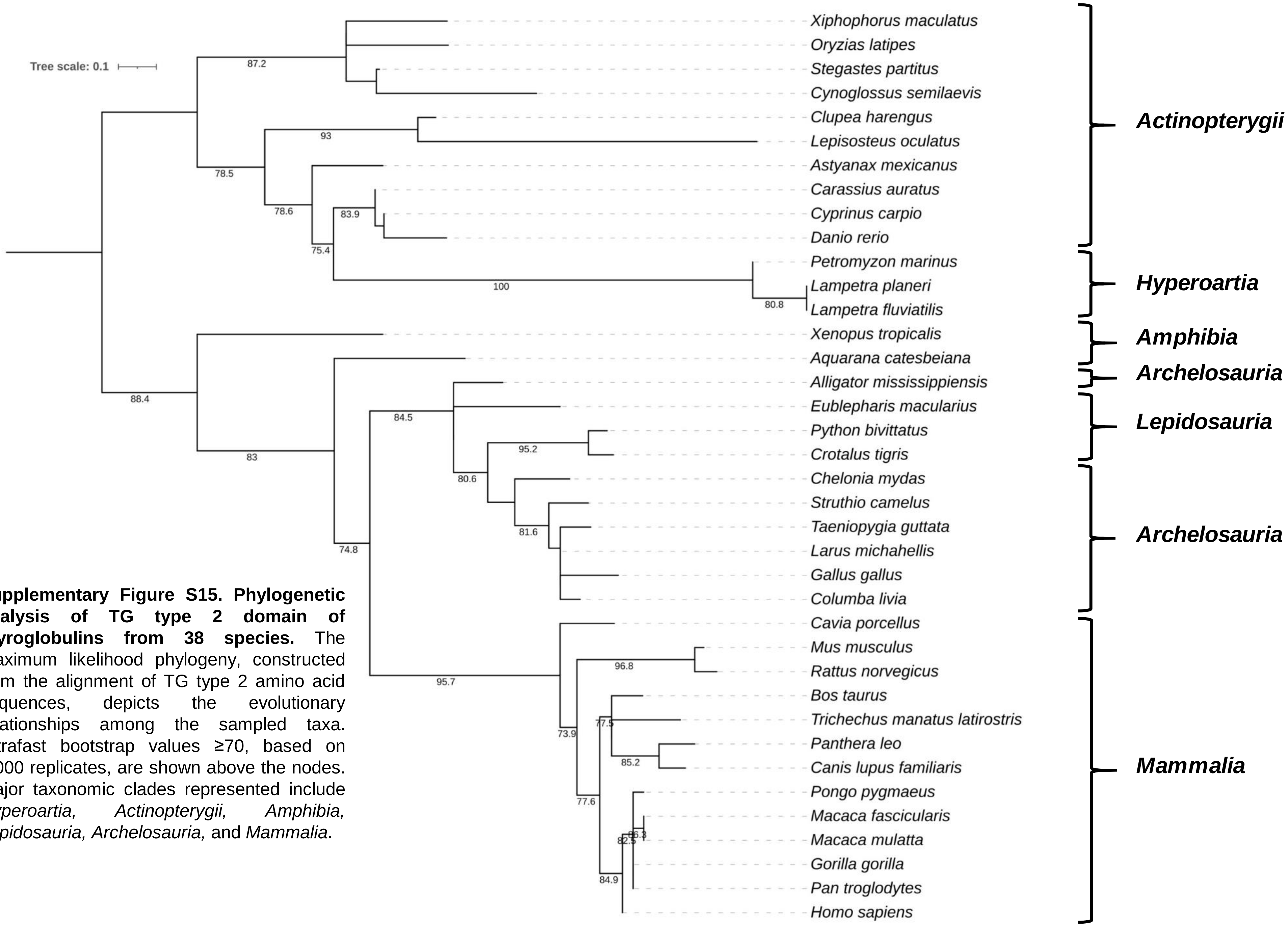


**Supplementary Figure S15. Phylogenetic analysis of TG type 2 domain of thyroglobulins from 38 species.** The maximum likelihood phylogeny, constructed from the alignment of TG type 2 amino acid sequences, depicts the evolutionary relationships among the sampled taxa. Ultrafast bootstrap values ≥70, based on 2,000 replicates, are shown above the nodes. Major taxonomic clades represented include *Hyperoartia, Actinopterygii, Amphibia, Lepidosauria, Archelosauria,* and *Mammalia*.

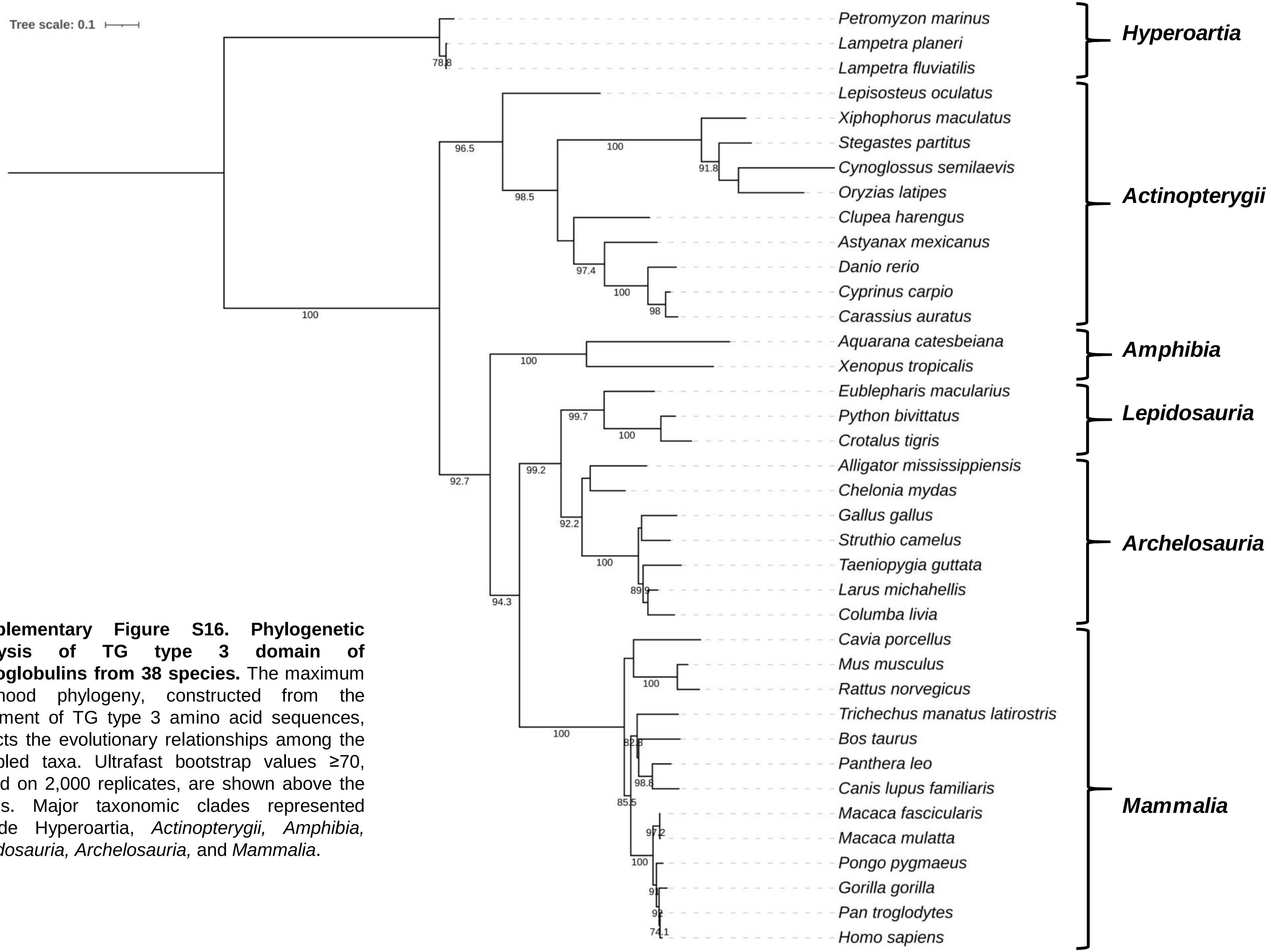


**Supplementary Figure S16. Phylogenetic analysis of TG type 3 domain of thyroglobulins from 38 species.** The maximum likelihood phylogeny, constructed from the alignment of TG type 3 amino acid sequences, depicts the evolutionary relationships among the sampled taxa. Ultrafast bootstrap values ≥70, based on 2,000 replicates, are shown above the nodes. Major taxonomic clades represented include Hyperoartia, *Actinopterygii, Amphibia, Lepidosauria, Archelosauria,* and *Mammalia*.

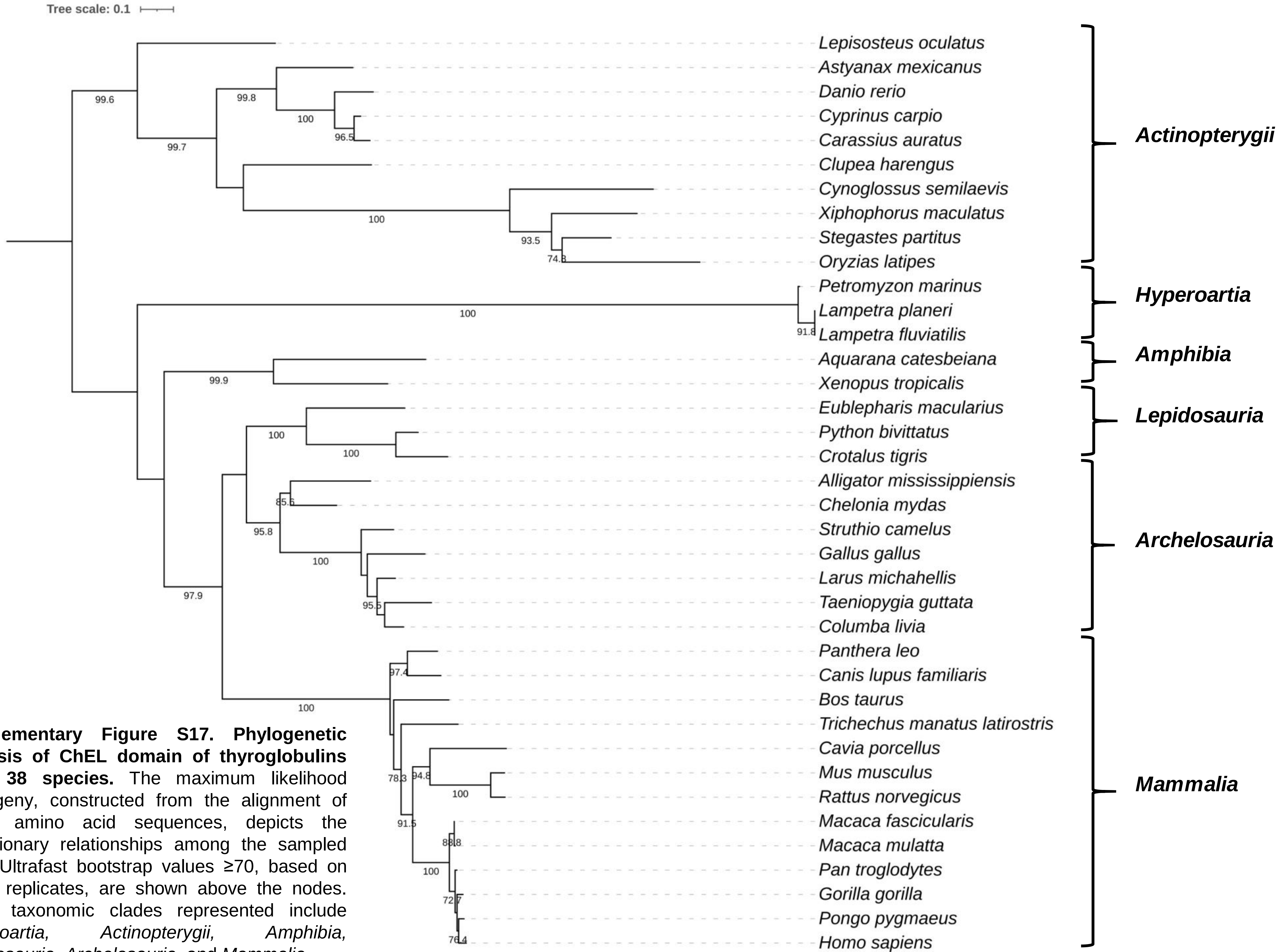


**Supplementary Figure S17. Phylogenetic analysis of ChEL domain of thyroglobulins from 38 species.** The maximum likelihood phylogeny, constructed from the alignment of ChEL amino acid sequences, depicts the evolutionary relationships among the sampled taxa. Ultrafast bootstrap values ≥70, based on 2,000 replicates, are shown above the nodes. Major taxonomic clades represented include *Hyperoartia, Actinopterygii, Amphibia, Lepidosauria, Archelosauria,* and *Mammalia*.

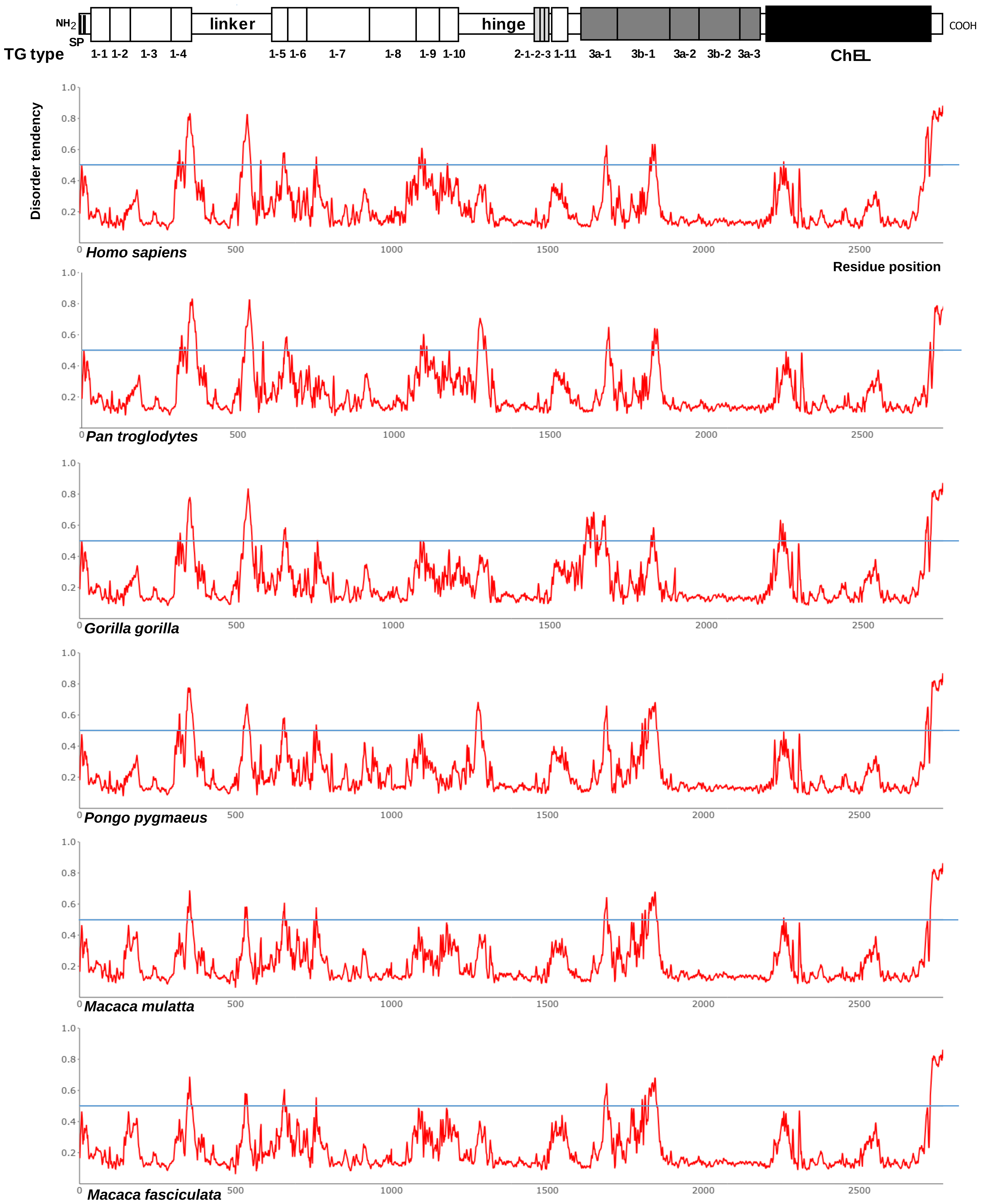


**Supplementary Figure 18. Protein disorder prediction for full-length thyroglobulin sequences from species within the *Mammalia* clade (a).** Disorder analysis was performed using the AIUPred web server with the "AIUPred-only disorder" setting. Residues with a predicted score ≥0.5 were classified as disordered, while those below this threshold were considered ordered. At the top, the classical model of thyroglobulin's primary structure is shown to scale, with the signal peptide (SP), TG type-1, TG type-2, and TG type-3 modules, linker and hinge domains, spacers 1, 2, and 3, and the cholinesterase-like (ChEL) homology domain represented as boxed elements.

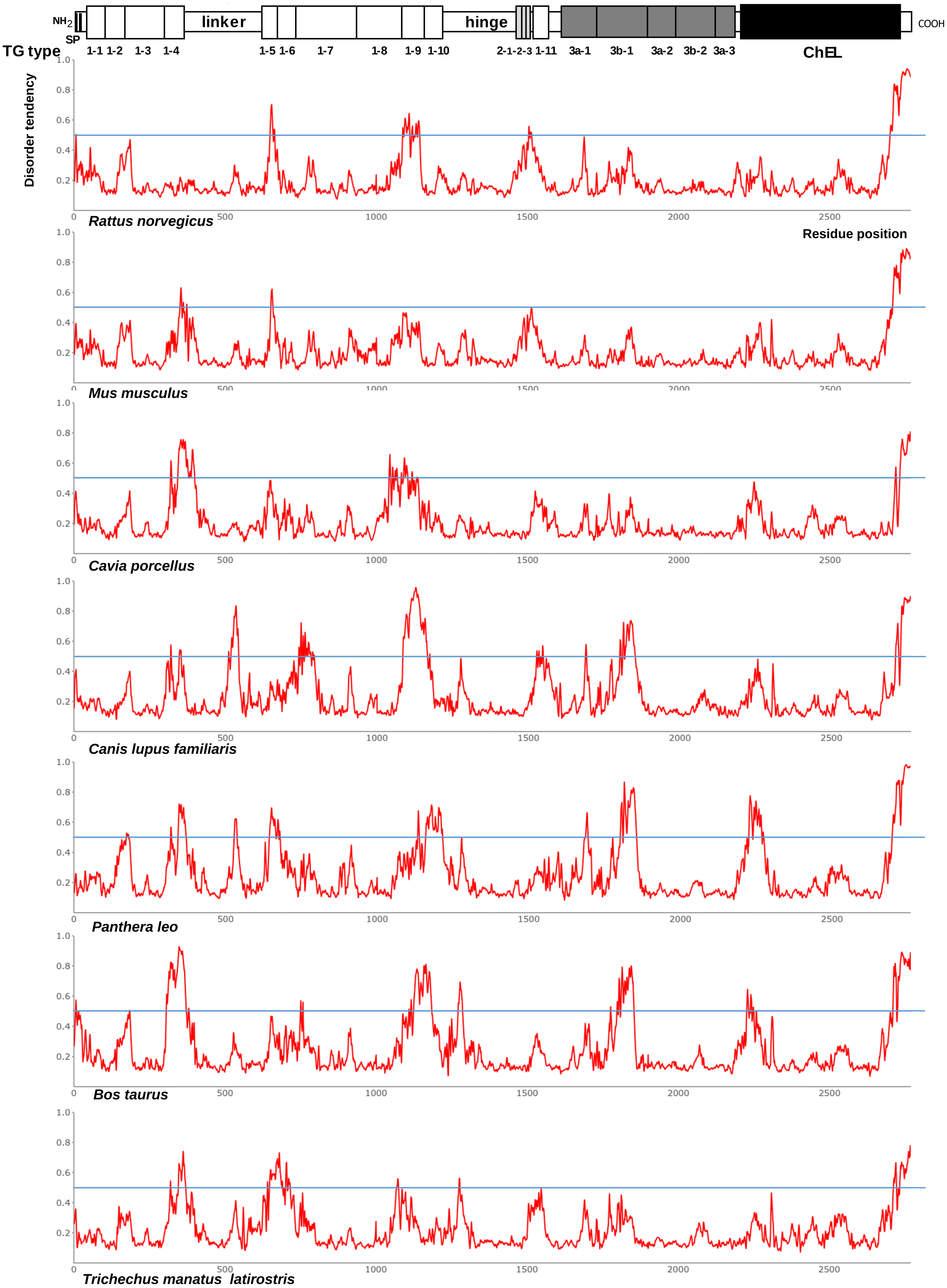
NH2
SP
linker
hinge
COOH
TG type
1-1 1-2 1-3 1-4 1-5 1-6 1-7 1-8 1-9 1-10 2-1-2-3 1-11 3a-1 3b-1 3a-2 3b-2 3a-3
ChEL
Disorder tendency
Residue position
Rattus norvegicus
Mus musculus
Cavia porcellus
Canis lupus familiaris
Panthera leo
Bos taurus
Trichechus manatus latirostris

**Supplementary Figure 19. Protein disorder prediction for full-length thyroglobulin sequences from species within the *Mammalia* clade (b).** Disorder analysis was performed using the AIUPred web server with the “AIUPred-only disorder” setting. Residues with a predicted score ≥0.5 were classified as disordered, while those below this threshold were considered ordered. At the top, the classical model of thyroglobulin’s primary structure is shown to scale, with the signal peptide (SP), TG type-1, TG type-2, and TG type-3 modules, linker and hinge domains, spacers 1, 2, and 3, and the cholinesterase-like (ChEL) homology domain represented as boxed elements.

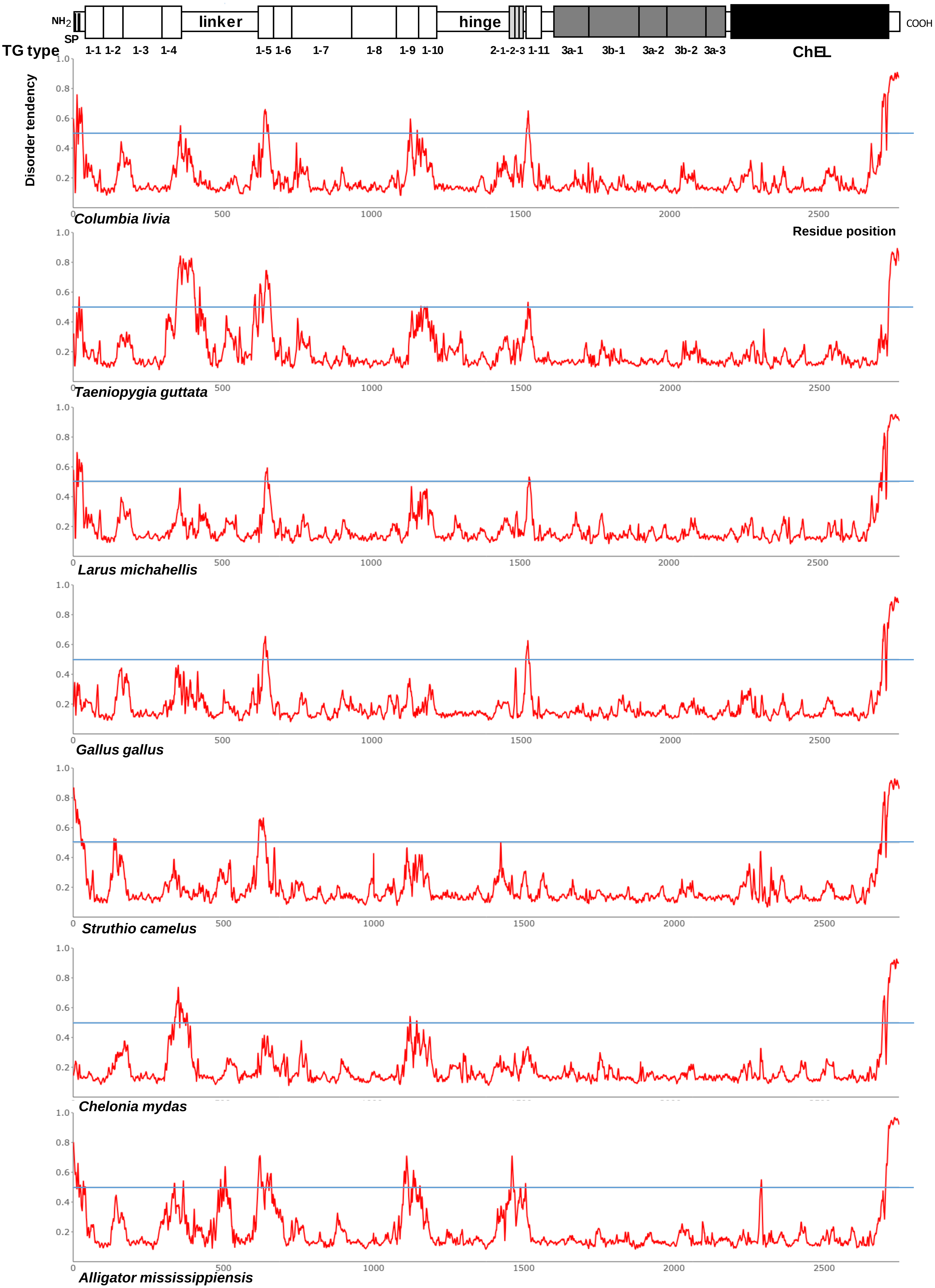
NH2
SP
TG type
1-1 1-2 1-3 1-4
linker
1-5 1-6 1-7 1-8 1-9 1-10
hinge
2-1-2-3 1-11
3a-1 3b-1 3a-2 3b-2 3a-3
ChEL
COOH
Disorder tendency
Residue position
Columbia livia
Taeniopygia guttata
Larus michahellis
Gallus gallus
Struthio camelus
Chelonia mydas
Alligator mississippiensis

**Supplementary Figure 20. Protein disorder prediction for full-length thyroglobulin sequences from species within the *Archelosauria* clade.** Disorder analysis was performed using the AIUPred web server with the “AIUPred-only disorder” setting. Residues with a predicted score ≥0.5 were classified as disordered, while those below this threshold were considered ordered. At the top, the classical model of thyroglobulin’s primary structure is shown to scale, with the signal peptide (SP), TG type-1, TG type-2, and TG type-3 modules, linker and hinge domains, spacers 1, 2, and 3, and the cholinesterase-like (ChEL) homology domain represented as boxed elements.

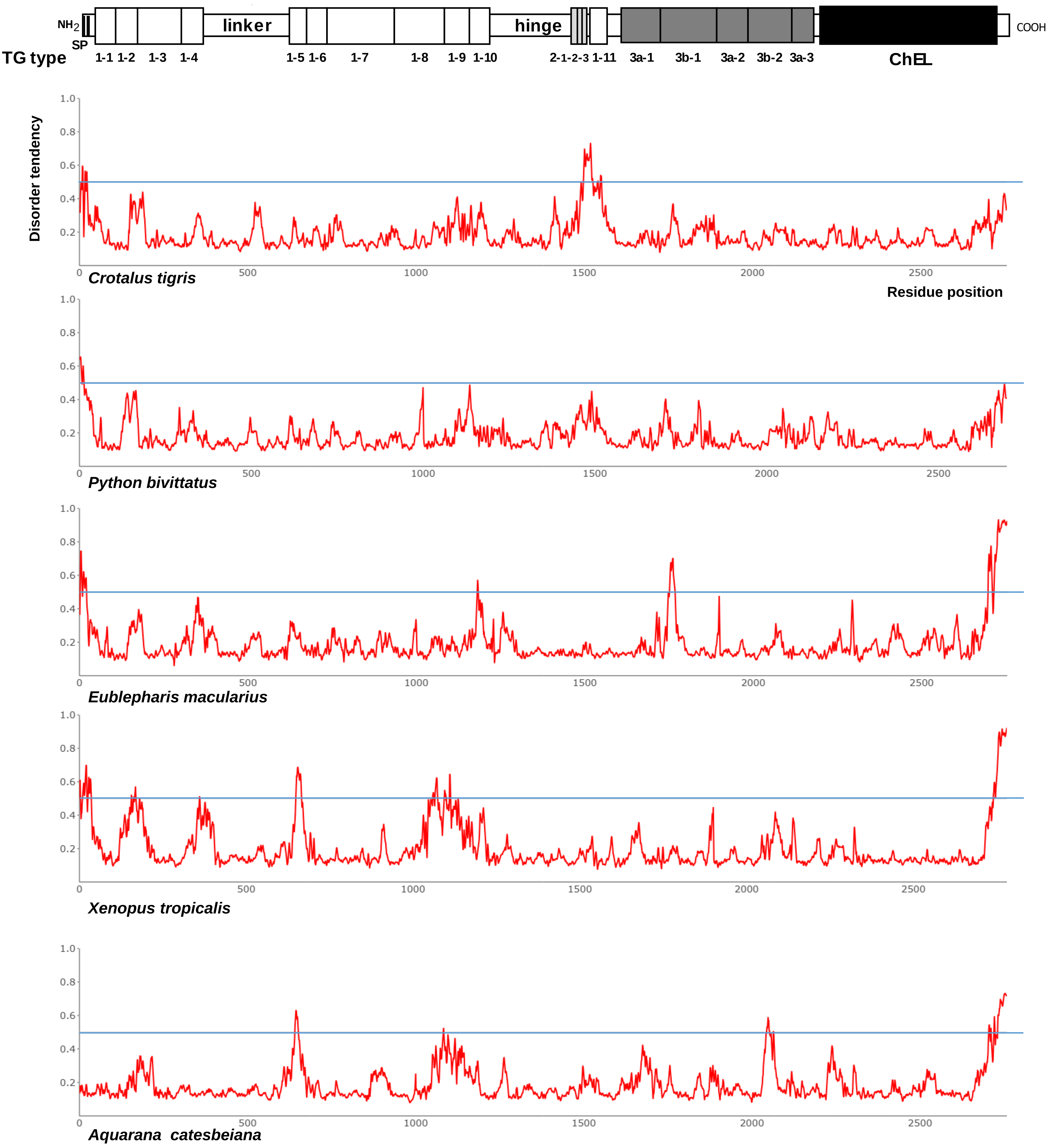


**Supplementary Figure 21. Protein disorder prediction for full-length thyroglobulin sequences from species within the *Lepidosauria* and *Amphibia* clades.** Disorder analysis was performed using the AIUPred web server with the "AIUPred-only disorder" setting. Residues with a predicted score ≥0.5 were classified as disordered, while those below this threshold were considered ordered. At the top, the classical model of thyroglobulin's primary structure is shown to scale, with the signal peptide (SP), TG type-1, TG type-2, and TG type-3 modules, linker and hinge domains, spacers 1, 2, and 3, and the cholinesterase-like (ChEL) homology domain represented as boxed elements.

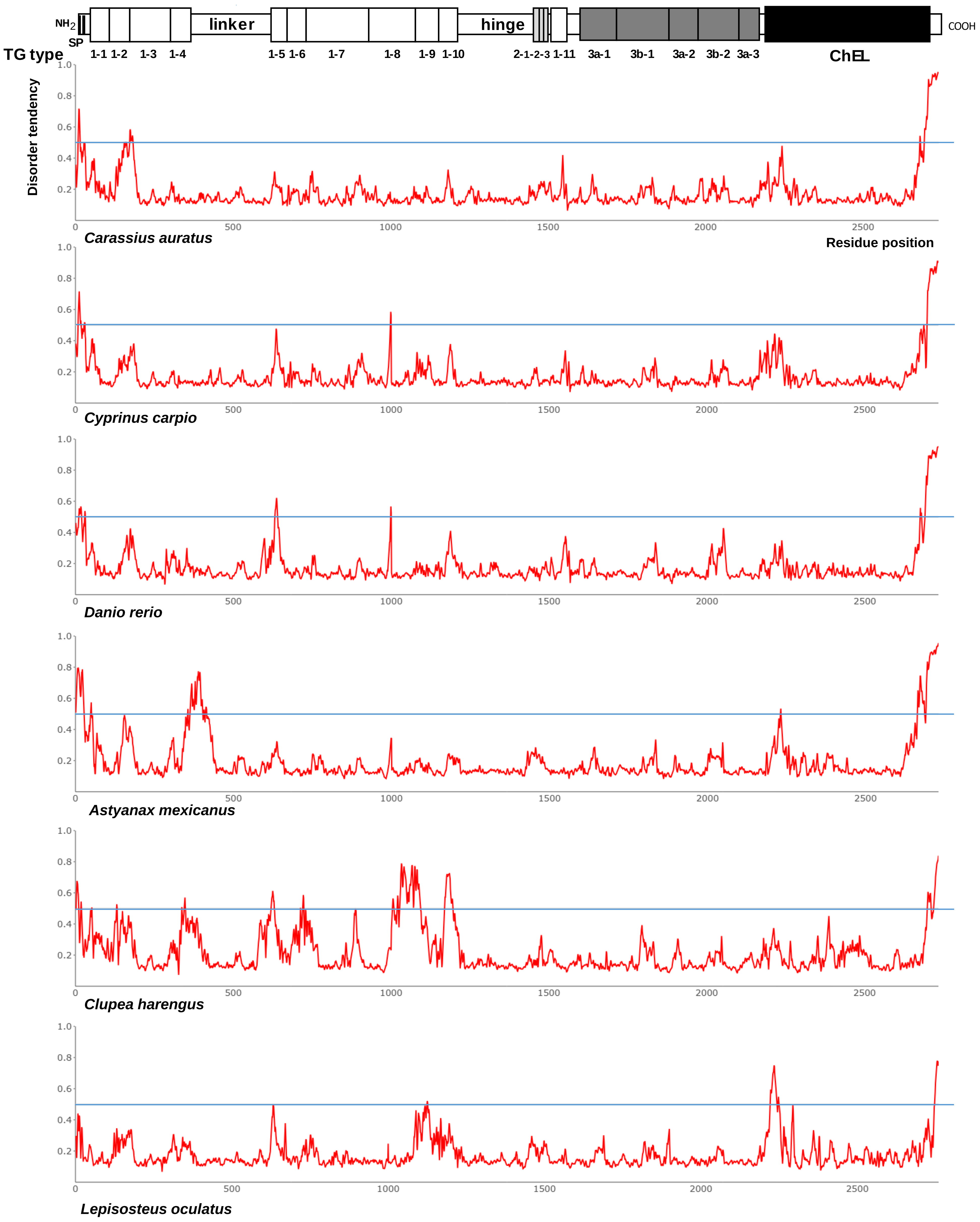


**Supplementary Figure 22. Protein disorder prediction for full-length thyroglobulin sequences from species within the *Actinopterygii* clade (a).** Disorder analysis was performed using the AIUPred web server with the "AIUPred-only disorder" setting. Residues with a predicted score ≥0.5 were classified as disordered, while those below this threshold were considered ordered. At the top, the classical model of thyroglobulin's primary structure is shown to scale, with the signal peptide (SP), TG type-1, TG type-2, and TG type-3 modules, linker and hinge domains, spacers 1, 2, and 3, and the cholinesterase-like (ChEL) homology domain represented as boxed elements.

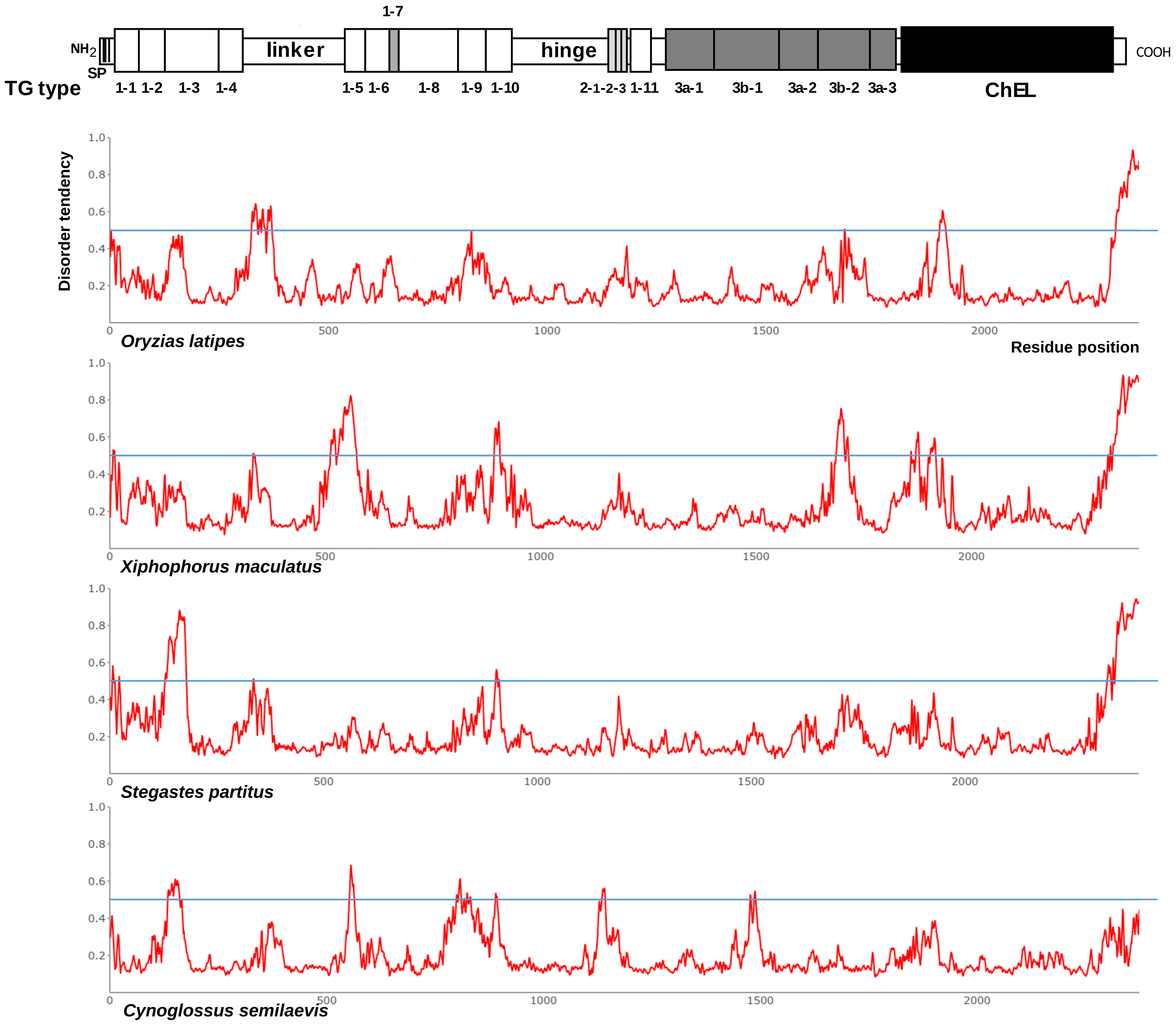


**Supplementary Figure 23. Protein disorder prediction for full-length thyroglobulin sequences from species within the *Actinopterygii* clade (b).** Disorder analysis was performed using the AIUPred web server with the “AIUPred-only disorder” setting. Residues with a predicted score ≥0.5 were classified as disordered, while those below this threshold were considered ordered. At the top, the classical model of thyroglobulin’s primary structure is shown to scale, with the signal peptide (SP), TG type-1, TG type-2, and TG type-3 modules, linker and hinge domains, spacers 1, 2, and 3, and the cholinesterase-like (ChEL) homology domain represented as boxed elements.

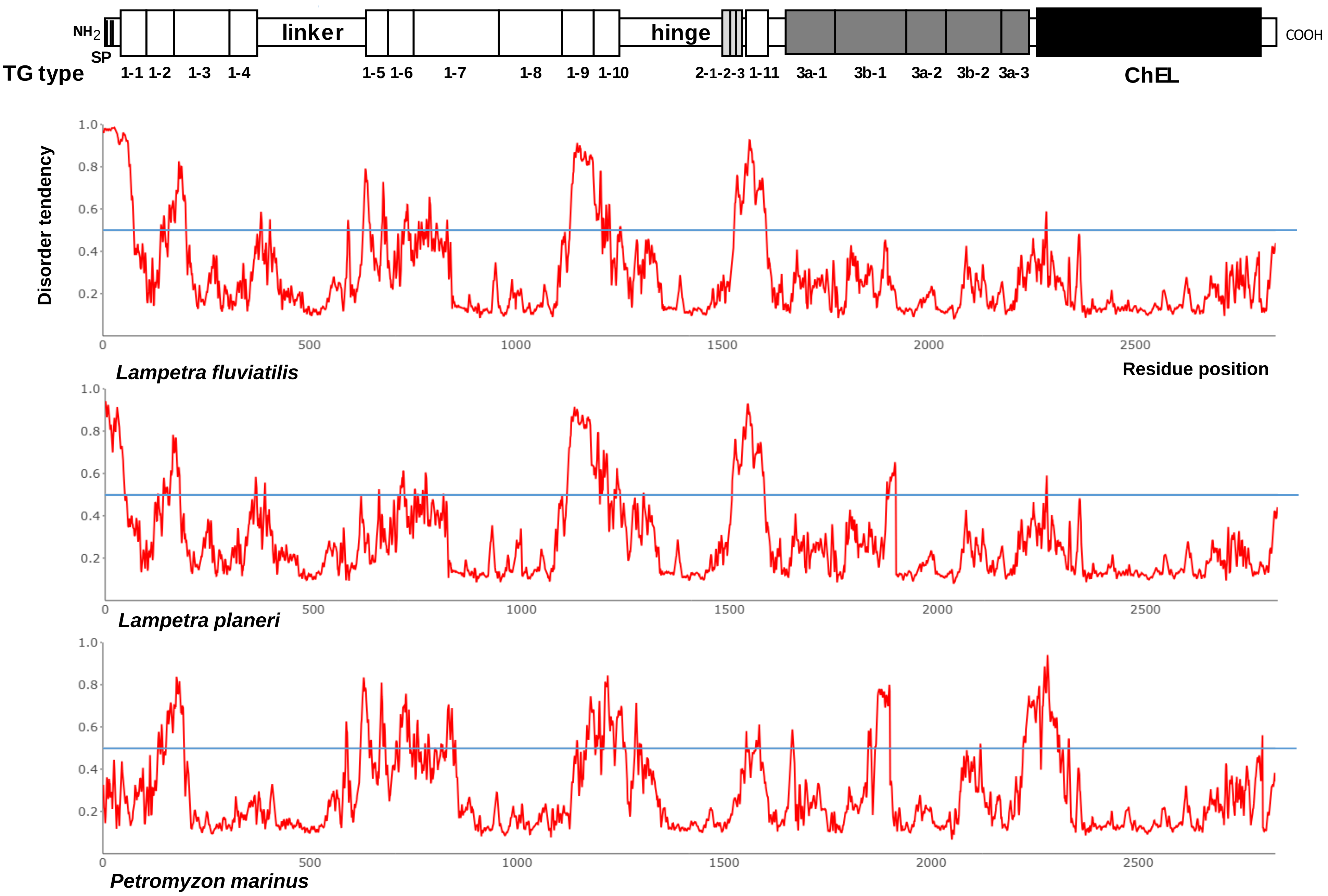


**Supplementary Figure 24. Protein disorder prediction for full-length thyroglobulin sequences from species within the *Hyperoartia* clade.** Disorder analysis was performed using the AIUPred web server with the "AIUPred-only disorder" setting. Residues with a predicted score ≥0.5 were classified as disordered, while those below this threshold were considered ordered. At the top, the classical model of thyroglobulin's primary structure is shown to scale, with the signal peptide (SP), TG type-1, TG type-2, and TG type-3 modules, linker and hinge domain, spacers 1, 2, and 3, and the cholinesterase-like (ChEL) homology domain represented as boxed elements.

.
