## Supplementary Video S1-Evolution of thyroglobulin for "Evolution of thyroglobulin: an integrated view of its origin and complexity from a structural perspective"

### Slide 1
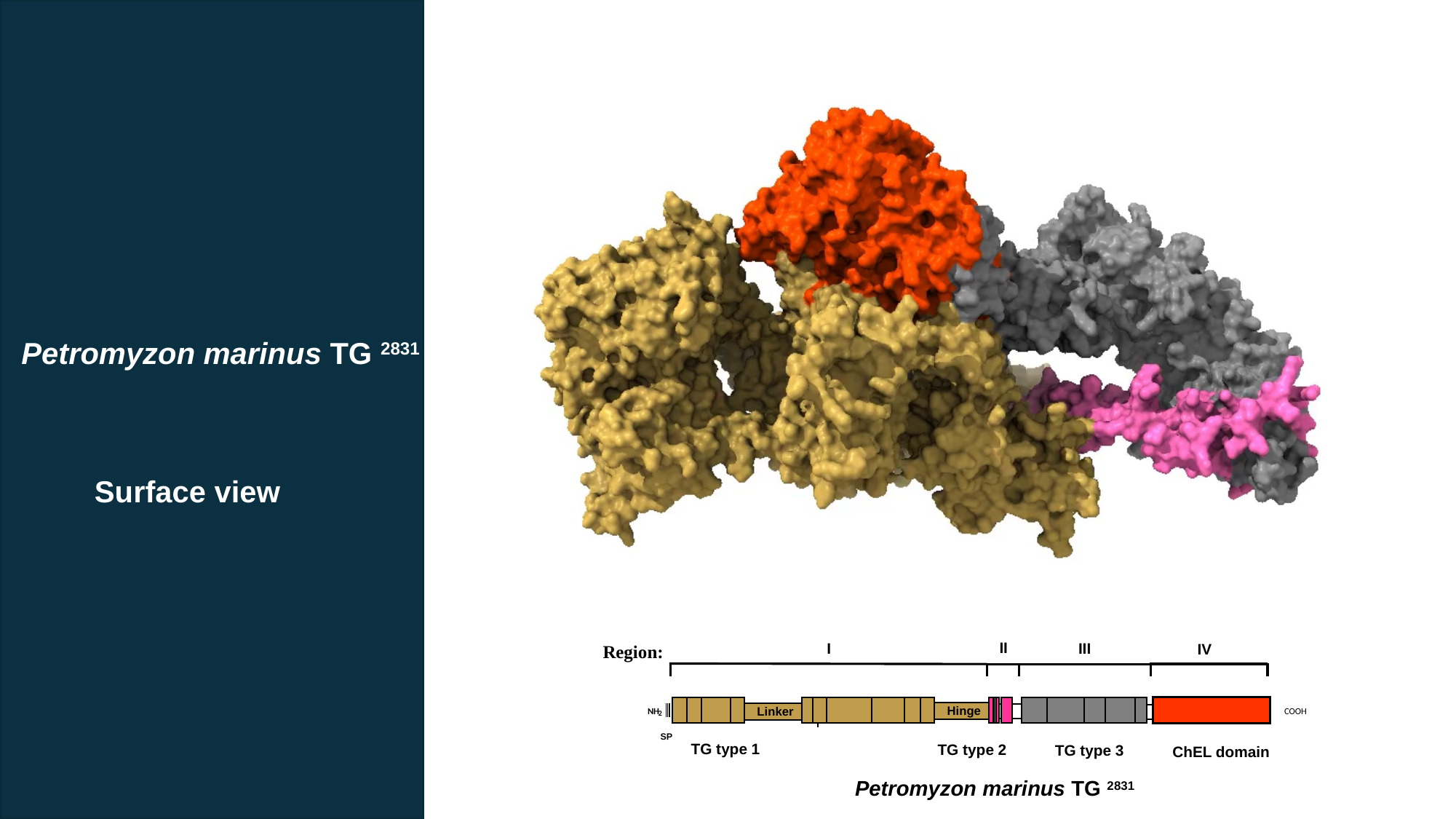

Petromyzon marinus TG ^2831^
Surface view
 II
 I
 III
IV
Region:
Hinge
Linker
NH
2
COOH
SP
TG type 1
TG type 2
TG type 3
ChEL domain
Petromyzon marinus TG ^2831^
