## Supplementary Video S2-Evolution of thyroglobulin for "Evolution of thyroglobulin: an integrated view of its origin and complexity from a structural perspective"

### Slide 1
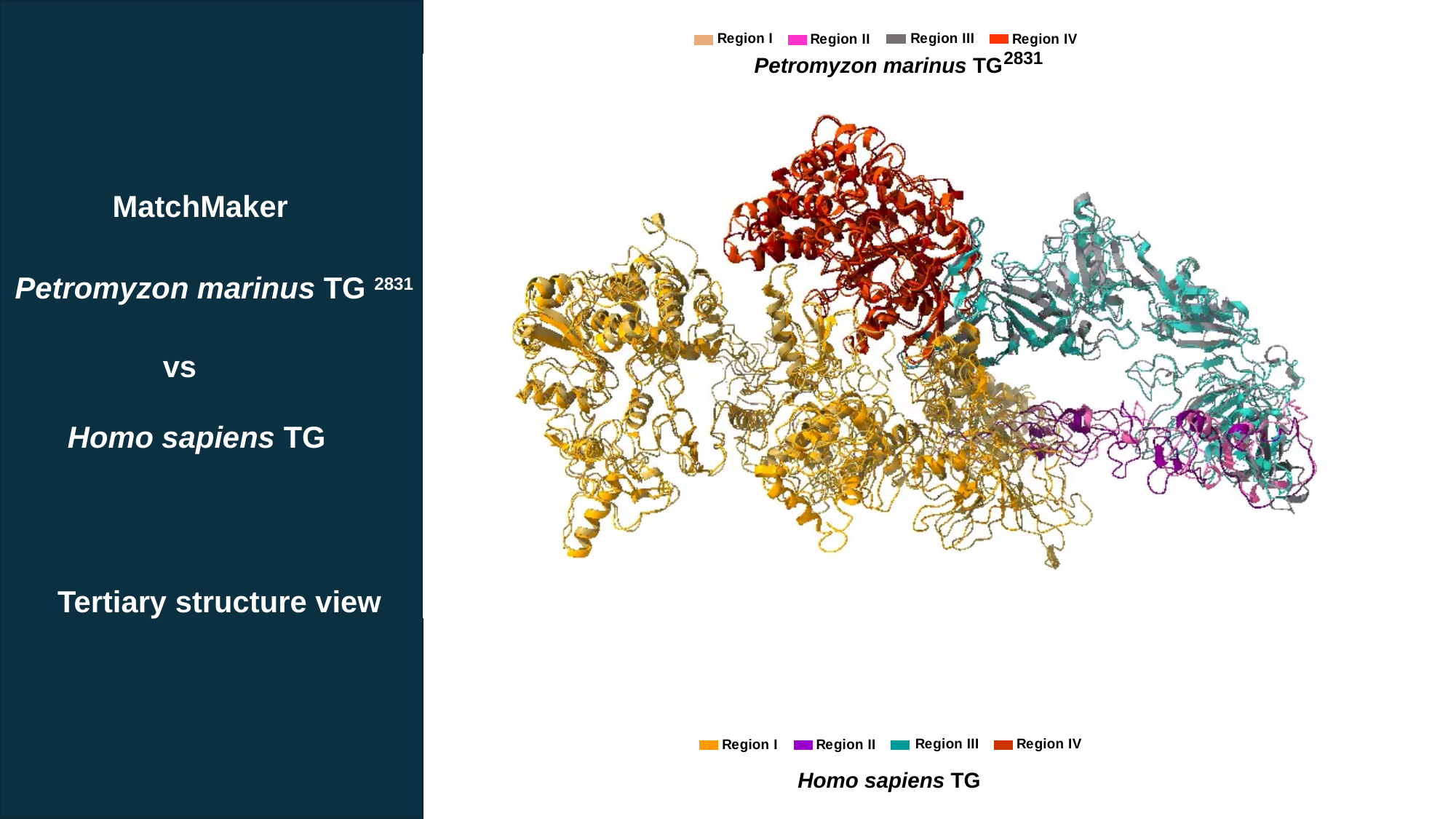

2831
Petromyzon marinus TG
MatchMaker
Petromyzon marinus TG ^2831^
vs
Homo sapiens TG
Tertiary structure view
Homo sapiens TG
