## Supplementary Video S3-Evolution of thyroglobulin for "Evolution of thyroglobulin: an integrated view of its origin and complexity from a structural perspective"

### Slide 1
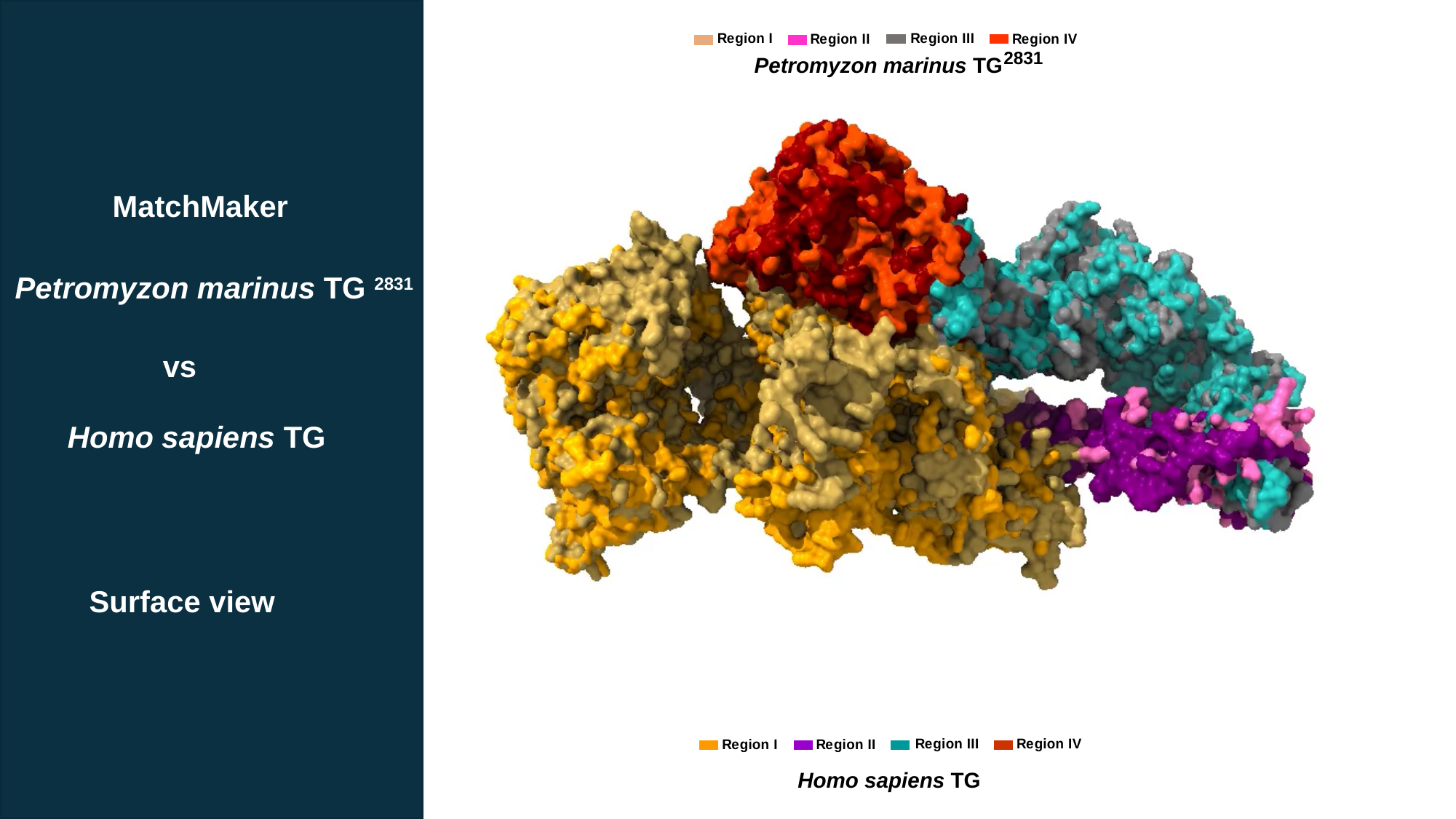

2831
Petromyzon marinus TG
MatchMaker
Petromyzon marinus TG ^2831^
vs
Homo sapiens TG
Surface view
Homo sapiens TG
